# Spatial and temporal transcriptomics reveal microglia-astroglia crosstalk in the amyloid-β plaque cell niche of Alzheimer’s disease

**DOI:** 10.1101/719930

**Authors:** Wei-Ting Chen, Ashley Lu, Katleen Craessaerts, Benjamin Pavie, Carlo Sala Frigerio, Renzo Mancuso, Xiaoyan Qian, Jana Lalakova, Malte Kühnemund, Iryna Voytyuk, Leen Wolfs, An Snellinx, Sebastian Munck, Aleksandra Jurek, Jose Fernandez Navarro, Takaomi C Saido, Joakim Lundeberg, Mark Fiers, Bart De Strooper

## Abstract

The linear cause-consequence relationship linking amyloid-β peptide (Aβ) accumulation to neuronal dysfunction in Alzheimer disease (AD) is gradually replaced by the concept that Aβ initiates complex inflammatory-like cellular alterations that progressively become Aβ independent and lead to brain dyshomeostasis. Little is known about the pathophysiology of this cellular phase of AD. We use here two orthogonal technologies, Spatial Transcriptomics and *in situ* sequencing, to analyse the transcriptome changes in cells in the amyloid-β plaque niche in a knock-in mouse model for AD. We identify a multicellular co-expressed gene network of 57 Plaque-Induced Genes (PIGs) that define a series of co-ordinated and spatially restricted microglia, astroglia and oligodendrocyte responses to progressing amyloid plaques encompassing complement, oxidative stress and inflammation. A separate oligodendrocyte network suggests abnormal myelination. Spatial Transcriptomics provides an unprecedented approach to untangle the dysregulated cellular network in the vicinity of pathogenic hallmarks of AD and other brain diseases.

## Introduction

Tremendous progress has been made over the last years to define cell states in physiological and pathological conditions using next generation sequencing approaches. For example, in the Alzheimer’ disease (AD) field, we know now that microglia respond with a stereotypical response to amyloid-β (Aβ) plaque formation (Keren-Shaul *et al.*, 2017; Krasemann *et al.*, 2017; Sala Frigerio *et al.*, 2019; Srinivasan *et al.*, 2019). Less is however known about neurons, astrocytes or oligodendrocytes. These cells are much more difficult to isolate than microglia, especially from the fragile aging brain. Isolating single nuclei from frozen brain might circumvent part of the problems (Mathys *et al.*, 2019), but it remains unclear how well all disease relevant transcriptomic changes are covered (Lake *et al.*, 2017). In addition, methods to isolate single cells or nuclei induce artificial changes in expression profiles (van den Brink *et al.*, 2017). A fundamental problem is the loss of most spatial information including the relationship to amyloid plaques. While the Aβ plaque niche has been investigated using laser capture microdissection (Rothman *et al.*, 2018; Xiong, Ge and Ma, 2019), these experiments involved pooling of the micro-dissected samples and information on original spatial localization, amyloid density, and cellular composition of the samples was lost. In contrast, the use of spatially barcoded arrays allows global unbiased transcriptome profiling maintaining the morphology of the tissue and the spatial localization of the sequenced molecules (Ståhl *et al.*, 2016; Rodriques *et al.*, 2019, Maniatis et al, 2019, Science).

The advent of these new technologies allows tackling in a novel way a central question in AD research, i.e., what is the relationship of amyloid plaques to the neurodegenerative process in brain? The question is particularly relevant given the emerging insight and worry that amyloid lowering therapies show little efficacy with regard to memory improvement (Gallardo and Holtzman, 2017; Makin, 2018). Some investigators have suggested that the plaques are inert and that soluble oligomeric Aβ drives neurotoxicity (Glabe, 2008; Ono, Condron and Teplow, 2009; Mucke and Selkoe, 2012). However, Aβ lowering drugs should affect amyloid plaques as well as oligomers and the failed amyloid therapy trials therefore suggests that the role of amyloid peptide in AD needs to be reconsidered. We proposed that amyloid plaques act as a trigger, not as a driver of the disease (Karran, Mercken and De Strooper, 2011) and that accumulation of Aβ in the brain initiates a complex multicellular neurodegenerative process that might become autonomous, i.e. independent of the presence of plaques (Strooper and Karran, 2016). GWAS studies show in fact that much of the risk of sporadic AD is associated with genes expressed in micro- and astroglia (Jansen *et al.*, 2019). These genes are strongly up- or downregulated when exposed to amyloid deposition (Salih *et al.*, 2018; Sala Frigerio *et al.*, 2019; Sierksma *et al.*, 2019). It is clear that we need more in depth study of the protective and harmful cellular reactions that occur in the course of AD. The study of this “cellular phase of AD” (Strooper and Karran, 2016) should lead to a comprehensive understanding of the complex interactions over time between neurons, glia cells and blood vessel cells, that determine the pathogenic outcome triggered by abnormal Aβ generation and deposition.

We used here “Spatial Transcriptomics” (ST, Figure 1A) (Ståhl et al., 2016), to measure *in situ*, in hundreds of small spots, close to and remote from amyloid plaques, generating transcriptome wide analysis in different brain areas. The volume of each small tissue domain (8×10^−5^ mm^3^) covers dozens of cells and is thus ideal to understand the putative dysfunctional intercellular network induced by amyloid-β plaques. We use the human APP knock-in *APP*^*NL-G-F*^ mouse model to understand the transcriptomic alterations through disease progression (Saito *et al.*, 2014). We obtained spatially resolved data and integrated all the information into one comprehensive analysis. We provide here a first analysis of the data, characterizing the molecular alterations in function of plaque load. We identified several gene co-expression networks, one involving oligodendrocytes and myelin genes and one identifying 57 amyloid <u>P</u>laque <u>I</u>nduced <u>G</u>enes (PIGs), which co-express only in the amyloid-plaque niches but not in control tissue domains. To investigate the cellular signature of the 57 PIGs, we performed *in situ* sequencing and *in situ* hybridization to reveal the spatial localization of PIGs with single cell and plaque resolution. Together, the 57 PIGs define a dysfunctional network reflecting intercellular crosstalk between plaque-associated microglia and astrocytes. The complement cascade appears to be one of the most dysregulated networks in this amyloid-β plaque cellular niche.

**Figure 1.**
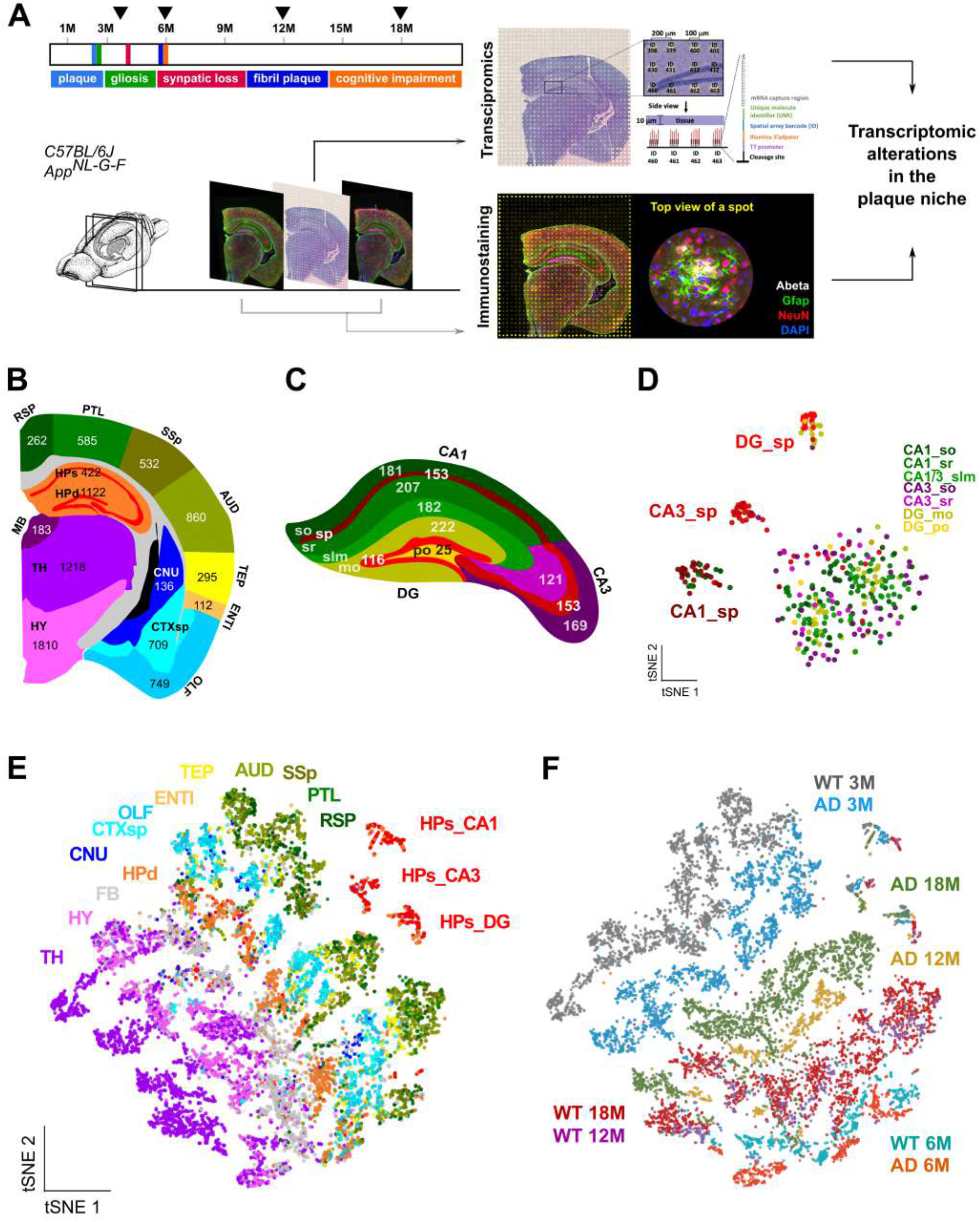
Spatially resolved transcriptomic profiles in adult brain. (A) Sequential 10 μm coronal sections (bregma −2.0 to −2.2) of *App*^*NL-G-F*^ and WT brain were collected at 3−, 6−, 12−, and 18-months of age. The middle section was used for spatial RNAseq and the two adjacent sections for immunostaining. (B-C) Total number of tissue domains per brain region (B) and in hippocampus (C) based on Allen Brain Atlas. (D) t-distributed stochastic neighbour embedding (t-SNE) plot visualizing 355 transcriptomic profiles in WT hippocampus at 3-months of age. Tissue domains cluster according to somatic layers CA1, CA3, and DG. (E and F) t-SNE plots visualizing the 10,327 transcriptomic profiles. Tissue domains were coloured according to brain region (E) or to genotype and age (F). Abbreviations: somatic layers (sp), stratum oriens (so), stratum radiatum (sr), stratum lacunosum-moleculare (slm), molecular layer (mo), polymorph layer (po), thalamus (TH), hypothalamus (HY), fiber tract (FB), dendritic layers of hippocampus (HPd), cerebral nuclei (CNU), cortical subplate (CTXsp), olfactory areas (OLF), entorhinal area (ENTI), temporal association area, ectorhinal area, and perirhinal area (TEP), auditory areas (AUD), primary somatosensory area (SSp), posterior parietal association areas (PLT), retrosplenial area (RSP).

## Results

We obtained three adjacent coronal sections per sample (Figure 1A) by cryo-sectioning frozen mouse brains at bregma stage −2.0 to −2.2 from *App*^*NL-G-F*^ and WT control *C57Black6J* mice at 3, 6, 12, and 18 months of age (Table S1 and Fig S1A). The two outer sections were used for immunostaining and amyloid plaque detection, while the middle section was processed for Spatial Transcriptomics (ST, see the Methods) (Ståhl et al., 2016).

We detected on average 31,283 ± 7441 unique molecular identifiers and 6,578 ± 987 unique genes per tissue domain defined by the circular spots of capture probes on the cover slip (mean ± SD, Figure S1). Every library contains information on >500 individual annotated tissue domains adding up to 10,327 tissue domains over 20 libraries. We subdivided the coronal sections into 15 anatomical brain regions (Figure 1B) according to the Allen Mouse Brain Atlas (Lein et al., 2007) and subsequently assigned each tissue domain to one of the brain regions (see the Methods). The number of tissue domains per brain region varies between 112 (entorhinal cortex) and 1810 (hypothalamus) (Figure 1B). We finally aligned the adjacent (stained) slides with the middle (ST) section to annotate each tissue domain with plaque load (as evaluated by mAb 6E10 staining), reactive astrocytes (GFAP), neuron quantity (NeuN) and nuclei quantity (DAPI) (see the Methods) (Figure 1A, 2A and S2).

**Figure 2.**
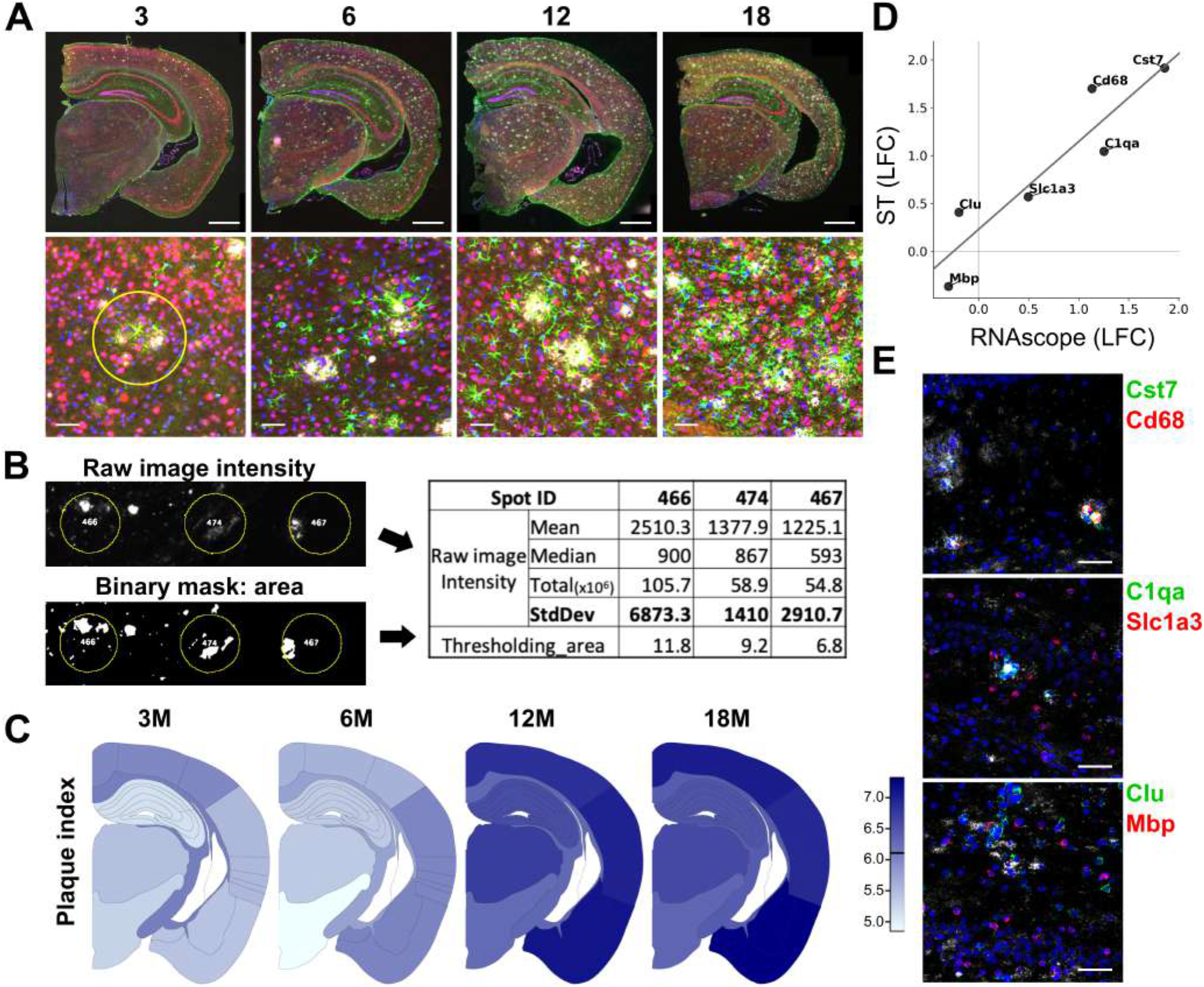
Linking gene expression alterations to amyloid plaque load. (A) Immunofluorescent staining of amyloid plaques (mAb 6E10, white), astrocytes (Gfap, green), neurons (NeuN, red), and nuclei (DAPI, blue) at the indicated ages. Scale bar: 500 μm. Bottom: zoom-ins of tissue domains (yellow circle: 100 μm in diameter) in the primary somatosensory area of the cortex. Scale bar: 25 μm. (B) Computerized pictures of amyloid plaques quantified with Image-J. Yellow circles indicate the associated tissue domains. From the five computed parameters, standard deviation of pixel intensities was found to have the highest correlation with independent expert analysis. This is used as amyloid plaque index of a tissue domain. (C) The mean of the amyloid plaque index per brain region is plotted at four time points. (D) Pearson correlation of ST and RNAscope analysis (as shown in panel E) of 6 selected transcripts at 18-month of age. Scatter plot shows the LFC of each target as detected via ST (y-axis) or RNAscope (x-axis). (E) RNAscope experiment shows the expression of the indicated genes (red or green). Plaques are immunostained with mAb 6E10 (white). Nuclei were visualized by DAPI (blue). Scale bar: 50 μm.

### Spatially resolved transcriptomic profiles in adult mouse brain

We wondered whether the 10,327 transcriptome profiles could be separated in a biological meaningful way using t-Distributed Stochastic Neighbor Embedding (t-SNE). Reassuringly, the tissue domains clustered according to mouse brain structure with remarkable specificity (Figure 1E). To see if ST can distinguish finer regions in the hippocampus, we applied a further division of the hippocampal tissue domains into 10 anatomical brain regions (Figure 1C) (Lein et al., 2007) and re-applied the t-SNE analysis to tissue domains from this region, limiting the analysis to WT hippocampus at 3-months of age to exclude genotype and age effects (Figure 1D). The 82 tissue domains covering the somatic layers of the hippocampus segregated clearly into CA1, CA3, and DG-subregions. Intriguingly, the 198 tissue domains of the different hippocampal dendritic layers intermingle with each other, suggesting that the basic gene expression in these neuropil layers are similar. We evaluated the expression of landmark genes in the hippocampus, and the results (Figure S1F and S1G) demonstrate that ST allows the differentiation of tissue domains to precise anatomic regions in the brain.

When considering tissue domains of brain slices from mice of different age or genotype, again good separation was observed (Figure 1F). The spots of wild-type mice of 12 and 18 months of age overlap (purple and red in Figure 1F), suggesting that the wild-type brain transcriptome does not change much anymore between these two time points. The *App*^*NL-G-F*^ transcriptional profile in contrast still changes between 12 and 18 months, in concordance with the progression of pathology over this period.

### Linking gene expression alterations to amyloid plaque load

Amyloid deposition in *App*^*NL-G-F*^ mice starts around 3 months, as shown by immunostaining with anti-Aβ_1-16_ antibody (6E10, Figure 2A). These plaques are largely ‘diffuse’. Thioflavin-S starts to appear in the amyloid core of 6E10-positive plaques from 6 months on (Figure S2D). On average, 1,565 ±167 plaques with an estimated diameter >10 μm (derived from a measured surface >78.5 μm^2^) are detected per coronal section of *App*^*NL-G-F*^ mouse at 18-months of age. Of these plaques, 98% have a diameter <50 μm (<1963 μm^2^ surface), which aligns well with previous measurements by others (Kuo *et al.*, 2001). The diameter of a tissue domain is 100 μm and the thickness of a single section is 10 μm (Figure 1A). Therefore, when we detect amyloid plaques in one of the two outer adjacent sections, it is reasonable to assume that adjacent cells in the central section were exposed to this amyloid plaque as well. We used the standard deviation of fluorescence intensity of pixels in a tissue domain, averaged between the two adjacent slides, as the plaque index for each tissue domain (see the Methods). The plaque index was used to quantify the amyloid propagation per region in the *App*^*NL-G-F*^ mice (Figure 2C), illustrating its stereotypically progression from the dorsal cortex towards ventral cortex, thalamus, and hippocampus (Figures 2A and 2C). Astrogliosis (Gfap staining) is selectively increased in the stratum lacunosum moleculare (CA_slm) of the hippocampus during normal aging, while it spreads all over the brain but especially towards the cortical regions during diseased aging (Figures S2).

To understand changes in gene expression we performed two differential expression analyses. The first model compares *App*^*NL-G-F*^ to WT (disease model), the second one models the effect of plaque load on gene expression (plaque model, see the Methods). We validated the model using single molecule fluorescence *in situ* hybridization (smFISH, RNAscope, Figure 2D and 2E) of six genes that accordingly to the plaque model were significantly dysregulated in *App*^*NL-G-F*^ at 18-months of age over a range of changes (*Cst7* (Log Fold-Changes (LFC): 1.91), *Cd68* (LFC: 1.70), *C1qa* (LFC: 1.04), *Slc1a3* (LFC: 0.57), *Clu* (LFC: 0.41), and *Mbp* (LFC: −0.37)). We measured the mean intensity of the hybridization per cell in the plaque niches (cells within 10 μm of the 6E10-positive areas) compared to that in tissue far from plaques (operationally defined as three expansions of 65 pixels or >64.6 μm). As shown in Figure 2D, the ST data highly correlates with the RNAscope data (Pearson correlation = 0.92, p-value = 0.009, see also Methods).

We compared both models by plotting LFC by genotype (the *disease* axis) against LFC by amyloid plaque exposure (the *plaque* axis). We observe major gene expression alterations according to genotype, amyloid plaque deposition and age (Figures 3A and 3B). Much can be learned from individual inspection of these genes. For example, *Sez6* (seizure-related gene 6), a physiological BACE1 substrate (Pigoni et al., 2016; Zhu et al., 2018) required for the appropriate development of neuronal dendrites and excitatory postsynaptic responses (Gunnersen et al., 2007), is significantly down-regulated in the disease-axis (LFC: −3, False Discovery Rates (FDR) ⋍ 0) but not in the plaque-axis (LFC: 0.20, FDR = 0.415) at 3-month of age. This suggests that this early change is driven by the mutated APP but not by visible early plaque deposition. The cellular signatures of the top 50 plaque reactive genes sorted by FDR at 3-month or at 18-month of age are summarized in Figures S3A and S3B. In the early stage (3-month of age), several oligodendrocyte-expressing genes are up-regulated (eg. *Plp1*, *Mobp*, *Mal*, *Cnp*, *Mbp*, *Mag*, *Apod*), while interneuron-expressing genes (eg. *Bex2*, *Scg2*, *Nap1l5*, *Resp18*, *Gprasp2*) are down-regulated in the plaque-axis (Figure S3A). In addition, *Il1a* (Interleukin 1 Alpha) with the ability to drive expression of proinflammatory tumor necrosis factor alpha (TNFa), interleukin-6 (Il-6), and complement components (Gerritsma *et al.*, 1996; Basu, Krady and Levison, 2004; Berge, Johnson and Berge, 2009), is significantly upregulated early in the plaque-axis. In contrast, many of the genes changed at 18 months are upregulated in function of combined disease- and plaque-axis, including those belonging to the well-known inflammatory signature of AD (Figure S3B). At this late stage, an oligodendroglial marker, *Plp1*, is significantly down-regulated in the plaque niches (LFC: −0.70, FDR: 2.03e^−14^, Figure S3B) but not in the disease-axis.

**Figure 3.**
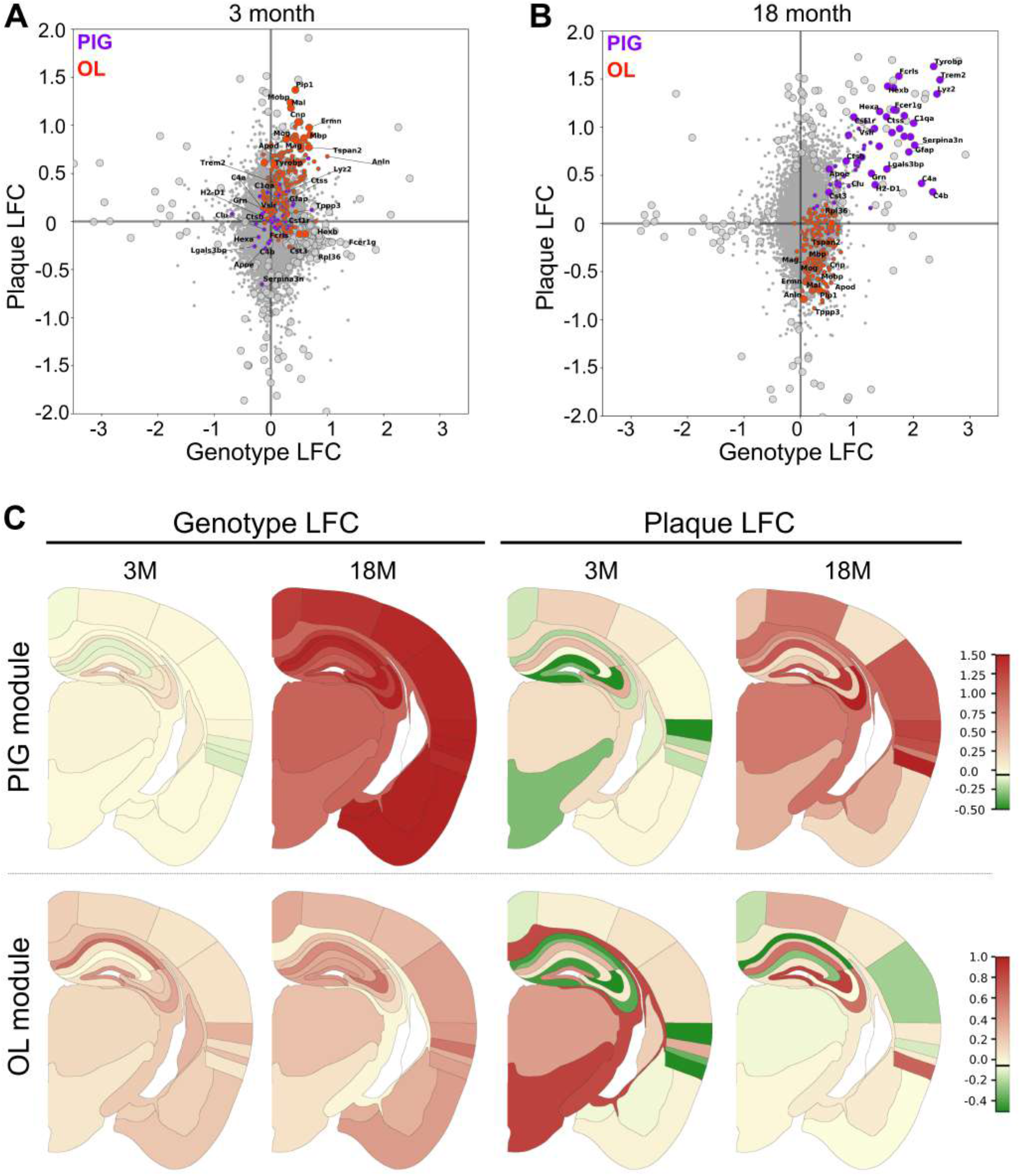
Multicellular gene co-expressing networks around amyloid plaques. (A and B) Scatter plots showing the plaque effect (y-axis) and the genotype effect (x-axis) on differential gene expression at 3− (A) and 18− (B) months of age. The y-axis (plaque-axis) represents the LFC of gene expression in the tissue domains in function of amyloid plaque index. The x-axis represents the LFC of gene expression in AD versus WT at 3 and 18 months of age. The 100 most significantly (FDR) differentially expressed genes are represented by enlarged spots. Individual genes of the OLigodendrocyte (OL module) (red spots) and the Plaque-Induced Gene (PIG) module (purple spots) are indicated. Notice strong upregulation of OL genes at 3 months and downregulation at 18 months while PIGs are relatively unchanged at 3 months but become very dominant at 18 months. (C) Regional plots of average differential expression changes along plaque- or genotype-axis for the PIG and the OL module over time. Each region is coloured according to the mean of the observed LFCs (see color bar) of the module across different brain regions.

We employed GORilla (Ashburner *et al.*, 2000; Eden *et al.*, 2009; Carbon *et al.*, 2019), and identified 13 functional super categories (Figure S4A, Table S2, see the Methods), including antigen processing, chemotaxis, lysosomal degradation and inflammation that are altered along both the plaque- and the disease-axis at 18 months. Confirming the observations discussed above, we find a clear switch in the sign of a myelin category, which is up at 3 months and down at 18 months along the plaque-axis (Figure S4A) but not in the disease-axis.

### Multicellular gene co-expressing networks around the amyloid plaques

Genes with similar expression patterns (co-expression) are likely to have similar functions, and can be grouped into modules by “Weighted Gene Co-expression Network Analysis (WGCNA)” (Zhang and Horvath, 2005). We investigated the 50% most variable genes across the 4 available time-points, over both genotypes and over all detectable brain regions and grouped highly co-fluctuating genes associated with amyloid plaques versus non-plaque environments. In total, we identified 12 modules (Table S3), and extracted the hypothetical biological functions (Table S3), the relation to amyloid plaque exposure, the disease window (Figure S4B), the cellular composition (Figure S4C), and the affected brain regions (see the Methods).

We focus here on the WGCNA modules that were most responsive to the amyloid plaques, which are the “red” and “purple” modules. The red module (Figures 3 and S4B) represents mature myelinating oligodendrocytes *(Mbp*, *Mobp*, *Mog*, and *Plp1)*, but also oligodendrocyte precursor cells (*Cnp*, *Sox10*, *Gpr37*, *Olig1*, and *Olig2)*. We refer to this module as “the oligodendrocyte (OL) module”. Agreeing with the initial ontology analysis above, this module goes up in early and down in late phase along the plaque-axis, while it has a trend to go up along the disease-axis. The result shows that oligodendroglial genes are initially upregulated especially in the vicinity of amyloid plaques, but become eventually depleted in the plaque niche while they remain relatively high far from plaques in the diseased brain. Interestingly, when considering the plaque-axis, we see a strong effect in the hippocampal-entorhinal neuronal circuit, where the OL module is already strongly down-regulated at 3 months of disease (Figure 3C lower panel, last two pictures). This might reflect specific cellular vulnerabilities in different brain regions.

The “purple” module, to which we will refer as the <u>P</u>laque-<u>I</u>nduced <u>G</u>enes (PIG) module, contains 57 genes. This module is the most reactive module up in both the plaque- and disease-axis at 18-months of age, while displaying minor dysregulation at 3-months of age (Figures 3, and S4B). The module represents a rather homogeneous, brain-wide response, showing little regional variation in contrast to the oligodendrocyte module (Figure 3C). The module starts to be up-regulated from 6-month of age in *App*^*NL-G-F*^ mice (Figure 4C) when the plaques become Thioflavin-S positive (Figure S2D) and increases exponentially between 6 and 12 months to stabilize broadly over the brain thereafter (Figure 3C). We identify a moderate but significant correlation between the amyloid accumulation and the PIG module expression among all tissue domains in AD mice at 18-month of age (Pearson correlation = 0.39, p-value ⋍ 0) (Figure 4D).

**Figure 4.**
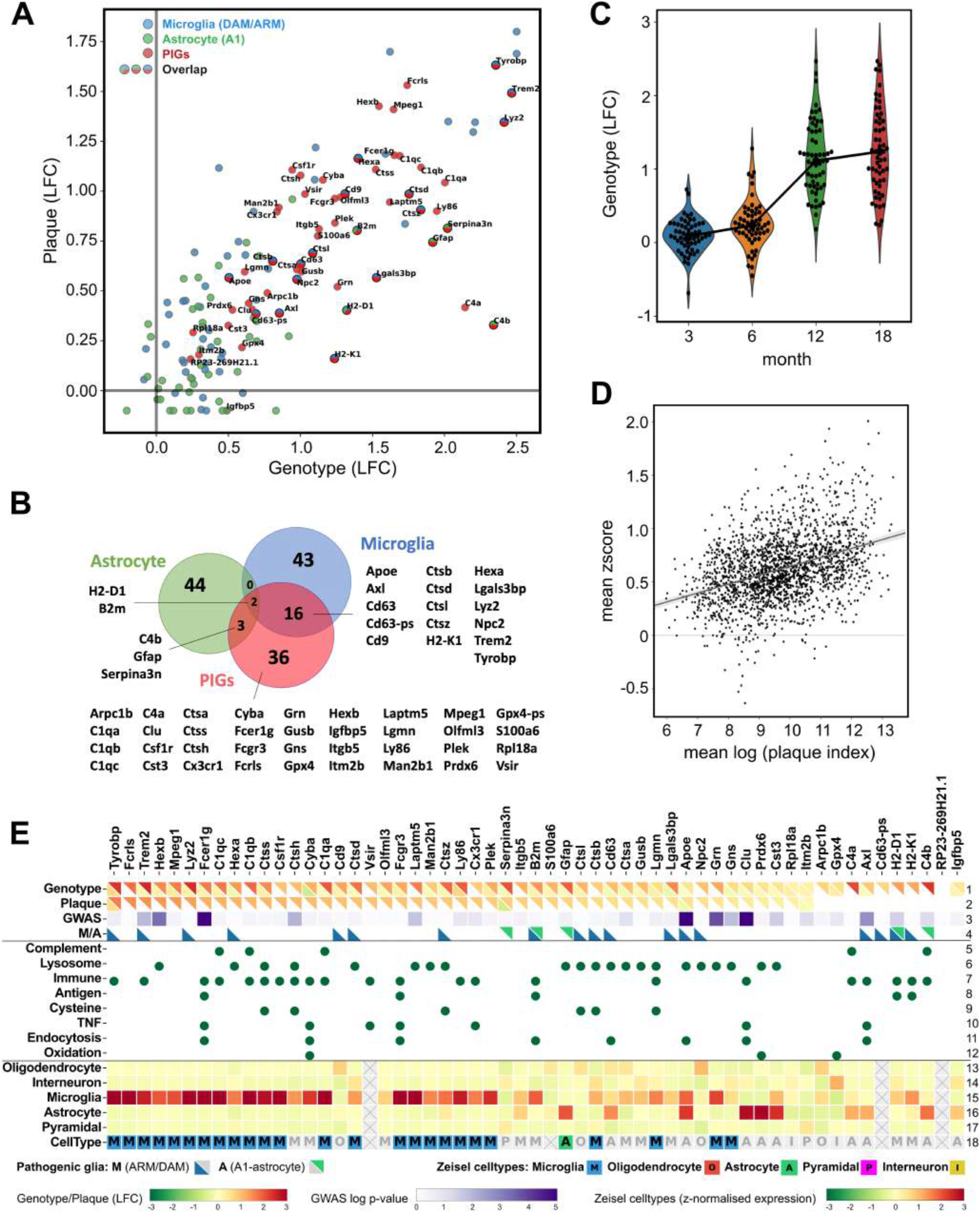
Characterization of the 57 Plaque-Induced Genes (PIGs). (A) Differential expression of 57 PIGs (red spots), 61 ARM/DAM microglia markers (blue spots) and 50 A1-astroglial markers (green spots) in the plaque-disease interaction plot at 18-month of age. (B) Venn diagram highlights the overlap of the PIGs with ARM/DAM microglia and A1-astrocytes, and 36 PIGs have not been associated with either phenotype before. The 61 activated microglial markers were the overlap between top 120 ARM genes and top 120 DAM genes sorted by FDR. (C) LFCs of 57 PIGs in the genotype model at the indicated age. (D) For every tissue domain the mean z-score of 57 PIGs (y-axis) is positively correlated with the mean of the log-transformed plaque index (x-axis). (E) PIGs are ranked from left to right according to their LFC in the plaque-axis at 18-month of age. Heatmap squares (top 2 rows) show LFCs of the genotype and the plaque effect at 3- (left bottom triangle) and 18- (right top triangle) months of age. The third row indicates the −log10 transformed GWAS AD risk p-values (see the Methods). ARM/DAM or A1 are highlighted in triangles (4th row). Genes in the significantly enriched GO terms are highlighted in green spots (row 5-12). Expression level of each gene in the indicated cell type calculated from single cell RNAseq data (Zeisel *et al.*, 2015) (row-13-17). Zeisel et al. defined (color background) / predicted cell type (grey background) genes are highlighted in the last row.

We investigated the cellular signatures of the PIG module (Zeisel *et al.*, 2015). The top plaque-reactive PIGs were mainly expressed by microglia, while astroglia are clearly also part of the module response (Figure 4E, rows 16-17). Both microglia and astrocytes undergo pronounced transformation after brain injury and disease (Zamanian *et al.*, 2012; Keren-Shaul *et al.*, 2017; Sala Frigerio *et al.*, 2019). We therefore further checked the 57 PIGs against the profiles (Figures 4A and 4B) of activated astrocytes and microglia. We selected the top 50 changes in LPS-induced inflammatory astrocytes (A1) (Zamanian *et al.*, 2012), and the top 61 changed genes in the overlap of the ARM and DAM genes in microglia (Sala Frigerio *et al.*, 2019; Keren-Shaul *et al.*, 2017). The 57 PIGs are among the strongly reactive (up-regulated) genes along either the plaque- or the disease-axis compared to most of the ARM/DAM- and A1-genes (Figure 4A). Five PIGs overlap with A1 markers and 18 PIGs with ARM/DAM genes (Figure 4B). Most interestingly there are 36 PIGs not known as disease-associated glia genes (Figure 4B). Based on GO term analysis (Figure 4E), we conclude that the PIGs module is involved in the activation of the classical complement cascade, endocytosis, lysosomal degradation, antigen processing and presentation, immune response and oxidation-reduction processes. One dominant group are known initiators of the classical complement cascade (e.g. *C1qa*, *C1qb*, *C1qc*, *C4a*).

### The PIGs module reflects micro- and astroglial crosstalk in the plaque niche

To evaluate directly with an independent approach which cell types contribute to the PIGs response in the plaque niches, we applied an *in situ* sequencing (ISS) method (Ke *et al.*, 2013) (see the Methods, Figures 5 and S5). We used customized probes to map the gene expression of PIGs together with cell type markers (*Itgam* for microglia, *Slc1a3* for astrocytes, *Syp* for neurons, and *Plp1* for oligodendrocytes). We generated two ISS libraries, one for each genotype, at 18-months of age (Figure 5). We quantified gene expression as number of fluorescent spots per gene per cell. We applied standard watershed segmentation to estimate cell borders and expanded the boundary by 10 μm (Figure 5D). After removal of low expressing cells (< 5 molecules per cell), we investigated the gene expression from 127,694 cells over the two coronal sections, detecting on average 13 ± 0.6 unique molecules per cell. We grouped the genes in 5 concentric rings around plaques. The first ring (ring1, the plaque niche) extends 10 μm around each plaque and consecutive rings expand with 18.2 μm (65 pixels) (Figure 5E).

**Figure 5.**
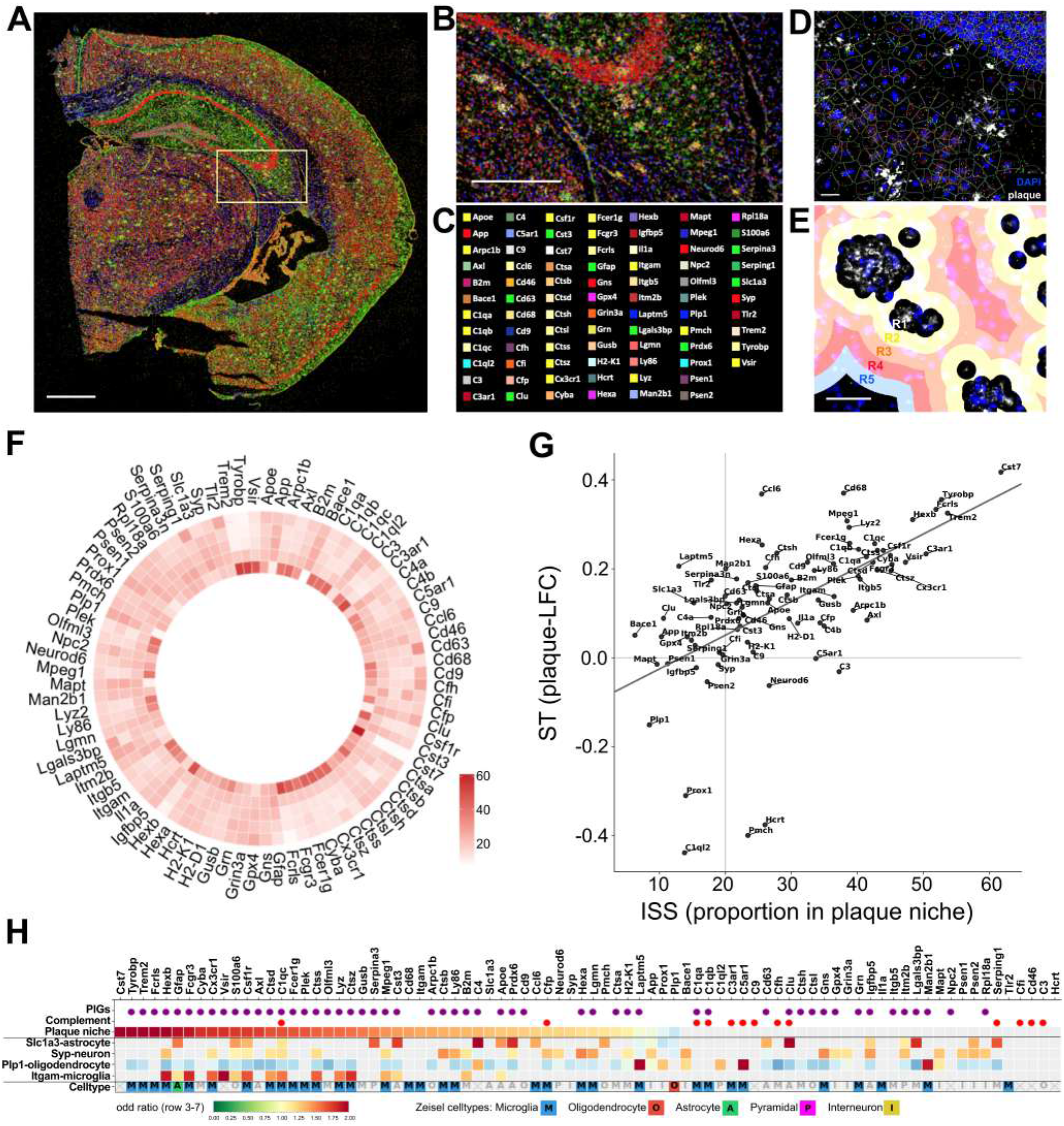
Validation of Spatial Transcriptomics analysis by *in situ* sequencing. (A, B) Distribution of 84 genes in the coronal section (A) and zoom into the hippocampus (B) of *App*^*NL-G-F*^ mice at 18-month of age demonstrated by *in situ* sequencing (ISS). Red neuronal (eg. *Syp*, *Neurod6*, *Grin3a*); yellow microglial (eg. *Itgam*, *Cx3cr1*, *Csf1r*); green astroglial (eg. *Slc1a3*, *Gfap*, *Serpina3n*) and blue oligodendroglial transcripts (eg. *Plp1*, *Laptm5*) are indicated and displayed in (C). (D) Cellular segmentation using watershed method based on DAPI-positive nuclei shown in hippocampus (DG_po). Color dots follow the same color code as in (C). Amyloid plaques are white while nuclei are blue. (E) Rings circumscribing amyloid plaques from R1 (ring 1, the most inner ring) to R5 (ring 5, the most outer ring). Scale bars: 800 μm (A), 400 μm (B), 20 μm (D), and 50 μm (E). (F) Circular heatmap shows gene expression in the indicated rings (inner circle: ring 1; most outer circle: ring 5). (G) Scatter plot shows the LFC of each target according to the plaque index as detected via ST (y-axis) or the expression of each target in ring 1 (the plaque niche) in ISS (x-axis) at 18-month of age. (H) Cellular signature of the 84 genes. PIGs (1^st^ row) and complement components (2^nd^ row) are highlighted. The significant odds ratio of an indicated gene being in plaque niches (ring 1) is displayed in row 3, and in the significant odds ratio of being in a selective cell type within the plaque niches (ring1) of the ISS result is displayed in row 4-7, non-significant odds are colored by grey. Zeisel et al. defined (color background) / predicted cell type (grey background) genes are highlighted in the last row.

Many genes show differential expression only in ring 1, instead of more gradual alterations across multiple rings (Figure 5F). We applied logistic regression for each gene which shows that of the 53 detected PIGs, 36 are significantly enriched in ring 1 (odds ratio>1, padj <0.05, see the Methods, Figure 5H), including 12 ARM/DAM microglial markers, 4 A1-astroglial markers and 21 other PIGs (Figure 5H, row3) while *Laptm5* was significantly depleted in ring 1 (odds ratio <1, padj < 0.05, Figures 5F and 5H). The result confirms that the detected gene expression alterations in ST are unique to the microenvironment of plaques. The data from the ISS (gene enrichment in the plaque niche) correlate well (cor = 0.6, p-value = 2.659e^−09^) with the plaque model derived from ST data (Figure 5G). An independent second run of ISS confirms the results.

We further analyzed which cell types contribute to the PIGs expression in the plaque niche (see Methods, Figure S5), which confirmed PIGs expressing in microglia and astrocytes (Figure 5H, row 4-7). In addition, oligodendrocytes express complement *C4* and lysosomal genes *Man2b1* and *Laptm5* (odds ratio >1, padj<0.05). Dysregulation of neuronal lysosomes in the plaque niches was also suggested (odds ratio >1, padj<0.05), including changes in *Gns* (catabolism of heparin and heparan sulfate), *Hexa* (degradation of ganglioside GM2 and hexosamines) and *Lgmn* and *Ctsb* (proteolysis). We finally validated the cellular signature of PIGs involved in the complement cascade, using RNAscope *in situ* hybridization on 3 different mice per genotype at 18-month of age. We confirmed that *C1qa* was specifically expressed by *Itgam*-positive microglia close to amyloid plaques (Figure 6B); *C4* was expressed by *Slc1a3*-positive astrocytes and *Mbp*-positive oligodendrocytes, while *Clu* was expressed by *Slc1a3*-positive astrocytes close to amyloid plaques in *App*^*NL-G-F*^ at 18-month of age (Figures 6).

**Figure 6.**
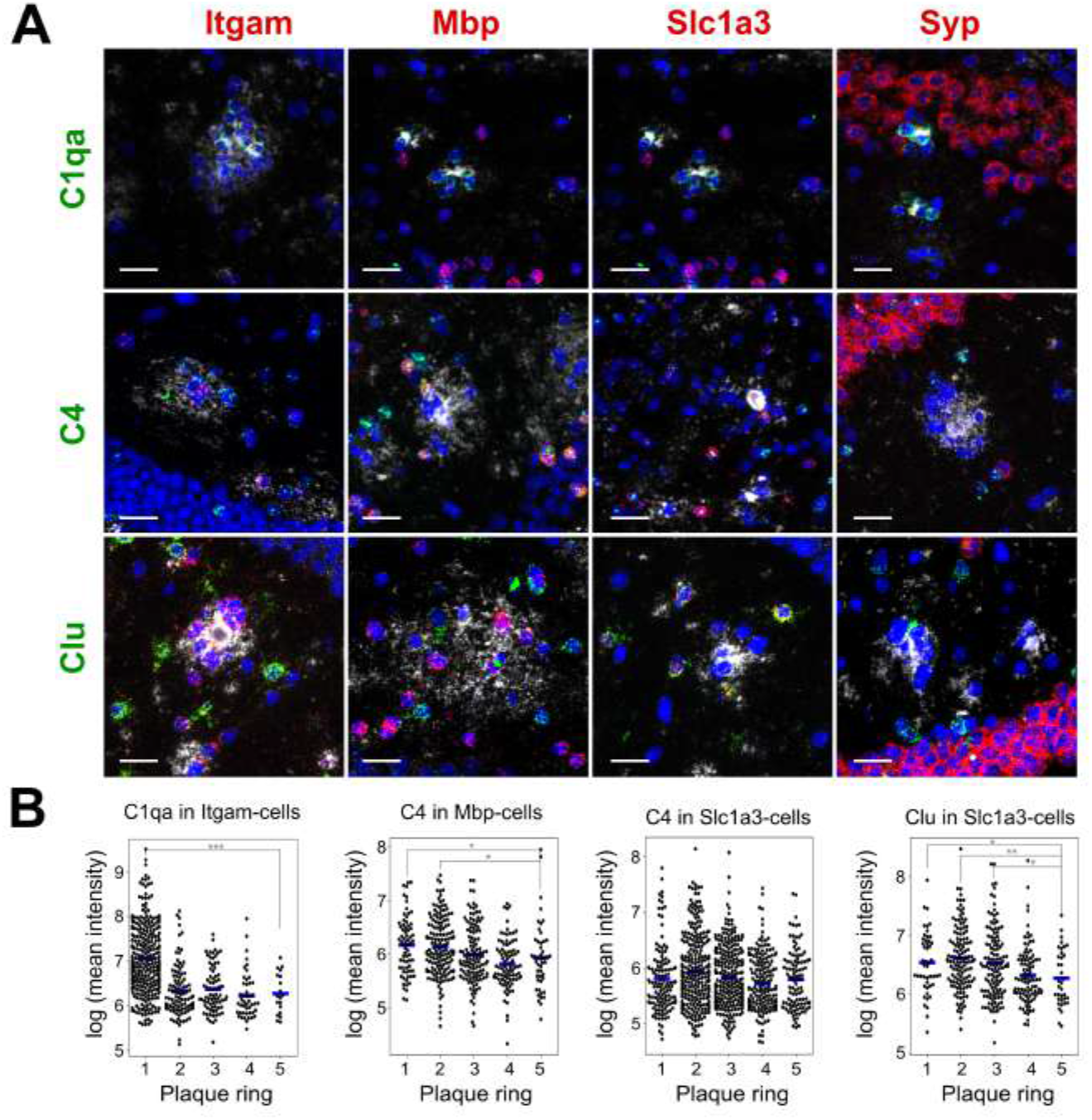
Microglia, astrocytes and oligodendrocytes are involved in the activation of the complement cascade around amyloid plaques. (A) Combined RNAscope and immunofluorescence analysis of complement components (*C1qa*, *C4* or *Clu* probes in green) expression by microglia (*Itgam*, red), oligodendrocytes (*Mbp*, red), astrocytes (*Slc1a3*, red), and neurons (*Syp*, red) in the vicinity of amyloid plaque (6E10, white) in the hippocampus of *App*^*NL-G-F*^ mice at 18-month of age. Nuclei are blue (DAPI). Scale bar: 25 μm. (B) Quantification of *C1qa*, *C4* and *Clu* staining intensity per cell, classified based on cell type and distance from a plaque. Measurements were made from >2900 single cells for each condition from 3 *App*^*NL-G-F*^ mice. Expression of *C1qa* in microglia, of *C4* in oligodendrocytes, and *Clu* in astrocytes were increased in the plaque niches. * p < 0.05, ** p < 0.005, *** p < 0.0005, Mann Whitney U test compared to the 5th ring.

### Astroglial PIGs and microglial PIGs are co-expressed exclusively in the amyloid plaque niches

A final question is to what extent the PIG response is a coordinated multi-cellular response to the amyloid plaques. We therefore recalculated co-expression matrix of the PIGs in different quartiles of plaque exposure by splitting the full database into one WT and four AD sets (Figure 7B). In WGCNA, the connectivity measures how strongly expression of a gene correlates with all other network genes of tissue domains from the same quartile (or WT). These results are visualized here by Circos plots in Figure 7A. The depth of color indicates the level of correlation between each two genes in a subset of tissue domains. In WT mice, the overall connectivity of the PIGs is relatively low and the PIGs split into 3 sub-networks, an astroglial (green), a microglial (blue) and finally a third group of genes that do not cluster nor co-express with other genes (grey) (Figures 7A and S6A). Subsequently, in the AD mice, we see a dose-dependent effect of plaque load on the strength of connectivity between the PIGs. In quartile 1 (Q1), calculated from the 25% tissue domains with the lowest amyloid exposure, the 3 sub-groups identified above appear somewhat co-expressed but the connectivity remains relatively weak. In Q4, with tissue domains exposed to the highest amyloid burden, a very strong co-expression pattern across almost all PIGs can be observed (Figures 7A and S6B). The PIGs in astrocytes and microglia thus only co-express in the tissue domains that encompass the amyloid plaques.

**Figure 7.**
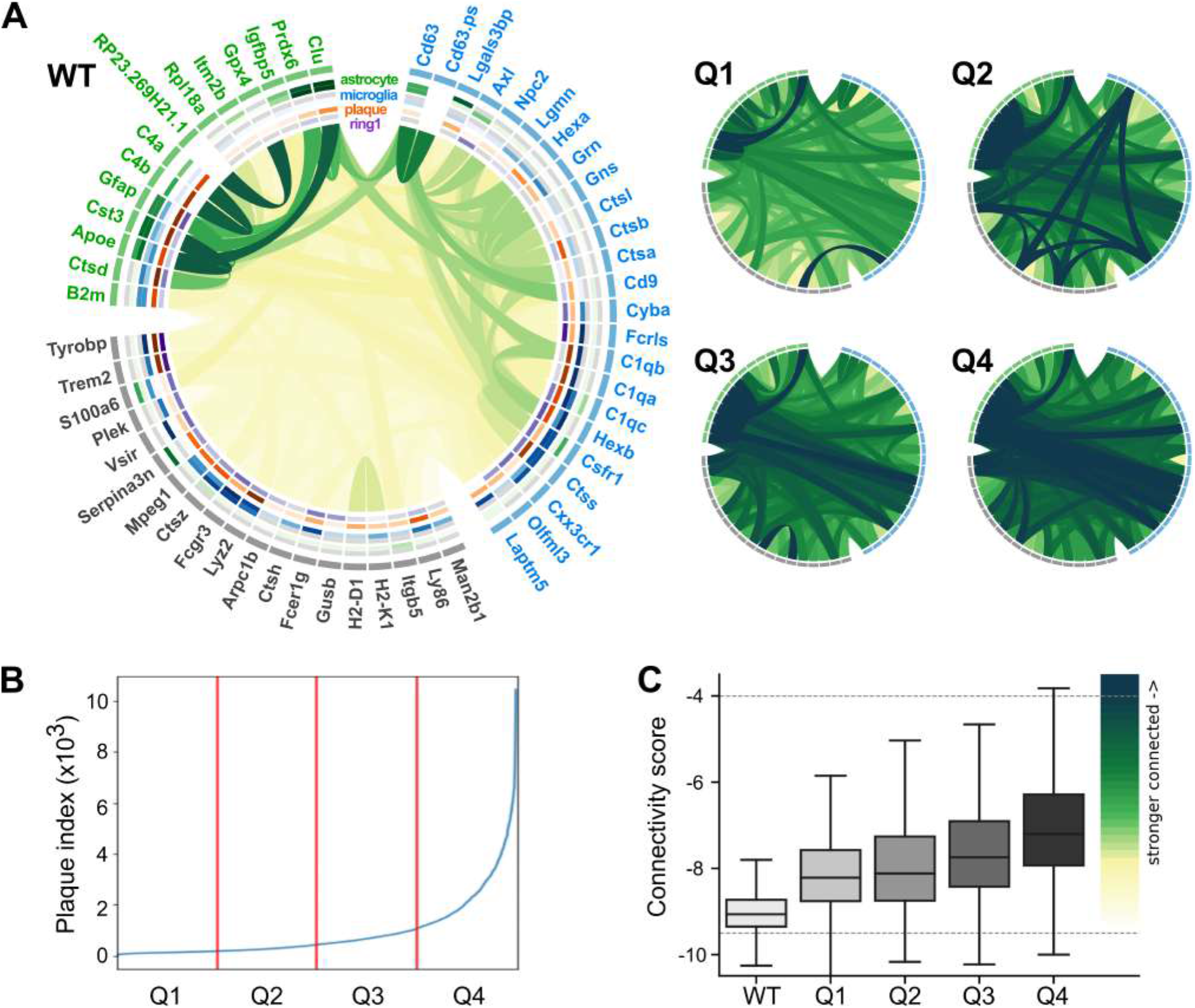
Astroglial PIGs and microglial PIGs are co-expressed in the plaque niches but not in the control tissue domains. (A) Circos plot: connectivity map derived from the pairwise correlation score derived from WGCNA adjacency matrix of PIGs in different subgroups. Notice the segregation into 3 subgroups (blue, green, and grey) in WT. Cell types are indicated using either average z-score from single-cell sequencing data (outer ring)(Zeisel *et al.*, 2015) or the odd ratios of the ISS results (inner ring) in the plaque niches. The strength of correlation of a gene expression with the plaque load across all AD tissue domains is indicated in the orange ring. The odd ratio of a gene presenting in ring1 is indicated in the most inner ring. The green lines indicate the strength of connectivity score between genes. Notice the dose sensitive increase in connection of the genes with increasing amyloid index from Q1 to Q4 with Q1 <25% lowest and Q4 >75% highest exposed tissue domains. (B) Quantile selection of AD tissue domains (x-axis: tissue domains sorted by plaque index) according to the plaque load (y-axis: plaque index). (C) The mean of the connectivity score of PIGs in the indicated tissue domains. The color depths of green curve in panel A is indicated.

## Discussion

The current work provides an extensive documentation of the gene expression alterations during the cellular phase of AD in a knock-in mouse model for amyloid plaques. The most striking finding is the gradual establishment of a multicellular co-expressed gene expression response towards amyloid plaques propagated by the different cell types in the amyloid plaque niche (Figure 7B). We identified 57 Plaque Induced Genes or PIGs that characterize this pathway and that are expressed in microglia, astroglia and oligodendrocytes (Figures 4 to 7). Furthermore, we found an unexpected response of the oligodendrocytes, separate from the PIGs response, that is characterized by an early increased expression of myelination genes at 3 months (Figure 3A) and a decreased expression at 18 months in the plaque niches (Figure 3B). The later response is already pronounced in hippocampus and entorhinal cortex in the early stage (Figure 3C), suggesting aspects of regional vulnerability, or at least brain region specific responses, to amyloid plaques.

### Spatial transcriptomics to probe the multicellular environment of the amyloid plaques

The data supporting our conclusions are derived from the spatial transcriptomics (ST) analysis of 10 control and 10 diseased coronal slices of different mice at different ages, encompassing >10.000 small brain volumes (8×10^−5^ mm^3^) covering whole coronal sections between bregma −2.0 to −2.2. As discussed in the result section, such volumes are ideally suited to probe the changes in gene expression in the cellular microenvironment of the amyloid plaques (the so-called amyloid plaque niches). To increase cellular resolution, we combined this unbiased approach with additional single molecule approaches, i.e. classical *in situ* hybridization using RNAscope and independent multi-molecule *in situ* sequencing experiments (Ke et a*l*., 2013). We notice the remarkable congruency of the data: the single molecule approaches confirm in general very well the predictions based on the unbiased spatial transcriptomic approach (Figures 2D and 5G). Thus, our experiments provide a strong validation of the ST technology as an excellent approach to evaluate gene expression changes in brain. In contrast to single nuclei (Mathys et al., 2019) or single cell approaches (Zeisel et al., 2015; Keren-Shaul et a*l*., 2017), ST does not require harsh dissociation procedures, maintains the spatial resolution of the data, and includes cytoplasmic, axonal or dendritic RNA. ST should be considered as the first choice for further systematic in depth analysis of the cellular phase in human AD samples. The major limitation of the technique at this moment, i.e. 100 μm resolution, is rapidly overcome by recent technical improvements such as Slide-seq (Rodriques et a*l*., 2019) or higher resolution arrays (Vickovic *et al.*, 2019).

### Complement as an important part of the intercellular crosstalk in the amyloid plaque niche

The 57 PIGs identified here reflect part of a multicellular response induced by the presence of amyloid plaques and involve parts of already known cellular reactions such as the activated astroglia (A1, Figure 4B) and microglia (DAM/ARM, Figure 4B). In addition, 36 PIGs are not known as part of the signature of disease-associated glia (Figure 4B). Based on the enrichment of GO terms in the PIGs module (Figure 4E), we conclude that the module is involved in the activation of the classical complement cascade (GO:0006956, e.g. *C1qa, C1qb, C1qc, C4a, C4b*), but also in effector mechanisms triggered by the complement cascade (Schmidt and Gessner, 2005; Thielens *et al.*, 2017) such as endocytosis (GO:0045807, e.g. *Fcer1g*, *Fcgr3*, *B2m*, *Cd63*, *Cyba*, *Apoe*, *Clu*, *Axl*), lysosomal degradation (GO:0005764, eg. *Hexa*, *Hexb*, *Ctsa*, *Ctsb*, *Ctsd*, *Ctsh*, *Ctsl*, *Ctss*, *Ctsz*, *Laptm5*, *Man2b1*, *Cd63*, *Gusb*, *Lgmn*, *Npc2*, *Grn*, *Gns*, *Prdx6*, *Cst3*), antigen processing and presentation (GO:0002474, e.g. *Fcer1g*, *Fcgr3*, *B2m*, *H2-D1*, *H2-K1*), immune response (GO:0002376, e.g. *Csf1r*, *Cx3cr1*, *Ly86*, *Trem2*, *Tyrobp*, *Vsir*), and oxidation-reduction processes (GO:0055114, e.g. *Cyba*, *Prdx6*, *Gpx4*). In addition, also some microglial PIGs are in fact homeostatic markers (e.g. *Cx3cr1*, *Olfml3*, *Csf1r*), confirming increased numbers of microglia in the plaque niche (Figure S5D). We propose that the network of 57 PIGs constitutes an intercellular crosstalk genetic network between astrocytes and microglia which results, among others, in the inappropriate control of the classical complement cascade.

Inappropriate control of the classical complement cascade causes unresolvable inflammation and disease (Hong *et al.*, 2016; Hansen, Hanson and Sheng, 2018; Morgan, 2018). The classical complement cascade is triggered by C1q-complex, and C1q is mostly expressed by microglia in the first cellular envelope around amyloid plaque, as shown in Figure 6 (Eikelenboom *et al.*, 1988; McGeer *et al.*, 1989). The release of C1q from microglia is necessary to induce neurotoxic, inflammatory astrocytes (A1) (Liddelow *et al.*, 2017), which also strongly upregulates genes involved in classical complement activation such *C4* (Figure 6) (Zamanian *et al.*, 2012). An exciting finding is illustrated in Figure 7, demonstrating how the overall connectivity between the PIGs is low in the wild type brains and increases with amyloid plaque load in the *App*^*NL-G-F*^ brains. In the 25% spots with the highest amyloid burden, almost all PIGs show strong correlation of expression. We found that genes associated with risk of AD as defined by GWAS studies were significantly enriched in this PIG module (odds ratio: 8.26, p-value = 0.009, Figure S4C, see the Methods). The PIGs thus show specific co-expression with GWAS hits such as *Apoe, Trem2* and *Clu* in the amyloid plaque niches (Figure 7), suggesting the interplay of ApoE-Trem2 (Krasemann et al., 2017; Parhizkar et a*l.*, 2019) with the complement pathway (Yin *et al.*, 2019).

### Oligodendrocytes and neurons are part of the cellular phase in the amyloid plaque niche

The oligodendrocyte co-expressed network response (OL) to amyloid plaques is of high interest. This OL appears to be upregulated in the early phase and downregulated in the later phase of the disease. Oligodendrocytes and myelin have received little attention in the field of AD research (Braak and Braak, 1996; Ettle, Schlachetzki and Winkler, 2016). Recently, Zhang et al. suggested that oligodendrocyte precursor cells (OPCs) surrounding amyloid plaques undergo senescence and stop differentiating into myelin-repairing oligodendrocytes. Instead, they release inflammatory molecules into the plaque environment (Zhang *et al.*, 2019), which may trigger or enhance the local activation of microglia and production of proinflammatory cytokines. In the current study, we confirm the depletion of myelin transcripts but also increased expression of complement C4 by Mbp-positive oligodendrocytes in the amyloid plaque niches at 18 months of disease. In contrast, we find increased myelin transcripts in the amyloid plaque niches at 3 months. We speculate that an initial remyelination reaction around the amyloid plaques could be part of a homeostatic response to keep the brain functional in the early phase of disease. When aging, OPC senescence induced by the amyloid plaques and dysregulated microglia phenotypes may stop remyelination directly or indirectly via inducing inflammation. Together with activated astrocytes and microglia, oligodendrocytes are thus clearly involved in the dysregulated environment of the amyloid plaque niches.

Space constraints do not allow us to discuss in much depth other salient features of the cellular response but we would like to highlight a few observations we made. For instance, we identified three neuronal networks by WGCNA (Figure S4: magenta, brown and pink). The interneuron module “pink” (136 genes including *Gabrg1*, *Baiap3*, *Glra3*, *Ahi1*, *Cartpt*, *Calb2*, *Camk2d*, *Camk2n2*, *Crebl2*) is down-regulated along the plaque-axis at 3-month of age, indicating the early vulnerability of interneurons in the amyloid plaque niches (Figure S4B). We also identified an astroglial-vascular module (black, 150 genes including *mt-Co1*, *mt-Nd1*, *Hba-a1*, *Hbb-bs*, *Aldoc*, *Slc7a10*, Figure S4, Table S4), mostly enriched in the CA_slm layers and up-regulated in the AD mice with aging, indicating the dysregulation of metabolism in the diseased aging hippocampus.

### Some current limitations of the approach

The current dataset is limited to the *App*^*NL-G-F*^ model. Although this is probably the best model for Aβ pathology at the moment, it comes short with regard to the development of neurofibrillary tangles, neurodegeneration, and memory deficits. The *App*^*NL-G-F*^ amyloid model is however excellent to study brain-inflammatory alterations as recently demonstrated in a study comparing human and mouse brain (Castillo *et al.*, 2017). Methodological limitations should be taken into account when analyzing these data. Notable is the fact that the number of reads captured per ST tissue domain is lower than in bulk RNA sequencing, and more on par with single cell transcriptomics. The available ST technology does also not reach “single-cell” level. This is complemented by *in-situ* sequencing experiments which strengthened the conclusions drawn from the ST analysis, but were impeded by low coverage per cell and the challenge to define cell identity and boundary. We observed for instance that there is only 22% overlap between *Cx3cr1*-positive microglia and *Csf1r*-positive microglia, and that 16% of Slc1a3-positive astrocytes express *Plp1*. Zeisel et al. have suggested that 3 marker genes are needed for a reliable cell type estimation (Zeisel *et al.*, 2018). However, increasing the number of targets for *in situ* requires higher magnification imaging to avoid optical crowding, at the expense of speed and throughput. Further improvement of cell type identification is however crucial.

## Conclusion

We present here a brain disease oriented application of the recently developed Spatial Transcriptomics technology (Ståhl *et al.*, 2016), demonstrating its strengths and limitations to study with unprecedented depth diverse cellular reactions triggered by the amyloid plaques. The data unequivocally demonstrate that amyloid plaques are not innocent bystanders of the disease as has been sometimes suggested (Robakis, 2010; Kametani and Hasegawa, 2018), but are in fact inducing strong and coordinated responses of all cells in the amyloid plaque niche. We also demonstrate that the action radius is limited to 10-20 μm from the plaque. Further work is needed to understand whether and when removal of amyloid plaques by for instance immunisation (Gallardo and Holtzman, 2017) is sufficient to reverse these ongoing cellular processes. It is tempting to speculate that antibody binding to amyloid plaques on its own would already influence these glia responses, which might be beneficial but also harmful for the progression of the disease.

This work is a first step into the direction of the human brain Alzheimer atlas (De Strooper and Karran, 2016), which should describe the different stages of the disease at the multi-omics level. Such resource would accelerate research into this dreadful disease significantly. Obviously, this will require the collaboration of several research groups and open access and data sharing will be essential to achieve this ultimate goal.

## Acknowledgments

This work was supported by INSTALZ, EU Joint Programme - Neurodegenerative Disease Research (JPND), Fonds voor Wetenschappelijk Onderzoek (FWO), Methusalem grant from the KU Leuven/Flemish Government, “Opening the Future” of the Leuven Universiteit Fonds (LUF), Stichting Alzheimer onderzoek (SAO), the Alzheimer Association (USA), MRC, Alzheimer Society (UK), Alzheimer Research UK, VIB Tech Watch, the Knut and Alice Wallenberg foundation, and the Thon foundation. W.T.C. is supported by Taiwan Ministry of Science and Technology (MOST-105-2917-I-564-081), and Marie Skłodowska-Curie No 665501. S.M. and the imaging is supported by FWO grants FWO I001818N, (AKUL/17/044); FWO I001719N and AKUL/13/39. B.D.S. is supported by an ERC-consolidator grant, the Geneeskundige Stichting Koningin Elisabeth, and the Bax-Vanluffelen Chair for Alzheimer’s Disease. The authors thank Nikky Corthout (VIB Image core). Sequencing was carried out by the VIB Nucleomics Core.

## Author Contributions

W.T.C., A.L., M.F., and B.D.S. conceptualized and designed the study. K.C., A.S., C.S.F., and W.T.C. optimized and performed spatial transcriptomics (ST) together with inventors A.J., J.F.N. and J.L.. X.Q., J.L., M.K. and W.T.C. optimized and performed *in situ* sequencing together with immunostaining. W.T.C. performed and analyzed *in situ* hybridization (RNAscope) with I.V. and L.W.. W.T.C. and B.P. optimized the image analysis of ST with J.F.N. and the image analysis of *in situ* sequencing with X.Q. along with data analysis input from M.F.. B.P. developed programs for all image analysis. S.M. setup the slide scanner imaging supported imaging and image registration. A.L. and M.F. conceptualized and executed all bioinformatics analysis with biological input of W.T.C., R.M. and B.D.S..T.C.S. contributed animals. B.D.S. is responsible for the funding. The first draft of the manuscript was written by W.T.C., A.L., M.F. and B.D.S., with input from all co-authors. All co-authors read and approved the final version of the manuscript.

## KEY RESOURCES TABLE

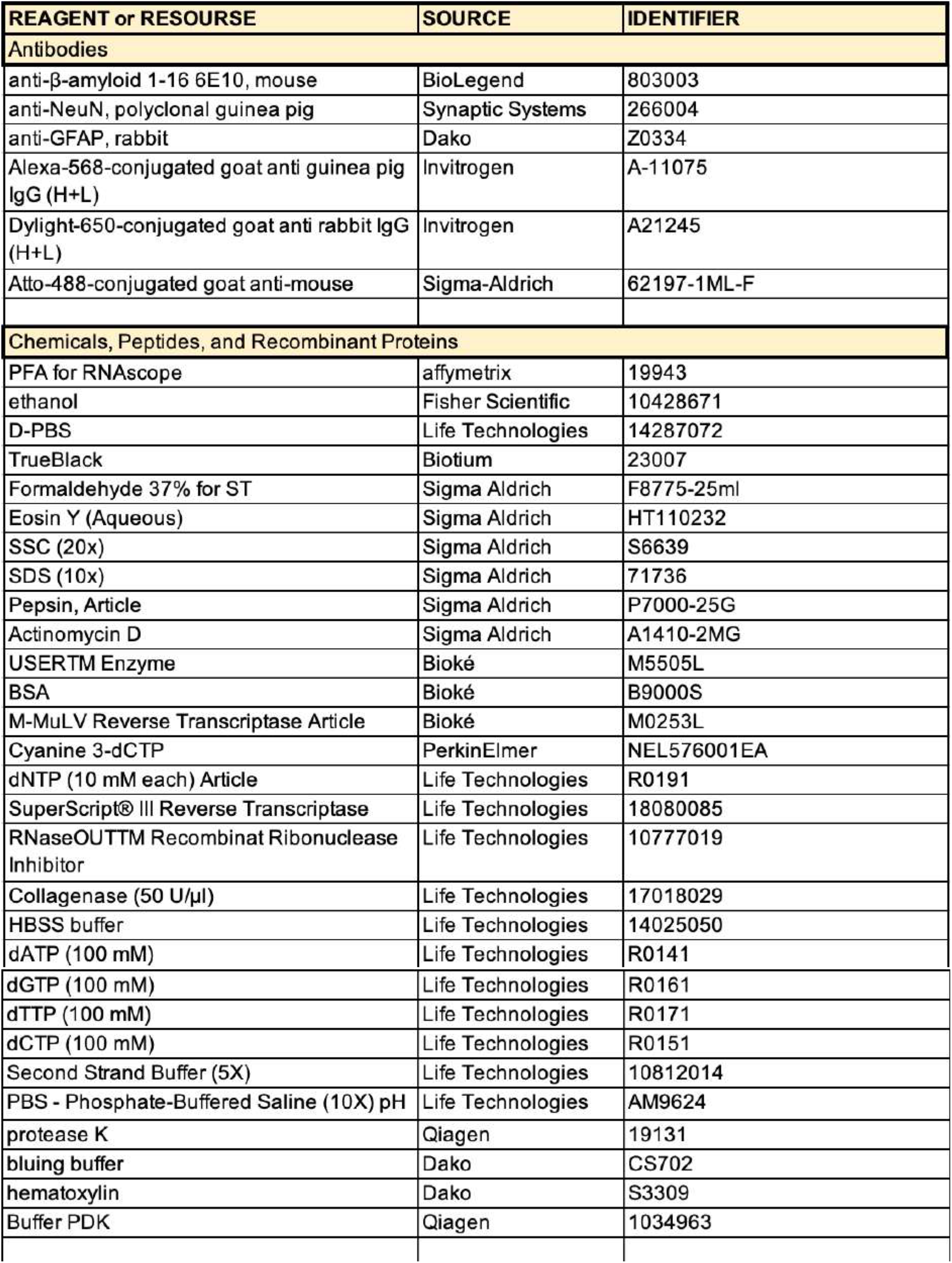
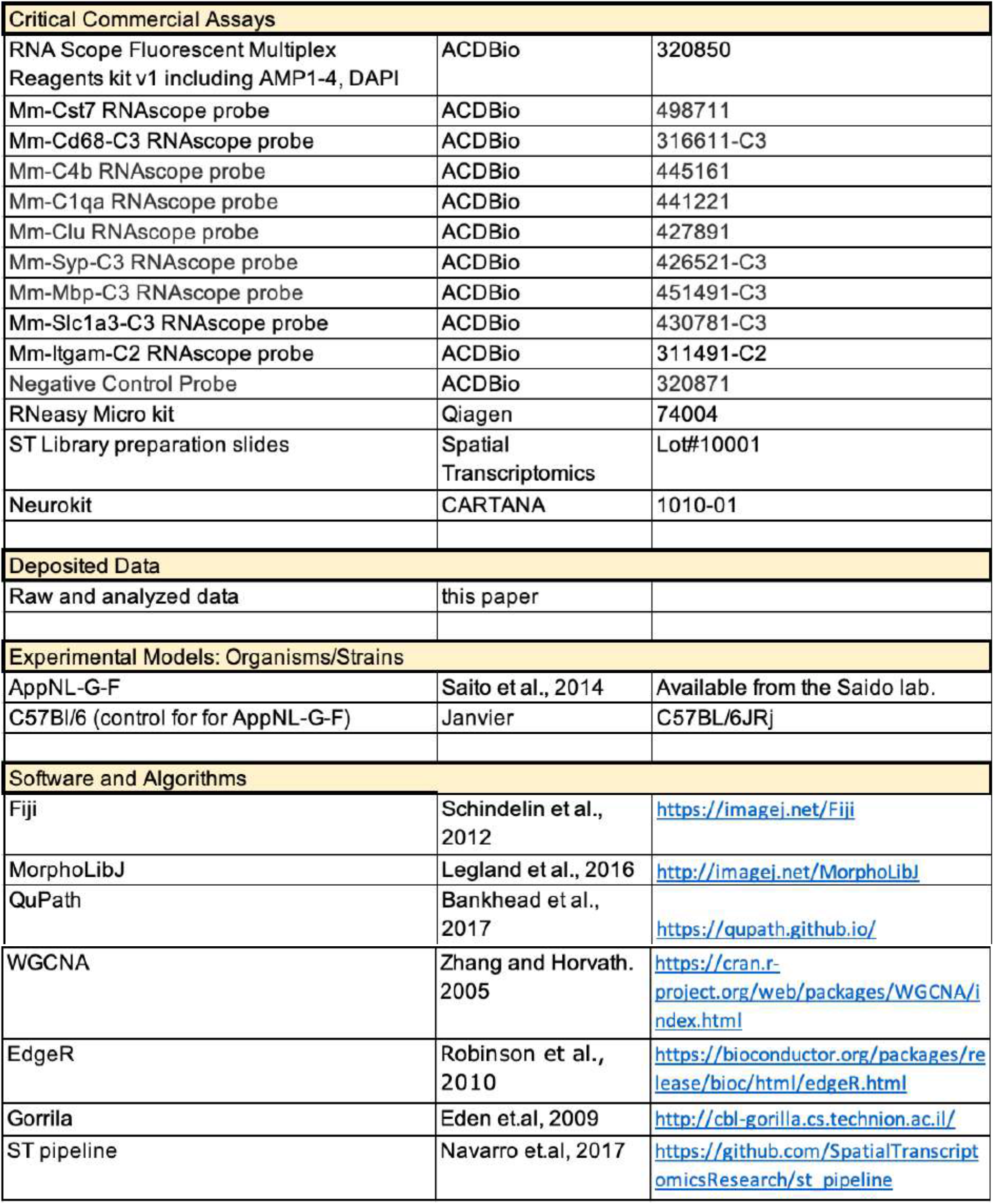

## CONTACT FOR REAGENT AND RESOURCE SHARING

Further information and requests for reagents may be directed to, and will be fulfilled by the corresponding author Bart De Strooper

## EXPERIMENTAL MODEL AND SUBJECT DETAILS

### Mice

All animal experiments were conducted according to protocols approved by the local Ethical Committee of Laboratory Animals of the KU Leuven (governmental license LA1210591, ECD project number P056-2016) following governmental and EU guidelines. *App*^*NL-G-F*^ knock-in (Saito et al., 2014) mice express Swedish (KM670/671NL), Beyreuther/Iberian (I716F), and Arctic (E693G) mutations in the *APP* gene under the endogenous promoter on the C57BL/6J background. *App*^*NL-G-F*^ mice were backcrossed for at least 2 generations with C57BL/6J mice in the De Strooper lab.

Male mice (*App*^*NL-G-F*^ KI and C57BL/6J controls) were sacrificed at 3.5 months (106 days), 6 months (average 183.5 days), 12 months (average, 367 days), or 18 months (average 551 days) of age, giving rise to 8 experimental groups (n=2 per group for 3.5− and 18-months of age; n=1 per group for 6− and 12-months of age; n=12 for total): WT_03, WT_06, WT_12, WT_18, AD_03, AD_06, AD_12, AD_18. Following cervical dislocation, left and right hemispheres were embedded in cold OCT separately and snap-frozen in liquid nitrogen. Samples were stored at −80°C.

## METHOD DETAILS

### Tissue collection for Spatial Transcriptomics

OCT-embedded hemispheres were cryosectioned coronally to a thickness of 10 μm (bregma: −2.0 to −2.2) using a CryoStar NX70 cryostat (ThermoFisher). We layered tissue sections onto a spatially barcoded array to collect *in situ* 2D-RNAseq of Spatial Transcriptomics (Lot#10001, Spatial Transcriptomics, Stockholm, Sweden) or to a regular glass slide for immunohistochemistry. Each spatially barcoded array has 1007 tissue domains, with a diameter of 100 μm and a center-to-center distance of 200 μm, over an area of 6.2 mm by 6.6 mm. One coronal section normally covers the area of 500 to 600 spots on the array, each defining on tissue domain. Each spot contains approximately 200 million barcoded reverse-transcription oligo(dT) primers allowing to get a global transcriptomic profile of a tissue domain with a volume of 0.00008 mm^3^. After cryosection, all sections were stored at −80°C before proceeding with experiments.

We collected one right and one left hemisphere for each experimental group at 3.5− and 18-months of age, and one right hemisphere for each experimental group at 6− and 12-months of age. We performed experiments of *App*^*NL-G-F*^ KI mice and C57BL/6J at the same age at the same time. RNA quality was checked by RNeasy Micro Kit (Qiagen, Hilden, Germany) and Agilent 2100 Bioanalyzer with RNA nano chips (Agilent Technologies, Inc., Santa Clara, CA, USA). RIN values of the tissues were between 8.6 to 9.45. Details of sample quality is described in Table S1.

### In situ 2D-RNAseq via Spatial Transcriptomics

Spatial Transcriptomics experiments were performed following the Library Preparation Manual (Spatial Transcriptomics, Stockholm, Sweden) (Ståhl et al., 2016). Briefly, cryosectioned tissues were fixed on a spatially barcoded array by 3.7% formaldehyde solution at room temperature for 10 min, and stained by hematoxylin for 7 min, bluing buffer for 2 min, and eosin for 20 sec at room temperature. We acquired HE images by Zeiss Axio Scan.Z1 slidescanner (Carl Zeiss AG, Oberkochen, Germany). After imaging, tissues were immediately permeabilized by collagenase in HBSS-BSA buffer for 20 min and 0.1% pepsin in 0.1M HCl for 6 min at 37°C, and followed by *in situ* reverse transcription by adding cDNA synthesis master mix at 42°C for 18-20 hours to get the stable cDNA attached on the array. Tissues on the array were then removed by the incubation in 2.5 mg/ml proteinase K in PDK buffer at 56°C for 1h with interval shaking (300 rpm, 15s shake, 15s rest), which left cDNA coupled to the arrayed oligonucleotides on the slide. We collected the cDNA probes by probe cleavage using 100U/ml USER enzyme in 1X second strand buffer with dNTP and BSA. Library preparation of the released cDNA probes was performed in the laboratory of our collaborato led by Prof. Joakim Lundeberg at KTH Royal Institute of Technology, Sweden, including second strand synthesis, in vitro transcription, adapter ligation, second cDNA synthesis, qPCR quantification, and PCR amplification. The number of cycles to amplify the final libraries is between 8-11 cycles. Library quality was checked on an Agilent BioAnalyser DNA High Sensitivity chip. We selected two out of six libraries per mouse according to lower amplification cycles, better tissue morphology, higher RIN value, and similar length of cDNA to perform paired end sequencing on an Illumina NextSeq500 sequencer at the VIB Nucleomics Core (Leuven, Belgium). To determine the exact localization and quality of each of the 1007 tissue domains on the array, a fluorescent Cyanine-3 probe was hybridized to the remaining DNA capture probes and the arrays were scanned at 548 nm by Zeiss Axio Scan.Z1 slidescanner (Carl Zeiss AG, Oberkochen, Germany). The image was used together with the HE-image to annotate the spatial localization of each tissue domain using the Allan Brain atlas as reference.

### Immunohistochemistry of Spatial Transcriptomics

Immunohistochemistry was performed on the two adjacent sections to the one used for sequencing. After fixation in 4% ice-cold paraformaldehyde (PFA) for 20 min and a wash with PBS, we performed antigen retrieval by microwave boiling the tissue 3 times in 10 mM sodium citrate at pH 6.0 to expose antigenic sites. After cooling down to room temperature for 20 min, brain tissues were washed and blocked in TBS-buffer solution containing 0.5% Triton X-100 and 5% normal goat serum for 2h. The serum-blocked tissues were then stained with mouse Alexa Fluor 488 anti-Aβ_1-16_ antibody, 6E10 (803013, BioLegend) at 4°C overnight, and guinea pig anti-NeuN antibody (266004, Millipore) and rabbit anti-Gfap antibody (Z0334, DAKO) in blocking buffer at 4°C overnight. The immuno-stained tissues were then incubated with goat Alexa 594 anti-guinea pig IgG (H+L) antibody (106-585-003, Jackson Immunolabs) and goat Alexa 647 anti-rabbit IgG (H+L) antibody (A21245, Invitrogen) for 1.5 h at RT. After incubation with DAPI and mounting with glycerol, imaging was carried out on Zeiss Axio Scan.Z1 slidescanner (Carl Zeiss AG, Oberkochen, Germany) using a 20X objective. Volume images were acquired with 16-bit depth to allow a broad range of intensity values and rendered using Fiji (Schindelin *et al.*, 2012).

### In situ sequencing by CARTANA

OCT-embedded hemispheres of mice at 18-month of age were cryosectioned coronally into 14 μm (bregma −2.0 to −2.2) and layered onto SuperFrost Plus glass slides (ThermoFisher) and further stored at −80°C before experiments. Samples were shipped on dry ice to CARTANA (Solna, Sweden) for tissue fixation, reverse transcription, probe ligation, rolling cycle amplification with reagents and according to the procedures supplied in the Neurokit (1010-01, CARTANA, Sweden), followed by fluorescence labeling, and sequencing by sequential images at 20X objective (Ke *et al.*, 2013). Five probes were designed for each gene, except *Itgam*, which has 10 customized probes to increase the detection sensitivity. We included probes for 7 additional genes that do not belong to the PIG module but which significantly react to the presence of amyloid plaques at 18-months of age in the ST analysis: *Cst7*, *Cd68*, *Ccl6*, *Prox1*, *Hcrt*, *Pmch*, and *C1ql2* (LFC: 1.91, 1.70, 1.69, −1.42, −1.72, −1.83, and −2.01, respectively). We removed afterwards probes *H2-D1*, *Cd63-ps* and *RP23-269H21.1* because of design issues and renamed the probes *C4a*, *C4b*, *Serpina3n*, *Lyz2* as *C4*, *Serpina3* and *Lyz* because they cross react with related genes. To reduce lipofuscin autofluorescence, 1X TrueBlack (Biotium, Fremont, CA) was applied for 30 sec before fluorescence labeling. The result table of the spatial coordinates of each molecule of 84 targets together with the reference DAPI image per sample were provided by CARTANA. After in situ sequencing, samples were shipped back to the lead laboratory in Leuven for further immunostaining of amyloid plaques on the same tissue. Briefly, after removing coverslips, the brain tissues were washed and blocked in TBS-buffer solution containing 0.5% Triton X-100 and 5% normal goat serum for 2h. The serum-blocked tissues were then stained with mouse Alexa Fluor 488 anti-Aβ_1-16_ antibody, 6E10 (803013, BioLegend) at 4°C overnight. After immunostaining, sections were then incubated in 1X TrueBlack (Biotium, Fremont, CA) solution for 30 sec to reduce lipofuscin autofluorescence. After staining with DAPI (Sigma-Aldrich) and mounting with FluorSave Reagent (Merck Millipore, Burlington, MA). Imaging was carried out on MÄRZHÄUSER SlideExpress 2 with 20X objective and Hamamatsu ORCA Flash4.0 camera. Volume images were acquired with 16-bit depth to allow for a broad range of intensity values and rendered using Fiji.

### Multiplexing RNAscope and immunohistochemistry

OCT-embedded hemispheres of mice were cryosectioned coronally into 14 μm (bregma −2.0 to −2.2) and layered onto SuperFrost Plus glass slides (ThermoFisher) and further stored at −80°C before experiments. RNAscope experiments were performed using the Manual Fluorescent Multiplex kit v1 (Advanced Cell Diagnostics, Newark, CA) following manufacturer’s recommendations with minor adjustments. Briefly, after fixation and protease digestion, probe hybridization was carried out at 40°C for 2 h with the indicated probe sets. Probes were all from Advanced Cell Diagnostics: Mm-Cst7 (498711), Mm-Cd68-C3 (316611-C3), Mm-C4 (445161), Mm-C1qa (44221), Mm-Clu (427891), Mm-Syp-C3 (426521-C3), Mm-Mbp-C3 (451491-C3), Mm-Slc1a3-C3 (430781-C3), and Mm-Itgam-C2 (311491-C2). After amplification steps to obtain the RNAscope signals, we immediately performed immunohistochemistry to acquire the immunofluorescence picture of amyloid-beta plaques in the tissues. Briefly, the sections were blocked for 1 hour at RT in PBS containing 0.3% Triton X-100 and 5% normal goat serum and immunostained with the anti-Aβ_1-16_ primary antibody (6E10, BioLegend, San Diego, CA) at 4°C overnight and then with an Atto-488-conjugated goat anti-mouse secondary antibody (Sigma- Aldrich, Saint Louis, MO) at RT for 1h. After immunostaining, sections were incubated in 1X TrueBlack (Biotium, Fremont, CA) solution for 30 sec to reduce lipofuscin autofluorescence. After staining with DAPI (Sigma-Aldrich) and mounting with FluorSave Reagent (Merck Millipore, Burlington, MA), sections were imaged via a Leica TCS SP8 X confocal microscope (Leica Microsystems, Wetzlar, Germany) using a 40X objective.

## QUANTIFICATION AND STATISTICAL ANALYSIS

### Image analysis

#### Metadata of Spatial Transcriptomics

HE and Cy3-spot images were acquired from the middle ST sections. Fluorescent amyloid-beta, astrocyte, neuron, and nuclei images were acquired from the two adjacent sections stained by 6E10, anti-Gfap antibody, anti-NeuN antibody, and DAPI. Manually aligned HE and Cy3-spot images were used to bridge the transcriptomics picture with the immunostaining pictures. To de-barcode the spatial localization of each transcriptomic profile, we converted the pixel coordinates of the 1007 tissue domains on the Cy3-spot image into the theoretical coordinates described in the ID-file of the spatially barcoded array. To acquire the spatially corresponding amyloid and cellular information per tissue domain, we manually aligned and transformed fluorescence images into the corresponding HE images. To annotate the anatomic brain regions, we manually aligned and transformed the reference atlas from Allen Brain Institute into the corresponding HE images. More details of the reference atlas for each sample are described in Table S1. Image alignment was processed using the Fiji plugin “Landmark correspondences” (Legland, Arganda-Carreras and Andrey, 2016) and the precision of all aligned images were checked before analysis.

We developed a Fiji groovy script package to automate the image processing and analysis. Quality of the image has been checked before computation. We annotated the unreliable areas (eg. damaged tissue, out of focus, or dirt on the image) by manual assignment of region of interest (ROI) by Fiji. For each tissue domain, the package computed the percentage of damaged area, coverage area of tissue, coverage area of Cy3-detectable spot, and the coverage area of individual brain region according to the Allan Brain atlas images. We filtered the spots by the coverage area of tissue > 90%, damaged area of tissue <30%, and the coverage area of Cy3-detectable spot >90%. This results in 500-600 useful tissue domains for each sample.

To measure the immunostainings, our home-derived package also computed 5 parameters for the Aβ, the Gfap, the NeuN and the DAPI staining within each tissue domain: (1) mean pixel intensity, (2) median pixel intensity, (3) sum of pixel intensity, (4) standard deviation of pixel intensity, and (5) percentage of area of the computed positive signals per tissue domain. Together, the package generated the metadata with 146 observations for each tissue domain.

#### Quantification of immunostainings

To assign a single measure to a tissue domain for each of the immunostainings, we computed 5 statistics for each tissue domain and staining (Aβ, Gfap, NeuN and DAPI): (1) mean pixel intensity, (2) median pixel intensity, (3) sum of pixel intensity, (4) standard deviation of pixel intensity, and (5) percentage of area of the computed positive signals per tissue domain. To select the most representative statistics, a group of 8 experts were employed in a random ranking approach. In this exercise, two images of random tissue domains (showing the same staining) were shown side by side, and we asked the experts to identify the tissue domain image which, according to their best estimate, contains the highest amount of stained material. For each staining we scored a large number of pairs (Aβ: 1672, DAPI: 1685, NeuN: 1464, Gfap: 2271). Subsequently, we calculated for every scored pair and for each calculated parameter, the difference between the two tissue domains. We then used the Mann Whitney U test (MWU, p-value) to identify which parameter has the best power to distinguish between the two tissue domains, and, which parameter is most often in concordance with the expert. Based on this approach, we picked the percentage of area of the computed positive signals for Gfap (80.14% correct, MWU p=10^−297^), DAPI (79.35% correct, MWU p=10^−167^) and NeuN (72.13% correct, MWU p=10^−121^). Remarkably, the standard deviation of pixel intensity worked best for the Aβ (66.51%, correct, MWU p=10^−92^). We experimented with normalization per slide, but this did not improve the predictive power, so we proceeded without normalization. The final score taken per staining/tissue domain is the mean of the corresponding spot in the two adjacent slides.

#### Metadata of *in situ* sequencing

We developed a Fiji groovy script package to automate the metadata of *in situ* sequencing, including number of signals per targeted gene and spatial distance to amyloid plaque. To get the single cell resolution, we performed a white top-hat filtering followed by a Gaussian filtering and converted the DAPI signals into binary masks by using a histogram-derived threshold method, Triangle, in Fiji. Watershed segmentation on the DAPI staining was performed to separate the binary mask of DAPI. The segmented single cell/nucleus was expanded by 10 μm to identify the cytoplasm (Qian *et al.*, 2018; Sala Frigerio *et al.*, 2019). Within each ROI per cell, we computed the number of each target gene based on their spatial coordinates per molecule provided by CARTANA. To get the spatial relevance to amyloid plaque, we first manually aligned the amyloid immunostaining with the DAPI reference. As *in situ* sequencing and immunostaining were carried out on the same tissue, the precision of alignment is very high and has been checked before analysis. We converted the amyloid-positive signals into binary masks by using a histogram-derived threshold method, Triangle, in Image-J. The ROI of the plaque niche (ring 1) is based on the area mask with boundary expansion by 10 μm. We compute 5 co-centroid circles (donuts) from the ROI of ring1 in the plaque niche to the ROI of ring5 far from plaque with 18.2 μm extension per ring without overlap between plaques. The assignment of the spatial distance to plaque per cell is based on this classification of 5 rings. The metadata generated by this image analysis was further applied in the data analysis of the *in situ* sequencing experiment. We applied QuPath to overlay the spatial images of multiple targets, immunostainings, DAPI staining, and ROIs of cells and plaques, and to generate the representative images shown in the figures (Bankhead *et al.*, 2017).

#### RNAscope quantification

Three male *App*^*NL-G-F*^ KI mice at 18 month of age per experimental condition were used for quantification. For each mouse, 6 images of the hippocampus (two of CA1, two of CA3, and two of DG) were acquired and stacked in Fiji. NIS-elements software (Nikon, Amsterdam, Netherlands) was used to detect nuclei, microglia, astrocyte, neuron, oligodendrocyte, and plaque using a custom-made protocol (Sala Frigerio *et al.*, 2019). Nuclei and cell body perimeter were established using the DAPI signal; microglia, astrocytes, neurons, and oligodendrocytes were identified by RNAscope puncta from the *Itgam*, *Slc1a3*, *Syp*, and *Mbp* probes, respectively. All parameters were kept constant between images to allow unbiased detection. Around each plaque, five concentric circles were drawn, for each circle we counted microglia, astrocytes, neurons, and oligodendrocytes, and for each cell we measured the intensity of the signal (eg. *Cst7*, *Cd68*, *C1qa*, *C4*). To quantify the degree of change around amyloid plaques, we first log transformed the target intensity per cell, and classified their expression level per cell to the distance to amyloid plaques (ring).

### Sequencing Data Analysis

#### Generation of raw counts, cpm counts

Sequencing data were pre-processed with the ST pipeline (Navarro *et al.*, 2017), which filtered low quality bases, mapped against the mouse genome (Ensembl 88), and generated a count matrix. The count matrix was further filtered by removing spots with tissue coverage less than 30% in the HE image. The EdgeR “cpm” function was used for library size normalization and the output log-cpm matrix was used for the rest of the analyses.

#### Differential expression analysis

DE analysis was conducted by fitting two separate GLM models using plaque intensity and genotype information respectively. Each GLM model was tested for differential expression by using EdgeR quasi-likelihood F-test which accounts for the uncertainty in dispersion estimation at the age of 3 months and 18 months, separately. The plaque model represents transcriptional changes in the vicinity of amyloid plaques, which models the log transformed plaque index as a continuous variable, and its LFC indicates the changes in gene expression per unit change in plaque index. For a more straightforward interpretation, we multiply all LFC by a constant 4.59 (the difference between the maximum observed plaques and the minimum observed plaque index across the database). The corrected LFC represents the amount of changes in gene expression from the minimum to the maximum observed plaque load, and is used throughout the paper. The genotype model assesses transcriptional changes between WT and TG mice.

#### WGCNA

WGCNA package in R (Zhang and Horvath, 2005) was used to build signed co-expression networks. The set of genes with the highest 50% standard deviation was selected using the “varFilter” package from the Bioconductor (Huber *et al.*, 2015). Soft power 14 was chosen by WGCNA’s “pickSoftThreshold” function to calculate the adjacency matrix, and the module identification was performed by the “cutreeDynamic” function by selecting deepSplit =4. The adjacency matrix is calculated using the “adjacency.fromsimilary” function using the signed network and soft thresholding power 14. The mean of the connectivity score of a given module is calculated by first taking the row sum of the adjacency matrix as the intra modular connectivity score per gene, and then the average of the intra modular connective score of all genes in the given module was calculated.

#### Functional Enrichment

Functional annotation of the DE analysis was performed by GOrilla using the “Single ranked list of genes” model. For each DE analysis, two ranks are generated using LFCs, one from the most negative to most positive, and vice versa. The software will search for GO terms that are enriched in the top of the list compared to the rest of the list using the mHG statistics. Total of 8 GOrilla analyses were performed for each age group (3 months and 8 months) and Genotype and Plaque, respectively. Bonferroni correction was performed on all Gorilla analyses based on the total number of comparisons (21825 GO terms * 8 ranks = 174600).

Functional annotation of each module was performed by GOrilla using the “Two unranked lists of genes” model. Each module is used as the target list and the total of 36715 genes expressed in our dataset were used as the background set. The software searches for GO terms that are enriched in the target set compared to the background set using the standard Hypergeometric statistics.

To merge similar GO terms into cluster, we first generated a similarity matrix between all significantly enriched GO terms based on the number of the genes overlapping between two GO terms. Next, we performed hierarchical clustering using the complete linkage method, with tree height 3.3 which grouped all significant GO terms into 12 clusters.

#### Logistic regression

Logistic regression was used to predict plaque niche enrichment and cell type enrichment of the *in situ* sequencing data. For plaque niche enrichment, each gene is fitted as a continuous explanatory variable in a single logistic regression, with the proportion of cells in ring 1 over cells in non-ring 1 used as the logit response variable. An odd ratio is calculated by taking the exponential of the coefficient calculated from each regression, which indicates the ratio of the probability of a given gene in ring1 (plaque niche) and the probability in non-ring 1. For cell type enrichment, each gene is used as explanatory variable the same way as the plaque niche enrichment, but the logit is calculated as the log of the proportion of cells expressing at least 1 copy of a given cell type marker over cells that do not express that cell type marker. The p-values from each regression model were corrected using Bonferroni correction.

## References

Ashburner, M. et al. (2000). Gene ontology: Tool for the unification of biology. Nature Genetics. 25, 25–29.

Bankhead, P. et al. (2017). QuPath: Open source software for digital pathology image analysis. Scientific Reports. 7(1), 16878.

Basu, A., Krady, J. K. and Levison, S. W. (2004). Interleukin-1: A master regulator of neuroinflammation. Journal of Neuroscience Research. 78(2), 151–6.

Berge, V., Johnson, E. and Berge, K. E. (2009). Interleukin-1α, interleukin 6 and tumor necrosis factor α increase the synthesis and expression of the functional alternative and terminal complement pathways by human umbilical vein endothelial cells in vitro. APMIS. 104(3), 213–9.

Borowsky, I. W. and Collins, R. C. (1989). Histochemical changes in enzymes of energy metabolism in the dentate gyrus accompany deafferentation and synaptic reorganization. Neuroscience. 33(2), 253–262.

Braak, H. and Braak, E. (1996). Development of Alzheimer-related neurofibrillary changes in the neocortex inversely recapitulates cortical myelogenesis. Acta Neuropathologica. 92(2), 197–201.

van den Brink, S. C. et al. (2017). Single-cell sequencing reveals dissociation-induced gene expression in tissue subpopulations. Nature Methods. 14(10), 935–936.

Carbon, S. et al. (2019). The Gene Ontology Resource: 20 years and still GOing strong. Nucleic Acids Research. 47(D1), 330–338.

Castillo, E. et al. (2017). Comparative profiling of cortical gene expression in Alzheimer’s disease patients and mouse models demonstrates a link between amyloidosis and neuroinflammation. Scientific Reports. 7(1), 17762.

Eden, E. et al. (2009). GOrilla: a tool for discovery and visualization of enriched GO terms in ranked gene lists. BMC bioinformatics. 10(1), 48.

Eikelenboom, P. et al. (1988). Complement activation in amyloid plaques in Alzheimer’s dementia. Virchows Archiv B Cell Pathology Including Molecular Pathology. 56(4), 259–262.

Ettle, B., Schlachetzki, J. C. M. and Winkler, J. (2016). Oligodendroglia and Myelin in Neurodegenerative Diseases: More Than Just Bystanders? Molecular Neurobiology. 53(5), 3046–3062.

Gallardo, G. and Holtzman, D. M. (2017). Antibody therapeutics targeting Aβ and tau. Cold Spring Harbor Perspectives in Medicine. 7(10), a024331.

Gerritsma, J. S. J. et al. (1996). Interleukin-1α enhances the biosynthesis of complement C3 and factor B by human kidney proximal tubular epithelial cells in vitro. Molecular Immunology. 33(10), 847–854.

Glabe, C. G. (2008). Structural classification of toxic amyloid oligomers. Journal of Biological Chemistry. 283(44), 29639–43.

Hansen, D. V., Hanson, J. E. and Sheng, M. (2018). Microglia in Alzheimer’s disease. Journal of Cell Biology. 217(2), 459–472.

Hong, S. et al. (2016). Complement and microglia mediate early synapse loss in Alzheimer mouse models. Science. 352(6286), 712–716.

Huber, W. et al. (2015). Orchestrating high-throughput genomic analysis with Bioconductor. Nature Methods. 12(2), 115–21.

Jansen, I. E. et al. (2019). Genome-wide meta-analysis identifies new loci and functional pathways influencing Alzheimer’s disease risk. Nature Genetics. 51(3), 404–413.

Kametani, F. and Hasegawa, M. (2018). Reconsideration of amyloid hypothesis and tau hypothesis in Alzheimer’s disease. Frontiers in Neuroscience. 12, 25.

Karran, E., Mercken, M. and De Strooper, B. (2011). The amyloid cascade hypothesis for Alzheimer’s disease: an appraisal for the development of therapeutics. Nat Rev Drug Discov. 10(9), 698–712.

Ke, R. et al. (2013) ‘In situ sequencing for RNA analysis in preserved tissue and cells. Nature Methods. 10(9), 857–860.

Keren-Shaul, H. et al. (2017). A Unique Microglia Type Associated with Restricting Development of Alzheimer’s Disease. Cell. 169(7), 1276–1290.

Krasemann, S. et al. (2017). The TREM2-APOE Pathway Drives the Transcriptional Phenotype of Dysfunctional Microglia in Neurodegenerative Diseases. Immunity. 47(3), 566–581.

Kuo, Y. M. et al. (2001). Comparative Analysis of Amyloid-β Chemical Structure and Amyloid Plaque Morphology of Transgenic Mouse and Alzheimer’s Disease Brains. Journal of Biological Chemistry. 276(16), 12991–8.

Lake, B. B. et al. (2017). A comparative strategy for single-nucleus and single-cell transcriptomes confirms accuracy in predicted cell-type expression from nuclear RNA. Scientific Reports. 7(1), 6031.

Legland, D., Arganda-Carreras, I. and Andrey, P. (2016). MorphoLibJ: Integrated library and plugins for mathematical morphology with ImageJ. Bioinformatics. 32(22), 3532–3534.

Liddelow, S. A. et al. (2017). Neurotoxic reactive astrocytes are induced by activated microglia. Nature. 541(7638), 481–487.

Makin, S. (2018). The amyloid hypothesis on trial. Nature. 559(7715), S4–S7.

Maniatis, S. et al. (2019). Spatiotemporal dynamics of molecular pathology in amyotrophic lateral sclerosis. Science. 364, 89–93

Mathys, H. et al. (2019). Single-cell transcriptomic analysis of Alzheimer’s disease. Nature. 570(7761), 332–337.

McGeer, P. L. et al. (1989). Activation of the classical complement pathway in brain tissue of Alzheimer patients. Neuroscience Letters. 107(1-3), 341–346.

McKenzie, A. T. et al. (2018). Brain Cell Type Specific Gene Expression and Co-expression Network Architectures. Scientific Reports. 8(1), 8868.

Morgan, B. P. (2018). Complement in the pathogenesis of Alzheimer’s disease. Seminars in Immunopathology. 40(1), 113–124.

Mucke, L. and Selkoe, D. J. (2012). Neurotoxicity of amyloid β-protein: Synaptic and network dysfunction. Cold Spring Harbor Perspectives in Medicine. 2(7), a006338.

Ono, K., Condron, M. M. and Teplow, D. B. (2009). Structure-neurotoxicity relationships of amyloid beta-protein oligomers. Proceedings of the National Academy of Sciences. 106(35), 14745–50.

Parhizkar, S. et al. (2019). Loss of TREM2 function increases amyloid seeding but reduces plaque-associated ApoE. Nature Neuroscience. 22(2), 191–204.

Qian, X. et al. (2018). A spatial atlas of inhibitory cell types in mouse hippocampus. https://www.biorxiv.org/content/10.1101/431957v1

Robakis, N. K. (2010). Are Aβ and its derivatives causative agents or innocent bystanders in AD? Neurodegenerative Diseases. 7(1-3), 32–7.

Rodriques, S. G. et al. (2019). Slide-seq: A scalable technology for measuring genome-wide expression at high spatial resolution. Science. 363(6434), 1463–1467.

Rothman, S. M. et al. (2018). Human Alzheimer’s disease gene expression signatures and immune profile in APP mouse models: A discrete transcriptomic view of Aβ plaque pathology. Journal of Neuroinflammation. 15(1), 256.

Saito, T. et al. (2014). Single App knock-in mouse models of Alzheimer’s disease. Nature neuroscience. 17(5), 661–663.

Sala Frigerio, C. et al. (2019). The Major Risk Factors for Alzheimer’s Disease: Age, Sex, and Genes Modulate the Microglia Response to Aβ Plaques. Cell Reports. 27(4), 1293–1306.e6.

Salih, D. A. et al. (2018). Genetic variability in response to Aβ deposition influences Alzheimer’s risk. https://www.biorxiv.org/content/10.1101/437657v1

Schindelin, J. et al. (2012). Fiji: An open-source platform for biological-image analysis. Nature Methods. 9(7), 676–82.

Schmidt, R. E. and Gessner, J. E. (2005). Fc receptors and their interaction with complement in autoimmunity. Immunology Letters. 100(1), 56–57.

Shimada, M. et al. (1992). Microvessel and astroglial cell densities in the mouse hippocampus. Journal of anatomy. 180 (Pt 1), 89–95.

Sierksma, A. et al. (2019). Novel Alzheimer risk genes determine the microglia response to amyloid-β but not to TAU pathology. https://www.biorxiv.org/content/10.1101/491902v2

Srinivasan, K. et al. (2019). Alzheimer’s patient brain myeloid cells exhibit enhanced aging and unique transcriptional activation. https://www.biorxiv.org/content/10.1101/610345v1?rss=1

Ståhl, P. L. et al. (2016). Visualization and analysis of gene expression in tissue sections by spatial transcriptomics. Science. 353(6294), 78–82.

Strooper, B. De and Karran, E. (2016). The Cellular Phase of Alzheimer’s Disease. Cell. 64(4), 603–615.

Thielens, N. M. et al. (2017). C1q: A fresh look upon an old molecule. Molecular Immunology. 89, 73–83.

Vickovic, S. et al. (2019). High-density spatial transcriptomics arrays for in situ tissue profiling. https://www.biorxiv.org/content/10.1101/563338v2

Xiong, F., Ge, W. and Ma, C. (2019). Quantitative proteomics reveals distinct composition of amyloid plaques in Alzheimer’s disease. Alzheimer’s and Dementia. 15(3), 429–440.

Yin, C. et al. (2019). ApoE attenuates unresolvable inflammation by complex formation with activated C1q. Nature Medicine. 25(3), 496–506.

Zamanian, J. L. et al. (2012). Genomic Analysis of Reactive Astrogliosis. Journal of Neuroscience. 32(18), 6391–6410.

Zeisel, A. et al. (2015). Cell types in the mouse cortex and hippocampus revealed by single-cell RNA-seq. Science. 347(6226), 1138–42.

Zeisel, A. et al. (2018). Molecular architecture of the mouse nervous system. Cell. 174(4), 999–1014.e22.

Zhang, B. and Horvath, S. (2005). A General Framework for Weighted Gene Co-Expression Network Analysis. Statistical Applications in Genetics and Molecular Biology. 4, Article 17.

Zhang, P. et al. (2019). Senolytic therapy alleviates Aβ-associated oligodendrocyte progenitor cell senescence and cognitive deficits in an Alzheimer’s disease model. Nature Neuroscience. 22(5), 719–728.

