## Supplementary Figures and Tables for "Spatial and temporal transcriptomics reveal microglia-astroglia crosstalk in the amyloid-β plaque cell niche of Alzheimer’s disease"

### Supplementary Information

#### Figure and Table Legends

**Figure S1. Experimental setup and data quality.** (A) Summary of sample size, number of mice, number of ST experiments and total number of tissue domains passing quality control per age and genotype group. (B, C) The distribution of the number of genes or reads per tissue domains. (D) Each dot represents a gene, and total 46454 genes were detected in full database. The scatter plot shows the distribution of total number of reads per gene (x-axis) and the percentage of tissue domains expressing each gene (y-axis). Among 46454 genes, 1970 genes only have one copy; 25665 genes were lowly expressed (LogRead <5); and 40 genes were highly expressed (LogRead > 13) such as *Plp1, Mbp, Sez6,* and *Apoe*. (E) Each dot represents a transcriptomic profile (tissue domain), in total 10327 tissue domains are available. The scatter plot shows the distribution of total number of reads per tissue domain (x-axis) and the percentage of expressing gene per tissue domain (y-axis). (F, G) The same t-SNE plot as Figure 1D, but tissue domains are colored according to gene expression of landmark genes in somatic layers (F) or metabolic genes in dendritic layers (G). As shown in panel F, *Dsp* and *Prox1* were localized in the granule cell layer of Dentate Gyrus (DG_sp), while *Fibcd1* and *Wfs1* were localized in the pyramidal layer of CA1 (CA1_sp). In addition, we also identified the enrichment of *mt-Co1* (cytochrome c oxidase), *Aldoc* (aldolase c), *Hba-a1* (hemoglobin subunit A1), and *Gfap* (glial fibrillary acidic protein) in the dendritic layer of hippocampus in panel G. This observation is in line with previous reports that the dendritic layer serves as the metabolic center of the hippocampus, containing the most microvessels, astroglial (Shimada *et al*., 1992) and highest activity of metabolic enzymes, including *mt-Co1* or *Aldoc* encoding proteins (Borowsky and Collins, 1989).

**Figure S2. Amyloid deposition and astrogliosis** (A and B) Immunofluorescent staining of amyloid plaques (mAb 6E10, white), astrocytes (Gfap, green), neurons (NeuN, red), and nuclei (DAPI, blue) in coronal sections of C57Black6J (A) and *APP*$^{NL-G-F}$ (B) brain at the indicated age. Selective zoom-ins in of neocortex, hippocampus, and thalamus are indicated. Scale bar: 500 µm in full coronal section, 100 µm or 20 µm in the zoom-ins. CX_SSp: primary somatosensory area of cortex, HP_CA1_sr: field CA1, stratum radiatum of hippocampus; TH_GENc: Ganiculate group, ventral thalamus. (C) Regional plots show the expression levels of GFAP protein at the indicated region, age and genotype. (D) Amyloid plaques are visualized with antibody 6E10 (red). Amyloid fibrils are visualized with Thioflavin-S (green) (age of mice is indicated). Notice that Thioflavin S-positive amyloid fibrils are detectable in the 6E10-positive plaques after 6-month of age. Scale bar: 200 µm and 100 µm in the zoom-ins.

**Figure S3. Gene expression alterations around amyloid plaques.** (A and B) The 50 most significantly changed genes sorted by FDR at 3- (A) and 18-month (B) of age. It shows the genes sorted by LFC according to plaque-model. The heatmap squares (top 2 rows) show the LFCs of the genotype effect and the plaque effect at 3- (left bottom triangle) and 18- (right top triangle) month of age, and the -log10 calculated from GWAS AD risk p-values (3rd row). ARM/DAM and A1 markers are highlighted in triangles (4th row). Average expression profile in single-cell sequencing data from wild-type (CD-1) mice by Zeisel et al. for each cell type is plotted as heatmap squares (row 5-9), and Zeisel et al. defined (color background) / predicted cell type (grey background) genes are highlighted in the last row. Notice that both *Plp1* and *Il1a* are among the top 10 up-regulated genes, while many interneuronal genes (eg. *Bex2, Scg2, Napl15, Resp18, Gprasp2*) are down-regulated in the plaque-axis at 3-months of age (A). The top up-regulated genes in the plaque-axis at the 18-months of age are mostly microglial genes, while *Plp1* is down-regulated among the top significantly changed 50 genes at 18-months of age (B).

**Figure S4. Gene ontology analysis and 12 co-expression networks defined by WGCNA.** (A and B) Summary of the dynamic alteration of gene expression in the plaque niche during disease progression by GO analysis and co-expression modules. Panel A shows the significantly changed 13 super GO-categories, while panel B shows the alteration of 12 WGCNA-identified modules in both plaque- and genotype-axis in the 3-months (blue) and 18-months (purple) of age. The plaque-disease interaction plot takes the average of LFC of genes in each GO-category (A) and in each module (B) of the indicated axis and age. (C) Summary of the cellular composition (column 2-9) and GWAS enrichment (1$^{st}$ column) per co-expression module. The first column shows the significant odds ratio of each module associated with top 255 Marioni's GWAS AD hits (*pval* < 0.0005). The 2$^{nd}$ to 9$^{th}$ columns show the significant odds ratio of each module associated with the indicated cell type markers derived from single cell sequencing data from wild-type (CD-1) mice by Zeisel et al.. The last column summarizes the significant GO category for each module.

**Figure S5. Cellular distribution revealed by *in situ* sequencing.** (A and B) Amyloid plaque (6E10, white) and DAPI staining (A) was coupled with *in situ* sequencing (B) in the AD mice at 18-month of age. (B) Selective cell type markers shown in the same coronal section (neuron in red: *Syp, App, Neurod6*; microglia in yellow: *C1qa, Arpc1b, Csf1r, Ctsz, Fcgr3, Itgam, Lyz, Vsir*; astrocyte in green: *Cd63, Clu, Cst3, Gfap, Prdx6, Serpina3n, Slc1a3*; oligodendrocyte in blue: *Plp1, Laptm5*). Scale bar: 800 µm of panel A, 1000 µm of panel B (C) Cell population as detected with different markers in WT brain and AD brain. Notice that there is no major shifts in cell composition between $APP^{NL\text{-}G\text{-}F}$ and the control with around 41% *Syp*-positive neurons, 23% *Plp1*-positive oligodendrocytes, 10% *Slc1a3*-positive astrocyte, and 1.5% positive *Itgam*-positive microglia. Since the expression of Itgam is low, this has likely lead to an under-estimation of the microglia population. We therefore included 2 additional microglia markers from the PIGs series (*Csf1r* and *Cx3cr1*). Again, we do not see major overall shifts in the number of microglial cells with 13.2% *Csf1r*-positive microglia and 9.8% *Cx3cr1*-positive microglia. While the general astroglia marker (*Slc1a3*) did not change, the percentage of *Gfap*-positive cells increased dramatically in the AD mice (26.6%) compared to WT mice (7.3%), confirming the strong astrogliosis in the former. (D) Cell population in the rings around the amyloid plaques as indicated, and normalized to ring1. Notice that the percentage of *Gfap*-positive cells was high in ring 1 and ring 2 and decreased away from the plaque, while the percentage of *Plp1*-positive oligodendrocytes was low in ring 1 but increased by distance to the plaque. The *Slc1a3*-positive astrocytes are somewhat enriched in the second ring, while *Itgam*-positive (or *Csf1r*-positive, *Cx3cr1*-positive) microglia are enriched in the first ring. The ISS result confirms the depletion of *Plp1*-positive oligodendrocytes in the plaque niche at 18 months, as we observed also with the ST approach.

**Figure S6. Cluster dendrogram and module assignment of PIGs in WT and Q4 using WGCNA.** Genes were clustered based on a dissimilarity measure (1-TOM). The branches correspond to modules of highly interconnected groups of genes. Colors in the horizontal bar represent the modules.

**Table S1. Sample quality.** For each ST library, we report the age, hemisphere, RIN value, PCR cycle, bregma, and the ID according to the Allen Brain atlas of each coronal section of each mouse.

**Table S2. List of enriched GO term in functions of plaque and genotype at 3- and 18-months of age.** Detailled information of GO terms within each of the 13 super categories. The significance (P-Bonferroni), the enrichment score, and the leading gene of each GO term per age (M03 or M18), and per direction of the plaque-disease interaction plot (GD: down-regulation in the genotype/disease model; GU: up-regulation in the genotype/disease model; PD: down-regulation in the plaque-model; PU: up-regulation in the plaque-model) are included.

**Table S3. List of genes of each co-expression network defined by WGCNA.**

**Table S4. The significantly enriched GO term in each co-expression network**. GO number, description, enrichment score, and significance (p-value and adjusted FDR) per GO term for each WGCNA module is included.

Figure S1

A

| | WT: C57Black6J | | | | AD: APP$^{NL-G-F}$ | | | |
|---|---|---|---|---|---|---|---|---|
| Age (month) | 3 | 6 | 12 | 18 | 3 | 6 | 12 | 18 |
| (mouse) | 2 | 1 | 1 | 2 | 2 | 1 | 1 | 2 |
| (section) | 4 | 1 | 1 | 4 | 4 | 1 | 1 | 4 |
| Spot | 2277 | 537 | 447 | 2057 | 2015 | 478 | 488 | 2028 |

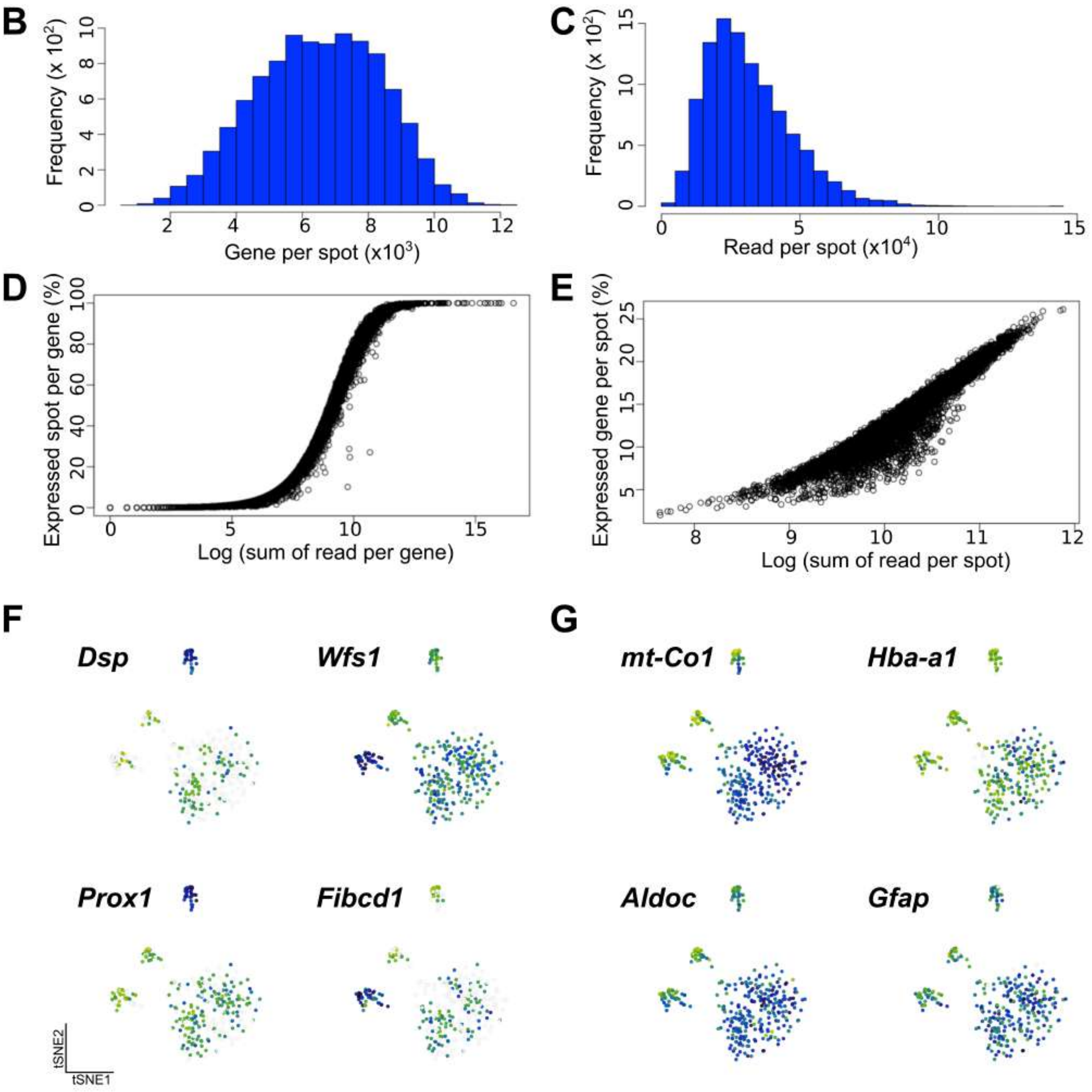

Figure S2

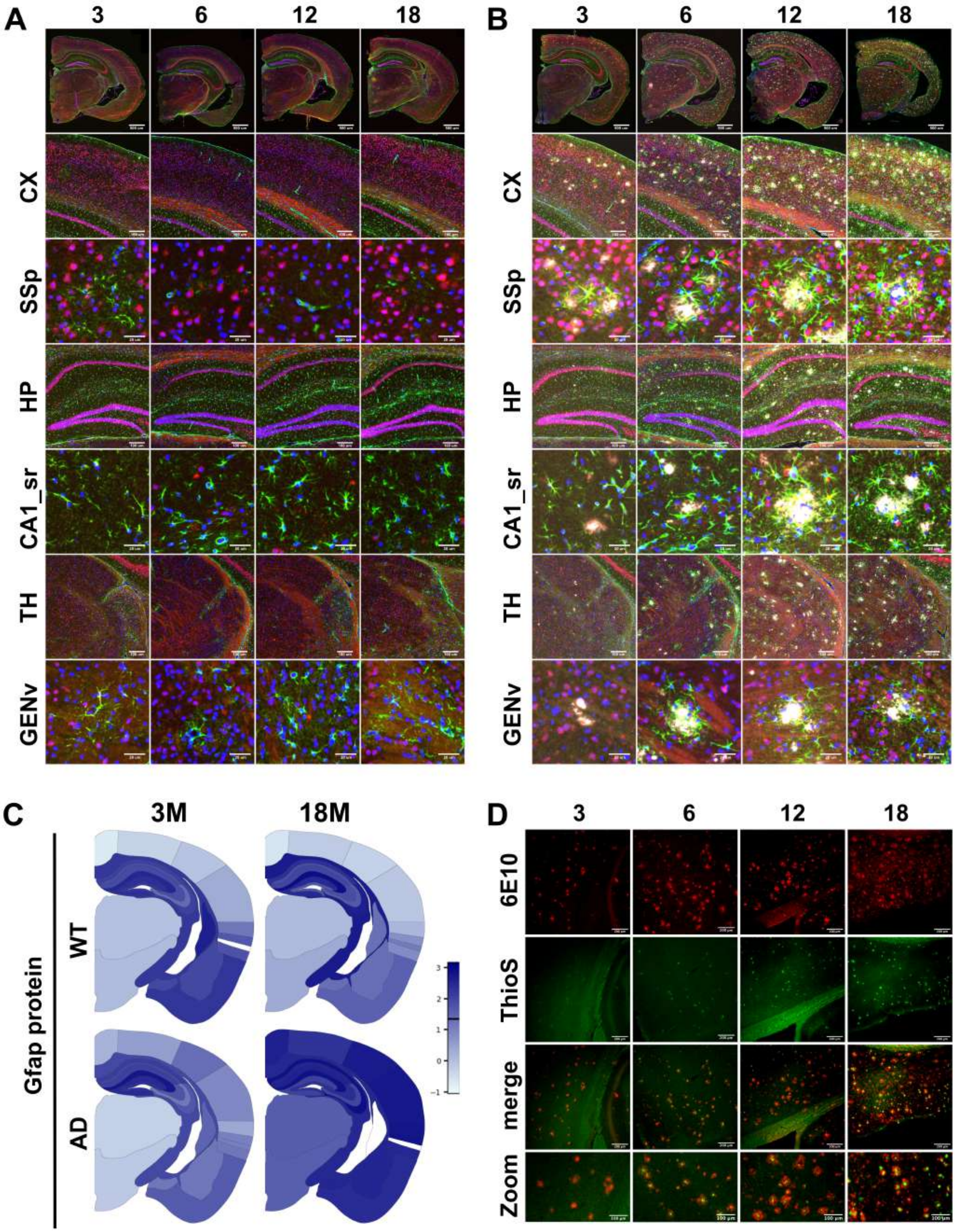

A
3
6
12
18
CX
SSp
HP
CA1_sr
TH
GENv
B
3
6
12
18
CX
SSp
HP
CA1_sr
TH
GENv
C
3M
18M
Gfap protein
WT
AD
3
2
1
0
-1
D
3
6
12
18
6E10
ThioS
merge
Zoom

Figure S3

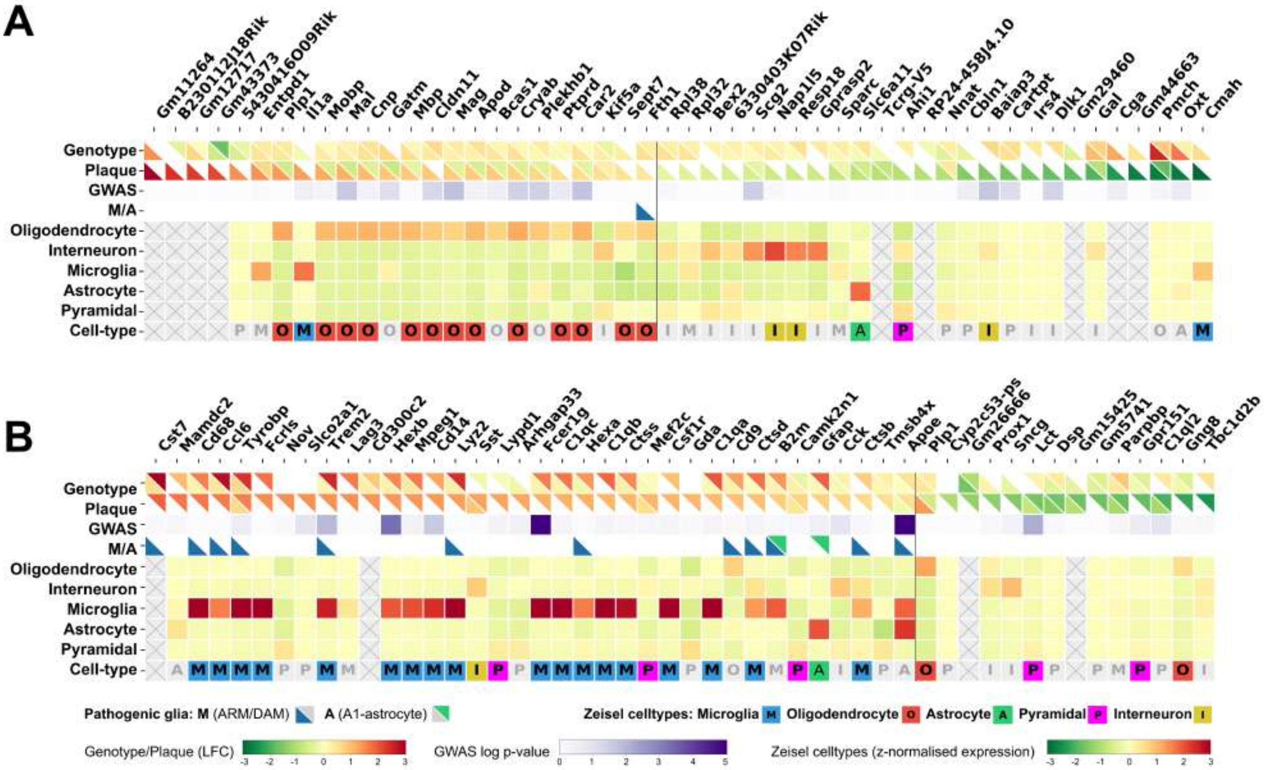
A
B
Genotype
Plaque
GWAS
M/A
Oligodendrocyte
Interneuron
Microglia
Astrocyte
Pyramidal
Cell-type
Pathogenic glia: M (ARM/DAM) A (A1-astrocyte)
Zeisel celltypes: Microglia Oligodendrocyte Astrocyte Pyramidal Interneuron
Genotype/Plaque (LFC)
GWAS log p-value
Zeisel celltypes (z-normalised expression)

Figure S4

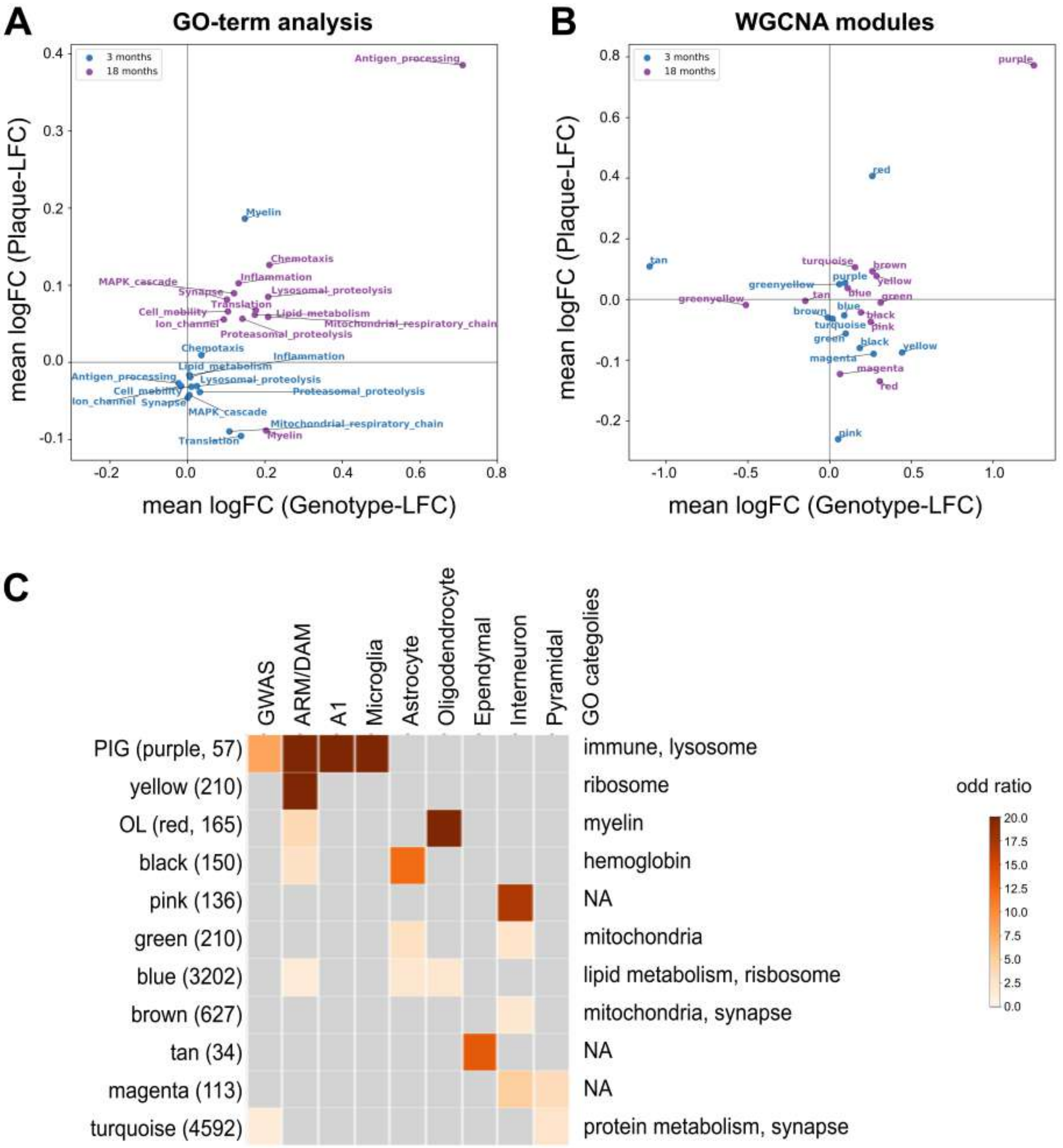
A
GO-term analysis
3 months
18 months
mean logFC (Plaque-LFC)
mean logFC (Genotype-LFC)
Antigen_processing
Myelin
Chemotaxis
Inflammation
MAPK_cascade
Synapse
Lysosomal_proteolysis
Translation
Cell_mobility
Lipid_metabolism
Ion_channel
Mitochondrial_respiratory_chain
Proteasomal_proteolysis
B
WGCNA modules
purple
red
tan
turquoise
brown
greenyellow
yellow
blue
green
black
pink
magenta
C
GWAS
ARM/DAM
A1
Microglia
Astrocyte
Oligodendrocyte
Ependymal
Interneuron
Pyramidal
GO categolies
PIG (purple, 57)
yellow (210)
OL (red, 165)
black (150)
pink (136)
green (210)
blue (3202)
brown (627)
tan (34)
magenta (113)
turquoise (4592)
immune, lysosome
ribosome
myelin
hemoglobin
NA
mitochondria
lipid metabolism, risbosome
mitochondria, synapse
NA
NA
protein metabolism, synapse
odd ratio

Figure S5

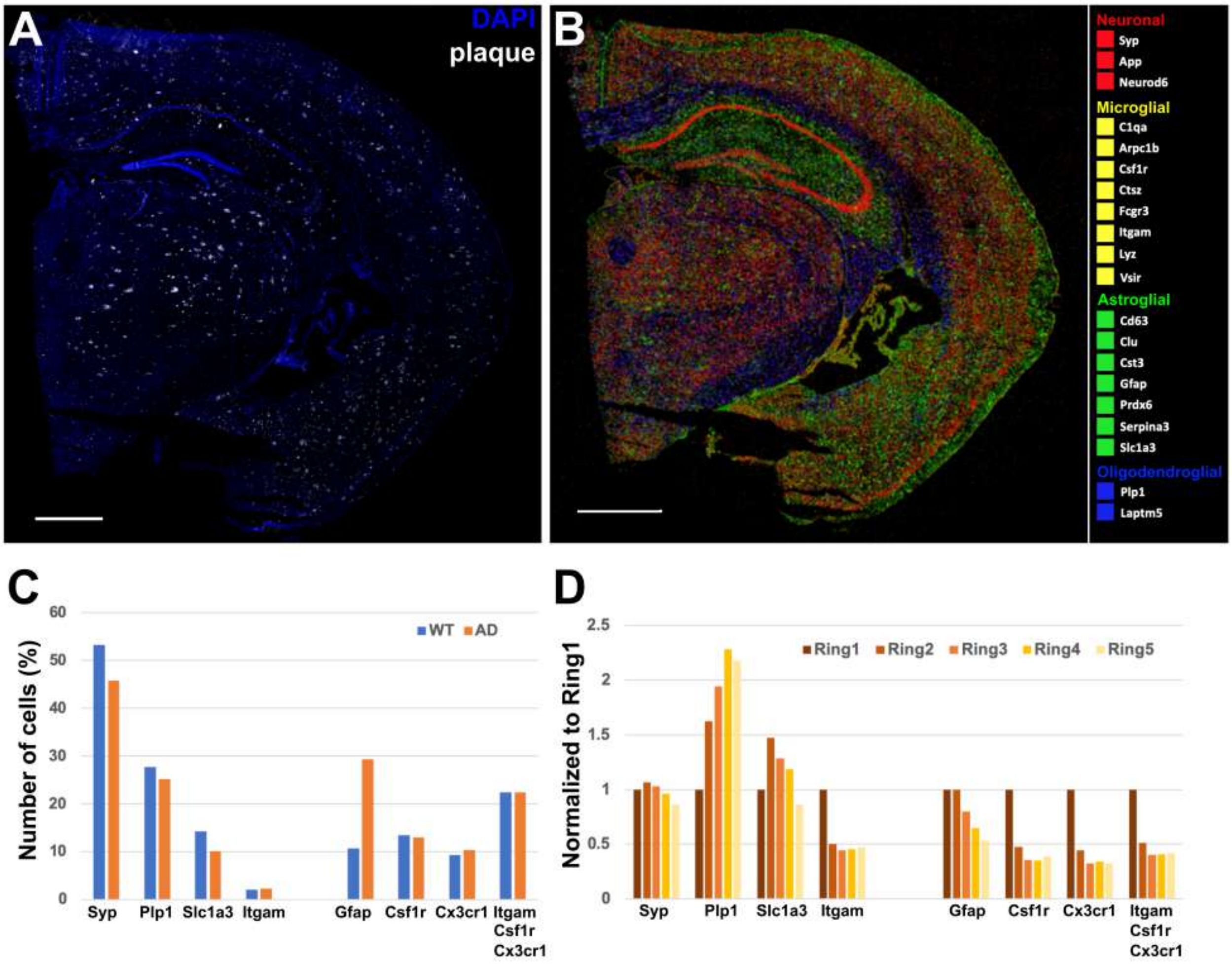
A
DAPI
plaque
B
Neuronal
Syp
App
Neurod6
Microglial
C1qa
Arpc1b
Csf1r
Ctsz
Fcgr3
Itgam
Lyz
Vsir
Astroglial
Cd63
Clu
Cst3
Gfap
Prdx6
Serpina3
Slc1a3
Oligodendroglial
Plp1
Laptm5
C
Number of cells (%)
WT
AD
0
10
20
30
40
50
60
Syp
Plp1
Slc1a3
Itgam
Gfap
Csf1r
Cx3cr1
Itgam Csf1r Cx3cr1
D
Normalized to Ring1
Ring1
Ring2
Ring3
Ring4
Ring5
0
0.5
1
1.5
2
2.5
Syp
Plp1
Slc1a3
Itgam
Gfap
Csf1r
Cx3cr1
Itgam Csf1r Cx3cr1

Figure S6

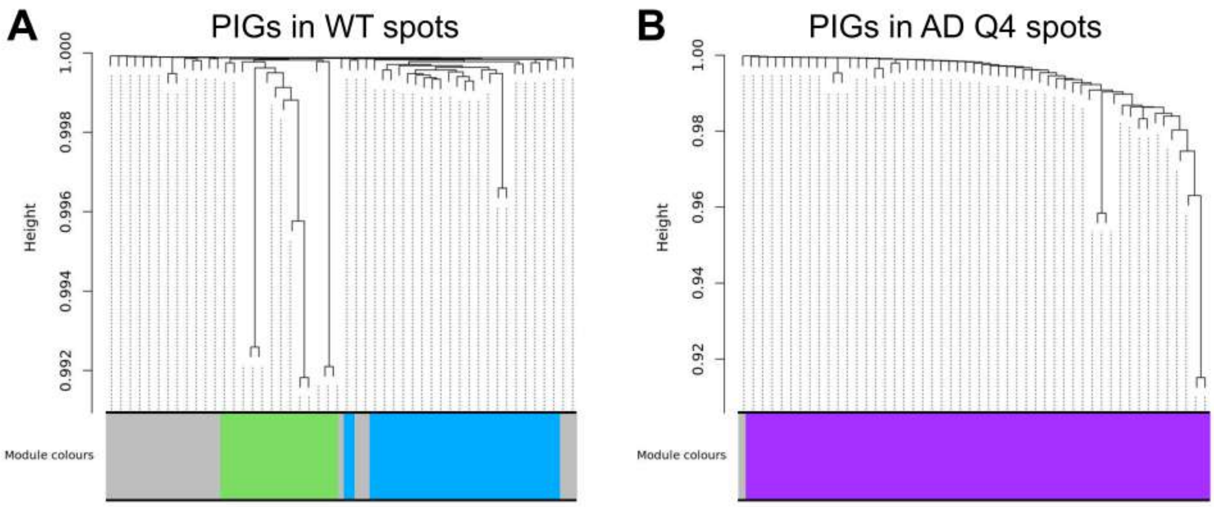
A
PIGs in WT spots
Height
1.000
0.998
0.996
0.994
0.992
Module colours
B
PIGs in AD Q4 spots
Height
1.00
0.98
0.96
0.94
0.92
Module colours

Table S1

| Sample id | Sample name | Genotype | Hemisphere | Birthday | Brain frozen | Age (day) | Age (month) | Bregma | Interaural | Slide | RIN | PCR cycle | Library Index | Ave_length _cDNA (bp) | Allen Atlas ID |
|---|---|---|---|---|---|---|---|---|---|---|---|---|---|---|---|
| N02_D1 | APPNLGF14 | APPNLGF | R | 22/06/2016 | 06/10/2016 | 107 | 3 | -2.18 | 1.74 | Lot#10001; ID#034 | 9.3 | 8 | 9- CAAGCAGAAGACGGCATACGAGATCTGATCGTGACTGGAGTTC | 548 | 100960092 |
| N03_D2 | APPNLGF15 | APPNLGF | L | 22/06/2016 | 06/10/2016 | 107 | 3 | -2.06 | 1.74 | Lot#10001; ID#034 | 9.3 | 8 | 10- CAAGCAGAAGACGGCATACGAGATAAGCTAGTGACTGGAGTTC | 541 | 100960084 |
| N02_C1 | APPNLGF14 | APPNLGF | R | 22/06/2016 | 06/10/2016 | 107 | 3 | -2.18 | 1.74 | Lot#10001; ID#034 | 9.3 | 8 | 7- CAAGCAGAAGACGGCATACGAGATGATCTGGTGACTGGAGTTC | 522 | 100960092 |
| N03_C2 | APPNLGF15 | APPNLGF | L | 22/06/2016 | 06/10/2016 | 107 | 3 | -2.06 | 1.74 | Lot#10001; ID#034 | 9.3 | 8 | 8- CAAGCAGAAGACGGCATACGAGATTCAAGTGTGACTGGAGTTC | 509 | 100960084 |
| B02_D1 | C57Black12 | C57Black6 | R | 22/06/2016 | 06/10/2016 | 107 | 3 | -2.18 | 1.62 | Lot#10001; ID#027 | 9.45 | 8 | 3- CAAGCAGAAGACGGCATACGAGATGCCTAAGTGACTGGAGTTC | 441 | 100960092 |
| B02_E1 | C57Black12 | C57Black6 | R | 22/06/2016 | 06/10/2016 | 107 | 3 | -2.18 | 1.62 | Lot#10001; ID#027 | 9.45 | 9 | 5- CAAGCAGAAGACGGCATACGAGATCACTGTGTGACTGGAGTTC | 491 | 100960092 |
| B03_C2 | C57Black13 | C57Black6 | L | 22/06/2016 | 06/10/2016 | 107 | 3 | -2.18 | 1.62 | Lot#10001; ID#027 | 9.25 | 8 | 2- CAAGCAGAAGACGGCATACGAGATACATCGGTGACTGGAGTTC | 489 | 100960092 |
| B03_D2 | C57Black13 | C57Black6 | L | 22/06/2016 | 06/10/2016 | 107 | 3 | -2.18 | 1.62 | Lot#10001; ID#027 | 9.25 | 9 | 4- CAAGCAGAAGACGGCATACGAGATTGGTCAGTGACTGGAGTTC | 535 | 100960092 |
| N04_D1 | APPNLGF20 | APPNLGF | R | 01/08/2015 | 09/02/2017 | 559 | 18 | -2.30 | 1.50 | Lot#10001; ID#062 | 9.25 | 9 | 3- CAAGCAGAAGACGGCATACGAGATGCCTAAGTGACTGGAGTTC | 517 | 100960088 |
| N04_E1 | APPNLGF20 | APPNLGF | R | 01/08/2015 | 09/02/2017 | 559 | 18 | -2.30 | 1.50 | Lot#10001; ID#062 | 9.25 | 10 | 5- CAAGCAGAAGACGGCATACGAGATCACTGTGTGACTGGAGTTC | 504 | 100960088 |
| N05_C2 | APPNLGF21 | APPNLGF | L | 02/08/2015 | 09/02/2017 | 558 | 18 | -2.18 | 1.62 | Lot#10001; ID#062 | 9 | 10 | 2- CAAGCAGAAGACGGCATACGAGATACATCGGTGACTGGAGTTC | 495 | 100960092 |
| N05_D2 | APPNLGF21 | APPNLGF | L | 02/08/2015 | 09/02/2017 | 558 | 18 | -2.18 | 1.62 | Lot#10001; ID#062 | 9 | 9 | 4- CAAGCAGAAGACGGCATACGAGATTGGTCAGTGACTGGAGTTC | 504 | 100960092 |
| B04_D1 | C57Black32 | C57Black6 | R | 28/10/2015 | 25/04/2017 | 546 | 18 | -2.18 | 1.62 | Lot#10001; ID#063 | 8.8 | 10 | 9- CAAGCAGAAGACGGCATACGAGATCTGATCGTGACTGGAGTTC | 435 | 100960092 |
| B04_E1 | C57Black32 | C57Black6 | R | 28/10/2015 | 25/04/2017 | 546 | 18 | -2.06 | 1.74 | Lot#10001; ID#063 | 8.8 | 10 | 11- CAAGCAGAAGACGGCATACGAGATGTAGCCGTGACTGGAGTTC | 447 | 100960092 |
| B05_D2 | C57Black31 | C57Black6 | L | 28/10/2015 | 25/04/2017 | 546 | 18 | -2.18 | 1.62 | Lot#10001; ID#063 | 8.4 | 10 | 10- CAAGCAGAAGACGGCATACGAGATAAGCTAGTGACTGGAGTTC | 450 | 100960092 |
| B05_E2 | C57Black31 | C57Black6 | L | 28/10/2015 | 25/04/2017 | 546 | 18 | -2.06 | 1.74 | Lot#10001; ID#063 | 8.4 | 10 | 12- CAAGCAGAAGACGGCATACGAGATTACAAGGTGACTGGAGTTC | 444 | 100960092 |
| B06_E1 | C57Black22 | C57Black6 | R | 10/02/2016 | 16/02/2017 | 373 | 12 | -2.18 | 1.62 | Lot#10001; ID#064 | 9.05 | 10 | 3- CAAGCAGAAGACGGCATACGAGATGCCTAAGTGACTGGAGTTC | 432 | 100960092 |
| N06_D2 | APPNLGF25 | APPNLGF | R | 27/02/2016 | 23/02/2017 | 363 | 12 | -2.06 | 1.74 | Lot#10001; ID#064 | 8.55 | 10 | 5- CAAGCAGAAGACGGCATACGAGATCACTGTGTGACTGGAGTTC | 433 | 100960092 |
| B07_C2 | C57Black53 | C57Black6 | R | 19/12/2016 | 19/6/2017 | 183 | 6 | -2.3 | 1.50 | Lot#10001; ID#065 | 9.35 | 11 | 10- CAAGCAGAAGACGGCATACGAGATAAGCTAGTGACTGGAGTTC | 406 | 100960088 |
| N07_C1 | APPNLGF49 | APPNLGF | R | 16/12/2016 | 19/6/2017 | 186 | 6 | -2.3 | 1.50 | Lot#10001; ID#065 | 9.6 | 10 | 7- CAAGCAGAAGACGGCATACGAGATGATCTGGTGACTGGAGTTC | 433 | 100960088 |

Table S2

| | Cluster | Description | Selected cluster | Name selected cluster | Min_PBonferroni |
|---|---|---|---|---|---|
| GO:0070372 | 12 | regulation of ERK1 and ERK2 cascade | 13 | MAPK_cascade | 0.007246616 |
| GO:0043410 | 12 | positive regulation of MAPK cascade | 13 | MAPK_cascade | 2.46876E-05 |
| GO:0043408 | 12 | regulation of MAPK cascade | 13 | MAPK_cascade | 0.000539197 |
| GO:0042327 | 12 | positive regulation of phosphorylation | 13 | MAPK_cascade | 1.20368E-07 |
| GO:0070374 | 12 | positive regulation of ERK1 and ERK2 cascade | 13 | MAPK_cascade | 0.026652808 |
| GO:0001934 | 12 | positive regulation of protein phosphorylation | 13 | MAPK_cascade | 2.38279E-07 |
| GO:0022604 | 12 | regulation of cell morphogenesis | 12 | Cell_mobility | 1.3265E-05 |
| GO:0033674 | 12 | positive regulation of kinase activity | 12 | Cell_mobility | 0.030214704 |
| GO:2000147 | 12 | positive regulation of cell motility | 12 | Cell_mobility | 0.000948201 |
| GO:0019900 | 12 | kinase binding | 12 | Cell_mobility | 0.027266928 |
| GO:2000145 | 12 | regulation of cell motility | 12 | Cell_mobility | 6.06751E-08 |
| GO:0032880 | 12 | regulation of protein localization | 12 | Cell_mobility | 0.0399178 |
| GO:0030334 | 12 | regulation of cell migration | 12 | Cell_mobility | 1.67041E-08 |
| GO:0030335 | 12 | positive regulation of cell migration | 12 | Cell_mobility | 0.000524458 |
| GO:0031344 | 12 | regulation of cell projection organization | 12 | Cell_mobility | 0.000126509 |
| GO:0031346 | 12 | positive regulation of cell projection organization | 12 | Cell_mobility | 0.007762477 |
| GO:0016477 | 12 | cell migration | 12 | Cell_mobility | 1.04155E-05 |
| GO:0060341 | 12 | regulation of cellular localization | 12 | Cell_mobility | 0.000778704 |
| GO:0051272 | 12 | positive regulation of cellular component movement | 12 | Cell_mobility | 0.001731818 |
| GO:0051270 | 12 | regulation of cellular component movement | 12 | Cell_mobility | 1.41248E-08 |
| GO:0051046 | 12 | regulation of secretion | 12 | Cell_mobility | 4.31112E-05 |
| GO:0050769 | 12 | positive regulation of neurogenesis | 12 | Cell_mobility | 0.003144294 |
| GO:0050767 | 12 | regulation of neurogenesis | 12 | Cell_mobility | 5.35513E-07 |
| GO:0050678 | 12 | regulation of epithelial cell proliferation | 12 | Cell_mobility | 0.046059 |
| GO:0048870 | 12 | cell motility | 12 | Cell_mobility | 6.68163E-06 |
| GO:0045664 | 12 | regulation of neuron differentiation | 12 | Cell_mobility | 3.91809E-05 |
| GO:0008284 | 12 | positive regulation of cell proliferation | 12 | Cell_mobility | 9.13811E-08 |
| GO:0008285 | 12 | negative regulation of cell proliferation | 12 | Cell_mobility | 8.90474E-08 |
| GO:0010769 | 12 | regulation of cell morphogenesis involved in differentiation | 12 | Cell_mobility | 0.000735716 |
| GO:0040017 | 12 | positive regulation of locomotion | 12 | Cell_mobility | 0.0030706 |
| GO:0040012 | 12 | regulation of locomotion | 12 | Cell_mobility | 4.69188E-07 |
| GO:0120035 | 12 | regulation of plasma membrane bounded cell projection organization | 12 | Cell_mobility | 0.000183008 |
| GO:0040011 | 12 | locomotion | 12 | Cell_mobility | 3.36538E-05 |
| GO:0010975 | 12 | regulation of neuron projection development | 12 | Cell_mobility | 0.006423695 |
| GO:0045596 | 12 | negative regulation of cell differentiation | 12 | Cell_mobility | 0.005748163 |
| GO:0010720 | 12 | positive regulation of cell development | 12 | Cell_mobility | 0.0010047 |
| GO:0097530 | 11 | granulocyte migration | 11 | Chemotaxis | 9.23636E-05 |
| GO:0034341 | 11 | response to interferon-gamma | 11 | Chemotaxis | 5.82186E-10 |
| GO:0060326 | 11 | cell chemotaxis | 11 | Chemotaxis | 0.001744101 |
| GO:0042330 | 11 | taxis | 11 | Chemotaxis | 0.006927274 |
| GO:0071346 | 11 | cellular response to interferon-gamma | 11 | Chemotaxis | 2.82495E-06 |
| GO:0071621 | 11 | granulocyte chemotaxis | 11 | Chemotaxis | 1.67041E-05 |
| GO:1990266 | 11 | neutrophil migration | 11 | Chemotaxis | 2.27224E-05 |
| GO:0008009 | 11 | chemokine activity | 11 | Chemotaxis | 0.0276354 |
| GO:0048247 | 11 | lymphocyte chemotaxis | 11 | Chemotaxis | 0.003181142 |
| GO:0030595 | 11 | leukocyte chemotaxis | 11 | Chemotaxis | 5.28143E-06 |
| GO:0030593 | 11 | neutrophil chemotaxis | 11 | Chemotaxis | 5.29371E-06 |
| GO:0050900 | 11 | leukocyte migration | 11 | Chemotaxis | 5.42882E-07 |
| GO:0006935 | 11 | chemotaxis | 11 | Chemotaxis | 0.006006094 |
| GO:0097529 | 11 | myeloid leukocyte migration | 11 | Chemotaxis | 8.21693E-05 |
| GO:0072676 | 11 | lymphocyte migration | 11 | Chemotaxis | 0.000443395 |
| GO:0070665 | 10 | positive regulation of leukocyte proliferation | 10 | Inflammation | 0.031074472 |
| GO:0071345 | 10 | cellular response to cytokine stimulus | 10 | Inflammation | 0.000652195 |
| GO:0071219 | 10 | cellular response to molecule of bacterial origin | 10 | Inflammation | 0.001163143 |
| GO:0002521 | 10 | leukocyte differentiation | 10 | Inflammation | 0.006878144 |
| GO:0070663 | 10 | regulation of leukocyte proliferation | 10 | Inflammation | 1.62128E-05 |
| GO:0071216 | 10 | cellular response to biotic stimulus | 10 | Inflammation | 0.007602806 |
| GO:0002237 | 10 | response to molecule of bacterial origin | 10 | Inflammation | 0.005723598 |
| GO:1903706 | 10 | regulation of hemopoiesis | 10 | Inflammation | 6.52195E-08 |
| GO:0002685 | 10 | regulation of leukocyte migration | 10 | Inflammation | 0.003205706 |
| GO:0002250 | 10 | adaptive immune response | 10 | Inflammation | 0.002603869 |
| GO:0002687 | 10 | positive regulation of leukocyte migration | 10 | Inflammation | 0.003967215 |
| GO:0002274 | 10 | myeloid leukocyte activation | 10 | Inflammation | 0.000507263 |
| GO:0001819 | 10 | positive regulation of cytokine production | 10 | Inflammation | 1.36335E-06 |
| GO:0001818 | 10 | negative regulation of cytokine production | 10 | Inflammation | 4.82698E-07 |
| GO:0001817 | 10 | regulation of cytokine production | 10 | Inflammation | 2.88636E-09 |
| GO:0001775 | 10 | cell activation | 10 | Inflammation | 2.28453E-07 |
| GO:0002695 | 10 | negative regulation of leukocyte activation | 10 | Inflammation | 0.020020312 |
| GO:0002697 | 10 | regulation of immune effector process | 10 | Inflammation | 0.000476557 |
| GO:0001664 | 10 | G protein-coupled receptor binding | 10 | Inflammation | 0.00859768 |
| GO:1901701 | 10 | cellular response to oxygen-containing compound | 10 | Inflammation | 0.005109478 |
| GO:0002683 | 10 | negative regulation of immune system process | 10 | Inflammation | 0.001240522 |
| GO:1903037 | 10 | regulation of leukocyte cell-cell adhesion | 10 | Inflammation | 0.014616056 |
| GO:0002699 | 10 | positive regulation of immune effector process | 10 | Inflammation | 0.009690814 |
| GO:0002703 | 10 | regulation of leukocyte mediated immunity | 10 | Inflammation | 0.000598153 |
| GO:0002819 | 10 | regulation of adaptive immune response | 10 | Inflammation | 0.000234594 |
| GO:0002821 | 10 | positive regulation of adaptive immune response | 10 | Inflammation | 0.003365378 |
| GO:0002822 | 10 | regulation of adaptive immune response based on somatic recombination of immune receptors built from immunoglobulin superfamily domains | 10 | Inflammation | 0.02640716 |
| GO:1903555 | 10 | regulation of tumor necrosis factor superfamily cytokine production | 10 | Inflammation | 3.9795E-08 |
| GO:1903557 | 10 | positive regulation of tumor necrosis factor superfamily cytokine production | 10 | Inflammation | 0.001694971 |

| | | | | | |
|---|---|---|---|---|---|
| GO:1903708 | 10 | positive regulation of hemopoiesis | 10 | Inflammation | 0.008364314 |
| GO:1902105 | 10 | regulation of leukocyte differentiation | 10 | Inflammation | 0.00013265 |
| GO:0030155 | 10 | regulation of cell adhesion | 10 | Inflammation | 2.40735E-06 |
| GO:0032760 | 10 | positive regulation of tumor necrosis factor production | 10 | Inflammation | 0.001510735 |
| GO:0050727 | 10 | regulation of inflammatory response | 10 | Inflammation | 0.001707254 |
| GO:0050670 | 10 | regulation of lymphocyte proliferation | 10 | Inflammation | 6.28859E-05 |
| GO:0033993 | 10 | response to lipid | 10 | Inflammation | 0.003488202 |
| GO:0006954 | 10 | inflammatory response | 10 | Inflammation | 4.10232E-07 |
| GO:0031347 | 10 | regulation of defense response | 10 | Inflammation | 9.43288E-06 |
| GO:0045637 | 10 | regulation of myeloid cell differentiation | 10 | Inflammation | 1.60899E-05 |
| GO:0031349 | 10 | positive regulation of defense response | 10 | Inflammation | 0.000759052 |
| GO:0045321 | 10 | leukocyte activation | 10 | Inflammation | 2.34594E-06 |
| GO:0007162 | 10 | negative regulation of cell adhesion | 10 | Inflammation | 0.008659092 |
| GO:0032101 | 10 | regulation of response to external stimulus | 10 | Inflammation | 3.32853E-11 |
| GO:0043900 | 10 | regulation of multi-organism process | 10 | Inflammation | 0.024810448 |
| GO:0032102 | 10 | negative regulation of response to external stimulus | 10 | Inflammation | 0.001756383 |
| GO:0009615 | 10 | response to virus | 10 | Inflammation | 0.021862672 |
| GO:0032103 | 10 | positive regulation of response to external stimulus | 10 | Inflammation | 5.33056E-06 |
| GO:0032675 | 10 | regulation of interleukin-6 production | 10 | Inflammation | 0.015107352 |
| GO:0042129 | 10 | regulation of T cell proliferation | 10 | Inflammation | 0.007025533 |
| GO:0032680 | 10 | regulation of tumor necrosis factor production | 10 | Inflammation | 3.21799E-08 |
| GO:0032944 | 10 | regulation of mononuclear cell proliferation | 10 | Inflammation | 8.20464E-06 |
| GO:0019221 | 10 | cytokine-mediated signaling pathway | 10 | Inflammation | 6.54652E-06 |
| GO:0022407 | 10 | regulation of cell-cell adhesion | 10 | Inflammation | 0.000546567 |
| GO:0050863 | 10 | regulation of T cell activation | 10 | Inflammation | 0.000857312 |
| GO:0034097 | 10 | response to cytokine | 10 | Inflammation | 4.12689E-07 |
| GO:0051607 | 10 | defense response to virus | 10 | Inflammation | 0.003291683 |
| GO:0050870 | 10 | positive regulation of T cell activation | 10 | Inflammation | 0.040163448 |
| GO:0050921 | 10 | positive regulation of chemotaxis | 10 | Inflammation | 0.03868956 |
| GO:0050866 | 10 | negative regulation of cell activation | 10 | Inflammation | 0.001952902 |
| GO:0022409 | 10 | positive regulation of cell-cell adhesion | 10 | Inflammation | 0.036478728 |
| GO:0030545 | 10 | receptor regulator activity | 10 | Inflammation | 0.017318184 |
| GO:0009897 | 9 | external side of plasma membrane | 10 | Inflammation | 1.49845E-06 |
| GO:0051251 | 9 | positive regulation of lymphocyte activation | 10 | Inflammation | 0.017318184 |
| GO:0051249 | 9 | regulation of lymphocyte activation | 10 | Inflammation | 0.000491296 |
| GO:0002252 | 9 | immune effector process | 10 | Inflammation | 2.66528E-10 |
| GO:0045087 | 9 | innate immune response | 10 | Inflammation | 1.91605E-06 |
| GO:0002253 | 9 | activation of immune response | 10 | Inflammation | 0.004335687 |
| GO:0002694 | 9 | regulation of leukocyte activation | 10 | Inflammation | 6.49739E-08 |
| GO:0043207 | 9 | response to external biotic stimulus | 10 | Inflammation | 1.08822E-08 |
| GO:0009617 | 9 | response to bacterium | 10 | Inflammation | 2.53017E-06 |
| GO:0050867 | 9 | positive regulation of cell activation | 10 | Inflammation | 2.35822E-05 |
| GO:0098542 | 9 | defense response to other organism | 10 | Inflammation | 1.1828E-06 |
| GO:0050865 | 9 | regulation of cell activation | 10 | Inflammation | 3.23027E-09 |
| GO:0098552 | 9 | side of membrane | 10 | Inflammation | 5.99381E-08 |
| GO:0061024 | 9 | membrane organization | 10 | Inflammation | 0.009076694 |
| GO:0006955 | 9 | immune response | 10 | Inflammation | 7.82389E-18 |
| GO:0006952 | 9 | defense response | 10 | Inflammation | 8.1678E-11 |
| GO:0050778 | 9 | positive regulation of immune response | 10 | Inflammation | 1.48617E-08 |
| GO:0002696 | 9 | positive regulation of leukocyte activation | 10 | Inflammation | 5.88327E-05 |
| GO:0051704 | 9 | multi-organism process | 10 | Inflammation | 0.00010612 |
| GO:0051707 | 9 | response to other organism | 10 | Inflammation | 3.57418E-08 |
| GO:0009607 | 9 | response to biotic stimulus | 10 | Inflammation | 3.51277E-08 |
| GO:0002684 | 9 | positive regulation of immune system process | 10 | Inflammation | 4.49536E-18 |
| GO:0050776 | 9 | regulation of immune response | 10 | Inflammation | 8.13095E-10 |
| GO:0001012 | 8 | RNA polymerase II regulatory region DNA binding | | | 1.90377E-15 |
| GO:0003700 | 8 | DNA-binding transcription factor activity | | | 1.12507E-15 |
| GO:0003690 | 8 | double-stranded DNA binding | | | 1.78095E-13 |
| GO:0000978 | 8 | RNA polymerase II proximal promoter sequence-specific DNA binding | | | 6.77988E-09 |
| GO:0000981 | 8 | DNA-binding transcription factor activity, RNA polymerase II-specific | | | 1.02558E-09 |
| GO:0000976 | 8 | transcription regulatory region sequence-specific DNA binding | | | 2.00203E-16 |
| GO:0000987 | 8 | proximal promoter sequence-specific DNA binding | | | 9.22408E-09 |
| GO:0001067 | 8 | regulatory region nucleic acid binding | | | 9.87505E-19 |
| GO:0000977 | 8 | RNA polymerase II regulatory region sequence-specific DNA binding | | | 6.70619E-15 |
| GO:0001228 | 8 | DNA-binding transcription activator activity, RNA polymerase II-specific | | | 2.60387E-08 |
| GO:0044212 | 8 | transcription regulatory region DNA binding | | | 1.31422E-18 |
| GO:1990837 | 8 | sequence-specific double-stranded DNA binding | | | 6.66934E-16 |
| GO:0016651 | 7 | oxidoreductase activity, acting on NAD(P)H | 9 | Mitochondrial_respiratory | 0.003439072 |
| GO:1990204 | 7 | oxidoreductase complex | 9 | Mitochondrial_respiratory | 1.43704E-07 |
| GO:0005747 | 7 | mitochondrial respiratory chain complex I | 9 | Mitochondrial_respiratory | 2.77582E-05 |
| GO:0098800 | 7 | inner mitochondrial membrane protein complex | 9 | Mitochondrial_respiratory | 2.7881E-10 |
| GO:0098803 | 7 | respiratory chain complex | 9 | Mitochondrial_respiratory | 8.21693E-06 |
| GO:0030964 | 7 | NADH dehydrogenase complex | 9 | Mitochondrial_respiratory | 2.77582E-05 |
| GO:0045271 | 7 | respiratory chain complex I | 9 | Mitochondrial_respiratory | 2.77582E-05 |
| GO:0098798 | 7 | mitochondrial protein complex | 9 | Mitochondrial_respiratory | 1.17665E-17 |
| GO:0044455 | 7 | mitochondrial membrane part | 9 | Mitochondrial_respiratory | 2.32137E-11 |
| GO:0070469 | 7 | respiratory chain | 9 | Mitochondrial_respiratory | 3.94265E-05 |
| GO:0019886 | 7 | antigen processing and presentation of exogenous peptide antigen via MHC class II | 8 | Antigen_processing | 1.35106E-06 |
| GO:0019882 | 7 | antigen processing and presentation | 8 | Antigen_processing | 3.36538E-11 |
| GO:0002504 | 7 | antigen processing and presentation of peptide or polysaccharide antigen via MHC class II | 8 | Antigen_processing | 3.66016E-06 |
| GO:0048002 | 7 | antigen processing and presentation of peptide antigen | 8 | Antigen_processing | 3.2794E-12 |
| GO:0042590 | 7 | antigen processing and presentation of exogenous peptide antigen via MHC class I | 8 | Antigen_processing | 0.032425536 |
| GO:0019884 | 7 | antigen processing and presentation of exogenous antigen | 8 | Antigen_processing | 7.23433E-14 |
| GO:0042611 | 7 | MHC protein complex | 8 | Antigen_processing | 1.64584E-07 |
| GO:0002495 | 7 | antigen processing and presentation of peptide antigen via MHC class II | 8 | Antigen_processing | 3.66016E-06 |
| GO:0042613 | 7 | MHC class II protein complex | 8 | Antigen_processing | 0.004028627 |
| GO:0002478 | 7 | antigen processing and presentation of exogenous peptide antigen | 8 | Antigen_processing | 4.20058E-12 |

| | | | | | |
|---|---|---|---|---|---|
| GO:1903509 | 6 | liposaccharide metabolic process | 7 | Lipid_metabolism | 0.000314429 |
| GO:0046486 | 6 | glycerolipid metabolic process | 7 | Lipid_metabolism | 0.008192361 |
| GO:0046488 | 6 | phosphatidylinositol metabolic process | 7 | Lipid_metabolism | 0.008352032 |
| GO:0044255 | 6 | cellular lipid metabolic process | 7 | Lipid_metabolism | 5.94468E-10 |
| GO:0006644 | 6 | phospholipid metabolic process | 7 | Lipid_metabolism | 1.48617E-05 |
| GO:0008654 | 6 | phospholipid biosynthetic process | 7 | Lipid_metabolism | 0.000169497 |
| GO:0008610 | 6 | lipid biosynthetic process | 7 | Lipid_metabolism | 1.30193E-10 |
| GO:0008289 | 6 | lipid binding | 7 | Lipid_metabolism | 0.011705127 |
| GO:0008203 | 6 | cholesterol metabolic process | 7 | Lipid_metabolism | 0.000102312 |
| GO:0008202 | 6 | steroid metabolic process | 7 | Lipid_metabolism | 0.000280039 |
| GO:0006695 | 6 | cholesterol biosynthetic process | 7 | Lipid_metabolism | 0.004937525 |
| GO:0006694 | 6 | steroid biosynthetic process | 7 | Lipid_metabolism | 0.004691877 |
| GO:0006672 | 6 | ceramide metabolic process | 7 | Lipid_metabolism | 0.003488202 |
| GO:0006665 | 6 | sphingolipid metabolic process | 7 | Lipid_metabolism | 0.000594468 |
| GO:0006664 | 6 | glycolipid metabolic process | 7 | Lipid_metabolism | 0.000224768 |
| GO:0006650 | 6 | glycerophospholipid metabolic process | 7 | Lipid_metabolism | 0.016704064 |
| GO:0006643 | 6 | membrane lipid metabolic process | 7 | Lipid_metabolism | 5.94468E-07 |
| GO:0016125 | 6 | sterol metabolic process | 7 | Lipid_metabolism | 2.62843E-06 |
| GO:0016126 | 6 | sterol biosynthetic process | 7 | Lipid_metabolism | 3.75841E-07 |
| GO:0098852 | 6 | lytic vacuole membrane | 6 | Lysosomal_proteolysis | 3.697E-09 |
| GO:1905369 | 6 | endopeptidase complex | 6 | Lysosomal_proteolysis | 4.65503E-06 |
| GO:1905368 | 6 | peptidase complex | 6 | Lysosomal_proteolysis | 8.60996E-05 |
| GO:0045807 | 6 | positive regulation of endocytosis | 6 | Lysosomal_proteolysis | 1.1742E-05 |
| GO:0044437 | 6 | vacuolar part | 6 | Lysosomal_proteolysis | 1.24052E-09 |
| GO:0061919 | 6 | process utilizing autophagic mechanism | 6 | Lysosomal_proteolysis | 2.83723E-06 |
| GO:0044440 | 6 | endosomal part | 6 | Lysosomal_proteolysis | 0.000959255 |
| GO:0007033 | 6 | vacuole organization | 6 | Lysosomal_proteolysis | 0.00486383 |
| GO:0006914 | 6 | autophagy | 6 | Lysosomal_proteolysis | 2.83723E-06 |
| GO:0006897 | 6 | endocytosis | 6 | Lysosomal_proteolysis | 0.004986654 |
| GO:0005774 | 6 | vacuolar membrane | 6 | Lysosomal_proteolysis | 1.38791E-09 |
| GO:0005773 | 6 | vacuole | 6 | Lysosomal_proteolysis | 4.40938E-26 |
| GO:0005770 | 6 | late endosome | 6 | Lysosomal_proteolysis | 0.009899614 |
| GO:0005769 | 6 | early endosome | 6 | Lysosomal_proteolysis | 0.013142168 |
| GO:0005768 | 6 | endosome | 6 | Lysosomal_proteolysis | 8.52399E-15 |
| GO:0005765 | 6 | lysosomal membrane | 6 | Lysosomal_proteolysis | 3.88124E-09 |
| GO:0005764 | 6 | lysosome | 6 | Lysosomal_proteolysis | 3.13201E-30 |
| GO:0016239 | 6 | positive regulation of macroautophagy | 6 | Lysosomal_proteolysis | 0.01965184 |
| GO:0016241 | 6 | regulation of macroautophagy | 6 | Lysosomal_proteolysis | 0.00017441 |
| GO:0000323 | 6 | lytic vacuole | 6 | Lysosomal_proteolysis | 3.13201E-30 |
| GO:0030100 | 6 | regulation of endocytosis | 6 | Lysosomal_proteolysis | 0.000217398 |
| GO:0010506 | 6 | regulation of autophagy | 6 | Lysosomal_proteolysis | 0.002935494 |
| GO:0034976 | 6 | response to endoplasmic reticulum stress | | | 0.000379526 |
| GO:0036503 | 6 | ERAD pathway | | | 2.75126E-05 |
| GO:0030176 | 6 | integral component of endoplasmic reticulum membrane | | | 3.94265E-05 |
| GO:0030433 | 6 | ubiquitin-dependent ERAD pathway | | | 0.000144932 |
| GO:0031227 | 6 | intrinsic component of endoplasmic reticulum membrane | | | 7.97128E-05 |
| GO:0042552 | 6 | myelination | 5 | Myelin | 8.35203E-07 |
| GO:0032291 | 6 | axon ensheathment in central nervous system | 5 | Myelin | 0.001220871 |
| GO:0043209 | 6 | myelin sheath | 5 | Myelin | 3.04604E-10 |
| GO:0007272 | 6 | ensheathment of neurons | 5 | Myelin | 8.5117E-08 |
| GO:0008366 | 6 | axon ensheathment | 5 | Myelin | 8.5117E-08 |
| GO:0022010 | 6 | central nervous system myelination | 5 | Myelin | 0.001220871 |
| GO:0019911 | 6 | structural constituent of myelin sheath | 5 | Myelin | 0.001940619 |
| GO:0006412 | 5 | translation | 4 | Translation | 5.17089E-29 |
| GO:0044391 | 5 | ribosomal subunit | 4 | Translation | 1.41248E-45 |
| GO:0043043 | 5 | peptide biosynthetic process | 4 | Translation | 1.38791E-28 |
| GO:0044445 | 5 | cytosolic part | 4 | Translation | 8.06954E-36 |
| GO:1901566 | 5 | organonitrogen compound biosynthetic process | 4 | Translation | 2.28453E-23 |
| GO:0015934 | 5 | large ribosomal subunit | 4 | Translation | 6.46054E-25 |
| GO:0005840 | 5 | ribosome | 4 | Translation | 6.60793E-40 |
| GO:0043604 | 5 | amide biosynthetic process | 4 | Translation | 5.6499E-26 |
| GO:0043603 | 5 | cellular amide metabolic process | 4 | Translation | 3.38994E-29 |
| GO:0005198 | 5 | structural molecule activity | 4 | Translation | 1.19139E-20 |
| GO:0034645 | 5 | cellular macromolecule biosynthetic process | 4 | Translation | 5.92012E-15 |
| GO:1990904 | 5 | ribonucleoprotein complex | 4 | Translation | 1.05015E-18 |
| GO:0009059 | 5 | macromolecule biosynthetic process | 4 | Translation | 8.29062E-12 |
| GO:0006518 | 5 | peptide metabolic process | 4 | Translation | 6.59565E-25 |
| GO:0022625 | 5 | cytosolic large ribosomal subunit | 4 | Translation | 1.43704E-26 |
| GO:0003735 | 5 | structural constituent of ribosome | 4 | Translation | 2.05116E-47 |
| GO:0009260 | 4 | ribonucleotide biosynthetic process | | | 0.020634432 |
| GO:0019693 | 4 | ribose phosphate metabolic process | | | 0.020143136 |
| GO:0006753 | 4 | nucleoside phosphate metabolic process | | | 2.24768E-05 |
| GO:0009117 | 4 | nucleotide metabolic process | | | 0.000146161 |
| GO:0019637 | 4 | organophosphate metabolic process | | | 4.96209E-11 |
| GO:0019438 | 4 | aromatic compound biosynthetic process | | | 0.025547392 |
| GO:0009165 | 4 | nucleotide biosynthetic process | | | 0.03746132 |
| GO:0018130 | 4 | heterocycle biosynthetic process | | | 0.009727661 |
| GO:0046390 | 4 | ribose phosphate biosynthetic process | | | 0.001473888 |
| GO:1901135 | 4 | carbohydrate derivative metabolic process | | | 1.26509E-13 |
| GO:1901137 | 4 | carbohydrate derivative biosynthetic process | | | 1.52302E-05 |
| GO:1901362 | 4 | organic cyclic compound biosynthetic process | | | 0.046795944 |
| GO:0090407 | 4 | organophosphate biosynthetic process | | | 2.92321E-08 |
| GO:0055086 | 4 | nucleobase-containing small molecule metabolic process | | | 8.74507E-05 |
| GO:0043161 | 3 | proteasome-mediated ubiquitin-dependent protein catabolic process | 3 | Proteasomal_proteolysis | 0.000168269 |
| GO:0043632 | 3 | modification-dependent macromolecule catabolic process | 3 | Proteasomal_proteolysis | 1.98975E-06 |
| GO:0010498 | 3 | proteasomal protein catabolic process | 3 | Proteasomal_proteolysis | 3.94265E-05 |
| GO:0019941 | 3 | modification-dependent protein catabolic process | 3 | Proteasomal_proteolysis | 4.79014E-07 |
| GO:0051603 | 3 | proteolysis involved in cellular protein catabolic process | 3 | Proteasomal_proteolysis | 1.90377E-09 |

| | | | | | |
|---|---|---|---|---|---|
| GO:0030163 | 3 | protein catabolic process | 3 | Proteasomal_proteolysis | 1.19999E-08 |
| GO:0006511 | 3 | ubiquitin-dependent protein catabolic process | 3 | Proteasomal_proteolysis | 1.03909E-05 |
| GO:0044265 | 3 | cellular macromolecule catabolic process | 3 | Proteasomal_proteolysis | 3.25484E-07 |
| GO:0009057 | 3 | macromolecule catabolic process | 3 | Proteasomal_proteolysis | 1.85464E-08 |
| GO:1901565 | 3 | organonitrogen compound catabolic process | 3 | Proteasomal_proteolysis | 9.93646E-12 |
| GO:0043269 | 2 | regulation of ion transport | 2 | Synapse | 5.60077E-09 |
| GO:0072503 | 2 | cellular divalent inorganic cation homeostasis | 2 | Synapse | 0.011287526 |
| GO:2001257 | 2 | regulation of cation channel activity | 2 | Synapse | 2.00203E-05 |
| GO:0043235 | 2 | receptor complex | 2 | Synapse | 0.005084914 |
| GO:0097060 | 2 | synaptic membrane | 2 | Synapse | 4.06547E-05 |
| GO:0043197 | 2 | dendritic spine | 2 | Synapse | 0.029723408 |
| GO:0043025 | 2 | neuronal cell body | 2 | Synapse | 5.26915E-08 |
| GO:0042391 | 2 | regulation of membrane potential | 2 | Synapse | 0.018055128 |
| GO:0072507 | 2 | divalent inorganic cation homeostasis | 2 | Synapse | 0.029969056 |
| GO:0098589 | 2 | membrane region | 2 | Synapse | 0.000151074 |
| GO:0044297 | 2 | cell body | 2 | Synapse | 2.61615E-10 |
| GO:0010959 | 2 | regulation of metal ion transport | 2 | Synapse | 9.50658E-05 |
| GO:0044309 | 2 | neuron spine | 2 | Synapse | 0.020634432 |
| GO:0007626 | 2 | locomotory behavior | 2 | Synapse | 0.02272244 |
| GO:0007613 | 2 | memory | 2 | Synapse | 3.10745E-05 |
| GO:0007611 | 2 | learning or memory | 2 | Synapse | 2.03888E-07 |
| GO:0007610 | 2 | behavior | 2 | Synapse | 5.35513E-06 |
| GO:0045121 | 2 | membrane raft | 2 | Synapse | 2.59159E-05 |
| GO:0045211 | 2 | postsynaptic membrane | 2 | Synapse | 0.000289865 |
| GO:0051924 | 2 | regulation of calcium ion transport | 2 | Synapse | 0.01351064 |
| GO:0050890 | 2 | cognition | 2 | Synapse | 5.40426E-07 |
| GO:0050801 | 2 | ion homeostasis | 2 | Synapse | 0.040654744 |
| GO:0050804 | 2 | modulation of chemical synaptic transmission | 2 | Synapse | 0.007762477 |
| GO:0044057 | 2 | regulation of system process | 2 | Synapse | 0.00609207 |
| GO:0050807 | 2 | regulation of synapse organization | 2 | Synapse | 0.000380754 |
| GO:0098793 | 2 | presynapse | 2 | Synapse | 0.00332853 |
| GO:0098797 | 2 | plasma membrane protein complex | 2 | Synapse | 5.35513E-05 |
| GO:0023052 | 2 | signaling | 2 | Synapse | 0.03439072 |
| GO:0022898 | 2 | regulation of transmembrane transporter activity | 2 | Synapse | 0.000118525 |
| GO:1904062 | 2 | regulation of cation transmembrane transport | 2 | Synapse | 0.002210832 |
| GO:0030425 | 2 | dendrite | 2 | Synapse | 0.000180551 |
| GO:0031644 | 2 | regulation of neurological system process | 2 | Synapse | 0.017441008 |
| GO:0032409 | 2 | regulation of transporter activity | 2 | Synapse | 4.36025E-05 |
| GO:0032412 | 2 | regulation of ion transmembrane transporter activity | 2 | Synapse | 5.35513E-05 |
| GO:0019725 | 2 | cellular homeostasis | 2 | Synapse | 0.00050235 |
| GO:0033267 | 2 | axon part | 2 | Synapse | 0.00552708 |
| GO:0098794 | 2 | postsynapse | 2 | Synapse | 0.007332593 |
| GO:0099572 | 2 | postsynaptic specialization | 2 | Synapse | 1.07839E-06 |
| GO:0034762 | 2 | regulation of transmembrane transport | 2 | Synapse | 6.49739E-06 |
| GO:0034765 | 2 | regulation of ion transmembrane transport | 2 | Synapse | 1.80551E-06 |
| GO:0099240 | 2 | intrinsic component of synaptic membrane | 2 | Synapse | 7.16064E-07 |
| GO:0099699 | 2 | integral component of synaptic membrane | 2 | Synapse | 2.46876E-06 |
| GO:0099177 | 2 | regulation of trans-synaptic signaling | 2 | Synapse | 0.008253773 |
| GO:0014069 | 2 | postsynaptic density | 2 | Synapse | 1.75638E-06 |
| GO:0098857 | 2 | membrane microdomain | 2 | Synapse | 2.80039E-05 |
| GO:0098889 | 2 | intrinsic component of presynaptic membrane | 2 | Synapse | 0.004065474 |
| GO:0098936 | 2 | intrinsic component of postsynaptic membrane | 2 | Synapse | 2.21083E-07 |
| GO:0098978 | 2 | glutamatergic synapse | 2 | Synapse | 2.14942E-09 |
| GO:0099055 | 2 | integral component of postsynaptic membrane | 2 | Synapse | 1.54758E-06 |
| GO:0099056 | 2 | integral component of presynaptic membrane | 2 | Synapse | 0.022968088 |
| GO:0008324 | 1 | cation transmembrane transporter activity | 1 | Ion_channel | 1.91605E-07 |
| GO:0015318 | 1 | inorganic molecular entity transmembrane transporter activity | 1 | Ion_channel | 2.1617E-07 |
| GO:0034220 | 1 | ion transmembrane transport | 1 | Ion_channel | 1.15086E-05 |
| GO:0034702 | 1 | ion channel complex | 1 | Ion_channel | 0.010047003 |
| GO:0005216 | 1 | ion channel activity | 1 | Ion_channel | 0.020143136 |
| GO:1902495 | 1 | transmembrane transporter complex | 1 | Ion_channel | 0.006951838 |
| GO:0055085 | 1 | transmembrane transport | 1 | Ion_channel | 9.81364E-07 |
| GO:0015077 | 1 | monovalent inorganic cation transmembrane transporter activity | 1 | Ion_channel | 0.001916054 |
| GO:0046873 | 1 | metal ion transmembrane transporter activity | 1 | Ion_channel | 3.37766E-06 |
| GO:0034703 | 1 | cation channel complex | 1 | Ion_channel | 0.016704064 |
| GO:0022838 | 1 | substrate-specific channel activity | 1 | Ion_channel | 0.03930368 |
| GO:0022857 | 1 | transmembrane transporter activity | 1 | Ion_channel | 3.79526E-08 |
| GO:0022890 | 1 | inorganic cation transmembrane transporter activity | 1 | Ion_channel | 1.15455E-07 |
| GO:0015075 | 1 | ion transmembrane transporter activity | 1 | Ion_channel | 3.50048E-06 |
| GO:1990351 | 1 | transporter complex | 1 | Ion_channel | 0.011582303 |
| GO:0005261 | 1 | cation channel activity | 1 | Ion_channel | 0.010587429 |
| GO:0006812 | 1 | cation transport | 1 | Ion_channel | 0.000249333 |
| GO:0098656 | 6 | anion transmembrane transport | | | 0.027881048 |
| GO:0120039 | 6 | plasma membrane bounded cell projection morphogenesis | | | 0.02640716 |
| GO:0098657 | 6 | import into cell | | | 7.11151E-06 |
| GO:0042277 | 6 | peptide binding | | | 0.000567447 |
| GO:0120036 | 6 | plasma membrane bounded cell projection organization | | | 9.28549E-08 |
| GO:0042274 | 6 | ribosomal small subunit biogenesis | | | 0.000499894 |
| GO:0035567 | 6 | non-canonical Wnt signaling pathway | | | 0.001202447 |
| GO:0099503 | 6 | secretory vesicle | | | 0.000696412 |
| GO:0098982 | 6 | GABA-ergic synapse | | | 0.013879112 |
| GO:0098858 | 6 | actin-based cell projection | | | 0.0245648 |
| GO:0034660 | 6 | ncRNA metabolic process | | | 0.008401162 |
| GO:0098805 | 6 | whole membrane | | | 1.01084E-18 |
| GO:0042176 | 6 | regulation of protein catabolic process | | | 0.003365378 |
| GO:0098742 | 6 | cell-cell adhesion via plasma-membrane adhesion molecules | | | 5.10948E-07 |
| GO:0099568 | 6 | cytoplasmic region | | | 0.003107447 |
| GO:1901136 | 6 | carbohydrate derivative catabolic process | | | 0.001510735 |

| | | | | | |
|---|---|---|---|---|---|
| GO:0034613 | 6 | cellular protein localization | | | 0.006165765 |
| GO:1990234 | 6 | transferase complex | | | 0.000210029 |
| GO:1905039 | 6 | carboxylic acid transmembrane transport | | | 0.022353968 |
| GO:0030182 | 6 | neuron differentiation | | | 0.012528048 |
| GO:0030246 | 6 | carbohydrate binding | | | 0.000433569 |
| GO:1903825 | 6 | organic acid transmembrane transport | | | 0.022353968 |
| GO:0030888 | 6 | regulation of B cell proliferation | | | 0.030951648 |
| GO:0031012 | 6 | extracellular matrix | | | 0.000691499 |
| GO:0031175 | 6 | neuron projection development | | | 0.000881876 |
| GO:0034622 | 6 | cellular protein-containing complex assembly | | | 2.43192E-09 |
| GO:0031329 | 6 | regulation of cellular catabolic process | | | 6.30087E-08 |
| GO:1902653 | 6 | secondary alcohol biosynthetic process | | | 0.000894159 |
| GO:0031966 | 6 | mitochondrial membrane | | | 1.24052E-17 |
| GO:0031968 | 6 | organelle outer membrane | | | 0.004053192 |
| GO:0031974 | 6 | membrane-enclosed lumen | | | 0.005920117 |
| GO:1902652 | 6 | secondary alcohol metabolic process | | | 2.10029E-05 |
| GO:1901617 | 6 | organic hydroxy compound biosynthetic process | | | 0.005870987 |
| GO:1901615 | 6 | organic hydroxy compound metabolic process | | | 2.40735E-06 |
| GO:0032940 | 6 | secretion by cell | | | 0.010120698 |
| GO:0032989 | 6 | cellular component morphogenesis | | | 0.000541654 |
| GO:0033218 | 6 | amide binding | | | 2.41963E-06 |
| GO:0140101 | 6 | catalytic activity, acting on a tRNA | | | 0.03316248 |
| GO:0031300 | 6 | intrinsic component of organelle membrane | | | 0.007344875 |
| GO:0042788 | 6 | polysomal ribosome | | | 2.98462E-05 |
| GO:0043233 | 6 | organelle lumen | | | 0.005920117 |
| GO:0098609 | 6 | cell-cell adhesion | | | 1.48617E-07 |
| GO:0055114 | 6 | oxidation-reduction process | | | 9.93646E-08 |
| GO:0045503 | 6 | dynein light chain binding | | | 0.002493327 |
| GO:0045505 | 6 | dynein intermediate chain binding | | | 0.028617992 |
| GO:0046165 | 6 | alcohol biosynthetic process | | | 0.003512766 |
| GO:0051959 | 6 | dynein light intermediate chain binding | | | 0.001473888 |
| GO:0046982 | 6 | protein heterodimerization activity | | | 0.037829792 |
| GO:0048037 | 6 | cofactor binding | | | 0.000741857 |
| GO:0048193 | 6 | Golgi vesicle transport | | | 0.021248552 |
| GO:0048468 | 6 | cell development | | | 5.99381E-11 |
| GO:0048471 | 6 | perinuclear region of cytoplasm | | | 3.9058E-05 |
| GO:0048511 | 6 | rhythmic process | | | 0.000178095 |
| GO:0048598 | 6 | embryonic morphogenesis | | | 9.05213E-07 |
| GO:0048609 | 6 | multicellular organismal reproductive process | | | 0.001645842 |
| GO:0051186 | 6 | cofactor metabolic process | | | 0.001866925 |
| GO:0051015 | 6 | actin filament binding | | | 0.000417602 |
| GO:0048646 | 6 | anatomical structure formation involved in morphogenesis | | | 0.00250561 |
| GO:0048660 | 6 | regulation of smooth muscle cell proliferation | | | 1.65812E-05 |
| GO:0048666 | 6 | neuron development | | | 0.047532888 |
| GO:0048729 | 6 | tissue morphogenesis | | | 0.01780948 |
| GO:0048731 | 6 | system development | | | 2.8618E-06 |
| GO:0048812 | 6 | neuron projection morphogenesis | | | 0.012773696 |
| GO:0048858 | 6 | cell projection morphogenesis | | | 0.03807544 |
| GO:0050662 | 6 | coenzyme binding | | | 1.35106E-05 |
| GO:0050839 | 6 | cell adhesion molecule binding | | | 0.000120613 |
| GO:0044447 | 6 | axoneme part | | | 0.040777568 |
| GO:0042803 | 6 | protein homodimerization activity | | | 0.000589555 |
| GO:0044441 | 6 | ciliary part | | | 0.021002904 |
| GO:0098588 | 6 | bounding membrane of organelle | | | 5.68675E-15 |
| GO:0097485 | 6 | neuron projection guidance | | | 0.007541394 |
| GO:0072655 | 6 | establishment of protein localization to mitochondrion | | | 0.003697002 |
| GO:0072594 | 6 | establishment of protein localization to organelle | | | 0.00012528 |
| GO:0071855 | 6 | neuropeptide receptor binding | | | 0.00342679 |
| GO:0043254 | 6 | regulation of protein complex assembly | | | 0.004139169 |
| GO:0071826 | 6 | ribonucleoprotein complex subunit organization | | | 0.021248552 |
| GO:0043436 | 6 | oxoacid metabolic process | | | 1.83008E-08 |
| GO:0070727 | 6 | cellular macromolecule localization | | | 0.007025533 |
| GO:0070585 | 6 | protein localization to mitochondrion | | | 0.009187235 |
| GO:0044087 | 6 | regulation of cellular component biogenesis | | | 0.016458416 |
| GO:0044089 | 6 | positive regulation of cellular component biogenesis | | | 0.006128918 |
| GO:0070382 | 6 | exocytic vesicle | | | 0.000119508 |
| GO:0044283 | 6 | small molecule biosynthetic process | | | 3.13201E-08 |
| GO:0070180 | 6 | large ribosomal subunit rRNA binding | | | 2.00203E-05 |
| GO:0070013 | 6 | intracellular organelle lumen | | | 0.004974372 |
| GO:0044389 | 6 | ubiquitin-like protein ligase binding | | | 0.03500484 |
| GO:0044429 | 6 | mitochondrial part | | | 1.69497E-21 |
| GO:0062023 | 6 | collagen-containing extracellular matrix | | | 0.000716064 |
| GO:0060589 | 6 | nucleoside-triphosphatase regulator activity | | | 0.015721472 |
| GO:0060429 | 6 | epithelium development | | | 0.006006094 |
| GO:0044431 | 6 | Golgi apparatus part | | | 2.80039E-05 |
| GO:0044432 | 6 | endoplasmic reticulum part | | | 1.1484E-23 |
| GO:0044433 | 6 | cytoplasmic vesicle part | | | 1.80551E-06 |
| GO:0030162 | 6 | regulation of proteolysis | | | 0.00368472 |
| GO:0000028 | 6 | ribosomal small subunit assembly | | | 0.000283723 |
| GO:0007389 | 6 | pattern specification process | | | 7.75019E-05 |
| GO:0007264 | 6 | small GTPase mediated signal transduction | | | 0.024073504 |
| GO:0007218 | 6 | neuropeptide signaling pathway | | | 0.000130193 |
| GO:0007167 | 6 | enzyme linked receptor protein signaling pathway | | | 0.014001936 |
| GO:0007159 | 6 | leukocyte cell-cell adhesion | | | 0.002210832 |
| GO:0007157 | 6 | heterophilic cell-cell adhesion via plasma membrane cell adhesion molecules | | | 0.041514512 |
| GO:0007155 | 6 | cell adhesion | | | 1.47389E-10 |
| GO:0007017 | 6 | microtubule-based process | | | 0.002186267 |
| GO:0007010 | 6 | cytoskeleton organization | | | 0.002468762 |

| | | | | | |
|---|---|---|---|---|---|
| GO:0007005 | 6 | mitochondrion organization | | | 0.002321374 |
| GO:0006886 | 6 | intracellular protein transport | | | 2.05116E-08 |
| GO:0007399 | 6 | nervous system development | | | 1.19999E-06 |
| GO:0006865 | 6 | amino acid transport | | | 0.039549328 |
| GO:0007411 | 6 | axon guidance | | | 0.011127854 |
| GO:0007631 | 6 | feeding behavior | | | 6.33772E-07 |
| GO:0009798 | 6 | axis specification | | | 0.013756288 |
| GO:0002577 | 6 | regulation of antigen processing and presentation | | | 0.006509672 |
| GO:0008569 | 6 | ATP-dependent microtubule motor activity, minus-end-directed | | | 0.019283368 |
| GO:0008360 | 6 | regulation of cell shape | | | 0.011078725 |
| GO:0002579 | 6 | positive regulation of antigen processing and presentation | | | 0.000437253 |
| GO:0008283 | 6 | cell proliferation | | | 0.014984528 |
| GO:0008217 | 6 | regulation of blood pressure | | | 0.013756288 |
| GO:0008047 | 6 | enzyme activator activity | | | 0.001387911 |
| GO:0008021 | 6 | synaptic vesicle | | | 0.000173182 |
| GO:0007507 | 6 | heart development | | | 2.48104E-05 |
| GO:0006839 | 6 | mitochondrial transport | | | 0.003721567 |
| GO:0006836 | 6 | neurotransmitter transport | | | 0.000241963 |
| GO:0006732 | 6 | coenzyme metabolic process | | | 0.039672152 |
| GO:0004553 | 6 | hydrolase activity, hydrolyzing O-glycosyl compounds | | | 0.006681626 |
| GO:0005179 | 6 | hormone activity | | | 3.47592E-07 |
| GO:0005789 | 6 | endoplasmic reticulum membrane | | | 0.000137563 |
| GO:0005184 | 6 | neuropeptide hormone activity | | | 8.90474E-06 |
| GO:0005743 | 6 | mitochondrial inner membrane | | | 3.86896E-16 |
| GO:0005741 | 6 | mitochondrial outer membrane | | | 0.007737912 |
| GO:0005539 | 6 | glycosaminoglycan binding | | | 0.048269832 |
| GO:0005201 | 6 | extracellular matrix structural constituent | | | 0.006976403 |
| GO:0003774 | 6 | motor activity | | | 0.000393037 |
| GO:0003712 | 6 | transcription coregulator activity | | | 0.046550296 |
| GO:0005911 | 6 | cell-cell junction | | | 0.040654744 |
| GO:0005975 | 6 | carbohydrate metabolic process | | | 4.15145E-05 |
| GO:0030133 | 6 | transport vesicle | | | 6.44826E-08 |
| GO:0009887 | 6 | animal organ morphogenesis | | | 3.15658E-06 |
| GO:0002902 | 6 | regulation of B cell apoptotic process | | | 0.03316248 |
| GO:0006520 | 6 | cellular amino acid metabolic process | | | 0.048761128 |
| GO:0006457 | 6 | protein folding | | | 6.80445E-08 |
| GO:0006082 | 6 | organic acid metabolic process | | | 1.10173E-07 |
| GO:0003006 | 6 | developmental process involved in reproduction | | | 0.000431112 |
| GO:0006066 | 6 | alcohol metabolic process | | | 1.30193E-08 |
| GO:0003341 | 6 | cilium movement | | | 0.000361103 |
| GO:0003682 | 6 | chromatin binding | | | 0.000289865 |
| GO:0009888 | 6 | tissue development | | | 0.00022354 |
| GO:0008092 | 6 | cytoskeletal protein binding | | | 3.73385E-05 |
| GO:0009896 | 6 | positive regulation of catabolic process | | | 0.001375629 |
| GO:0016192 | 6 | vesicle-mediated transport | | | 9.92418E-13 |
| GO:0002181 | 6 | cytoplasmic translation | | | 6.79217E-12 |
| GO:0016491 | 6 | oxidoreductase activity | | | 6.48511E-06 |
| GO:0001505 | 6 | regulation of neurotransmitter levels | | | 0.007160639 |
| GO:0009894 | 6 | regulation of catabolic process | | | 1.02681E-08 |
| GO:0019752 | 6 | carboxylic acid metabolic process | | | 2.98462E-08 |
| GO:0019843 | 6 | rRNA binding | | | 6.58337E-05 |
| GO:0019866 | 6 | organelle inner membrane | | | 1.46161E-14 |
| GO:0015935 | 6 | small ribosomal subunit | | | 2.06344E-20 |
| GO:0019867 | 6 | outer membrane | | | 0.004053192 |
| GO:0022402 | 6 | cell cycle process | | | 0.004360252 |
| GO:0000902 | 6 | cell morphogenesis | | | 0.008081819 |
| GO:0000502 | 6 | proteasome complex | | | 1.87921E-05 |
| GO:0022411 | 6 | cellular component disassembly | | | 0.014247584 |
| GO:0000122 | 6 | negative regulation of transcription by RNA polymerase II | | | 2.41963E-05 |
| GO:0022610 | 6 | biological adhesion | | | 1.40019E-10 |
| GO:0022627 | 6 | cytosolic small ribosomal subunit | | | 4.43395E-24 |
| GO:0022804 | 6 | active transmembrane transporter activity | | | 0.000191605 |
| GO:0030030 | 6 | cell projection organization | | | 1.46161E-08 |
| GO:0015631 | 6 | tubulin binding | | | 0.00404091 |
| GO:0005509 | 6 | calcium ion binding | | | 1.54758E-06 |
| GO:0010817 | 6 | regulation of hormone levels | | | 0.021862672 |
| GO:0009986 | 6 | cell surface | | | 1.55986E-11 |
| GO:0015291 | 6 | secondary active transmembrane transporter activity | | | 0.021125728 |
| GO:0002443 | 7 | leukocyte mediated immunity | | | 1.21104E-05 |
| GO:0002460 | 7 | adaptive immune response based on somatic recombination of immune receptors built from immunoglobulin superfamily domains | | | 3.38994E-06 |
| GO:0019724 | 7 | B cell mediated immunity | | | 7.68878E-05 |
| GO:0002449 | 7 | lymphocyte mediated immunity | | | 1.31422E-06 |
| GO:0016064 | 7 | immunoglobulin mediated immune response | | | 2.1617E-05 |
| GO:2001233 | 12 | regulation of apoptotic signaling pathway | | | 0.019283368 |
| GO:1902533 | 12 | positive regulation of intracellular signal transduction | | | 3.32853E-07 |
| GO:1903532 | 12 | positive regulation of secretion by cell | | | 0.04483076 |
| GO:0032386 | 12 | regulation of intracellular transport | | | 0.002444198 |
| GO:1902532 | 12 | negative regulation of intracellular signal transduction | | | 0.049866544 |
| GO:1903530 | 12 | regulation of secretion by cell | | | 2.3705E-05 |
| GO:0070201 | 12 | regulation of establishment of protein localization | | | 0.017318184 |
| GO:0060627 | 12 | regulation of vesicle-mediated transport | | | 1.05383E-05 |
| GO:0051962 | 12 | positive regulation of nervous system development | | | 0.003242554 |
| GO:0051347 | 12 | positive regulation of transferase activity | | | 0.00250561 |
| GO:0051345 | 12 | positive regulation of hydrolase activity | | | 0.021248552 |
| GO:0051336 | 12 | regulation of hydrolase activity | | | 0.005993811 |
| GO:1901214 | 12 | regulation of neuron death | | | 0.000386896 |
| GO:0010942 | 12 | positive regulation of cell death | | | 0.03561896 |

### Table S3

| Black | Black_continue_ | Green | Green_continue_ | Green_continue_ | Greenyellow | Magenta | Magenta_continue_ | Pink | Pink_continue_1 | Purple | Red | Red_continue_1 | Red_continue_2 | Tan | Yellow | Yellow_continue | Yellow_continue |
|---|---|---|---|---|---|---|---|---|---|---|---|---|---|---|---|---|---|
| 2310022B05Rik | Mgp | 1500009C09Rik | Gm2a | Prpsap1 | AA474408 | 1700019D03Rik | Rap1gds1 | 2210013O21Rik | Itih3 | Apoe | 2810468N07Rik | Mal | Tmem98 | 2810433D01Rik | Chchd2 | Gm4997 | Rpl34-ps1 |
| AI464131 | Mgst1 | 2310061I04Rik | Gm3756 | Prr13 | Arhgap5 | 1810041L15Rik | Rasd1 | 6330403K07Rik | Klhl13 | Arpc1b | 5031439G07Rik | Mapk8ip1 | Tnfaip6 | Ace | Eef1a1 | Gm5075 | Rpl34-ps2 |
| Abhd4 | Mid1ip1 | 2410015M20Rik | Gm43712 | Prune2 | BC001981 | Adarb1 | Rgs16 | A730017C20Rik | Lonrf2 | Axl | Abca2 | Mapt | Tppp3 | Akr7a5 | Fau | Gm5093 | Rpl35 |
| Aco2 | Mlc1 | Acss2 | Gm6166 | Psmb10 | Cachd1 | Adarb2 | Rgs4 | AW551984 | Ly6h | B2m | Anln | Mbp | Tprn | Asxl2 | Ftl1 | Gm5453 | Rpl35a |
| Acot11 | Mt1 | Acyp2 | Gm6682 | Psmc5 | Crlf1 | Adcy8 | Rgs8 | Abat | Nap1l5 | C1qa | Apod | Megf10 | Trf | Atp1a3 | Ftl1-ps1 | Gm5735 | Rpl37 |
| Acsbg1 | Mt2 | Agpat5 | Gm8566 | Ptpa | Emilin2 | Adgra1 | Rims3 | Ahi1 | Ncam1 | C1qb | Arhgap23 | Mobp | Trim59 | Bloc1s3 | Gas5 | Gm5822 | Rpl37rt |
| Acsl3 | Myoc | Aig1 | Gm9803 | RP24-91J7.1 | Fbn1 | Adgrf5 | Rnf152 | Amigo2 | Ndn | C1qc | Arhgef10 | Mog | Tsc22d4 | Clic6 | Gm10036 | Gm5963 | Rpl38 |
| Adcyap1r1 | Ndrg2 | Aktip | Gpr137 | Rab11fip5 | Gcn1l1 | Adra1b | Rora | Ankrd34b | Nenf | C4a | Arrdc3 | Myrf | Tspan2 | Cnot3 | Gm10059 | Gm6030 | Rpl38-ps1 |
| Adgrg1 | Nefh | Aldh5a1 | Haghl | Rab4a | Gm11410 | Amotl1 | Samd5 | Ap1s2 | Nnat | C4b | Arsg | Nacc2 | Tspyl4 | Col9a3 | Gm10073 | Gm6134 | Rpl38-ps2 |
| Agt | Ntm | Anapc2 | Hdhd2 | Ranbp6 | Gm12155 | Ankrd34c | Sema4g | Araf | Nrip2 | Cd63 | Aspa | Ndrg1 | Ttll7 | Enpp2 | Gm10076 | Gm6170 | Rpl39-ps |
| Aldh1a1 | Ntsr2 | Arl2 | Hmga1-rs1 | Rgs10 | Gm15564 | Arhgap44 | Sema5a | Arhgdig | Nrsn2 | Cd63-ps | Atp1b3 | Neat1 | Ttyh2 | Folr1 | Gm10086 | Gm6222 | Rpl41 |
| Aldoc | Paqr8 | Arl4c | Hs6st2 | Rida | Gm20594 | Atp6ap2 | Sgpp2 | Arxes2 | Pafah1b3 | Cd9 | Bcas1 | Nkx6-2 | Tubb4a | Foxn3 | Gm10126 | Gm6272 | Rpl9 |
| Anxa5 | Pink1 | Arl6ip4 | Hsd17b12 | Rsrc1 | Gm24245 | Atpaf1 | Sh3d19 | B630019K06Rik | Pak3 | Clu | Bin1 | Nmral1 | Ugt8a | Gm10801 | Gm10154 | Gm6548 | Rplp1 |
| Apln | Pla2g7 | Asb13 | Inafm1 | Rtcb | Gm24270 | Avpi1 | Shank3 | Baiap3 | Pcbd1 | Csf1r | Bnip3l | Olig1 | Wnk1 | Gm15728 | Gm10180 | Gm6745 | Rplp2 |
| Aqp4 | Ppp1r9b | Atp5f1 | Inpp5f | Sacm1l | Gm25911 | B4galt5 | Shox2 | Bend6 | Pcdh17 | Cst3 | Car2 | Olig2 | Wscd1 | Gm43841 | Gm10263 | Gm6767 | Rps10 |
| Asrgl1 | Rpl26 | BC029722 | Insig1 | Sall2 | Gm26917 | Bok | Slc29a1 | Bex1 | Pcsk1n | Ctsa | Carhsp1 | Pacs2 | | Gm4750 | Gm10275 | Gm6905 | Rps10-ps1 |
| Atp1a2 | Rpl31 | B230118H07Rik | Ipo4 | Scn1a | Gm28439 | Cacna1g | Spock3 | Bex2 | Pdxk | Ctsb | Cd81 | Padi2 | | Hmgcl | Gm10288 | Gm6977 | Rps10-ps2 |
| Atp1b2 | Rpn2 | Babam1 | Iscu | Sdr39u1 | Gm42418 | Ccdc136 | Sptbn1 | Bex3 | Peg3 | Ctsd | Cd82 | Pde8a | | Igf2 | Gm10343 | Gm7079 | Rps11 |
| Bcan | Rps27 | Bcap31 | Kansl3 | Slc20a1 | Gm43743 | Cd47 | St3gal4 | Bex4 | Pgrmc1 | Ctsh | Cdc42ep1 | Pdlim2 | | Igfbp2 | Gm10443 | Gm7123 | Rps12 |
| Bsg | S100a1 | Bcat1 | Kcnh2 | Slc22a23 | Ift46 | Chrna4 | Synpo2 | Blcap | Plcxd2 | Ctsl | Cdc42ep2 | Phldb1 | | Kif15 | Gm10689 | Gm7206 | Rps12-ps10 |
| Btbd17 | S1pr1 | Bpgm | Kctd2 | Slc25a33 | Kif5a | Cit | Syt9 | Cacna2d2 | Pnck | Ctss | Cers2 | Pip4k2a | | Kif3a | Gm10736 | Gm7290 | Rps12-ps19 |
| Cacng4 | Sash1 | Cat | L1cam | Slc25a39 | Kif5b | Clmn | Tanc1 | Cacybp | Pnma3 | Ctsz | Cldn11 | Pkp4 | | Kl | Gm11425 | Gm7363 | Rps12-ps24 |
| Cadm4 | Scg3 | Ccdc92 | Lamtor2 | Slc32a1 | Lars2 | Cpne9 | Tcf7l2 | Calb2 | Pnmal2 | Cx3cr1 | Clic4 | Plekhb1 | | Lrrc17 | Gm11694 | Gm7384 | Rps12-ps5 |
| Cbln1 | Sec62 | Cd99l2 | Lbh | Slc38a1 | Lbx1 | Ctnnb1 | Thsd7a | Caly | Podxl2 | Cyba | Cnp | Plin3 | | Oaz1 | Gm11878 | Gm7429 | Rps12-ps9 |
| Cd59a | Sema6a | Cdc123 | Lgi2 | Smarcd3 | Malat1 | Dbndd1 | Tmem163 | Camk2d | Prkacb | Fcer1g | Cntn2 | Pllp | | Pip5k1a | Gm11942 | Gm7504 | Rps12l1 |
| Chga | Serpine2 | Cept1 | Limk1 | Smim12 | Mios | Edil3 | Tmem41a | Camk2n2 | Prkar1a | Fcgr3 | Cryab | Plod1 | | Prlr | Gm12013 | Gm7536 | Rps15 |
| Ckb | Slc16a1 | Cfl2 | Limk2 | Smim19 | Mir6236 | Elmo1 | Tnnt1 | Canx | Prmt2 | Fcrls | Csrp1 | Plp1 | | Sez6 | Gm12034 | Gm7589 | Rps15-ps2 |
| Ckb-ps1 | Slc17a6 | Chst10 | Lrrc24 | Snrk | Mt3 | Eml1 | Tpd52l1 | Cartpt | Rab9b | Gfap | Dbi | Plxnb3 | | Slc13a4 | Gm12191 | Gm7618 | Rps16 |
| Cntfr | Slc25a18 | Clcn6 | Lrrc59 | Snrpe | Nop9 | Enho | Trnp1 | Cbln2 | Rcn1 | Gns | Ddr1 | Pmp22 | | Sostdc1 | Gm12350 | Gm7867 | Rps16-ps2 |
| Cox7a2l | Slc2a1 | Cmas | Lrrc8a | Snx1 | Plpp3 | Eps8l2 | Tspan33 | Cbln4 | Rcn2 | Gpx4 | Degs1 | Ppp1r14a | | Sulf1 | Gm12380 | Gm7990 | Rps17 |
| Ctnnbip1 | Slc39a12 | Col4a3bp | Lsm6 | Snx5 | Ptprz1 | Fam20c | Tspan9 | Cd200 | Resp18 | Grn | Desi1 | Prdx1 | | Supt7l | Gm12466 | Gm8129 | Rps18 |
| Ctxn3 | Slc6a1 | Coprs | Manf | Sod1 | Purb | Fndc4 | Tunar | Cd83 | Rgag4 | Gusb | Dnajb2 | Prr18 | | Timm22 | Gm12918 | Gm8213 | Rps19 |
| Cyp2j9 | Slc6a11 | Cops3 | Mfsd6 | Srpr | Rn18s-rs5 | Gabbr2 | Vav3 | Celf6 | Rgs17 | H2-D1 | Dusp26 | Prr5l | | Ttr | Gm12943 | Gm8292 | Rps19-ps13 |
| Ddah1 | Slc6a9 | Creg1 | Miga2 | Srxn1 | Sept5 | Gabra4 | Vwc2 | Chd3os | Rit2 | H2-K1 | Dync1li2 | Psat1 | | Vps8 | Gm12967 | Gm8451 | Rps19-ps2 |
| Eef2 | Slc7a10 | Csdc2 | Mir124-2hg | Tango2 | Slc1a2 | Gdpd5 | Xylt2 | Crebl2 | Rpl22l1 | Hexa | Efhd1 | Ptgds | | | Gm13192 | Gm8618 | Rps19-ps3 |
| Epas1 | Sparc | Cuta | Mroh1 | Tceal1 | Slc46a3 | Gnal | Zfp804a | Cxx1a | Rwdd2a | Hexb | Efnb3 | Ptprd | | | Gm13202 | Gm8623 | Rps19-ps4 |
| Etnppl | Sparcl1 | Cxxc4 | Mrpl12 | Timm23 | mt-Nd6 | Gpr153 | Zic1 | Cxx1b | Samd14 | Igfbp5 | Elavl3 | Qdpr | | | Gm13456 | Gm8624 | Rps19-ps5 |
| Fads1 | Timp3 | Cyp51 | Mrpl15 | Tkt | mt-Rnr1 | Grm1 | Zmat4 | Cygb | Scg2 | Itgb5 | Elovl1 | Qk | | | Gm13623 | Gm8692 | Rps19-ps6 |
| Fam107a | Tmcc2 | Ddt | Mrpl18 | Tm2d3 | mt-Rnr2 | Grm4 | | Dcaf12l1 | Sdc3 | Itm2b | Elovl7 | Rassf2 | | | Gm13862 | Gm8731 | Rps19-ps7 |
| Fam213a | Tmem176b | Ddx25 | Mrpl34 | Tmed3 | | Hdac9 | | Disp2 | Slc22a17 | Laptm5 | Endod1 | Rcan2 | | | Gm14044 | Gm8805 | Rps19-ps9 |
| Fgfr3 | Trp53inp2 | Ddx3y | Mrpl52 | Tmem246 | | Hlf | | Dner | Smim17 | Lgals3bp | Epb41l3 | Rhog | | | Gm14303 | Gm9396 | Rps20 |
| Foxp2 | Tsc22d3 | Dhrs1 | Mrps6 | Tmem41b | | Ildr2 | | Dnm3 | Stmn1 | Lgmn | Ermn | Rnf13 | | | Gm14407 | Gm9701 | Rps21 |
| Gja1 | Ttyh1 | Dnajc15 | Msrb1 | Tmem60 | | Inpp5a | | Dpysl5 | Tac1 | Ly86 | Evi2a | Rpl36 | | | Gm14439 | Gm9703 | Rps23 |
| Gjb6 | Tuba1a | Dpysl3 | Mtfr1l | Tmx3 | | Kcnc2 | | Dzank1 | Tagln3 | Lyz2 | Fa2h | Rps27a | | | Gm14586 | Gm9843 | Rps23-ps1 |
| Glud1 | Vamp1 | Echs1 | Mypop | Tomm5 | | Kcnd2 | | Elavl2 | Timp2 | Man2b1 | Fgfr2 | Rps4x | | | Gm14633 | Pmch | Rps23-ps2 |
| Glul | Zfhx3 | Eef1b2 | Nap1l2 | Tram1l1 | | Kcnip4 | | Elavl4 | Tmem130 | Mpeg1 | Fnbp1 | Rps9 | | | Gm14859 | RP23-13B8.12 | Rps24 |
| Gm10925 | mt-Atp6 | Eml2 | Nap1l3 | Tsn | | Kitl | | Epm2aip1 | Tmem255a | Npc2 | Frmd8 | Rpsa | | | Gm15198 | RP23-246F14.1 | Rps24-ps2 |
| Gm11407 | mt-Atp8 | Entpd4 | Ndufaf2 | Tuba1b | | Kndc1 | | Fgf12 | Tmem91 | Olfml3 | Fth1 | Rpsa-ps9 | | | Gm15427 | RP23-2N7.4 | Rps24-ps3 |
| Gm11512 | mt-Co1 | Esrra | Ndufaf5 | Tuba1c | | Lef1 | | Fhl1 | Tmsb10 | Plek | Gab1 | S100a16 | | | Gm15464 | RP23-464F18.3 | Rps28 |
| Gm12892 | mt-Co2 | Esrrg | Ndufb7 | Ube2o | | Lhfp | | Flywch2 | Tmx4 | Prdx6 | Gal3st1 | S100b | | | Gm15590 | RP23-58B7.2 | Rps29 |
| Gm13339 | mt-Co3 | Fabp5 | Nova1 | Usp8 | | Lrtm2 | | Fstl5 | Tox2 | RP23-269H21.1 | Gatm | S1pr5 | | | Gm15710 | RP23-70B19.5 | Rps3 |
| Gm13340 | mt-Cytb | Fam134b | Nr1d2 | Vamp7 | | Mb21d2 | | Fxyd6 | Tspyl2 | Rpl18a | Gjb1 | Sccpdh | | | Gm15727 | RP24-142I9.2 | Rps5 |
| Gm13341 | mt-Nd1 | Fam173a | Nrbp1 | Vapb | | Medag | | Gaa | Usp11 | S100a6 | Gjc2 | Scd1 | | | Gm15793 | RP24-502M22.1 | Rps7 |
| Gm28437 | mt-Nd2 | Fbxo3 | Nxph1 | Vegfb | | Megf9 | | Gabrg1 | Usp29 | Serpina3n | Gjc3 | Scd2 | | | Gm15796 | Rpl11 | Tpt1 |
| Gm28438 | mt-Nd3 | Fem1c | Oaz1-ps | Vps39 | | Nell1 | | Galnt16 | Vat1 | Trem2 | Gltp | Sec11c | | | Gm15843 | Rpl12 | Tpt1-ps3 |
| Gm28661 | mt-Nd4 | Fez2 | Olfm3 | Wdr37 | | Nmnat2 | | Gap43 | Vat1l | Tyrobp | Gm15440 | Selenop | | | Gm16005 | Rpl13 | Tpt1-ps5 |
| Gm29216 | mt-Nd4l | Fgf9 | Oprl1 | Wrb | | Nrip3 | | Glra2 | Wdr6 | Vsir | Gm15772 | Sept4 | | | Gm16020 | Rpl13-ps3 | Tpt1-ps6 |
| Gm5514 | mt-Nd5 | Fh1 | Osbpl2 | Wwp1 | | Ntng1 | | Glra3 | Zcchc12 | | Gm5905 | Serpinb1a | | | Gm17511 | Rpl13a | Uba52 |
| Gpld1 | mt-Ta | Fkbp4 | Paics | Zcchc17 | | Osbpl5 | | Gm11223 | Zcchc18 | | Gna12 | Sez6l2 | | | Gm17786 | Rpl17 | Wdr89 |
| Gpr17 | mt-Tc | Fndc5 | Pam16 | Zfas1 | | Patj | | Gm15421 | Zdbf2 | | Gnai2 | Sirt2 | | | Gm19287 | Rpl17-ps10 | Zbed5 |
| Gpr37l1 | mt-Te | Fuca2 | Paqr4 | | | Pcp4 | | Gm16053 | Zwint | | Gng11 | Slain1 | | | Gm19353 | Rpl17-ps4 | |
| Gstm1 | mt-Tf | Fundc1 | Paqr7 | | | Pcp4l1 | | Gm1673 | | | Gpr37 | Slc12a2 | | | Gm19366 | Rpl17-ps5 | |
| Hba-a1 | mt-Th | Fundc2 | Pcdh9 | | | Pdp1 | | Gm9844 | | | Gpr62 | Slc38a2 | | | Gm20430 | Rpl17-ps9 | |
| Hba-a2 | mt-Ti | G6pc3 | Pcx | | | Pitpnc1 | | Gnas | | | Gprc5b | Slc44a1 | | | Gm20568 | Rpl19 | |
| Hbb-bs | mt-Tl1 | Gas6 | Pepd | | | Plcb4 | | Gnl3l | | | Grb14 | Slc48a1 | | | Gm2735 | Rpl27a | |
| Hbb-bt | mt-Tl2 | Glo1 | Pfdn2 | | | Plekha7 | | Gprasp1 | | | Gsn | Sox10 | | | Gm2830 | Rpl29 | |
| Hcn2 | mt-Tm | Glrx | Pgm2 | | | Plekhg1 | | Gprasp2 | | | Hapln2 | Sox2ot | | | Gm29201 | Rpl30 | |
| Hsd11b1 | mt-Tn | Glrx5 | Pgrmc2 | | | Plxdc1 | | Grb10 | | | Il33 | Stmn4 | | | Gm3511 | Rpl30-ps1 | |
| Htra1 | mt-Tp | Gm12231 | Pigk | | | Prkcd | | Grin3a | | | Itgb4 | Syt11 | | | Gm36964 | Rpl30-ps10 | |
| Id3 | mt-Tq | Gm14150 | Plekha6 | | | Prox1 | | Hap1 | | | Jam3 | Taldo1 | | | Gm37713 | Rpl30-ps11 | |
| Jam2 | mt-Tr | Gm14328 | Pmvk | | | Ptpn3 | | Hdac11 | | | Josd2 | Tmbim1 | | | Gm38247 | Rpl30-ps3 | |
| Kcnj10 | mt-Ts2 | Gm14892 | Ppp2r5d | | | Ptpn4 | | Hpcal1 | | | Lamp1 | Tmcc3 | | | Gm4332 | Rpl30-ps8 | |
| Laptm4a | mt-Tt | Gm15920 | Pptc7 | | | Rab3c | | Htr2c | | | Lgi3 | Tmem229a | | | Gm4468 | Rpl30-ps9 | |
| Lcat | mt-Tv | Gm17018 | Prelid3b | | | Ramp3 | | Igsf1 | | | Lpar1 | Tmem63a | | | Gm4705 | Rpl31-ps12 | |
| Ldhb | mt-Tw | Gm21685 | Prmt1 | | | Rap1gap | | Impact | | | Mag | Tmem88b | | | Gm4995 | Rpl34 | |

Table S4

| module | GO Term | Description | P-value | FDR q-value | Enrichment |
|---|---|---|---|---|---|
| brown | GO:0043209 | myelin sheath | 7.429999999999999e-73 | 1.4099999999999997e-69 | 14.71 |
| brown | GO:0044444 | cytoplasmic part | 9.409999999999999e-61 | 8.929999999999999e-58 | 1.86 |
| brown | GO:0097458 | neuron part | 2.7299999999999997e-48 | 1.73e-45 | 3.39 |
| brown | GO:0016020 | membrane | 9.509999999999999e-44 | 4.51e-41 | 1.77 |
| brown | GO:0098800 | inner mitochondrial membrane protein complex | 1.6099999999999997e-42 | 6.099999999999999e-40 | 13.97 |
| brown | GO:0098796 | membrane protein complex | 3.04e-40 | 9.599999999999999e-38 | 3.89 |
| brown | GO:0098803 | respiratory chain complex | 5.1299999999999994e-39 | 1.3899999999999998e-36 | 18.73 |
| brown | GO:0044464 | cell part | 1.56e-38 | 3.7e-36 | 1.28 |
| brown | GO:0044424 | intracellular part | 2.91e-38 | 6.139999999999999e-36 | 1.38 |
| brown | GO:0044455 | mitochondrial membrane part | 7.95e-38 | 1.51e-35 | 9.63 |
| brown | GO:0044456 | synapse part | 5.12e-36 | 8.84e-34 | 4.33 |
| brown | GO:0006810 | transport | 8.81e-36 | 1.34e-31 | 2.37 |
| brown | GO:0031090 | organelle membrane | 2.62e-33 | 4.14e-31 | 3.26 |
| brown | GO:0051234 | establishment of localization | 8.509999999999999e-35 | 6.49e-31 | 2.31 |
| brown | GO:0051179 | localization | 8.09e-34 | 4.11e-30 | 2.17 |
| brown | GO:0043005 | neuron projection | 1.9600000000000001e-31 | 2.86e-29 | 3.36 |
| brown | GO:0070469 | respiratory chain | 3.62e-31 | 4.91e-29 | 19.12 |
| brown | GO:0031966 | mitochondrial membrane | 7.57e-30 | 9.58e-28 | 4.95 |
| brown | GO:0005743 | mitochondrial inner membrane | 8.489999999999999e-30 | 1.0100000000000001e-27 | 6.06 |
| brown | GO:0098798 | mitochondrial protein complex | 2.8300000000000005e-29 | 3.16e-27 | 7.24 |
| brown | GO:0033267 | axon part | 1.21e-28 | 1.27e-26 | 5.37 |
| brown | GO:1990204 | oxidoreductase complex | 1.7699999999999999e-28 | 1.7600000000000002e-26 | 12.5 |
| brown | GO:0044463 | cell projection part | 8.2e-28 | 7.410000000000001e-26 | 2.93 |
| brown | GO:0120038 | plasma membrane bounded cell projection part | 8.2e-28 | 7.779999999999999e-26 | 2.93 |
| brown | GO:0019866 | organelle inner membrane | 9.06e-28 | 7.81e-26 | 5.57 |
| brown | GO:0044429 | mitochondrial part | 1.7399999999999998e-27 | 1.4300000000000003e-25 | 3.93 |
| brown | GO:0043226 | organelle | 1.8800000000000002e-27 | 1.49e-25 | 1.41 |
| brown | GO:0043229 | intracellular organelle | 2.65e-27 | 2.02e-25 | 1.42 |
| brown | GO:0005739 | mitochondrion | 3.0500000000000003e-27 | 2.23e-25 | 2.77 |
| brown | GO:0044297 | cell body | 4.3e-27 | 3.0199999999999997e-25 | 4.1 |
| brown | GO:0044446 | intracellular organelle part | 1.59e-26 | 1.08e-24 | 1.65 |
| brown | GO:0044422 | organelle part | 1.74e-26 | 1.1399999999999999e-24 | 1.62 |
| brown | GO:0005515 | protein binding | 3.03e-27 | 1.36e-23 | 1.53 |
| brown | GO:0120025 | plasma membrane bounded cell projection | 1.13e-24 | 7.15e-23 | 2.62 |
| brown | GO:0044425 | membrane part | 5.43e-24 | 3.33e-22 | 1.61 |
| brown | GO:0005747 | mitochondrial respiratory chain complex I | 1.11e-23 | 6.21e-22 | 18.75 |
| brown | GO:0045271 | respiratory chain complex I | 1.11e-23 | 6.4e-22 | 18.75 |
| brown | GO:0030964 | NADH dehydrogenase complex | 1.11e-23 | 6.59e-22 | 18.75 |
| brown | GO:0042995 | cell projection | 2.0100000000000003e-23 | 1.0900000000000001e-21 | 2.43 |
| brown | GO:0051649 | establishment of localization in cell | 7.2e-25 | 2.7399999999999996e-21 | 2.94 |
| brown | GO:0006091 | generation of precursor metabolites and energy | 1.03e-24 | 3.15e-21 | 6.99 |
| brown | GO:0043227 | membrane-bounded organelle | 9.12e-23 | 4.8100000000000005e-21 | 1.42 |
| brown | GO:0032991 | protein-containing complex | 2.59e-21 | 1.33e-19 | 1.7 |
| brown | GO:0043025 | neuronal cell body | 3.73e-21 | 1.86e-19 | 3.9 |
| brown | GO:0070382 | exocytic vesicle | 4.9e-21 | 2.39e-19 | 7.27 |
| brown | GO:0031982 | vesicle | 5.2600000000000004e-21 | 2.49e-19 | 2.47 |
| brown | GO:0046907 | intracellular transport | 1.8000000000000001e-22 | 4.58e-19 | 3.02 |
| brown | GO:0022900 | electron transport chain | 3.5200000000000004e-22 | 7.67e-19 | 13.44 |
| brown | GO:0045202 | synapse | 1.79e-20 | 8.3e-19 | 3.69 |
| brown | GO:0022904 | respiratory electron transport chain | 4.86e-22 | 9.27e-19 | 14.21 |
| brown | GO:0009117 | nucleotide metabolic process | 9.66e-22 | 1.6400000000000001e-18 | 4.76 |
| brown | GO:0031410 | cytoplasmic vesicle | 4.62e-20 | 2.0899999999999997e-18 | 2.5 |
| brown | GO:0097708 | intracellular vesicle | 5.900000000000001e-20 | 2.61e-18 | 2.49 |
| brown | GO:0006753 | nucleoside phosphate metabolic process | 2.49e-21 | 3.8e-18 | 4.67 |
| brown | GO:0019899 | enzyme binding | 3.76e-21 | 4.23e-18 | 2.25 |
| brown | GO:0015077 | monovalent inorganic cation transmembrane transporter activity | 1.9699999999999998e-21 | 4.42e-18 | 5.08 |
| brown | GO:0015078 | proton transmembrane transporter activity | 3.1699999999999998e-21 | 4.75e-18 | 9.44 |
| brown | GO:0008021 | synaptic vesicle | 2.3e-19 | 9.93e-18 | 7.55 |
| brown | GO:0099503 | secretory vesicle | 2.76e-19 | 1.16e-17 | 4.22 |
| brown | GO:0030133 | transport vesicle | 4.2299999999999994e-19 | 1.74e-17 | 6.01 |
| brown | GO:0009150 | purine ribonucleotide metabolic process | 1.83e-20 | 2.54e-17 | 5.53 |
| brown | GO:0051641 | cellular localization | 2.11e-20 | 2.67e-17 | 2.44 |
| brown | GO:0055086 | nucleobase-containing small molecule metabolic process | 4.42e-20 | 5.19e-17 | 4.24 |
| brown | GO:0009259 | ribonucleotide metabolic process | 7.59e-20 | 8.26e-17 | 5.34 |
| brown | GO:0006163 | purine nucleotide metabolic process | 1.73e-19 | 1.76e-16 | 5.23 |
| brown | GO:0005874 | microtubule | 7.64e-18 | 3.0800000000000003e-16 | 4.28 |
| brown | GO:0019693 | ribose phosphate metabolic process | 5.7000000000000005e-19 | 5.429999999999999e-16 | 5.07 |
| brown | GO:0003824 | catalytic activity | 8.709999999999998e-19 | 7.84e-16 | 1.62 |
| brown | GO:0016818 | hydrolase activity, acting on acid anhydrides, in phosphorus-containing | 1.7000000000000002e-18 | 9.56e-16 | 3.11 |
| brown | GO:0016462 | pyrophosphatase activity | 1.39e-18 | 1.04e-15 | 3.12 |
| brown | GO:0016817 | hydrolase activity, acting on acid anhydrides | 1.7000000000000002e-18 | 1.09e-15 | 3.11 |
| brown | GO:0005737 | cytoplasm | 4.0899999999999997e-17 | 1.6199999999999999e-15 | 1.53 |
| brown | GO:0046034 | ATP metabolic process | 2.19e-18 | 1.97e-15 | 8.82 |
| brown | GO:0072521 | purine-containing compound metabolic process | 2.76e-18 | 2.3399999999999998e-15 | 4.76 |
| brown | GO:1902600 | proton transmembrane transport | 6.15e-18 | 4.94e-15 | 11.47 |
| brown | GO:0009167 | purine ribonucleoside monophosphate metabolic process | 7.190000000000001e-18 | 5.2200000000000004e-15 | 7.68 |
| brown | GO:0005829 | cytosol | 1.39e-16 | 5.39e-15 | 1.9 |
| brown | GO:0017111 | nucleoside-triphosphatase activity | 1.09e-17 | 5.46e-15 | 3.13 |
| brown | GO:0009205 | purine ribonucleoside triphosphate metabolic process | 7.190000000000001e-18 | 5.48e-15 | 7.68 |
| brown | GO:0009126 | purine nucleoside monophosphate metabolic process | 8.96e-18 | 6.21e-15 | 7.62 |
| brown | GO:0009987 | cellular process | 1.28e-17 | 8.479999999999999e-15 | 1.24 |
| brown | GO:0019637 | organophosphate metabolic process | 1.46e-17 | 9.28e-15 | 3.28 |
| brown | GO:0009161 | ribonucleoside monophosphate metabolic process | 1.71e-17 | 1e-14 | 7.45 |
| brown | GO:0009199 | ribonucleoside triphosphate metabolic process | 1.71e-17 | 1.04e-14 | 7.45 |
| brown | GO:0043231 | intracellular membrane-bounded organelle | 3.06e-16 | 1.16e-14 | 1.37 |
| brown | GO:0005488 | binding | 4.1e-17 | 1.85e-14 | 1.27 |
| brown | GO:0006796 | phosphate-containing compound metabolic process | 3.3e-17 | 1.86e-14 | 2.36 |
| brown | GO:0065008 | regulation of biological quality | 3.47e-17 | 1.89e-14 | 1.84 |
| brown | GO:0098793 | presynapse | 5.2e-16 | 1.94e-14 | 5.88 |
| brown | GO:0009144 | purine nucleoside triphosphate metabolic process | 3.93e-17 | 2.07e-14 | 7.24 |
| brown | GO:0019829 | cation-transporting ATPase activity | 9.75e-17 | 3.13e-14 | 11.84 |
| brown | GO:0042625 | ATPase coupled ion transmembrane transporter activity | 9.75e-17 | 3.37e-14 | 11.84 |
| brown | GO:0022853 | active ion transmembrane transporter activity | 9.75e-17 | 3.66e-14 | 11.84 |
| brown | GO:0022890 | inorganic cation transmembrane transporter activity | 9.21e-17 | 3.77e-14 | 3.6 |
| brown | GO:0006793 | phosphorus metabolic process | 8.340000000000001e-17 | 4.24e-14 | 2.33 |
| brown | GO:0051049 | regulation of transport | 8.94e-17 | 4.4e-14 | 2.25 |
| brown | GO:0009123 | nucleoside monophosphate metabolic process | 1.57e-16 | 7.5e-14 | 6.89 |

| | | | | | |
|---|---|---|---|---|---|
| brown | GO:0032879 | regulation of localization | 2.0499999999999998e-16 | 9.47e-14 | 1.96 |
| brown | GO:0016469 | proton-transporting two-sector ATPase complex | 3.8299999999999995e-15 | 1.4e-13 | 15.88 |
| brown | GO:0015318 | inorganic molecular entity transmembrane transporter activity | 7.28e-16 | 2.18e-13 | 3.04 |
| brown | GO:0030426 | growth cone | 7.89e-15 | 2.8300000000000006e-13 | 5.76 |
| brown | GO:0009141 | nucleoside triphosphate metabolic process | 8.32e-16 | 3.73e-13 | 6.49 |
| brown | GO:0098660 | inorganic ion transmembrane transport | 8.93e-16 | 3.8900000000000006e-13 | 3.95 |
| brown | GO:0015672 | monovalent inorganic cation transport | 9.68e-16 | 4.1e-13 | 4.57 |
| brown | GO:0050804 | modulation of chemical synaptic transmission | 1.03e-15 | 4.2299999999999996e-13 | 4.01 |
| brown | GO:0099513 | polymeric cytoskeletal fiber | 1.26e-14 | 4.35e-13 | 3.26 |
| brown | GO:0030427 | site of polarized growth | 1.24e-14 | 4.3700000000000006e-13 | 5.67 |
| brown | GO:0099177 | regulation of trans-synaptic signaling | 1.13e-15 | 4.53e-13 | 4.0 |
| brown | GO:0030424 | axon | 1.56e-14 | 5.290000000000001e-13 | 3.8 |
| brown | GO:0044769 | ATPase activity, coupled to transmembrane movement of ions, rotation | 2.1399999999999997e-15 | 6.02e-13 | 16.38 |
| brown | GO:0019900 | kinase binding | 2.44e-15 | 6.440000000000001e-13 | 2.96 |
| brown | GO:0008324 | cation transmembrane transporter activity | 3.2799999999999997e-15 | 8.179999999999999e-13 | 3.32 |
| brown | GO:0034220 | ion transmembrane transport | 2.36e-15 | 9.220000000000001e-13 | 3.44 |
| brown | GO:0033036 | macromolecule localization | 2.8e-15 | 1.04e-12 | 2.22 |
| brown | GO:0006886 | intracellular protein transport | 2.77e-15 | 1.06e-12 | 3.24 |
| brown | GO:0006754 | ATP biosynthetic process | 3.0899999999999997e-15 | 1.12e-12 | 10.13 |
| brown | GO:0008104 | protein localization | 3.15e-15 | 1.12e-12 | 2.23 |
| brown | GO:0016043 | cellular component organization | 3.4200000000000004e-15 | 1.18e-12 | 1.65 |
| brown | GO:0009206 | purine ribonucleoside triphosphate biosynthetic process | 4.33e-15 | 1.47e-12 | 9.29 |
| brown | GO:0071840 | cellular component organization or biogenesis | 4.7100000000000004e-15 | 1.56e-12 | 1.63 |
| brown | GO:0009145 | purine nucleoside triphosphate biosynthetic process | 5.71e-15 | 1.85e-12 | 9.17 |
| brown | GO:0070069 | cytochrome complex | 5.61e-14 | 1.8700000000000003e-12 | 17.47 |
| brown | GO:0043168 | anion binding | 9.21e-15 | 2.18e-12 | 1.91 |
| brown | GO:0098662 | inorganic cation transmembrane transport | 9.29e-15 | 2.95e-12 | 4.01 |
| brown | GO:0051648 | vesicle localization | 9.52e-15 | 2.96e-12 | 7.1 |
| brown | GO:0009201 | ribonucleoside triphosphate biosynthetic process | 9.829999999999999e-15 | 2.9999999999999997e-12 | 8.95 |
| brown | GO:0099512 | supramolecular fiber | 1.0699999999999999e-13 | 3.4299999999999997e-12 | 3.02 |
| brown | GO:0099081 | supramolecular polymer | 1.0699999999999999e-13 | 3.4900000000000003e-12 | 3.02 |
| brown | GO:0099080 | supramolecular complex | 1.1400000000000002e-13 | 3.6000000000000004e-12 | 3.01 |
| brown | GO:0030425 | dendrite | 1.4099999999999998e-13 | 4.4e-12 | 3.34 |
| brown | GO:0009152 | purine ribonucleotide biosynthetic process | 1.61e-14 | 4.8e-12 | 6.33 |
| brown | GO:0015075 | ion transmembrane transporter activity | 2.27e-14 | 5.1e-12 | 2.83 |
| brown | GO:0098655 | cation transmembrane transport | 2.41e-14 | 7.08e-12 | 3.82 |
| brown | GO:0019901 | protein kinase binding | 3.39e-14 | 7.27e-12 | 2.99 |
| brown | GO:0005215 | transporter activity | 3.84e-14 | 7.85e-12 | 2.5 |
| brown | GO:0017144 | drug metabolic process | 3.04e-14 | 8.439999999999999e-12 | 3.54 |
| brown | GO:0010970 | transport along microtubule | 3.04e-14 | 8.59e-12 | 6.77 |
| brown | GO:0099111 | microtubule-based transport | 3.04e-14 | 8.75e-12 | 6.77 |
| brown | GO:0097060 | synaptic membrane | 4.4799999999999996e-13 | 1.3699999999999999e-11 | 3.97 |
| brown | GO:0022857 | transmembrane transporter activity | 7.19e-14 | 1.41e-11 | 2.61 |
| brown | GO:0043933 | protein-containing complex subunit organization | 5.49e-14 | 1.5e-11 | 2.26 |
| brown | GO:0016192 | vesicle-mediated transport | 5.66e-14 | 1.51e-11 | 2.65 |
| brown | GO:0036094 | small molecule binding | 8.09e-14 | 1.52e-11 | 1.94 |
| brown | GO:0060341 | regulation of cellular localization | 6.98e-14 | 1.83e-11 | 2.79 |
| brown | GO:0009168 | purine ribonucleoside monophosphate biosynthetic process | 7.36e-14 | 1.84e-11 | 8.15 |
| brown | GO:0009260 | ribonucleotide biosynthetic process | 7.16e-14 | 1.85e-11 | 5.97 |
| brown | GO:0009127 | purine nucleoside monophosphate biosynthetic process | 7.36e-14 | 1.87e-11 | 8.15 |
| brown | GO:1902494 | catalytic complex | 6.999999999999999e-13 | 2.1100000000000003e-11 | 2.29 |
| brown | GO:0003954 | NADH dehydrogenase activity | 1.35e-13 | 2.4199999999999998e-11 | 19.06 |
| brown | GO:0006164 | purine nucleotide biosynthetic process | 9.84e-14 | 2.4199999999999998e-11 | 5.9 |
| brown | GO:0042886 | amide transport | 1.15e-13 | 2.78e-11 | 2.45 |
| brown | GO:0032981 | mitochondrial respiratory chain complex I assembly | 1.23e-13 | 2.89e-11 | 13.1 |
| brown | GO:0010257 | NADH dehydrogenase complex assembly | 1.23e-13 | 2.9399999999999997e-11 | 13.1 |
| brown | GO:0015031 | protein transport | 1.29e-13 | 2.99e-11 | 2.48 |
| brown | GO:0015833 | peptide transport | 1.4e-13 | 3.2e-11 | 2.46 |
| brown | GO:0009142 | nucleoside triphosphate biosynthetic process | 1.4800000000000003e-13 | 3.31e-11 | 7.89 |
| brown | GO:0009156 | ribonucleoside monophosphate biosynthetic process | 1.85e-13 | 3.98e-11 | 7.8 |
| brown | GO:0072522 | purine-containing compound biosynthetic process | 1.83e-13 | 3.98e-11 | 5.75 |
| brown | GO:0055085 | transmembrane transport | 1.8199999999999998e-13 | 4.01e-11 | 2.71 |
| brown | GO:0051640 | organelle localization | 1.98e-13 | 4.2e-11 | 3.74 |
| brown | GO:0045184 | establishment of protein localization | 2.04e-13 | 4.2700000000000005e-11 | 2.41 |
| brown | GO:0046390 | ribose phosphate biosynthetic process | 2.13e-13 | 4.39e-11 | 5.72 |
| brown | GO:0030705 | cytoskeleton-dependent intracellular transport | 2.5599999999999997e-13 | 5.19e-11 | 6.19 |
| brown | GO:0071705 | nitrogen compound transport | 2.6400000000000004e-13 | 5.289999999999999e-11 | 2.25 |
| brown | GO:0051656 | establishment of organelle localization | 4.33e-13 | 8.56e-11 | 4.4 |
| brown | GO:0050136 | NADH dehydrogenase (quinone) activity | 5.91e-13 | 9.49e-11 | 20.23 |
| brown | GO:0008137 | NADH dehydrogenase (ubiquinone) activity | 5.91e-13 | 9.85e-11 | 20.23 |
| brown | GO:0023052 | signaling | 5.04e-13 | 9.859999999999999e-11 | 3.64 |
| brown | GO:0036442 | proton-exporting ATPase activity | 5.809999999999999e-13 | 1.01e-10 | 15.14 |
| brown | GO:0022607 | cellular component assembly | 6.1e-13 | 1.18e-10 | 1.98 |
| brown | GO:0071702 | organic substance transport | 6.24e-13 | 1.19e-10 | 2.09 |
| brown | GO:0000166 | nucleotide binding | 8.229999999999999e-13 | 1.23e-10 | 1.99 |
| brown | GO:1901265 | nucleoside phosphate binding | 8.229999999999999e-13 | 1.28e-10 | 1.99 |
| brown | GO:0009055 | electron transfer activity | 1.03e-12 | 1.5e-10 | 9.58 |
| brown | GO:0009124 | nucleoside monophosphate biosynthetic process | 8.32e-13 | 1.5699999999999997e-10 | 7.26 |
| brown | GO:0046961 | proton-transporting ATPase activity, rotational mechanism | 1.28e-12 | 1.8000000000000002e-10 | 19.21 |
| brown | GO:0051650 | establishment of vesicle localization | 9.75e-13 | 1.8100000000000002e-10 | 6.8 |
| brown | GO:0097367 | carbohydrate derivative binding | 1.6e-12 | 2.18e-10 | 1.96 |
| brown | GO:0033178 | proton-transporting two-sector ATPase complex, catalytic domain | 7.83e-12 | 2.3199999999999997e-10 | 24.19 |
| brown | GO:0015405 | P-P-bond-hydrolysis-driven transmembrane transporter activity | 2.0399999999999998e-12 | 2.62e-10 | 6.57 |
| brown | GO:0015399 | primary active transmembrane transporter activity | 2.0399999999999998e-12 | 2.7000000000000005e-10 | 6.57 |
| brown | GO:0065003 | protein-containing complex assembly | 1.4999999999999999e-12 | 2.75e-10 | 2.27 |
| brown | GO:0032553 | ribonucleotide binding | 2.4e-12 | 2.99e-10 | 2.04 |
| brown | GO:0031625 | ubiquitin protein ligase binding | 3.0499999999999997e-12 | 3.71e-10 | 4.11 |
| brown | GO:0033108 | mitochondrial respiratory chain complex assembly | 2.41e-12 | 4.3700000000000004e-10 | 9.14 |
| brown | GO:0044281 | small molecule metabolic process | 2.59e-12 | 4.6399999999999995e-10 | 2.19 |
| brown | GO:0016787 | hydrolase activity | 4.05e-12 | 4.79e-10 | 1.84 |
| brown | GO:0098590 | plasma membrane region | 2.05e-11 | 5.98e-10 | 2.38 |
| brown | GO:0042626 | ATPase activity, coupled to transmembrane movement of substances | 5.740000000000001e-12 | 6.62e-10 | 6.61 |
| brown | GO:0007268 | chemical synaptic transmission | 4.669999999999999e-12 | 8.18e-10 | 4.4 |
| brown | GO:0030276 | clathrin binding | 7.3e-12 | 8.210000000000001e-10 | 10.28 |
| brown | GO:0098916 | anterograde trans-synaptic signaling | 4.669999999999999e-12 | 8.269999999999999e-10 | 4.4 |
| brown | GO:0032555 | purine ribonucleotide binding | 7.94e-12 | 8.71e-10 | 2.02 |
| brown | GO:1901564 | organonitrogen compound metabolic process | 5.0599999999999999e-12 | 8.77e-10 | 1.59 |
| brown | GO:0051050 | positive regulation of transport | 5.43e-12 | 9.300000000000001e-10 | 2.43 |
| brown | GO:0009165 | nucleotide biosynthetic process | 5.6999999999999995e-12 | 9.66e-10 | 4.82 |
| brown | GO:0035639 | purine ribonucleoside triphosphate binding | 1.05e-11 | 1.1199999999999999e-09 | 2.03 |
| brown | GO:0017076 | purine nucleotide binding | 1.1499999999999999e-11 | 1.2e-09 | 2.0 |
| brown | GO:0016887 | ATPase activity | 1.2699999999999999e-11 | 1.3e-09 | 3.43 |
| brown | GO:0044389 | ubiquitin-like protein ligase binding | 1.39e-11 | 1.3900000000000002e-09 | 3.89 |
| brown | GO:0044877 | protein-containing complex binding | 1.43e-11 | 1.4000000000000001e-09 | 2.31 |

| | | | | | |
|---|---|---|---|---|---|
| brown | GO:0043492 | ATPase activity, coupled to movement of substances | 1.56e-11 | 1.5e-09 | 5.96 |
| brown | GO:0099003 | vesicle-mediated transport in synapse | 1.17e-11 | 1.95e-09 | 7.22 |
| brown | GO:1901135 | carbohydrate derivative metabolic process | 1.2099999999999999e-11 | 2e-09 | 2.81 |
| brown | GO:0006812 | cation transport | 1.24e-11 | 2.04e-09 | 2.78 |
| brown | GO:1901293 | nucleoside phosphate biosynthetic process | 1.31e-11 | 2.12e-09 | 4.66 |
| brown | GO:0099537 | trans-synaptic signaling | 1.63e-11 | 2.59e-09 | 4.2 |
| brown | GO:0099536 | synaptic signaling | 1.63e-11 | 2.62e-09 | 4.2 |
| brown | GO:0006811 | ion transport | 1.7699999999999998e-11 | 2.7500000000000002e-09 | 2.33 |
| brown | GO:0007017 | microtubule-based process | 1.76e-11 | 2.7699999999999997e-09 | 2.97 |
| brown | GO:0044433 | cytoplasmic vesicle part | 1.0500000000000001e-10 | 3.02e-09 | 3.14 |
| brown | GO:0022804 | active transmembrane transporter activity | 3.36e-11 | 3.15e-09 | 3.68 |
| brown | GO:0072524 | pyridine-containing compound metabolic process | 2.32e-11 | 3.5700000000000003e-09 | 6.53 |
| brown | GO:0006996 | organelle organization | 2.7e-11 | 4.120000000000001e-09 | 1.92 |
| brown | GO:0072384 | organelle transport along microtubule | 2.93e-11 | 4.42e-09 | 8.6 |
| brown | GO:0009060 | aerobic respiration | 3.62e-11 | 5.41e-09 | 13.1 |
| brown | GO:0005751 | mitochondrial respiratory chain complex IV | 1.9199999999999998e-10 | 5.45e-09 | 23.29 |
| brown | GO:0098588 | bounding membrane of organelle | 2.1499999999999998e-10 | 6e-09 | 2.52 |
| brown | GO:0005753 | mitochondrial proton-transporting ATP synthase complex | 2.4e-10 | 6.42e-09 | 18.5 |
| brown | GO:0045277 | respiratory chain complex IV | 2.4e-10 | 6.520000000000001e-09 | 18.5 |
| brown | GO:0044430 | cytoskeletal part | 2.4799999999999997e-10 | 6.55e-09 | 2.05 |
| brown | GO:0045259 | proton-transporting ATP synthase complex | 2.4e-10 | 6.609999999999999e-09 | 18.5 |
| brown | GO:0051128 | regulation of cellular component organization | 4.73e-11 | 7.0099999999999996e-09 | 1.8 |
| brown | GO:0055114 | oxidation-reduction process | 5.18e-11 | 7.6e-09 | 2.54 |
| brown | GO:0008092 | cytoskeletal protein binding | 9.000000000000001e-11 | 8.26e-09 | 2.41 |
| brown | GO:0046496 | nicotinamide nucleotide metabolic process | 6.54e-11 | 9.490000000000001e-09 | 6.57 |
| brown | GO:0005856 | cytoskeleton | 4.8e-10 | 1.25e-08 | 2.05 |
| brown | GO:0048167 | regulation of synaptic plasticity | 8.879999999999999e-11 | 1.28e-08 | 4.45 |
| brown | GO:0019362 | pyridine nucleotide metabolic process | 9.340000000000001e-11 | 1.33e-08 | 6.44 |
| brown | GO:0017158 | regulation of calcium ion-dependent exocytosis | 9.599999999999999e-11 | 1.3600000000000001e-08 | 8.73 |
| brown | GO:0043167 | ion binding | 1.73e-10 | 1.56e-08 | 1.46 |
| brown | GO:0006101 | citrate metabolic process | 1.2399999999999999e-10 | 1.73e-08 | 11.98 |
| brown | GO:0005886 | plasma membrane | 6.84e-10 | 1.75e-08 | 1.55 |
| brown | GO:0090662 | ATP hydrolysis coupled transmembrane transport | 2.0799999999999998e-10 | 2.81e-08 | 13.25 |
| brown | GO:0016655 | oxidoreductase activity, acting on NAD(P)H, quinone or similar compou | 3.21e-10 | 2.8300000000000002e-08 | 12.81 |
| brown | GO:0015988 | energy coupled proton transmembrane transport, against electrochemi | 2.0799999999999998e-10 | 2.84e-08 | 13.25 |
| brown | GO:0099132 | ATP hydrolysis coupled cation transmembrane transport | 2.0799999999999998e-10 | 2.8599999999999998e-08 | 13.25 |
| brown | GO:0042623 | ATPase activity, coupled | 3.32e-10 | 2.8700000000000002e-08 | 3.55 |
| brown | GO:0099131 | ATP hydrolysis coupled ion transmembrane transport | 2.0799999999999998e-10 | 2.89e-08 | 13.25 |
| brown | GO:0051186 | cofactor metabolic process | 2.3199999999999997e-10 | 3.11e-08 | 3.23 |
| brown | GO:0098805 | whole membrane | 1.36e-09 | 3.4500000000000005e-08 | 2.72 |
| brown | GO:0048471 | perinuclear region of cytoplasm | 1.61e-09 | 4.0199999999999996e-08 | 2.67 |
| brown | GO:0098693 | regulation of synaptic vesicle cycle | 3.1399999999999995e-10 | 4.16e-08 | 8.89 |
| brown | GO:0120035 | regulation of plasma membrane bounded cell projection organization | 3.6799999999999997e-10 | 4.8399999999999997e-08 | 2.56 |
| brown | GO:0042391 | regulation of membrane potential | 3.77e-10 | 4.92e-08 | 3.18 |
| brown | GO:0008144 | drug binding | 5.93e-10 | 5.03e-08 | 1.96 |
| brown | GO:0006732 | coenzyme metabolic process | 4.06e-10 | 5.24e-08 | 3.91 |
| brown | GO:0023051 | regulation of signaling | 4.51e-10 | 5.78e-08 | 1.66 |
| brown | GO:0006099 | tricarboxylic acid cycle | 4.84e-10 | 6.15e-08 | 12.4 |
| brown | GO:0030054 | cell junction | 2.51e-09 | 6.18e-08 | 2.17 |
| brown | GO:0031344 | regulation of cell projection organization | 6.01e-10 | 7.58e-08 | 2.53 |
| brown | GO:0010646 | regulation of cell communication | 6.14e-10 | 7.68e-08 | 1.66 |
| brown | GO:0072350 | tricarboxylic acid metabolic process | 7.26e-10 | 9e-08 | 10.48 |
| brown | GO:0015986 | ATP synthesis coupled proton transport | 8.68e-10 | 1.06e-07 | 16.55 |
| brown | GO:0015985 | energy coupled proton transport, down electrochemical gradient | 8.68e-10 | 1.0699999999999999e-07 | 16.55 |
| brown | GO:0006733 | oxidoreduction coenzyme metabolic process | 9.070000000000001e-10 | 1.1e-07 | 5.67 |
| brown | GO:0097479 | synaptic vesicle localization | 1e-09 | 1.2e-07 | 7.49 |
| brown | GO:0008088 | axo-dendritic transport | 1.1e-09 | 1.3e-07 | 8.15 |
| brown | GO:0043269 | regulation of ion transport | 1.19e-09 | 1.4e-07 | 2.57 |
| brown | GO:0009166 | nucleotide catabolic process | 1.2e-09 | 1.41e-07 | 6.82 |
| brown | GO:0045333 | cellular respiration | 1.3300000000000002e-09 | 1.55e-07 | 8.91 |
| brown | GO:0007267 | cell-cell signaling | 1.92e-09 | 2.22e-07 | 3.31 |
| brown | GO:0099643 | signal release from synapse | 2.18e-09 | 2.5e-07 | 7.76 |
| brown | GO:0007018 | microtubule-based movement | 2.23e-09 | 2.53e-07 | 3.98 |
| brown | GO:1901137 | carbohydrate derivative biosynthetic process | 2.6300000000000002e-09 | 2.9699999999999997e-07 | 3.35 |
| brown | GO:0015991 | ATP hydrolysis coupled proton transport | 2.8199999999999998e-09 | 3.16e-07 | 12.48 |
| brown | GO:0051130 | positive regulation of cellular component organization | 3.07e-09 | 3.42e-07 | 2.09 |
| brown | GO:0090407 | organophosphate biosynthetic process | 3.71e-09 | 4.1e-07 | 3.16 |
| brown | GO:0009056 | catabolic process | 4.26e-09 | 4.6799999999999996e-07 | 1.93 |
| brown | GO:0044306 | neuron projection terminus | 2.61e-08 | 6.36e-07 | 5.21 |
| brown | GO:0046434 | organophosphate catabolic process | 6.2e-09 | 6.75e-07 | 4.82 |
| brown | GO:0034622 | cellular protein-containing complex assembly | 6.27e-09 | 6.77e-07 | 2.5 |
| brown | GO:0015631 | tubulin binding | 8.240000000000001e-09 | 6.86e-07 | 3.35 |
| brown | GO:0010975 | regulation of neuron projection development | 7.0099999999999996e-09 | 7.520000000000001e-07 | 2.65 |
| brown | GO:0042221 | response to chemical | 7.33e-09 | 7.809999999999999e-07 | 1.7 |
| brown | GO:0044437 | vacuolar part | 3.41e-08 | 8.179999999999999e-07 | 4.37 |
| brown | GO:1901292 | nucleoside phosphate catabolic process | 8.199999999999999e-09 | 8.68e-07 | 6.01 |
| brown | GO:0007154 | cell communication | 8.62e-09 | 9.06e-07 | 2.63 |
| brown | GO:0005750 | mitochondrial respiratory chain complex III | 4.18e-08 | 9.81e-07 | 23.29 |
| brown | GO:0045275 | respiratory chain complex III | 4.18e-08 | 9.93e-07 | 23.29 |
| brown | GO:0051960 | regulation of nervous system development | 9.670000000000002e-09 | 1.01e-06 | 2.18 |
| brown | GO:0015002 | heme-copper terminal oxidase activity | 1.46e-08 | 1.15e-06 | 15.53 |
| brown | GO:0016676 | oxidoreductase activity, acting on a heme group of donors, oxygen as | 1.46e-08 | 1.17e-06 | 15.53 |
| brown | GO:0004129 | cytochrome-c oxidase activity | 1.46e-08 | 1.19e-06 | 15.53 |
| brown | GO:0032386 | regulation of intracellular transport | 1.16e-08 | 1.2e-06 | 3.15 |
| brown | GO:0098794 | postsynapse | 5.59e-08 | 1.29e-06 | 3.9 |
| brown | GO:0045664 | regulation of neuron differentiation | 1.29e-08 | 1.32e-06 | 2.39 |
| brown | GO:0050905 | neuromuscular process | 1.31e-08 | 1.34e-06 | 5.45 |
| brown | GO:0033177 | proton-transporting two-sector ATPase complex, proton-transporting do | 6.29e-08 | 1.44e-06 | 13.31 |
| brown | GO:0015980 | energy derivation by oxidation of organic compounds | 1.51e-08 | 1.53e-06 | 5.4 |
| brown | GO:0019904 | protein domain specific binding | 2.0299999999999996e-08 | 1.58e-06 | 2.31 |
| brown | GO:0032590 | dendrite membrane | 7.84e-08 | 1.77e-06 | 9.19 |
| brown | GO:0016491 | oxidoreductase activity | 2.46e-08 | 1.85e-06 | 2.34 |
| brown | GO:0016675 | oxidoreductase activity, acting on a heme group of donors | 2.46e-08 | 1.87e-06 | 14.71 |
| brown | GO:0032940 | secretion by cell | 1.98e-08 | 1.99e-06 | 3.08 |
| brown | GO:0010769 | regulation of cell morphogenesis involved in differentiation | 2.09e-08 | 2.1000000000000002e-06 | 3.21 |
| brown | GO:0044248 | cellular catabolic process | 2.21e-08 | 2.2e-06 | 1.96 |
| brown | GO:0023061 | signal release | 2.46e-08 | 2.43e-06 | 4.67 |
| brown | GO:0032559 | adenyl ribonucleotide binding | 3.44e-08 | 2.54e-06 | 1.9 |
| brown | GO:0005875 | microtubule associated complex | 1.1900000000000001e-07 | 2.66e-06 | 4.46 |
| brown | GO:0098589 | membrane region | 1.23e-07 | 2.72e-06 | 2.97 |
| brown | GO:0005794 | Golgi apparatus | 1.37e-07 | 2.99e-06 | 1.94 |
| brown | GO:0031256 | leading edge membrane | 1.48e-07 | 3.19e-06 | 4.4 |
| brown | GO:0051588 | regulation of neurotransmitter transport | 3.41e-08 | 3.35e-06 | 5.12 |
| brown | GO:0030554 | adenyl nucleotide binding | 4.6200000000000003e-08 | 3.35e-06 | 1.89 |

| | | | | | |
|---|---|---|---|---|---|
| brown | GO:0019725 | cellular homeostasis | 3.75e-08 | 3.67e-06 | 2.31 |
| brown | GO:0034404 | nucleobase-containing small molecule biosynthetic process | 3.88e-08 | 3.77e-06 | 5.08 |
| brown | GO:0044237 | cellular metabolic process | 4.08e-08 | 3.94e-06 | 1.31 |
| brown | GO:0045121 | membrane raft | 1.85e-07 | 3.95e-06 | 2.98 |
| brown | GO:0098857 | membrane microdomain | 1.97e-07 | 4.07e-06 | 2.97 |
| brown | GO:0014069 | postsynaptic density | 1.97e-07 | 4.11e-06 | 3.3 |
| brown | GO:0099572 | postsynaptic specialization | 1.97e-07 | 4.16e-06 | 3.3 |
| brown | GO:0006120 | mitochondrial electron transport, NADH to ubiquinone | 4.38e-08 | 4.2000000000000004e-06 | 17.47 |
| brown | GO:0030030 | cell projection organization | 4.54e-08 | 4.33e-06 | 2.23 |
| brown | GO:0005524 | ATP binding | 6.21e-08 | 4.44e-06 | 1.91 |
| brown | GO:1901575 | organic substance catabolic process | 4.7800000000000005e-08 | 4.53e-06 | 1.95 |
| brown | GO:0032589 | neuron projection membrane | 2.28e-07 | 4.66e-06 | 6.55 |
| brown | GO:0008022 | protein C-terminus binding | 6.67e-08 | 4.69e-06 | 3.6 |
| brown | GO:0006887 | exocytosis | 5.1799999999999994e-08 | 4.87e-06 | 4.26 |
| brown | GO:0030672 | synaptic vesicle membrane | 2.46e-07 | 4.92e-06 | 7.25 |
| brown | GO:0099501 | exocytic vesicle membrane | 2.46e-07 | 4.97e-06 | 7.25 |
| brown | GO:0042981 | regulation of apoptotic process | 5.68e-08 | 5.31e-06 | 1.87 |
| brown | GO:0030658 | transport vesicle membrane | 2.73e-07 | 5.39e-06 | 6.45 |
| brown | GO:0046933 | proton-transporting ATP synthase activity, rotational mechanism | 8e-08 | 5.54e-06 | 16.3 |
| brown | GO:0043204 | perikaryon | 2.9199999999999997e-07 | 5.72e-06 | 4.43 |
| brown | GO:0001505 | regulation of neurotransmitter levels | 6.92e-08 | 6.44e-06 | 3.38 |
| brown | GO:0031175 | neuron projection development | 7.299999999999999e-08 | 6.7e-06 | 3.28 |
| brown | GO:2000300 | regulation of synaptic vesicle exocytosis | 7.28e-08 | 6.73e-06 | 10.84 |
| brown | GO:0032880 | regulation of protein localization | 7.480000000000001e-08 | 6.83e-06 | 2.12 |
| brown | GO:0005905 | clathrin-coated pit | 3.6700000000000004e-07 | 7.11e-06 | 6.99 |
| brown | GO:0016651 | oxidoreductase activity, acting on NAD(P)H | 1.14e-07 | 7.79e-06 | 6.31 |
| brown | GO:0061024 | membrane organization | 9.050000000000001e-08 | 8.21e-06 | 2.64 |
| brown | GO:0043067 | regulation of programmed cell death | 9.449999999999999e-08 | 8.53e-06 | 1.85 |
| brown | GO:0003924 | GTPase activity | 1.34e-07 | 8.96e-06 | 3.27 |
| brown | GO:0046903 | secretion | 1e-07 | 9e-06 | 2.67 |
| brown | GO:0019646 | aerobic electron transport chain | 1.0199999999999999e-07 | 9.05e-06 | 20.96 |
| brown | GO:0006123 | mitochondrial electron transport, cytochrome c to oxygen | 1.0199999999999999e-07 | 9.1e-06 | 20.96 |
| brown | GO:0044459 | plasma membrane part | 5.2e-07 | 9.96e-06 | 1.57 |
| brown | GO:0042802 | identical protein binding | 1.55e-07 | 1.0199999999999999e-05 | 1.74 |
| brown | GO:1902495 | transmembrane transporter complex | 5.520000000000001e-07 | 1.05e-05 | 3.04 |
| brown | GO:0010941 | regulation of cell death | 1.2199999999999998e-07 | 1.0800000000000002e-05 | 1.8 |
| brown | GO:0016999 | antibiotic metabolic process | 1.24e-07 | 1.09e-05 | 4.99 |
| brown | GO:0030003 | cellular cation homeostasis | 1.32e-07 | 1.15e-05 | 2.49 |
| brown | GO:0044325 | ion channel binding | 1.87e-07 | 1.22e-05 | 4.57 |
| brown | GO:0031346 | positive regulation of cell projection organization | 1.42e-07 | 1.23e-05 | 2.72 |
| brown | GO:0006734 | NADH metabolic process | 1.44e-07 | 1.24e-05 | 12.15 |
| brown | GO:0045211 | postsynaptic membrane | 6.6e-07 | 1.24e-05 | 3.28 |
| brown | GO:0006090 | pyruvate metabolic process | 1.58e-07 | 1.3500000000000001e-05 | 6.76 |
| brown | GO:0043270 | positive regulation of ion transport | 1.68e-07 | 1.4300000000000002e-05 | 3.14 |
| brown | GO:0010976 | positive regulation of neuron projection development | 1.85e-07 | 1.57e-05 | 2.98 |
| brown | GO:0032990 | cell part morphogenesis | 1.9e-07 | 1.59e-05 | 3.21 |
| brown | GO:1902803 | regulation of synaptic vesicle transport | 1.8899999999999999e-07 | 1.59e-05 | 9.83 |
| brown | GO:1990351 | transporter complex | 8.590000000000001e-07 | 1.6e-05 | 2.97 |
| brown | GO:1901566 | organonitrogen compound biosynthetic process | 2.07e-07 | 1.7199999999999998e-05 | 2.18 |
| brown | GO:0006122 | mitochondrial electron transport, ubiquinol to cytochrome c | 2.19e-07 | 1.82e-05 | 19.06 |
| brown | GO:0043648 | dicarboxylic acid metabolic process | 2.25e-07 | 1.85e-05 | 5.5 |
| brown | GO:0048168 | regulation of neuronal synaptic plasticity | 2.28e-07 | 1.87e-05 | 6.55 |
| brown | GO:0006836 | neurotransmitter transport | 2.35e-07 | 1.91e-05 | 4.5 |
| brown | GO:0098563 | intrinsic component of synaptic vesicle membrane | 1.08e-06 | 2e-05 | 9.64 |
| brown | GO:0006873 | cellular ion homeostasis | 2.52e-07 | 2.0300000000000002e-05 | 2.43 |
| brown | GO:0048812 | neuron projection morphogenesis | 2.5e-07 | 2.0300000000000002e-05 | 3.35 |
| brown | GO:0050770 | regulation of axonogenesis | 2.7e-07 | 2.14e-05 | 3.71 |
| brown | GO:0043523 | regulation of neuron apoptotic process | 2.7e-07 | 2.15e-05 | 3.34 |
| brown | GO:0006897 | endocytosis | 2.68e-07 | 2.15e-05 | 2.93 |
| brown | GO:0015630 | microtubule cytoskeleton | 1.51e-06 | 2.7600000000000003e-05 | 3.77 |
| brown | GO:1903827 | regulation of cellular protein localization | 3.6299999999999995e-07 | 2.8699999999999996e-05 | 2.57 |
| brown | GO:0007610 | behavior | 3.7700000000000005e-07 | 2.9600000000000005e-05 | 2.33 |
| brown | GO:0097480 | establishment of synaptic vesicle localization | 3.85e-07 | 2.9999999999999997e-05 | 6.26 |
| brown | GO:0120039 | plasma membrane bounded cell projection morphogenesis | 3.87e-07 | 2.9999999999999997e-05 | 3.28 |
| brown | GO:0048489 | synaptic vesicle transport | 3.85e-07 | 3.01e-05 | 6.26 |
| brown | GO:1903305 | regulation of regulated secretory pathway | 3.94e-07 | 3.0299999999999998e-05 | 4.9 |
| brown | GO:0022604 | regulation of cell morphogenesis | 4.1e-07 | 3.14e-05 | 2.52 |
| brown | GO:0050767 | regulation of neurogenesis | 4.15e-07 | 3.16e-05 | 2.08 |
| brown | GO:0017157 | regulation of exocytosis | 4.43e-07 | 3.3600000000000004e-05 | 3.9 |
| brown | GO:0060284 | regulation of cell development | 4.61e-07 | 3.48e-05 | 1.99 |
| brown | GO:0008017 | microtubule binding | 5.77e-07 | 3.71e-05 | 3.42 |
| brown | GO:0048858 | cell projection morphogenesis | 5.14e-07 | 3.84e-05 | 3.22 |
| brown | GO:0007010 | cytoskeleton organization | 5.13e-07 | 3.85e-05 | 2.16 |
| brown | GO:0043679 | axon terminus | 2.13e-06 | 3.86e-05 | 4.94 |
| brown | GO:0000149 | SNARE binding | 6.389999999999999e-07 | 4.05e-05 | 4.72 |
| brown | GO:0055080 | cation homeostasis | 5.48e-07 | 4.07e-05 | 2.3 |
| brown | GO:0051716 | cellular response to stimulus | 5.57e-07 | 4.1200000000000005e-05 | 1.58 |
| brown | GO:0031253 | cell projection membrane | 2.33e-06 | 4.18e-05 | 2.96 |
| brown | GO:0012506 | vesicle membrane | 2.43e-06 | 4.31e-05 | 3.36 |
| brown | GO:0046928 | regulation of neurotransmitter secretion | 6.32e-07 | 4.66e-05 | 5.99 |
| brown | GO:0065009 | regulation of molecular function | 6.37e-07 | 4.6700000000000003e-05 | 1.59 |
| brown | GO:0055082 | cellular chemical homeostasis | 6.45e-07 | 4.71e-05 | 2.28 |
| brown | GO:1903829 | positive regulation of cellular protein localization | 7.259999999999999e-07 | 5.27e-05 | 3.08 |
| brown | GO:0045261 | proton-transporting ATP synthase complex, catalytic core F(1) | 3.25e-06 | 5.65e-05 | 27.95 |
| brown | GO:0000275 | mitochondrial proton-transporting ATP synthase complex, catalytic core | 3.25e-06 | 5.7100000000000006e-05 | 27.95 |
| brown | GO:0050801 | ion homeostasis | 7.970000000000001e-07 | 5.7600000000000004e-05 | 2.19 |
| brown | GO:0033176 | proton-transporting V-type ATPase complex | 3.36e-06 | 5.7999999999999994e-05 | 13.1 |
| brown | GO:0051668 | localization within membrane | 8.079999999999999e-07 | 5.8099999999999996e-05 | 4.64 |
| brown | GO:0015662 | ATPase activity, coupled to transmembrane movement of ions, phosph | 9.41e-07 | 5.88e-05 | 8.27 |
| brown | GO:0034762 | regulation of transmembrane transport | 8.64e-07 | 6.19e-05 | 2.41 |
| brown | GO:0070201 | regulation of establishment of protein localization | 8.86e-07 | 6.31e-05 | 2.23 |
| brown | GO:0030659 | cytoplasmic vesicle membrane | 3.78e-06 | 6.47e-05 | 3.53 |
| brown | GO:0009108 | coenzyme biosynthetic process | 9.85e-07 | 6.979999999999999e-05 | 4.3 |
| brown | GO:0098771 | inorganic ion homeostasis | 1e-06 | 7.09e-05 | 2.24 |
| brown | GO:0008289 | lipid binding | 1.25e-06 | 7.730000000000001e-05 | 2.2 |
| brown | GO:0045055 | regulated exocytosis | 1.11e-06 | 7.829999999999999e-05 | 5.22 |
| brown | GO:0019905 | syntaxin binding | 1.36e-06 | 8.290000000000001e-05 | 5.59 |
| brown | GO:0035418 | protein localization to synapse | 1.19e-06 | 8.31e-05 | 8.06 |
| brown | GO:0006096 | glycolytic process | 1.19e-06 | 8.340000000000001e-05 | 8.06 |
| brown | GO:0031594 | neuromuscular junction | 5.19e-06 | 8.8e-05 | 5.41 |
| brown | GO:0005244 | voltage-gated ion channel activity | 1.49e-06 | 8.83e-05 | 3.61 |
| brown | GO:0022832 | voltage-gated channel activity | 1.49e-06 | 8.950000000000001e-05 | 3.61 |
| brown | GO:0006107 | oxaloacetate metabolic process | 1.32e-06 | 9.17e-05 | 14.97 |
| brown | GO:0051952 | regulation of amine transport | 1.42e-06 | 9.719999999999999e-05 | 4.44 |

| | | | | | |
|---|---|---|---|---|---|
| brown | GO:0098657 | import into cell | 1.41e-06 | 9.719999999999999e-05 | 2.6 |
| brown | GO:0006757 | ATP generation from ADP | 1.5e-06 | 0.000102 | 7.86 |
| brown | GO:0006875 | cellular metal ion homeostasis | 1.5e-06 | 0.000103 | 2.42 |
| brown | GO:0008152 | metabolic process | 1.63e-06 | 0.00011 | 1.25 |
| brown | GO:0051962 | positive regulation of nervous system development | 1.77e-06 | 0.00011899999999999999 | 2.27 |
| brown | GO:0101031 | chaperone complex | 7.42e-06 | 0.000125 | 11.65 |
| brown | GO:0034702 | ion channel complex | 8.11e-06 | 0.000135 | 2.83 |
| brown | GO:0035254 | glutamate receptor binding | 2.48e-06 | 0.000145 | 5.82 |
| brown | GO:0051646 | mitochondrion localization | 2.33e-06 | 0.000156 | 7.49 |
| brown | GO:0033180 | proton-transporting V-type ATPase, V1 domain | 9.52e-06 | 0.000157 | 23.29 |
| brown | GO:0022839 | ion gated channel activity | 2.83e-06 | 0.00016299999999999998 | 2.85 |
| brown | GO:1901214 | regulation of neuron death | 2.45e-06 | 0.000164 | 2.74 |
| brown | GO:1903421 | regulation of synaptic vesicle recycling | 2.5e-06 | 0.000166 | 10.63 |
| brown | GO:0070507 | regulation of microtubule cytoskeleton organization | 2.61e-06 | 0.000173 | 3.48 |
| brown | GO:0030285 | integral component of synaptic vesicle membrane | 1.06e-05 | 0.000173 | 11.03 |
| brown | GO:0006165 | nucleoside diphosphate phosphorylation | 2.63e-06 | 0.000174 | 6.47 |
| brown | GO:0051188 | cofactor biosynthetic process | 2.73e-06 | 0.00018 | 3.61 |
| brown | GO:0042866 | pyruvate biosynthetic process | 2.87e-06 | 0.000188 | 7.31 |
| brown | GO:0070887 | cellular response to chemical stimulus | 2.92e-06 | 0.00019099999999999998 | 1.73 |
| brown | GO:0050772 | positive regulation of axonogenesis | 3.08e-06 | 0.0002 | 4.78 |
| brown | GO:0005783 | endoplasmic reticulum | 1.24e-05 | 0.00020099999999999998 | 1.68 |
| brown | GO:0030117 | membrane coat | 1.29e-05 | 0.000207 | 5.46 |
| brown | GO:0044093 | positive regulation of molecular function | 3.25e-06 | 0.00021 | 1.76 |
| brown | GO:1903530 | regulation of secretion by cell | 3.28e-06 | 0.00021099999999999998 | 2.1 |
| brown | GO:0047497 | mitochondrion transport along microtubule | 3.36e-06 | 0.00021400000000000002 | 13.1 |
| brown | GO:0045666 | positive regulation of neuron differentiation | 3.39e-06 | 0.000215 | 2.49 |
| brown | GO:0034643 | establishment of mitochondrion localization, microtubule-mediated | 3.36e-06 | 0.000215 | 13.1 |
| brown | GO:0005768 | endosome | 1.39e-05 | 0.00022200000000000003 | 2.02 |
| brown | GO:0051046 | regulation of secretion | 3.54e-06 | 0.00022400000000000002 | 2.03 |
| brown | GO:0046939 | nucleotide phosphorylation | 3.72e-06 | 0.000234 | 6.24 |
| brown | GO:0098797 | plasma membrane protein complex | 1.53e-05 | 0.000241 | 2.21 |
| brown | GO:0043065 | positive regulation of apoptotic process | 4.09e-06 | 0.000256 | 2.22 |
| brown | GO:0055065 | metal ion homeostasis | 4.11e-06 | 0.00025699999999999996 | 2.25 |
| brown | GO:0006119 | oxidative phosphorylation | 4.22e-06 | 0.000262 | 17.47 |
| brown | GO:0007612 | learning | 4.24e-06 | 0.000262 | 3.67 |
| brown | GO:0007611 | learning or memory | 4.22e-06 | 0.000263 | 2.94 |
| brown | GO:0051899 | membrane depolarization | 4.28e-06 | 0.000263 | 6.99 |
| brown | GO:0022836 | gated channel activity | 4.73e-06 | 0.000269 | 2.77 |
| brown | GO:0048878 | chemical homeostasis | 4.41e-06 | 0.00027 | 1.91 |
| brown | GO:0019894 | kinesin binding | 5.19e-06 | 0.000292 | 6.84 |
| brown | GO:0031345 | negative regulation of cell projection organization | 4.79e-06 | 0.000292 | 3.34 |
| brown | GO:0048268 | clathrin coat assembly | 5.07e-06 | 0.000308 | 12.33 |
| brown | GO:0007269 | neurotransmitter secretion | 5.11e-06 | 0.000309 | 7.99 |
| brown | GO:0043068 | positive regulation of programmed cell death | 5.15e-06 | 0.00031 | 2.2 |
| brown | GO:0072525 | pyridine-containing compound biosynthetic process | 5.18e-06 | 0.00031099999999999997 | 6.02 |
| brown | GO:0007005 | mitochondrion organization | 5.26e-06 | 0.000315 | 2.99 |
| brown | GO:0005871 | kinesin complex | 2.0399999999999998e-05 | 0.000321 | 5.82 |
| brown | GO:2001023 | regulation of response to drug | 5.87e-06 | 0.00035 | 4.22 |
| brown | GO:0050807 | regulation of synapse organization | 6.04e-06 | 0.00035800000000000003 | 3.42 |
| brown | GO:0043232 | intracellular non-membrane-bounded organelle | 2.35e-05 | 0.000362 | 1.44 |
| brown | GO:0032984 | protein-containing complex disassembly | 6.16e-06 | 0.000364 | 4.5 |
| brown | GO:0043228 | non-membrane-bounded organelle | 2.35e-05 | 0.000365 | 1.44 |
| brown | GO:0046031 | ADP metabolic process | 6.26e-06 | 0.000368 | 6.69 |
| brown | GO:0005525 | GTP binding | 6.67e-06 | 0.00037 | 2.66 |
| brown | GO:0060627 | regulation of vesicle-mediated transport | 6.44e-06 | 0.00037799999999999997 | 2.41 |
| brown | GO:0098930 | axonal transport | 7.51e-06 | 0.00043799999999999997 | 6.55 |
| brown | GO:0032550 | purine ribonucleoside binding | 8.2e-06 | 0.00045 | 2.63 |
| brown | GO:0010720 | positive regulation of cell development | 7.79e-06 | 0.000454 | 2.16 |
| brown | GO:0032409 | regulation of transporter activity | 7.9e-06 | 0.000458 | 3.01 |
| brown | GO:0019674 | NAD metabolic process | 7.96e-06 | 0.000458 | 7.55 |
| brown | GO:0016079 | synaptic vesicle exocytosis | 7.96e-06 | 0.00045999999999999996 | 7.55 |
| brown | GO:0042592 | homeostatic process | 8.59e-06 | 0.000492 | 1.72 |
| brown | GO:0043195 | terminal bouton | 3.37e-05 | 0.000516 | 4.47 |
| brown | GO:0090148 | membrane fission | 9.52e-06 | 0.000544 | 23.29 |
| brown | GO:0001883 | purine nucleoside binding | 1.01e-05 | 0.000545 | 2.6 |
| brown | GO:0021549 | cerebellum development | 9.84e-06 | 0.000558 | 7.36 |
| brown | GO:1901998 | toxin transport | 9.84e-06 | 0.00056 | 7.36 |
| brown | GO:0008553 | proton-exporting ATPase activity, phosphorylative mechanism | 1.06e-05 | 0.00056 | 11.03 |
| brown | GO:0032549 | ribonucleoside binding | 1.06e-05 | 0.000566 | 2.59 |
| brown | GO:0035383 | thioester metabolic process | 1.0199999999999999e-05 | 0.0005740000000000001 | 5.06 |
| brown | GO:0006637 | acyl-CoA metabolic process | 1.0199999999999999e-05 | 0.000576 | 5.06 |
| brown | GO:0005975 | carbohydrate metabolic process | 1.04e-05 | 0.000582 | 2.54 |
| brown | GO:0051654 | establishment of mitochondrion localization | 1.06e-05 | 0.000589 | 11.03 |
| brown | GO:0006084 | acetyl-CoA metabolic process | 1.07e-05 | 0.00059 | 8.73 |
| brown | GO:0072583 | clathrin-dependent endocytosis | 1.06e-05 | 0.000591 | 11.03 |
| brown | GO:0034703 | cation channel complex | 3.9e-05 | 0.000592 | 2.98 |
| brown | GO:0051129 | negative regulation of cellular component organization | 1.0800000000000002e-05 | 0.000597 | 2.06 |
| brown | GO:0010033 | response to organic substance | 1.09e-05 | 0.0006 | 1.59 |
| brown | GO:0060548 | negative regulation of cell death | 1.13e-05 | 0.000618 | 1.84 |
| brown | GO:0044283 | small molecule biosynthetic process | 1.1400000000000001e-05 | 0.000621 | 2.34 |
| brown | GO:0051239 | regulation of multicellular organismal process | 1.15e-05 | 0.000625 | 1.44 |
| brown | GO:0042734 | presynaptic membrane | 4.2e-05 | 0.0006320000000000001 | 4.79 |
| brown | GO:0030132 | clathrin coat of coated pit | 4.24e-05 | 0.000634 | 17.47 |
| brown | GO:0016471 | vacuolar proton-transporting V-type ATPase complex | 4.47e-05 | 0.000663 | 11.65 |
| brown | GO:0005200 | structural constituent of cytoskeleton | 1.29e-05 | 0.000674 | 5.46 |
| brown | GO:0051954 | positive regulation of amine transport | 1.26e-05 | 0.000684 | 6.17 |
| brown | GO:0071786 | endoplasmic reticulum tubular network organization | 1.27e-05 | 0.000684 | 14.56 |
| brown | GO:0022898 | regulation of transmembrane transporter activity | 1.4300000000000002e-05 | 0.000771 | 2.99 |
| brown | GO:0009135 | purine nucleoside diphosphate metabolic process | 1.49e-05 | 0.000793 | 6.05 |
| brown | GO:0009179 | purine ribonucleoside diphosphate metabolic process | 1.49e-05 | 0.000796 | 6.05 |
| brown | GO:0019359 | nicotinamide nucleotide biosynthetic process | 1.49e-05 | 0.000798 | 6.05 |
| brown | GO:0055074 | calcium ion homeostasis | 1.59e-05 | 0.0008449999999999999 | 2.38 |
| brown | GO:0010942 | positive regulation of cell death | 1.6e-05 | 0.0008449999999999999 | 2.07 |
| brown | GO:0001882 | nucleoside binding | 1.73e-05 | 0.000893 | 2.52 |
| brown | GO:0050769 | positive regulation of neurogenesis | 1.7100000000000002e-05 | 0.0009029999999999999 | 2.2 |
| brown | GO:0019001 | guanyl nucleotide binding | 1.8100000000000003e-05 | 0.000915 | 2.51 |
| brown | GO:0017156 | calcium ion regulated exocytosis | 1.75e-05 | 0.000918 | 5.93 |
| brown | GO:0000226 | microtubule cytoskeleton organization | 1.76e-05 | 0.000924 | 2.57 |
| brown | GO:0032561 | guanyl ribonucleotide binding | 1.8100000000000003e-05 | 0.000926 | 2.51 |
| brown | GO:0047496 | vesicle transport along microtubule | 1.7800000000000002e-05 | 0.0009310000000000001 | 6.82 |
| brown | GO:0010243 | response to organonitrogen compound | 1.84e-05 | 0.000959 | 2.11 |
| brown | GO:0016052 | carbohydrate catabolic process | 1.9e-05 | 0.000985 | 4.74 |
| brown | GO:0044070 | regulation of anion transport | 1.93e-05 | 0.0009960000000000001 | 4.06 |
| yellow | GO:0044391 | ribosomal subunit | 2.3399999999999997e-76 | 4.4399999999999996e-73 | 77.5 |

| | | | | | |
|---|---|---|---|---|---|
| yellow | GO:0044445 | cytosolic part | 4.2599999999999999e-73 | 4.04e-70 | 65.83 |
| yellow | GO:0003735 | structural constituent of ribosome | 1.1699999999999998e-63 | 5.28e-60 | 88.25 |
| yellow | GO:0005840 | ribosome | 6.33e-56 | 4.01e-53 | 70.14 |
| yellow | GO:1990904 | ribonucleoprotein complex | 1.65e-52 | 7.84e-50 | 20.49 |
| yellow | GO:0022625 | cytosolic large ribosomal subunit | 7.67e-43 | 2.91e-40 | 109.85 |
| yellow | GO:0022627 | cytosolic small ribosomal subunit | 1.56e-42 | 4.9299999999999995e-40 | 152.24 |
| yellow | GO:0005198 | structural molecule activity | 2.51e-43 | 5.65e-40 | 24.97 |
| yellow | GO:0015935 | small ribosomal subunit | 2.74e-38 | 7.429999999999999e-36 | 102.71 |
| yellow | GO:0015934 | large ribosomal subunit | 3.85e-37 | 9.14e-35 | 65.75 |
| yellow | GO:0006412 | translation | 1.9900000000000002e-36 | 3.04e-32 | 40.29 |
| yellow | GO:0043043 | peptide biosynthetic process | 1.09e-35 | 8.27e-32 | 37.83 |
| yellow | GO:0043604 | amide biosynthetic process | 1.32e-32 | 6.709999999999999e-29 | 29.0 |
| yellow | GO:0006518 | peptide metabolic process | 5.08e-32 | 1.9399999999999998e-28 | 27.56 |
| yellow | GO:0043603 | cellular amide metabolic process | 3.6099999999999994e-27 | 1.1000000000000001e-23 | 17.98 |
| yellow | GO:0002181 | cytoplasmic translation | 6.46e-27 | 1.6399999999999998e-23 | 126.3 |
| yellow | GO:0005622 | intracellular | 1.02e-24 | 2.15e-22 | 14.44 |
| yellow | GO:1901566 | organonitrogen compound biosynthetic process | 1.0199999999999999e-23 | 2.23e-20 | 13.19 |
| yellow | GO:0032991 | protein-containing complex | 4.69e-22 | 8.91e-20 | 3.58 |
| yellow | GO:0043232 | intracellular non-membrane-bounded organelle | 2.71e-18 | 4.29e-16 | 4.91 |
| yellow | GO:0043228 | non-membrane-bounded organelle | 2.71e-18 | 4.68e-16 | 4.91 |
| yellow | GO:0042788 | polysomal ribosome | 1.5e-17 | 2.2e-15 | 125.99 |
| yellow | GO:0044446 | intracellular organelle part | 4.25e-17 | 5.77e-15 | 2.75 |
| yellow | GO:0044444 | cytoplasmic part | 1.32e-16 | 1.67e-14 | 2.45 |
| yellow | GO:0044422 | organelle part | 2.1800000000000002e-16 | 2.58e-14 | 2.65 |
| yellow | GO:0019843 | rRNA binding | 1.1199999999999999e-15 | 1.68e-12 | 57.99 |
| yellow | GO:0042274 | ribosomal small subunit biogenesis | 3.99e-15 | 7.61e-12 | 177.61 |
| yellow | GO:0006364 | rRNA processing | 7.41e-15 | 1.26e-11 | 28.66 |
| yellow | GO:0016072 | rRNA metabolic process | 3.43e-14 | 4.75e-11 | 25.24 |
| yellow | GO:0034645 | cellular macromolecule biosynthetic process | 3.39e-14 | 5.17e-11 | 4.85 |
| yellow | GO:0022618 | ribonucleoprotein complex assembly | 5.26e-14 | 6.679999999999999e-11 | 24.36 |
| yellow | GO:0009059 | macromolecule biosynthetic process | 5.97e-14 | 6.999999999999999e-11 | 4.74 |
| yellow | GO:0071826 | ribonucleoprotein complex subunit organization | 1.1199999999999999e-13 | 1.2200000000000002e-10 | 22.87 |
| yellow | GO:0003723 | RNA binding | 7.13e-13 | 8.01e-10 | 7.8 |
| yellow | GO:0044267 | cellular protein metabolic process | 8.329999999999999e-13 | 8.47e-10 | 4.49 |
| yellow | GO:0000028 | ribosomal small subunit assembly | 1.9900000000000002e-12 | 1.89e-09 | 143.28 |
| yellow | GO:0044271 | cellular nitrogen compound biosynthetic process | 3.7300000000000005e-12 | 3.3399999999999996e-09 | 4.22 |
| yellow | GO:0022613 | ribonucleoprotein complex biogenesis | 5.09e-12 | 4.31e-09 | 33.83 |
| yellow | GO:0034470 | ncRNA processing | 8.4e-12 | 6.74e-09 | 15.87 |
| yellow | GO:0044085 | cellular component biogenesis | 1.33e-11 | 1.01e-08 | 30.45 |
| yellow | GO:0044249 | cellular biosynthetic process | 6.85e-11 | 4.9699999999999995e-08 | 3.57 |
| yellow | GO:1901576 | organic substance biosynthetic process | 1.42e-10 | 9.86e-08 | 3.46 |
| yellow | GO:0034660 | ncRNA metabolic process | 1.8499999999999998e-10 | 1.23e-07 | 12.15 |
| yellow | GO:0009058 | biosynthetic process | 2.1499999999999998e-10 | 1.37e-07 | 3.4 |
| yellow | GO:0034641 | cellular nitrogen compound metabolic process | 3.71e-10 | 2.26e-07 | 3.07 |
| yellow | GO:0019538 | protein metabolic process | 6.31e-10 | 3.7e-07 | 3.39 |
| yellow | GO:0044424 | intracellular part | 3.69e-08 | 4.12e-06 | 1.53 |
| yellow | GO:0034622 | cellular protein-containing complex assembly | 6.02e-08 | 3.4e-05 | 7.25 |
| yellow | GO:0006396 | RNA processing | 6.43e-08 | 3.5e-05 | 7.21 |
| yellow | GO:1901564 | organonitrogen compound metabolic process | 1.1800000000000001e-07 | 6.21e-05 | 2.68 |
| yellow | GO:0044260 | cellular macromolecule metabolic process | 1.99e-07 | 0.000101 | 2.45 |
| yellow | GO:0043170 | macromolecule metabolic process | 2.3100000000000002e-07 | 0.00011399999999999999 | 2.23 |
| yellow | GO:0070180 | large ribosomal subunit rRNA binding | 4.89e-07 | 0.00044 | 173.98 |
| yellow | GO:0030490 | maturation of SSU-rRNA | 9.759999999999998e-07 | 0.000465 | 52.38 |
| yellow | GO:0000027 | ribosomal large subunit assembly | 1.61e-06 | 0.000745 | 46.4 |
| yellow | GO:0043229 | intracellular organelle | 7.22e-06 | 0.0007610000000000001 | 1.59 |
| yellow | GO:0022626 | cytosolic ribosome | 7.699999999999999e-06 | 0.000769 | 76.12 |
| blue | GO:0044424 | intracellular part | 6.529999999999998e-127 | 1.2399999999999998e-123 | 1.32 |
| blue | GO:0043226 | organelle | 5.699999999999999e-102 | 5.409999999999999e-99 | 1.36 |
| blue | GO:0043229 | intracellular organelle | 1.0799999999999998e-101 | 6.849999999999998e-99 | 1.37 |
| blue | GO:0044464 | cell part | 8.099999999999997e-91 | 3.8399999999999997e-88 | 1.21 |
| blue | GO:0043231 | intracellular membrane-bounded organelle | 6.979999999999998e-88 | 2.649999999999999e-85 | 1.41 |
| blue | GO:0043227 | membrane-bounded organelle | 4.809999999999985e-84 | 1.5199999999999996e-81 | 1.37 |
| blue | GO:0044446 | intracellular organelle part | 1.0499999999999997e-69 | 2.8599999999999997e-67 | 1.47 |
| blue | GO:0044422 | organelle part | 3.4299999999999996e-69 | 8.129999999999998e-67 | 1.45 |
| blue | GO:0044237 | cellular metabolic process | 1.5999999999999998e-67 | 2.4399999999999993e-63 | 1.45 |
| blue | GO:0008152 | metabolic process | 3.0499999999999995e-60 | 2.3199999999999997e-56 | 1.38 |
| blue | GO:0044444 | cytoplasmic part | 1.9099999999999998e-58 | 4.0199999999999994e-56 | 1.37 |
| blue | GO:0071704 | organic substance metabolic process | 7.440000000000001e-56 | 3.779999999999999e-52 | 1.38 |
| blue | GO:0044238 | primary metabolic process | 1.73e-53 | 6.59e-50 | 1.39 |
| blue | GO:0006807 | nitrogen compound metabolic process | 2.4299999999999996e-49 | 7.41e-46 | 1.4 |
| blue | GO:0005634 | nucleus | 2.41e-41 | 4.57e-39 | 1.41 |
| blue | GO:0034641 | cellular nitrogen compound metabolic process | 2.1199999999999998e-42 | 5.3899999999999994e-39 | 1.52 |
| blue | GO:0044260 | cellular macromolecule metabolic process | 1.71e-40 | 3.72e-37 | 1.45 |
| blue | GO:0043170 | macromolecule metabolic process | 6.96e-40 | 1.33e-36 | 1.39 |
| blue | GO:0044428 | nuclear part | 3.1e-38 | 5.3499999999999996e-36 | 1.52 |
| blue | GO:0005488 | binding | 6.81e-38 | 3.0599999999999997e-34 | 1.19 |
| blue | GO:0032991 | protein-containing complex | 7.42e-36 | 1.1699999999999999e-33 | 1.4 |
| blue | GO:0009058 | biosynthetic process | 1.32e-33 | 2.23e-30 | 1.52 |
| blue | GO:0046483 | heterocycle metabolic process | 4.0299999999999995e-33 | 6.15e-30 | 1.49 |
| blue | GO:1901576 | organic substance biosynthetic process | 5.16e-33 | 7.16e-30 | 1.52 |
| blue | GO:0006139 | nucleobase-containing compound metabolic process | 9.579999999999999e-33 | 1.2200000000000001e-29 | 1.5 |
| blue | GO:0006725 | cellular aromatic compound metabolic process | 7.21e-32 | 8.459999999999999e-29 | 1.47 |
| blue | GO:1901360 | organic cyclic compound metabolic process | 8.22e-32 | 8.95e-29 | 1.46 |
| blue | GO:0044249 | cellular biosynthetic process | 1.03e-31 | 1.05e-28 | 1.52 |
| blue | GO:1901363 | heterocyclic compound binding | 6.1e-31 | 1.37e-27 | 1.39 |
| blue | GO:0005829 | cytosol | 1.0800000000000002e-29 | 1.58e-27 | 1.53 |
| blue | GO:0097159 | organic cyclic compound binding | 1.24e-30 | 1.8600000000000004e-27 | 1.38 |
| blue | GO:0009987 | cellular process | 3.6e-30 | 3.43e-27 | 1.15 |
| blue | GO:0019222 | regulation of metabolic process | 3.09e-29 | 2.7699999999999996e-26 | 1.34 |
| blue | GO:0090304 | nucleic acid metabolic process | 7.76e-29 | 6.230000000000001e-26 | 1.51 |
| blue | GO:0016070 | RNA metabolic process | 7.480000000000001e-29 | 6.329999999999999e-26 | 1.56 |
| blue | GO:0010468 | regulation of gene expression | 4.13e-27 | 3.15e-24 | 1.43 |
| blue | GO:0044271 | cellular nitrogen compound biosynthetic process | 6.31e-27 | 4.37e-24 | 1.53 |
| blue | GO:0060255 | regulation of macromolecule metabolic process | 6.25e-27 | 4.5400000000000004e-24 | 1.34 |
| blue | GO:0009059 | macromolecule biosynthetic process | 9.88e-27 | 6.5499999999999995e-24 | 1.56 |
| blue | GO:0005515 | protein binding | 6.23e-27 | 7.01e-24 | 1.23 |
| blue | GO:0003676 | nucleic acid binding | 9.62e-27 | 8.660000000000001e-24 | 1.49 |
| blue | GO:0031323 | regulation of cellular metabolic process | 3.0900000000000004e-26 | 1.96e-23 | 1.33 |
| blue | GO:0071840 | cellular component organization or biogenesis | 5.05e-26 | 3.08e-23 | 1.37 |
| blue | GO:0034645 | cellular macromolecule biosynthetic process | 9.18e-26 | 5.38e-23 | 1.55 |
| blue | GO:0080090 | regulation of primary metabolic process | 1.84e-25 | 1.04e-22 | 1.33 |
| blue | GO:0005737 | cytoplasm | 1.65e-24 | 2.2299999999999996e-22 | 1.28 |
| blue | GO:0051171 | regulation of nitrogen compound metabolic process | 4.66e-24 | 2.54e-21 | 1.33 |
| blue | GO:0016043 | cellular component organization | 1.44e-23 | 7.58e-21 | 1.35 |

| | | | | | |
|---|---|---|---|---|---|
| blue | GO:2000112 | regulation of cellular macromolecule biosynthetic process | 1.7699999999999999e-22 | 9.01e-20 | 1.42 |
| blue | GO:0009889 | regulation of biosynthetic process | 3.21e-22 | 1.58e-19 | 1.39 |
| blue | GO:1901564 | organonitrogen compound metabolic process | 4.34e-22 | 2.0699999999999999e-19 | 1.36 |
| blue | GO:0043232 | intracellular non-membrane-bounded organelle | 3.15e-21 | 3.7299999999999996e-19 | 1.45 |
| blue | GO:0043228 | non-membrane-bounded organelle | 3.15e-21 | 3.98e-19 | 1.45 |
| blue | GO:0010556 | regulation of macromolecule biosynthetic process | 1.06e-21 | 4.9e-19 | 1.4 |
| blue | GO:0031326 | regulation of cellular biosynthetic process | 1.3699999999999998e-21 | 6.16e-19 | 1.39 |
| blue | GO:0018130 | heterocycle biosynthetic process | 5.660000000000001e-21 | 2.47e-18 | 1.5 |
| blue | GO:0044267 | cellular protein metabolic process | 9.19e-21 | 3.889999999999999e-18 | 1.48 |
| blue | GO:1901362 | organic cyclic compound biosynthetic process | 3.4e-20 | 1.4000000000000003e-17 | 1.48 |
| blue | GO:0034654 | nucleobase-containing compound biosynthetic process | 5.83e-20 | 2.34e-17 | 1.5 |
| blue | GO:0005654 | nucleoplasm | 2.2600000000000004e-19 | 2.52e-17 | 1.53 |
| blue | GO:0019438 | aromatic compound biosynthetic process | 7.89e-20 | 3.09e-17 | 1.49 |
| blue | GO:0019219 | regulation of nucleobase-containing compound metabolic process | 2.3600000000000003e-19 | 8.99e-17 | 1.38 |
| blue | GO:0003824 | catalytic activity | 1.35e-19 | 1.01e-16 | 1.27 |
| blue | GO:0051252 | regulation of RNA metabolic process | 4.66e-19 | 1.73e-16 | 1.39 |
| blue | GO:0032774 | RNA biosynthetic process | 7.740000000000001e-19 | 2.81e-16 | 1.53 |
| blue | GO:0006351 | transcription, DNA-templated | 1.79e-18 | 6.360000000000001e-16 | 1.53 |
| blue | GO:0097659 | nucleic acid-templated transcription | 1.96e-18 | 6.809999999999999e-16 | 1.52 |
| blue | GO:0043167 | ion binding | 1.22e-18 | 7.849999999999999e-16 | 1.28 |
| blue | GO:0048523 | negative regulation of cellular process | 1.4300000000000002e-17 | 4.85e-15 | 1.3 |
| blue | GO:0009892 | negative regulation of metabolic process | 2.37e-17 | 7.840000000000001e-15 | 1.42 |
| blue | GO:0010629 | negative regulation of gene expression | 3.04e-17 | 9.87e-15 | 1.57 |
| blue | GO:1990904 | ribonucleoprotein complex | 1.1399999999999998e-16 | 1.2e-14 | 1.78 |
| blue | GO:0048519 | negative regulation of biological process | 4.28e-17 | 1.36e-14 | 1.28 |
| blue | GO:0009890 | negative regulation of biosynthetic process | 7.78e-17 | 2.42e-14 | 1.59 |
| blue | GO:0003723 | RNA binding | 8.52e-17 | 4.79e-14 | 1.72 |
| blue | GO:0010605 | negative regulation of macromolecule metabolic process | 1.7e-16 | 5.18e-14 | 1.43 |
| blue | GO:0016740 | transferase activity | 1.38e-16 | 6.89e-14 | 1.45 |
| blue | GO:0010558 | negative regulation of macromolecule biosynthetic process | 2.62e-16 | 7.82e-14 | 1.6 |
| blue | GO:1902494 | catalytic complex | 8.710000000000001e-16 | 8.7e-14 | 1.6 |
| blue | GO:0048522 | positive regulation of cellular process | 3.78e-16 | 1.1100000000000001e-13 | 1.27 |
| blue | GO:2000113 | negative regulation of cellular macromolecule biosynthetic process | 5.14e-16 | 1.4800000000000003e-13 | 1.6 |
| blue | GO:0031327 | negative regulation of cellular biosynthetic process | 7.059999999999999e-16 | 1.9899999999999999e-13 | 1.58 |
| blue | GO:0043412 | macromolecule modification | 1.24e-15 | 3.43e-13 | 1.42 |
| blue | GO:0031090 | organelle membrane | 4.020000000000001e-15 | 3.81e-13 | 1.56 |
| blue | GO:0044451 | nucleoplasm part | 4.5799999999999995e-15 | 4.1399999999999995e-13 | 1.68 |
| blue | GO:0006355 | regulation of transcription, DNA-templated | 1.6899999999999999e-15 | 4.59e-13 | 1.36 |
| blue | GO:1903506 | regulation of nucleic acid-templated transcription | 1.92e-15 | 5.14e-13 | 1.36 |
| blue | GO:2001141 | regulation of RNA biosynthetic process | 2.85e-15 | 7.48e-13 | 1.36 |
| blue | GO:0031324 | negative regulation of cellular metabolic process | 5.82e-15 | 1.4999999999999999e-12 | 1.41 |
| blue | GO:0008104 | protein localization | 6.1e-15 | 1.55e-12 | 1.5 |
| blue | GO:0033036 | macromolecule localization | 7.2e-15 | 1.8000000000000002e-12 | 1.49 |
| blue | GO:0048518 | positive regulation of biological process | 1.05e-14 | 2.59e-12 | 1.23 |
| blue | GO:0036211 | protein modification process | 1.43e-14 | 3.3999999999999996e-12 | 1.42 |
| blue | GO:0006464 | cellular protein modification process | 1.43e-14 | 3.4500000000000004e-12 | 1.42 |
| blue | GO:0019538 | protein metabolic process | 1.53e-14 | 3.59e-12 | 1.33 |
| blue | GO:0009893 | positive regulation of metabolic process | 3.99e-14 | 9.21e-12 | 1.32 |
| blue | GO:0051172 | negative regulation of nitrogen compound metabolic process | 6.42e-14 | 1.46e-11 | 1.41 |
| blue | GO:0031325 | positive regulation of cellular metabolic process | 7.400000000000001e-14 | 1.6600000000000003e-11 | 1.33 |
| blue | GO:0051641 | cellular localization | 1.3699999999999997e-13 | 3.03e-11 | 1.46 |
| blue | GO:0005739 | mitochondrion | 4.1200000000000006e-13 | 3.5600000000000005e-11 | 1.47 |
| blue | GO:0051253 | negative regulation of RNA metabolic process | 2.61e-13 | 5.6e-11 | 1.56 |
| blue | GO:0010604 | positive regulation of macromolecule metabolic process | 2.59e-13 | 5.6299999999999996e-11 | 1.33 |
| blue | GO:0022625 | cytosolic large ribosomal subunit | 7.009999999999999e-13 | 5.79e-11 | 3.43 |
| blue | GO:0045934 | negative regulation of nucleobase-containing compound metabolic pro | 3.9399999999999994e-13 | 8.35e-11 | 1.52 |
| blue | GO:0009894 | regulation of catabolic process | 5.770000000000001e-13 | 1.2e-10 | 1.72 |
| blue | GO:0051173 | positive regulation of nitrogen compound metabolic process | 7.979999999999998e-13 | 1.64e-10 | 1.33 |
| blue | GO:0044391 | ribosomal subunit | 2.8700000000000005e-12 | 2.2700000000000002e-10 | 2.37 |
| blue | GO:1903507 | negative regulation of nucleic acid-templated transcription | 4.83e-12 | 9.83e-10 | 1.55 |
| blue | GO:0019899 | enzyme binding | 2.32e-12 | 1.0400000000000001e-09 | 1.38 |
| blue | GO:1902679 | negative regulation of RNA biosynthetic process | 5.3e-12 | 1.06e-09 | 1.55 |
| blue | GO:0009056 | catabolic process | 5.419999999999999e-12 | 1.07e-09 | 1.46 |
| blue | GO:0045892 | negative regulation of transcription, DNA-templated | 6.480000000000001e-12 | 1.27e-09 | 1.55 |
| blue | GO:1990234 | transferase complex | 1.68e-11 | 1.2800000000000001e-09 | 1.68 |
| blue | GO:0044248 | cellular catabolic process | 6.97e-12 | 1.3400000000000001e-09 | 1.49 |
| blue | GO:0010608 | posttranscriptional regulation of gene expression | 1.7e-11 | 3.25e-09 | 1.98 |
| blue | GO:0006357 | regulation of transcription by RNA polymerase II | 1.82e-11 | 3.42e-09 | 1.4 |
| blue | GO:0003677 | DNA binding | 9.92e-12 | 4.05e-09 | 1.39 |
| blue | GO:0051246 | regulation of protein metabolic process | 2.5999999999999997e-11 | 4.84e-09 | 1.33 |
| blue | GO:0051179 | localization | 3.7299999999999997e-11 | 6.859999999999999e-09 | 1.26 |
| blue | GO:0010628 | positive regulation of gene expression | 4.49e-11 | 8.15e-09 | 1.39 |
| blue | GO:0031329 | regulation of cellular catabolic process | 5.73e-11 | 1.03e-08 | 1.71 |
| blue | GO:0006996 | organelle organization | 6.61e-11 | 1.1699999999999999e-08 | 1.38 |
| blue | GO:0045184 | establishment of protein localization | 1.17e-10 | 2.06e-08 | 1.5 |
| blue | GO:0042886 | amide transport | 1.49e-10 | 2.58e-08 | 1.5 |
| blue | GO:0044445 | cytosolic part | 4.34e-10 | 3.17e-08 | 2.11 |
| blue | GO:0005794 | Golgi apparatus | 5.28e-10 | 3.46e-08 | 1.47 |
| blue | GO:0015934 | large ribosomal subunit | 5.14e-10 | 3.48e-08 | 2.54 |
| blue | GO:0006325 | chromatin organization | 2.1000000000000002e-10 | 3.59e-08 | 1.7 |
| blue | GO:0005815 | microtubule organizing center | 5.11e-10 | 3.59e-08 | 1.69 |
| blue | GO:0022607 | cellular component assembly | 2.41e-10 | 4.04e-08 | 1.36 |
| blue | GO:0006396 | RNA processing | 2.41e-10 | 4.08e-08 | 1.66 |
| blue | GO:1901575 | organic substance catabolic process | 2.4899999999999997e-10 | 4.13e-08 | 1.46 |
| blue | GO:0051716 | cellular response to stimulus | 3.11e-10 | 5.11e-08 | 1.32 |
| blue | GO:0006417 | regulation of translation | 3.46e-10 | 5.62e-08 | 2.05 |
| blue | GO:0034248 | regulation of cellular amide metabolic process | 3.75e-10 | 6.02e-08 | 1.97 |
| blue | GO:0046907 | intracellular transport | 3.9899999999999997e-10 | 6.34e-08 | 1.5 |
| blue | GO:0015833 | peptide transport | 4.59e-10 | 7.21e-08 | 1.49 |
| blue | GO:0051254 | positive regulation of RNA metabolic process | 5.609999999999999e-10 | 8.730000000000001e-08 | 1.4 |
| blue | GO:0044431 | Golgi apparatus part | 1.88e-09 | 1.1900000000000001e-07 | 1.77 |
| blue | GO:0015031 | protein transport | 8.55e-10 | 1.32e-07 | 1.49 |
| blue | GO:0006793 | phosphorus metabolic process | 1.23e-09 | 1.87e-07 | 1.39 |
| blue | GO:0032268 | regulation of cellular protein metabolic process | 1.52e-09 | 2.2899999999999997e-07 | 1.31 |
| blue | GO:1903311 | regulation of mRNA metabolic process | 1.65e-09 | 2.46e-07 | 2.15 |
| blue | GO:0006796 | phosphate-containing compound metabolic process | 1.75e-09 | 2.56e-07 | 1.39 |
| blue | GO:0044281 | small molecule metabolic process | 1.7300000000000002e-09 | 2.56e-07 | 1.41 |
| blue | GO:0045935 | positive regulation of nucleobase-containing compound metabolic proc | 1.8e-09 | 2.62e-07 | 1.37 |
| blue | GO:0019866 | organelle inner membrane | 4.4700000000000005e-09 | 2.74e-07 | 1.84 |
| blue | GO:0009062 | fatty acid catabolic process | 2.17e-09 | 3.12e-07 | 3.09 |
| blue | GO:0043169 | cation binding | 9.49e-10 | 3.2799999999999997e-07 | 1.25 |
| blue | GO:0016772 | transferase activity, transferring phosphorus-containing groups | 9.300000000000001e-10 | 3.4899999999999996e-07 | 1.54 |
| blue | GO:0003729 | mRNA binding | 1.27e-09 | 4.0799999999999995e-07 | 2.12 |
| blue | GO:0046872 | metal ion binding | 1.54e-09 | 4.6e-07 | 1.25 |

| | | | | | |
|---|---|---|---|---|---|
| blue | GO:0043603 | cellular amide metabolic process | 3.2299999999999994e-09 | 4.61e-07 | 1.66 |
| blue | GO:1901566 | organonitrogen compound biosynthetic process | 3.65e-09 | 5.15e-07 | 1.56 |
| blue | GO:0043168 | anion binding | 1.88e-09 | 5.29e-07 | 1.29 |
| blue | GO:0006518 | peptide metabolic process | 5.420000000000001e-09 | 7.58e-07 | 1.82 |
| blue | GO:0005783 | endoplasmic reticulum | 1.58e-08 | 9.359999999999999e-07 | 1.38 |
| blue | GO:0048193 | Golgi vesicle transport | 7.609999999999998e-09 | 1.06e-06 | 2.09 |
| blue | GO:0036094 | small molecule binding | 4.01e-09 | 1.06e-06 | 1.3 |
| blue | GO:0051960 | regulation of nervous system development | 8.86e-09 | 1.22e-06 | 1.48 |
| blue | GO:0005768 | endosome | 2.17e-08 | 1.25e-06 | 1.55 |
| blue | GO:0005912 | adherens junction | 2.23e-08 | 1.25e-06 | 1.96 |
| blue | GO:0051649 | establishment of localization in cell | 1.04e-08 | 1.42e-06 | 1.41 |
| blue | GO:0010557 | positive regulation of macromolecule biosynthetic process | 1.16e-08 | 1.57e-06 | 1.35 |
| blue | GO:0006635 | fatty acid beta-oxidation | 1.23e-08 | 1.64e-06 | 3.53 |
| blue | GO:0071705 | nitrogen compound transport | 1.27e-08 | 1.68e-06 | 1.39 |
| blue | GO:0000166 | nucleotide binding | 7.85e-09 | 1.86e-06 | 1.33 |
| blue | GO:1901265 | nucleoside phosphate binding | 7.85e-09 | 1.96e-06 | 1.33 |
| blue | GO:0070161 | anchoring junction | 3.68e-08 | 1.99e-06 | 1.92 |
| blue | GO:0005730 | nucleolus | 3.93e-08 | 2.07e-06 | 1.57 |
| blue | GO:0044429 | mitochondrial part | 4.3e-08 | 2.21e-06 | 1.53 |
| blue | GO:0033554 | cellular response to stress | 1.82e-08 | 2.39e-06 | 1.41 |
| blue | GO:0000122 | negative regulation of transcription by RNA polymerase II | 1.94e-08 | 2.53e-06 | 1.52 |
| blue | GO:0034613 | cellular protein localization | 2.12e-08 | 2.73e-06 | 1.52 |
| blue | GO:0046395 | carboxylic acid catabolic process | 2.41e-08 | 3.07e-06 | 2.15 |
| blue | GO:0016054 | organic acid catabolic process | 2.41e-08 | 3.09e-06 | 2.15 |
| blue | GO:0050767 | regulation of neurogenesis | 2.5499999999999997e-08 | 3.22e-06 | 1.49 |
| blue | GO:0016192 | vesicle-mediated transport | 2.71e-08 | 3.34e-06 | 1.48 |
| blue | GO:0070727 | cellular macromolecule localization | 2.67e-08 | 3.34e-06 | 1.51 |
| blue | GO:1903508 | positive regulation of nucleic acid-templated transcription | 2.7e-08 | 3.34e-06 | 1.37 |
| blue | GO:1902680 | positive regulation of RNA biosynthetic process | 2.8700000000000002e-08 | 3.5000000000000004e-06 | 1.37 |
| blue | GO:0045893 | positive regulation of transcription, DNA-templated | 3.6e-08 | 4.36e-06 | 1.37 |
| blue | GO:0003682 | chromatin binding | 1.94e-08 | 4.36e-06 | 1.64 |
| blue | GO:0031328 | positive regulation of cellular biosynthetic process | 3.91e-08 | 4.66e-06 | 1.33 |
| blue | GO:0050793 | regulation of developmental process | 3.8900000000000004e-08 | 4.67e-06 | 1.27 |
| blue | GO:0005840 | ribosome | 1.15e-07 | 5.73e-06 | 2.09 |
| blue | GO:0016071 | mRNA metabolic process | 5.0999999999999993e-08 | 6.03e-06 | 1.68 |
| blue | GO:0003712 | transcription coregulator activity | 3.32e-08 | 7.11e-06 | 1.72 |
| blue | GO:0005813 | centrosome | 1.5e-07 | 7.32e-06 | 1.65 |
| blue | GO:0006412 | translation | 6.55e-08 | 7.68e-06 | 1.93 |
| blue | GO:0140096 | catalytic activity, acting on a protein | 3.92e-08 | 8.02e-06 | 1.3 |
| blue | GO:0120035 | regulation of plasma membrane bounded cell projection organization | 7.31e-08 | 8.51e-06 | 1.53 |
| blue | GO:0016569 | covalent chromatin modification | 7.46e-08 | 8.62e-06 | 1.84 |
| blue | GO:0031965 | nuclear membrane | 1.8399999999999998e-07 | 8.71e-06 | 1.99 |
| blue | GO:0030030 | cell projection organization | 9.13e-08 | 1.05e-05 | 1.49 |
| blue | GO:0051128 | regulation of cellular component organization | 9.67e-08 | 1.1e-05 | 1.27 |
| blue | GO:0051234 | establishment of localization | 1.04e-07 | 1.18e-05 | 1.23 |
| blue | GO:0009057 | macromolecule catabolic process | 1.06e-07 | 1.18e-05 | 1.53 |
| blue | GO:0009891 | positive regulation of biosynthetic process | 1.1e-07 | 1.22e-05 | 1.31 |
| blue | GO:0042802 | identical protein binding | 6.929999999999999e-08 | 1.36e-05 | 1.32 |
| blue | GO:0044265 | cellular macromolecule catabolic process | 1.28e-07 | 1.4099999999999999e-05 | 1.57 |
| blue | GO:0032502 | developmental process | 1.32e-07 | 1.4499999999999998e-05 | 1.18 |
| blue | GO:0019395 | fatty acid oxidation | 1.4599999999999998e-07 | 1.59e-05 | 2.94 |
| blue | GO:0016604 | nuclear body | 3.44e-07 | 1.59e-05 | 1.54 |
| blue | GO:0071702 | organic substance transport | 1.57e-07 | 1.7e-05 | 1.32 |
| blue | GO:0030258 | lipid modification | 1.6100000000000003e-07 | 1.7199999999999998e-05 | 2.2 |
| blue | GO:0043043 | peptide biosynthetic process | 1.62e-07 | 1.73e-05 | 1.87 |
| blue | GO:0031344 | regulation of cell projection organization | 1.73e-07 | 1.82e-05 | 1.51 |
| blue | GO:0006810 | transport | 1.73e-07 | 1.83e-05 | 1.23 |
| blue | GO:0005856 | cytoskeleton | 4.15e-07 | 1.87e-05 | 1.35 |
| blue | GO:0072329 | monocarboxylic acid catabolic process | 1.8e-07 | 1.88e-05 | 2.59 |
| blue | GO:0042995 | cell projection | 4.37e-07 | 1.93e-05 | 1.28 |
| blue | GO:0034440 | lipid oxidation | 2.05e-07 | 2.13e-05 | 2.89 |
| blue | GO:0016570 | histone modification | 2.1800000000000002e-07 | 2.24e-05 | 1.81 |
| blue | GO:0003735 | structural constituent of ribosome | 1.3e-07 | 2.44e-05 | 2.19 |
| blue | GO:0140110 | transcription regulator activity | 1.41e-07 | 2.54e-05 | 1.37 |
| blue | GO:0044798 | nuclear transcription factor complex | 6.54e-07 | 2.7600000000000003e-05 | 1.99 |
| blue | GO:0005743 | mitochondrial inner membrane | 6.429999999999999e-07 | 2.7699999999999996e-05 | 1.73 |
| blue | GO:0032553 | ribonucleotide binding | 1.7699999999999998e-07 | 3.06e-05 | 1.31 |
| blue | GO:0005622 | intracellular | 7.529999999999999e-07 | 3.11e-05 | 1.48 |
| blue | GO:0071310 | cellular response to organic substance | 3.1e-07 | 3.18e-05 | 1.38 |
| blue | GO:0009896 | positive regulation of catabolic process | 3.36e-07 | 3.4200000000000005e-05 | 1.71 |
| blue | GO:0043604 | amide biosynthetic process | 3.46e-07 | 3.4899999999999995e-05 | 1.73 |
| blue | GO:0048471 | perinuclear region of cytoplasm | 9.22e-07 | 3.73e-05 | 1.53 |
| blue | GO:0044430 | cytoskeletal part | 9.709999999999999e-07 | 3.84e-05 | 1.33 |
| blue | GO:0008134 | transcription factor binding | 3.0399999999999997e-07 | 4.8899999999999996e-05 | 1.52 |
| blue | GO:0031984 | organelle subcompartment | 1.27e-06 | 4.92e-05 | 1.85 |
| blue | GO:0045664 | regulation of neuron differentiation | 4.91e-07 | 4.92e-05 | 1.49 |
| blue | GO:0017076 | purine nucleotide binding | 2.95e-07 | 4.92e-05 | 1.31 |
| blue | GO:0032555 | purine ribonucleotide binding | 3.38e-07 | 5.25e-05 | 1.31 |
| blue | GO:0007163 | establishment or maintenance of cell polarity | 5.429999999999999e-07 | 5.3700000000000004e-05 | 2.16 |
| blue | GO:0070647 | protein modification by small protein conjugation or removal | 5.389999999999999e-07 | 5.3700000000000004e-05 | 1.57 |
| blue | GO:0022618 | ribonucleoprotein complex assembly | 5.54e-07 | 5.45e-05 | 2.0 |
| blue | GO:0098791 | Golgi subcompartment | 1.5e-06 | 5.68e-05 | 1.89 |
| blue | GO:0050794 | regulation of cellular process | 6.690000000000001e-07 | 6.54e-05 | 1.09 |
| blue | GO:0045944 | positive regulation of transcription by RNA polymerase II | 7.2e-07 | 6.95e-05 | 1.37 |
| blue | GO:0006402 | mRNA catabolic process | 7.19e-07 | 6.979999999999999e-05 | 2.41 |
| blue | GO:0006974 | cellular response to DNA damage stimulus | 7.6e-07 | 7.290000000000001e-05 | 1.53 |
| blue | GO:0071826 | ribonucleoprotein complex subunit organization | 8.009999999999999e-07 | 7.63e-05 | 1.95 |
| blue | GO:0032868 | response to insulin | 8.13e-07 | 7.7e-05 | 2.3 |
| blue | GO:0005802 | trans-Golgi network | 2.08e-06 | 7.759999999999999e-05 | 2.01 |
| blue | GO:0031400 | negative regulation of protein modification process | 9.009999999999999e-07 | 8.48e-05 | 1.57 |
| blue | GO:0060284 | regulation of cell development | 9.5e-07 | 8.89e-05 | 1.4 |
| blue | GO:0031966 | mitochondrial membrane | 2.45e-06 | 8.94e-05 | 1.56 |
| blue | GO:0005777 | peroxisome | 2.87e-06 | 0.000103 | 2.17 |
| blue | GO:0044255 | cellular lipid metabolic process | 1.14e-06 | 0.000106 | 1.46 |
| blue | GO:0016020 | membrane | 3.08e-06 | 0.00010800000000000001 | 1.11 |
| blue | GO:0016567 | protein ubiquitination | 1.16e-06 | 0.00010800000000000001 | 1.66 |
| blue | GO:0032446 | protein modification by small protein conjugation | 1.2e-06 | 0.00011 | 1.62 |
| blue | GO:0017053 | transcriptional repressor complex | 3.44e-06 | 0.000117 | 2.5 |
| blue | GO:0042579 | microbody | 3.42e-06 | 0.000118 | 2.13 |
| blue | GO:0031331 | positive regulation of cellular catabolic process | 1.31e-06 | 0.00011999999999999999 | 1.74 |
| blue | GO:0043436 | oxoacid metabolic process | 1.37e-06 | 0.000125 | 1.45 |
| blue | GO:0022613 | ribonucleoprotein complex biogenesis | 1.46e-06 | 0.000132 | 2.35 |
| blue | GO:0031346 | positive regulation of cell projection organization | 1.47e-06 | 0.000132 | 1.62 |
| blue | GO:0034470 | ncRNA processing | 1.54e-06 | 0.00013700000000000002 | 1.75 |

| | | | | | |
|---|---|---|---|---|---|
| blue | GO:0051247 | positive regulation of protein metabolic process | 1.56e-06 | 0.000139 | 1.31 |
| blue | GO:0033043 | regulation of organelle organization | 1.69e-06 | 0.000149 | 1.36 |
| blue | GO:0120036 | plasma membrane bounded cell projection organization | 1.76e-06 | 0.000154 | 1.52 |
| blue | GO:0044282 | small molecule catabolic process | 1.88e-06 | 0.000162 | 1.78 |
| blue | GO:0031399 | regulation of protein modification process | 1.87e-06 | 0.00016299999999999998 | 1.29 |
| blue | GO:0050684 | regulation of mRNA processing | 1.91e-06 | 0.000164 | 2.18 |
| blue | GO:0010975 | regulation of neuron projection development | 1.93e-06 | 0.000165 | 1.53 |
| blue | GO:0019752 | carboxylic acid metabolic process | 1.95e-06 | 0.000166 | 1.45 |
| blue | GO:0097458 | neuron part | 5.25e-06 | 0.000175 | 1.27 |
| blue | GO:0016301 | kinase activity | 1.24e-06 | 0.00018600000000000002 | 1.46 |
| blue | GO:0006401 | RNA catabolic process | 2.35e-06 | 0.000199 | 2.19 |
| blue | GO:0016072 | rRNA metabolic process | 2.43e-06 | 0.000205 | 1.95 |
| blue | GO:0065007 | biological regulation | 2.48e-06 | 0.000207 | 1.08 |
| blue | GO:0070887 | cellular response to chemical stimulus | 2.55e-06 | 0.000212 | 1.31 |
| blue | GO:0042393 | histone binding | 1.48e-06 | 0.000215 | 2.03 |
| blue | GO:0051248 | negative regulation of protein metabolic process | 2.65e-06 | 0.00022 | 1.38 |
| blue | GO:0055114 | oxidation-reduction process | 2.76e-06 | 0.000227 | 1.43 |
| blue | GO:0033539 | fatty acid beta-oxidation using acyl-CoA dehydrogenase | 2.88e-06 | 0.000236 | 6.14 |
| blue | GO:0022604 | regulation of cell morphogenesis | 3.31e-06 | 0.00027 | 1.55 |
| blue | GO:0000956 | nuclear-transcribed mRNA catabolic process | 3.44e-06 | 0.000277 | 2.5 |
| blue | GO:0006629 | lipid metabolic process | 3.42e-06 | 0.000278 | 1.38 |
| blue | GO:0097708 | intracellular vesicle | 8.74e-06 | 0.000286 | 1.28 |
| blue | GO:0090090 | negative regulation of canonical Wnt signaling pathway | 3.59e-06 | 0.000288 | 2.23 |
| blue | GO:0035556 | intracellular signal transduction | 3.67e-06 | 0.000293 | 1.33 |
| blue | GO:0009968 | negative regulation of signal transduction | 3.73e-06 | 0.000297 | 1.35 |
| blue | GO:0031410 | cytoplasmic vesicle | 9.55e-06 | 0.000307 | 1.28 |
| blue | GO:0019637 | organophosphate metabolic process | 4.08e-06 | 0.000323 | 1.46 |
| blue | GO:0030054 | cell junction | 1.04e-05 | 0.00032900000000000003 | 1.35 |
| blue | GO:0006364 | rRNA processing | 4.53e-06 | 0.000354 | 1.99 |
| blue | GO:0006082 | organic acid metabolic process | 4.52e-06 | 0.00035499999999999996 | 1.42 |
| blue | GO:0035770 | ribonucleoprotein granule | 1.18e-05 | 0.000368 | 1.86 |
| blue | GO:0098727 | maintenance of cell number | 4.99e-06 | 0.000388 | 2.1 |
| blue | GO:2000026 | regulation of multicellular organismal development | 5.12e-06 | 0.000396 | 1.25 |
| blue | GO:0030055 | cell-substrate junction | 1.3000000000000001e-05 | 0.000398 | 1.98 |
| blue | GO:1901565 | organonitrogen compound catabolic process | 5.3e-06 | 0.00040599999999999995 | 1.49 |
| blue | GO:0006892 | post-Golgi vesicle-mediated transport | 5.29e-06 | 0.000407 | 2.5 |
| blue | GO:0035639 | purine ribonucleoside triphosphate binding | 2.8999999999999998e-06 | 0.000408 | 1.29 |
| blue | GO:0030029 | actin filament-based process | 5.44e-06 | 0.000414 | 1.66 |
| blue | GO:0042176 | regulation of protein catabolic process | 5.48e-06 | 0.00041600000000000003 | 1.65 |
| blue | GO:0034660 | ncRNA metabolic process | 5.58e-06 | 0.000421 | 1.61 |
| blue | GO:0007010 | cytoskeleton organization | 5.66e-06 | 0.000425 | 1.42 |
| blue | GO:0006886 | intracellular protein transport | 6.26e-06 | 0.000468 | 1.47 |
| blue | GO:0071495 | cellular response to endogenous stimulus | 6.31e-06 | 0.00047000000000000004 | 1.49 |
| blue | GO:0047485 | protein N-terminus binding | 3.55e-06 | 0.000484 | 2.18 |
| blue | GO:0009719 | response to endogenous stimulus | 6.55e-06 | 0.00048499999999999997 | 1.42 |
| blue | GO:0006520 | cellular amino acid metabolic process | 6.69e-06 | 0.000488 | 1.78 |
| blue | GO:0044085 | cellular component biogenesis | 6.62e-06 | 0.000488 | 2.18 |
| blue | GO:0035264 | multicellular organism growth | 6.69e-06 | 0.00049 | 2.23 |
| blue | GO:0051130 | positive regulation of cellular component organization | 6.77e-06 | 0.000492 | 1.33 |
| blue | GO:0090575 | RNA polymerase II transcription factor complex | 1.6899999999999997e-05 | 0.000509 | 1.94 |
| blue | GO:0019827 | stem cell population maintenance | 7.46e-06 | 0.000539 | 2.09 |
| blue | GO:0017016 | Ras GTPase binding | 4.56e-06 | 0.000603 | 1.62 |
| blue | GO:0016831 | carboxy-lyase activity | 5.16e-06 | 0.000611 | 3.49 |
| blue | GO:0097367 | carbohydrate derivative binding | 4.91e-06 | 0.000614 | 1.25 |
| blue | GO:0050662 | coenzyme binding | 4.83e-06 | 0.00062 | 1.75 |
| blue | GO:0008092 | cytoskeletal protein binding | 5.14e-06 | 0.000625 | 1.39 |
| blue | GO:1901605 | alpha-amino acid metabolic process | 8.82e-06 | 0.000634 | 1.91 |
| blue | GO:0032270 | positive regulation of cellular protein metabolic process | 9.37e-06 | 0.000671 | 1.29 |
| blue | GO:0051239 | regulation of multicellular organismal process | 9.42e-06 | 0.000671 | 1.19 |
| blue | GO:0043487 | regulation of RNA stability | 9.67e-06 | 0.000686 | 2.34 |
| blue | GO:0007264 | small GTPase mediated signal transduction | 1.0199999999999999e-05 | 0.000717 | 1.75 |
| blue | GO:0050789 | regulation of biological process | 1.0300000000000001e-05 | 0.000721 | 1.07 |
| blue | GO:0030554 | adenyl nucleotide binding | 6.43e-06 | 0.000742 | 1.3 |
| blue | GO:0005925 | focal adhesion | 2.5300000000000002e-05 | 0.0007509999999999999 | 1.99 |
| blue | GO:0004842 | ubiquitin-protein transferase activity | 6.87e-06 | 0.000772 | 1.62 |
| blue | GO:0005924 | cell-substrate adherens junction | 2.8100000000000002e-05 | 0.0008210000000000001 | 1.96 |
| blue | GO:0043434 | response to peptide hormone | 1.18e-05 | 0.000828 | 1.86 |
| blue | GO:0003713 | transcription coactivator activity | 7.9e-06 | 0.0008460000000000001 | 1.76 |
| blue | GO:0032559 | adenyl ribonucleotide binding | 7.73e-06 | 0.000848 | 1.3 |
| blue | GO:0061013 | regulation of mRNA catabolic process | 1.25e-05 | 0.000865 | 2.24 |
| blue | GO:0016458 | gene silencing | 1.25e-05 | 0.000868 | 2.2 |
| blue | GO:0019787 | ubiquitin-like protein transferase activity | 8.31e-06 | 0.000869 | 1.59 |
| blue | GO:0070013 | intracellular organelle lumen | 3.0299999999999998e-05 | 0.0008699999999999999 | 1.68 |
| blue | GO:0016773 | phosphotransferase activity, alcohol group as acceptor | 8.71e-06 | 0.0008900000000000001 | 1.44 |
| blue | GO:0045595 | regulation of cell differentiation | 1.32e-05 | 0.0009130000000000001 | 1.26 |
| blue | GO:0051093 | negative regulation of developmental process | 1.42e-05 | 0.000978 | 1.35 |
| magenta | GO:0016020 | membrane | 1.15e-09 | 2.19e-06 | 1.79 |
| magenta | GO:0097458 | neuron part | 1.15e-07 | 7.269999999999999e-05 | 2.97 |
| magenta | GO:0005886 | plasma membrane | 7.7e-08 | 7.31e-05 | 2.16 |
| magenta | GO:0120025 | plasma membrane bounded cell projection | 2.9699999999999997e-07 | 0.00014099999999999998 | 2.92 |
| magenta | GO:0043005 | neuron projection | 3.85e-07 | 0.000146 | 3.39 |
| magenta | GO:0030054 | cell junction | 9.740000000000001e-07 | 0.000308 | 3.48 |
| magenta | GO:0044459 | plasma membrane part | 1.2e-06 | 0.000324 | 2.37 |
| magenta | GO:0042995 | cell projection | 1.44e-06 | 0.000342 | 2.62 |
| magenta | GO:0030425 | dendrite | 4.1e-06 | 0.0008640000000000001 | 4.69 |
| red | GO:0007272 | ensheathment of neurons | 6.509999999999999e-27 | 4.96e-23 | 34.16 |
| red | GO:0008366 | axon ensheathment | 6.509999999999999e-27 | 9.930000000000001e-23 | 34.16 |
| red | GO:0042552 | myelination | 1.89e-25 | 9.6e-22 | 33.38 |
| red | GO:0043209 | myelin sheath | 2.53e-18 | 4.79e-15 | 14.02 |
| red | GO:0016020 | membrane | 7.570000000000001e-11 | 7.18e-08 | 1.7 |
| red | GO:0014013 | regulation of gliogenesis | 5.3e-11 | 2.0199999999999998e-07 | 12.22 |
| red | GO:0022010 | central nervous system myelination | 1.94e-10 | 4.93e-07 | 63.43 |
| red | GO:0032291 | axon ensheathment in central nervous system | 1.94e-10 | 5.92e-07 | 63.43 |
| red | GO:0044444 | cytoplasmic part | 3.87e-09 | 2.45e-06 | 1.59 |
| red | GO:0005886 | plasma membrane | 1.04e-08 | 4.93e-06 | 2.01 |
| red | GO:0048709 | oligodendrocyte differentiation | 2.9e-09 | 6.31e-06 | 29.6 |
| red | GO:0043218 | compact myelin | 1.85e-08 | 7.01e-06 | 101.49 |
| red | GO:0044464 | cell part | 3.12e-08 | 9.89e-06 | 1.24 |
| red | GO:0014015 | positive regulation of gliogenesis | 9.56e-09 | 1.82e-05 | 14.83 |
| red | GO:0010001 | glial cell differentiation | 1.2e-08 | 2.0399999999999998e-05 | 14.45 |
| red | GO:0005737 | cytoplasm | 9.58e-08 | 2.6000000000000002e-05 | 1.65 |
| red | GO:0019911 | structural constituent of myelin sheath | 6.9799999999999995e-09 | 3.14e-05 | 63.43 |
| red | GO:0033270 | paranode region of axon | 1.6399999999999999e-07 | 3.8900000000000004e-05 | 37.31 |
| red | GO:0031641 | regulation of myelination | 5.8499999999999994e-08 | 8.92e-05 | 19.73 |

| | | | | | |
|---|---|---|---|---|---|
| red | GO:0006633 | fatty acid biosynthetic process | 6.7699999999999999e-08 | 9.3799999999999999e-05 | 11.89 |
| red | GO:0031643 | positive regulation of myelination | 1.6399999999999999e-07 | 0.00020800000000000001 | 37.31 |
| red | GO:0045685 | regulation of glial cell differentiation | 2.72e-07 | 0.000319 | 12.38 |
| red | GO:0033269 | internode region of axon | 1.91e-06 | 0.00040300000000000004 | 95.15 |
| red | GO:0005515 | protein binding | 3.07e-07 | 0.00069000000000000001 | 1.48 |
| black | GO:0097458 | neuron part | 6.070000000000001e-08 | 5.7600000000000004e-05 | 3.05 |
| black | GO:0044444 | cytoplasmic part | 4.1100000000000004e-08 | 7.8e-05 | 1.68 |
| black | GO:0031838 | haptoglobin-hemoglobin complex | 2.3399999999999998e-07 | 8.89e-05 | 68.98 |
| black | GO:0005833 | hemoglobin complex | 1.5e-07 | 9.46e-05 | 75.88 |
| black | GO:0043005 | neuron projection | 2.3199999999999999e-07 | 0.00011 | 3.49 |
| black | GO:0044464 | cell part | 3.83e-07 | 0.000121 | 1.26 |
| black | GO:0043209 | myelin sheath | 7.43e-07 | 0.00020099999999999998 | 8.99 |
| black | GO:0015318 | inorganic molecular entity transmembrane transporter activity | 4.55e-07 | 0.000512 | 4.25 |
| black | GO:0022857 | transmembrane transporter activity | 6.02e-07 | 0.000542 | 3.75 |
| black | GO:0015075 | ion transmembrane transporter activity | 2.46e-07 | 0.000553 | 4.19 |
| black | GO:0005515 | protein binding | 4.41e-07 | 0.000661 | 1.58 |
| black | GO:0031720 | haptoglobin binding | 1.5e-07 | 0.00067300000000000001 | 75.88 |
| black | GO:0016020 | membrane | 3.25e-06 | 0.000772 | 1.61 |
| black | GO:0005215 | transporter activity | 1.1e-06 | 0.00082100000000000001 | 3.44 |
| green | GO:0044429 | mitochondrial part | 1.82e-09 | 1.73e-06 | 3.86 |
| green | GO:0005739 | mitochondrion | 1.14e-09 | 2.17e-06 | 2.76 |
| green | GO:0044424 | intracellular part | 4.94e-09 | 3.12e-06 | 1.31 |
| green | GO:0044444 | cytoplasmic part | 7.2199999999999995e-09 | 3.43e-06 | 1.53 |
| green | GO:0043227 | membrane-bounded organelle | 1.1200000000000001e-08 | 4.25e-06 | 1.43 |
| green | GO:0043231 | intracellular membrane-bounded organelle | 1.93e-08 | 6.09e-06 | 1.45 |
| green | GO:0043229 | intracellular organelle | 3.34e-07 | 9.05e-05 | 1.35 |
| green | GO:0098798 | mitochondrial protein complex | 5.9e-07 | 0.00014 | 5.65 |
| green | GO:0043226 | organelle | 6.7099999999999999e-07 | 0.00014199999999999998 | 1.33 |
| green | GO:0044464 | cell part | 3.68e-06 | 0.000699 | 1.19 |
| turquoise | GO:0044424 | intracellular part | 1.5199999999999993e-256 | 2.889999999999999e-253 | 1.34 |
| turquoise | GO:0043226 | organelle | 2.139999999999999e-202 | 2.0299999999999993e-199 | 1.38 |
| turquoise | GO:0043229 | intracellular organelle | 2.3999999999999986e-201 | 1.5199999999999992e-198 | 1.39 |
| turquoise | GO:0043227 | membrane-bounded organelle | 2.379999999999999e-194 | 1.1299999999999996e-191 | 1.42 |
| turquoise | GO:0043231 | intracellular membrane-bounded organelle | 1.5899999999999992e-191 | 6.0399999999999975e-189 | 1.45 |
| turquoise | GO:0044464 | cell part | 3.619999999999999e-175 | 1.1399999999999996e-172 | 1.22 |
| turquoise | GO:0044446 | intracellular organelle part | 4.719999999999999e-168 | 1.2799999999999995e-165 | 1.55 |
| turquoise | GO:0044422 | organelle part | 8.099999999999997e-161 | 1.9199999999999996e-158 | 1.52 |
| turquoise | GO:0044444 | cytoplasmic part | 8.189999999999997e-128 | 1.7299999999999995e-125 | 1.42 |
| turquoise | GO:0032991 | protein-containing complex | 1.6099999999999995e-114 | 3.059999999999999e-112 | 1.55 |
| turquoise | GO:0044428 | nuclear part | 2.2999999999999993e-105 | 3.9699999999999993e-103 | 1.65 |
| turquoise | GO:0005634 | nucleus | 9.999999999999997e-103 | 1.5899999999999997e-100 | 1.49 |
| turquoise | GO:0044237 | cellular metabolic process | 4.599999999999999e-96 | 7.0199999999999998e-92 | 1.4 |
| turquoise | GO:0043170 | macromolecule metabolic process | 8.8799999999999998e-95 | 6.769999999999999e-91 | 1.45 |
| turquoise | GO:0044260 | cellular macromolecule metabolic process | 1.87e-88 | 9.4999999999999997e-85 | 1.5 |
| turquoise | GO:0006807 | nitrogen compound metabolic process | 4.669999999999999e-84 | 1.7799999999999997e-80 | 1.39 |
| turquoise | GO:0005737 | cytoplasm | 1.3199999999999997e-79 | 1.93e-77 | 1.39 |
| turquoise | GO:0005488 | binding | 2.6399999999999995e-80 | 1.1899999999999998e-76 | 1.2 |
| turquoise | GO:1902494 | catalytic complex | 5.589999999999999e-78 | 7.5799999999999998e-76 | 2.06 |
| turquoise | GO:0044238 | primary metabolic process | 6.54e-79 | 1.9899999999999996e-75 | 1.35 |
| turquoise | GO:0071704 | organic substance metabolic process | 2.48e-75 | 6.2999999999999998e-72 | 1.33 |
| turquoise | GO:0008152 | metabolic process | 2.4899999999999992e-74 | 5.42e-71 | 1.31 |
| turquoise | GO:0005515 | protein binding | 6.6899999999999985e-74 | 1.4999999999999998e-70 | 1.29 |
| turquoise | GO:0044267 | cellular protein metabolic process | 2.7899999999999994e-69 | 5.309999999999999e-66 | 1.68 |
| turquoise | GO:0051641 | cellular localization | 2.3699999999999997e-67 | 4.009999999999999e-64 | 1.83 |
| turquoise | GO:0005829 | cytosol | 2.51e-59 | 3.17e-57 | 1.56 |
| turquoise | GO:0019538 | protein metabolic process | 3.4100000000000005e-59 | 5.2e-56 | 1.53 |
| turquoise | GO:1990234 | transferase complex | 2.8699999999999994e-57 | 3.4099999999999996e-55 | 2.23 |
| turquoise | GO:0033036 | macromolecule localization | 5.589999999999999e-57 | 7.75e-54 | 1.77 |
| turquoise | GO:0044451 | nucleoplasm part | 1.109999999999999e-55 | 1.24e-53 | 2.03 |
| turquoise | GO:0036211 | protein modification process | 6.149999999999999e-56 | 7.219999999999999e-53 | 1.66 |
| turquoise | GO:0006464 | cellular protein modification process | 6.149999999999999e-56 | 7.82e-53 | 1.66 |
| turquoise | GO:0008104 | protein localization | 8.9999999999999998e-56 | 9.8e-53 | 1.76 |
| turquoise | GO:0005654 | nucleoplasm | 9.5999999999999998e-53 | 1.0099999999999999e-50 | 1.68 |
| turquoise | GO:0009987 | cellular process | 4.620000000000001e-53 | 4.699999999999999e-50 | 1.15 |
| turquoise | GO:0043412 | macromolecule modification | 5.779999999999999e-53 | 5.51e-50 | 1.61 |
| turquoise | GO:0016070 | RNA metabolic process | 6.8e-53 | 6.1e-50 | 1.57 |
| turquoise | GO:0034641 | cellular nitrogen compound metabolic process | 2.64e-52 | 2.23e-49 | 1.43 |
| turquoise | GO:0046907 | intracellular transport | 5.749999999999999e-52 | 4.62e-49 | 1.93 |
| turquoise | GO:0019899 | enzyme binding | 4.8600000000000004e-52 | 7.29e-49 | 1.62 |
| turquoise | GO:0071840 | cellular component organization or biogenesis | 2.31e-51 | 1.76e-48 | 1.39 |
| turquoise | GO:0051649 | establishment of localization in cell | 2.66e-51 | 1.93e-48 | 1.84 |
| turquoise | GO:0016043 | cellular component organization | 2.2899999999999996e-49 | 1.5899999999999999e-46 | 1.39 |
| turquoise | GO:0051179 | localization | 2.76e-49 | 1.83e-46 | 1.44 |
| turquoise | GO:0090304 | nucleic acid metabolic process | 8.410000000000001e-48 | 5.34e-45 | 1.49 |
| turquoise | GO:0006139 | nucleobase-containing compound metabolic process | 1.83e-46 | 1.1099999999999999e-43 | 1.44 |
| turquoise | GO:0045184 | establishment of protein localization | 4.1499999999999994e-46 | 2.4299999999999997e-43 | 1.84 |
| turquoise | GO:0015031 | protein transport | 7.499999999999999e-46 | 4.24e-43 | 1.87 |
| turquoise | GO:0031323 | regulation of cellular metabolic process | 1.0800000000000002e-45 | 5.879999999999999e-43 | 1.33 |
| turquoise | GO:1901564 | organonitrogen compound metabolic process | 4.889999999999999e-45 | 2.5699999999999998e-42 | 1.39 |
| turquoise | GO:0046483 | heterocycle metabolic process | 1.5699999999999998e-44 | 7.969999999999999e-42 | 1.42 |
| turquoise | GO:0051128 | regulation of cellular component organization | 1.949999999999999e-44 | 9.5899999999999998e-42 | 1.54 |
| turquoise | GO:0019222 | regulation of metabolic process | 4.379999999999999e-44 | 2.0800000000000003e-41 | 1.31 |
| turquoise | GO:0015833 | peptide transport | 2.0800000000000002e-43 | 9.6299999999999998e-41 | 1.83 |
| turquoise | GO:0080090 | regulation of primary metabolic process | 2.99e-43 | 1.3400000000000001e-40 | 1.33 |
| turquoise | GO:0051171 | regulation of nitrogen compound metabolic process | 5.12e-43 | 2.23e-40 | 1.33 |
| turquoise | GO:0006725 | cellular aromatic compound metabolic process | 1.83e-42 | 7.74e-40 | 1.41 |
| turquoise | GO:1901363 | heterocyclic compound binding | 6.9800000000000005e-43 | 7.8499999999999998e-40 | 1.34 |
| turquoise | GO:0042886 | amide transport | 9.5e-42 | 3.91e-39 | 1.81 |
| turquoise | GO:0060255 | regulation of macromolecule metabolic process | 1.6699999999999997e-41 | 6.719999999999999e-39 | 1.32 |
| turquoise | GO:0006511 | ubiquitin-dependent protein catabolic process | 2.0800000000000003e-41 | 8.12e-39 | 2.4 |
| turquoise | GO:0097159 | organic cyclic compound binding | 1.35e-41 | 1.22e-38 | 1.33 |
| turquoise | GO:0003723 | RNA binding | 1.73e-41 | 1.3e-38 | 1.87 |
| turquoise | GO:0019941 | modification-dependent protein catabolic process | 3.7400000000000004e-40 | 1.4199999999999998e-37 | 2.35 |
| turquoise | GO:0043632 | modification-dependent macromolecule catabolic process | 9.98e-40 | 3.71e-37 | 2.32 |
| turquoise | GO:1901360 | organic cyclic compound metabolic process | 2.03e-39 | 7.3800000000000005e-37 | 1.38 |
| turquoise | GO:1990904 | ribonucleoprotein complex | 2.7999999999999995e-38 | 2.79e-36 | 1.9 |
| turquoise | GO:0051234 | establishment of localization | 8.909999999999999e-39 | 3.16e-36 | 1.42 |
| turquoise | GO:0071705 | nitrogen compound transport | 9.689999999999999e-39 | 3.3599999999999995e-36 | 1.69 |
| turquoise | GO:0070647 | protein modification by small protein conjugation or removal | 1.0499999999999998e-38 | 3.549999999999999e-36 | 2.15 |
| turquoise | GO:0006810 | transport | 1.56e-38 | 5.18e-36 | 1.43 |
| turquoise | GO:0006396 | RNA processing | 6.81e-38 | 2.2100000000000002e-35 | 2.02 |
| turquoise | GO:0019219 | regulation of nucleobase-containing compound metabolic process | 7.71e-38 | 2.4499999999999997e-35 | 1.4 |
| turquoise | GO:0097458 | neuron part | 5.31e-37 | 5.04e-35 | 1.59 |
| turquoise | GO:0006397 | mRNA processing | 7.99e-37 | 2.4799999999999997e-34 | 2.41 |

| | | | | | |
|---|---|---|---|---|---|
| turquoise | GO:0008380 | RNA splicing | 2.7099999999999994e-36 | 8.25e-34 | 2.6 |
| turquoise | GO:0044456 | synapse part | 1.26e-35 | 1.14e-33 | 1.9 |
| turquoise | GO:0044271 | cellular nitrogen compound biosynthetic process | 4.0999999999999995e-36 | 1.2300000000000001e-33 | 1.46 |
| turquoise | GO:0034645 | cellular macromolecule biosynthetic process | 6.47e-36 | 1.9e-33 | 1.49 |
| turquoise | GO:0051603 | proteolysis involved in cellular protein catabolic process | 7.769999999999998e-36 | 2.2399999999999996e-33 | 2.16 |
| turquoise | GO:0044265 | cellular macromolecule catabolic process | 1.5499999999999998e-35 | 4.2999999999999996e-33 | 2.05 |
| turquoise | GO:0016071 | mRNA metabolic process | 1.54e-35 | 4.3599999999999995e-33 | 2.19 |
| turquoise | GO:0043232 | intracellular non-membrane-bounded organelle | 9.009999999999998e-35 | 7.44e-33 | 1.43 |
| turquoise | GO:0043228 | non-membrane-bounded organelle | 9.009999999999998e-35 | 7.78e-33 | 1.43 |
| turquoise | GO:0016192 | vesicle-mediated transport | 7.919999999999999e-35 | 2.16e-32 | 1.81 |
| turquoise | GO:0009059 | macromolecule biosynthetic process | 9.73e-35 | 2.6e-32 | 1.47 |
| turquoise | GO:2000112 | regulation of cellular macromolecule biosynthetic process | 5.4399999999999996e-34 | 1.43e-31 | 1.39 |
| turquoise | GO:0051252 | regulation of RNA metabolic process | 5.92e-34 | 1.53e-31 | 1.4 |
| turquoise | GO:0010556 | regulation of macromolecule biosynthetic process | 1.3e-33 | 3.2899999999999996e-31 | 1.38 |
| turquoise | GO:0140096 | catalytic activity, acting on a protein | 2.13e-33 | 1.37e-30 | 1.5 |
| turquoise | GO:0009057 | macromolecule catabolic process | 7.21e-33 | 1.8e-30 | 1.91 |
| turquoise | GO:0010468 | regulation of gene expression | 7.549999999999999e-33 | 1.8599999999999997e-30 | 1.35 |
| turquoise | GO:0045202 | synapse | 3.1e-32 | 2.45e-30 | 1.96 |
| turquoise | GO:0016740 | transferase activity | 6.1199999999999994e-33 | 3.44e-30 | 1.49 |
| turquoise | GO:0031326 | regulation of cellular biosynthetic process | 3.8299999999999996e-32 | 9.26e-30 | 1.36 |
| turquoise | GO:0044249 | cellular biosynthetic process | 6.539999999999999e-32 | 1.5599999999999998e-29 | 1.38 |
| turquoise | GO:0006886 | intracellular protein transport | 8.31e-32 | 1.95e-29 | 1.96 |
| turquoise | GO:0009889 | regulation of biosynthetic process | 8.65e-32 | 2.0000000000000002e-29 | 1.35 |
| turquoise | GO:0070727 | cellular macromolecule localization | 9.18e-32 | 2.09e-29 | 1.82 |
| turquoise | GO:1901576 | organic substance biosynthetic process | 1.58e-31 | 3.5400000000000003e-29 | 1.37 |
| turquoise | GO:0034613 | cellular protein localization | 3.2200000000000003e-31 | 7.109999999999999e-29 | 1.82 |
| turquoise | GO:0009058 | biosynthetic process | 1.06e-30 | 2.31e-28 | 1.36 |
| turquoise | GO:0016604 | nuclear body | 4.2e-30 | 3.19e-28 | 1.93 |
| turquoise | GO:0006996 | organelle organization | 2.1599999999999998e-30 | 4.6399999999999995e-28 | 1.5 |
| turquoise | GO:0043167 | ion binding | 3.11e-30 | 1.56e-27 | 1.27 |
| turquoise | GO:0071702 | organic substance transport | 9.629999999999999e-30 | 2.04e-27 | 1.53 |
| turquoise | GO:0048522 | positive regulation of cellular process | 3.0799999999999997e-29 | 6.440000000000001e-27 | 1.28 |
| turquoise | GO:0032446 | protein modification by small protein conjugation | 3.38e-29 | 6.959999999999999e-27 | 2.11 |
| turquoise | GO:0051020 | GTPase binding | 2.5299999999999997e-28 | 1.14e-25 | 2.02 |
| turquoise | GO:0003676 | nucleic acid binding | 3.2000000000000003e-28 | 1.31e-25 | 1.37 |
| turquoise | GO:0005739 | mitochondrion | 2.13e-27 | 1.55e-25 | 1.52 |
| turquoise | GO:0032774 | RNA biosynthetic process | 8.43e-28 | 1.71e-25 | 1.48 |
| turquoise | GO:2001141 | regulation of RNA biosynthetic process | 1.33e-27 | 2.6700000000000002e-25 | 1.37 |
| turquoise | GO:0006351 | transcription, DNA-templated | 1.4699999999999998e-27 | 2.9e-25 | 1.48 |
| turquoise | GO:0097659 | nucleic acid-templated transcription | 1.71e-27 | 3.3399999999999997e-25 | 1.48 |
| turquoise | GO:1903506 | regulation of nucleic acid-templated transcription | 1.75e-27 | 3.3799999999999996e-25 | 1.37 |
| turquoise | GO:0006355 | regulation of transcription, DNA-templated | 2.27e-27 | 4.33e-25 | 1.37 |
| turquoise | GO:0018130 | heterocycle biosynthetic process | 2.53e-27 | 4.749999999999999e-25 | 1.43 |
| turquoise | GO:0034654 | nucleobase-containing compound biosynthetic process | 2.96e-27 | 5.51e-25 | 1.44 |
| turquoise | GO:0043005 | neuron projection | 9.87e-27 | 6.940000000000001e-25 | 1.61 |
| turquoise | GO:0048523 | negative regulation of cellular process | 6.23e-27 | 1.1399999999999999e-24 | 1.29 |
| turquoise | GO:0010498 | proteasomal protein catabolic process | 6.93e-27 | 1.2600000000000001e-24 | 2.35 |
| turquoise | GO:0031090 | organelle membrane | 3.7799999999999996e-26 | 2.56e-24 | 1.56 |
| turquoise | GO:0005681 | spliceosomal complex | 4.4499999999999995e-26 | 2.9100000000000002e-24 | 2.73 |
| turquoise | GO:0005794 | Golgi apparatus | 7.42e-26 | 4.700000000000001e-24 | 1.6 |
| turquoise | GO:0003824 | catalytic activity | 2.3400000000000003e-26 | 8.77e-24 | 1.24 |
| turquoise | GO:0019438 | aromatic compound biosynthetic process | 5.74e-26 | 1.0300000000000001e-23 | 1.42 |
| turquoise | GO:0016020 | membrane | 2.31e-25 | 1.4100000000000002e-23 | 1.19 |
| turquoise | GO:0048519 | negative regulation of biological process | 1.0300000000000002e-25 | 1.82e-23 | 1.26 |
| turquoise | GO:1901362 | organic cyclic compound biosynthetic process | 1.09e-25 | 1.91e-23 | 1.4 |
| turquoise | GO:0043161 | proteasome-mediated ubiquitin-dependent protein catabolic process | 2.8399999999999998e-25 | 4.92e-23 | 2.37 |
| turquoise | GO:0048518 | positive regulation of biological process | 4.51e-25 | 7.72e-23 | 1.23 |
| turquoise | GO:0030163 | protein catabolic process | 1.53e-24 | 2.5900000000000003e-22 | 2.12 |
| turquoise | GO:0016567 | protein ubiquitination | 1.77e-24 | 2.96e-22 | 2.06 |
| turquoise | GO:0033043 | regulation of organelle organization | 2.1200000000000002e-24 | 3.5099999999999997e-22 | 1.59 |
| turquoise | GO:0051130 | positive regulation of cellular component organization | 3.79e-24 | 6.22e-22 | 1.58 |
| turquoise | GO:0097060 | synaptic membrane | 3.59e-23 | 2.1299999999999998e-21 | 2.14 |
| turquoise | GO:0019787 | ubiquitin-like protein transferase activity | 1.87e-23 | 6.47e-21 | 2.03 |
| turquoise | GO:0044431 | Golgi apparatus part | 1.24e-22 | 7.1e-21 | 1.94 |
| turquoise | GO:0000151 | ubiquitin ligase complex | 1.33e-22 | 7.399999999999999e-21 | 2.26 |
| turquoise | GO:0042995 | cell projection | 1.69e-22 | 9.159999999999999e-21 | 1.42 |
| turquoise | GO:0009892 | negative regulation of metabolic process | 7.529999999999999e-23 | 1.22e-20 | 1.36 |
| turquoise | GO:0031324 | negative regulation of cellular metabolic process | 1.48e-22 | 2.37e-20 | 1.38 |
| turquoise | GO:0014069 | postsynaptic density | 7.040000000000001e-22 | 3.61e-20 | 2.25 |
| turquoise | GO:0099572 | postsynaptic specialization | 7.040000000000001e-22 | 3.7100000000000004e-20 | 2.25 |
| turquoise | GO:0000166 | nucleotide binding | 1.48e-22 | 4.4400000000000005e-20 | 1.42 |
| turquoise | GO:1901265 | nucleoside phosphate binding | 1.48e-22 | 4.76e-20 | 1.42 |
| turquoise | GO:0050794 | regulation of cellular process | 4.750000000000001e-22 | 7.55e-20 | 1.13 |
| turquoise | GO:0033365 | protein localization to organelle | 7.209999999999999e-22 | 1.1300000000000002e-19 | 1.92 |
| turquoise | GO:0009894 | regulation of catabolic process | 9.120000000000002e-22 | 1.4199999999999997e-19 | 1.71 |
| turquoise | GO:0005730 | nucleolus | 3.3799999999999995e-21 | 1.6899999999999998e-19 | 1.74 |
| turquoise | GO:0065007 | biological regulation | 1.2500000000000001e-21 | 1.92e-19 | 1.12 |
| turquoise | GO:0035639 | purine ribonucleoside triphosphate binding | 1.32e-21 | 3.72e-19 | 1.45 |
| turquoise | GO:0022607 | cellular component assembly | 2.49e-21 | 3.79e-19 | 1.4 |
| turquoise | GO:0010975 | regulation of neuron projection development | 3.46e-21 | 5.220000000000001e-19 | 1.81 |
| turquoise | GO:0050789 | regulation of biological process | 3.91e-21 | 5.85e-19 | 1.12 |
| turquoise | GO:0044433 | cytoplasmic vesicle part | 1.29e-20 | 6.26e-19 | 1.9 |
| turquoise | GO:0004842 | ubiquitin-protein transferase activity | 8.229999999999999e-21 | 2.18e-18 | 1.99 |
| turquoise | GO:0044087 | regulation of cellular component biogenesis | 1.83e-20 | 2.7199999999999998e-18 | 1.62 |
| turquoise | GO:0051173 | positive regulation of nitrogen compound metabolic process | 1.9699999999999998e-20 | 2.89e-18 | 1.32 |
| turquoise | GO:0010604 | positive regulation of macromolecule metabolic process | 2.08e-20 | 3.03e-18 | 1.31 |
| turquoise | GO:0098588 | bounding membrane of organelle | 6.92e-20 | 3.2800000000000003e-18 | 1.66 |
| turquoise | GO:0017076 | purine nucleotide binding | 1.6100000000000002e-20 | 4.03e-18 | 1.42 |
| turquoise | GO:0009893 | positive regulation of metabolic process | 3.4e-20 | 4.89e-18 | 1.29 |
| turquoise | GO:0045211 | postsynaptic membrane | 1.56e-19 | 7.2e-18 | 2.22 |
| turquoise | GO:0031325 | positive regulation of cellular metabolic process | 6.51e-20 | 9.28e-18 | 1.3 |
| turquoise | GO:0005768 | endosome | 2.15e-19 | 9.73e-18 | 1.67 |
| turquoise | GO:0032555 | purine ribonucleotide binding | 5.289999999999999e-20 | 1.25e-17 | 1.42 |
| turquoise | GO:0018193 | peptidyl-amino acid modification | 8.980000000000001e-20 | 1.27e-17 | 1.72 |
| turquoise | GO:0031267 | small GTPase binding | 8.17e-20 | 1.7499999999999998e-17 | 1.93 |
| turquoise | GO:0032553 | ribonucleotide binding | 7.81e-20 | 1.7600000000000002e-17 | 1.42 |
| turquoise | GO:0031344 | regulation of cell projection organization | 1.35e-19 | 1.8899999999999996e-17 | 1.67 |
| turquoise | GO:0120035 | regulation of plasma membrane bounded cell projection organization | 1.6199999999999999e-19 | 2.2399999999999998e-17 | 1.67 |
| turquoise | GO:0060341 | regulation of cellular localization | 1.87e-19 | 2.57e-17 | 1.63 |
| turquoise | GO:0120025 | plasma membrane bounded cell projection | 7.15e-19 | 3.15e-17 | 1.41 |
| turquoise | GO:0051172 | negative regulation of nitrogen compound metabolic process | 3.3399999999999997e-19 | 4.55e-17 | 1.36 |
| turquoise | GO:0010605 | negative regulation of macromolecule metabolic process | 5.839999999999999e-19 | 7.879999999999999e-17 | 1.34 |
| turquoise | GO:0031329 | regulation of cellular catabolic process | 6.75e-19 | 9.030000000000001e-17 | 1.71 |
| turquoise | GO:0008134 | transcription factor binding | 5.44e-19 | 1.11e-16 | 1.69 |

| | | | | | |
|---|---|---|---|---|---|
| turquoise | GO:0048193 | Golgi vesicle transport | 8.490000000000001e-19 | 1.1299999999999999e-16 | 2.24 |
| turquoise | GO:0044429 | mitochondrial part | 2.77e-18 | 1.2e-16 | 1.64 |
| turquoise | GO:0033554 | cellular response to stress | 9.28e-19 | 1.2199999999999999e-16 | 1.49 |
| turquoise | GO:0043933 | protein-containing complex subunit organization | 1.0699999999999999e-18 | 1.3999999999999998e-16 | 1.45 |
| turquoise | GO:0019904 | protein domain specific binding | 7.44e-19 | 1.45e-16 | 1.63 |
| turquoise | GO:0044432 | endoplasmic reticulum part | 4.1200000000000004e-18 | 1.74e-16 | 1.74 |
| turquoise | GO:0065009 | regulation of molecular function | 1.4200000000000001e-18 | 1.84e-16 | 1.34 |
| turquoise | GO:0016607 | nuclear speck | 5.3900000000000006e-18 | 2.22e-16 | 2.05 |
| turquoise | GO:0022604 | regulation of cell morphogenesis | 2.03e-18 | 2.6e-16 | 1.8 |
| turquoise | GO:0030425 | dendrite | 2.04e-17 | 8.22e-16 | 1.76 |
| turquoise | GO:0000398 | mRNA splicing, via spliceosome | 8.28e-18 | 1.04e-15 | 2.61 |
| turquoise | GO:0000377 | RNA splicing, via transesterification reactions with bulged adenosine as | 8.28e-18 | 1.05e-15 | 2.61 |
| turquoise | GO:0010769 | regulation of cell morphogenesis involved in differentiation | 9.56e-18 | 1.19e-15 | 1.98 |
| turquoise | GO:0000375 | RNA splicing, via transesterification reactions | 1.46e-17 | 1.8100000000000002e-15 | 2.59 |
| turquoise | GO:0006325 | chromatin organization | 1.92e-17 | 2.35e-15 | 1.7 |
| turquoise | GO:0072657 | protein localization to membrane | 1.93e-17 | 2.36e-15 | 1.94 |
| turquoise | GO:0072594 | establishment of protein localization to organelle | 2.05e-17 | 2.48e-15 | 2.17 |
| turquoise | GO:0045664 | regulation of neuron differentiation | 2.22e-17 | 2.6600000000000002e-15 | 1.63 |
| turquoise | GO:0045935 | positive regulation of nucleobase-containing compound metabolic proc | 2.64e-17 | 3.1399999999999997e-15 | 1.39 |
| turquoise | GO:0006508 | proteolysis | 3.5299999999999996e-17 | 4.17e-15 | 1.52 |
| turquoise | GO:0017016 | Ras GTPase binding | 3.03e-17 | 5.67e-15 | 1.87 |
| turquoise | GO:0030054 | cell junction | 3.22e-16 | 1.27e-14 | 1.49 |
| turquoise | GO:0098805 | whole membrane | 3.41e-16 | 1.32e-14 | 1.68 |
| turquoise | GO:0044440 | endosomal part | 5.87e-16 | 2.23e-14 | 2.17 |
| turquoise | GO:0065003 | protein-containing complex assembly | 2.36e-16 | 2.76e-14 | 1.44 |
| turquoise | GO:0036094 | small molecule binding | 1.61e-16 | 2.91e-14 | 1.32 |
| turquoise | GO:0043168 | anion binding | 2.15e-16 | 3.72e-14 | 1.3 |
| turquoise | GO:0005524 | ATP binding | 2.9900000000000003e-16 | 4.99e-14 | 1.43 |
| turquoise | GO:0071013 | catalytic step 2 spliceosome | 1.3900000000000001e-15 | 5.08e-14 | 2.95 |
| turquoise | GO:1905368 | peptidase complex | 1.3900000000000001e-15 | 5.16e-14 | 2.89 |
| turquoise | GO:0097708 | intracellular vesicle | 1.5699999999999999e-15 | 5.61e-14 | 1.38 |
| turquoise | GO:0044463 | cell projection part | 2.3e-15 | 7.8e-14 | 1.4 |
| turquoise | GO:0031410 | cytoplasmic vesicle | 2.26e-15 | 7.93e-14 | 1.38 |
| turquoise | GO:0120038 | plasma membrane bounded cell projection part | 2.3e-15 | 7.94e-14 | 1.4 |
| turquoise | GO:0098796 | membrane protein complex | 2.7600000000000003e-15 | 9.19e-14 | 1.46 |
| turquoise | GO:0003712 | transcription coregulator activity | 7.1299999999999995e-16 | 1.15e-13 | 1.78 |
| turquoise | GO:0030554 | adenyl nucleotide binding | 7.56e-16 | 1.17e-13 | 1.41 |
| turquoise | GO:0050767 | regulation of neurogenesis | 1.48e-15 | 1.7199999999999997e-13 | 1.53 |
| turquoise | GO:0050804 | modulation of chemical synaptic transmission | 1.5099999999999999e-15 | 1.74e-13 | 1.81 |
| turquoise | GO:0051246 | regulation of protein metabolic process | 1.6899999999999999e-15 | 1.9399999999999998e-13 | 1.29 |
| turquoise | GO:0010638 | positive regulation of organelle organization | 1.9100000000000003e-15 | 2.18e-13 | 1.69 |
| turquoise | GO:0099177 | regulation of trans-synaptic signaling | 1.94e-15 | 2.1899999999999998e-13 | 1.81 |
| turquoise | GO:0016570 | histone modification | 3.16e-15 | 3.5499999999999995e-13 | 1.93 |
| turquoise | GO:0032559 | adenyl ribonucleotide binding | 2.6600000000000002e-15 | 3.989999999999999e-13 | 1.4 |
| turquoise | GO:0046872 | metal ion binding | 2.8800000000000003e-15 | 4.18e-13 | 1.24 |
| turquoise | GO:0060284 | regulation of cell development | 4.07e-15 | 4.53e-13 | 1.48 |
| turquoise | GO:0043021 | ribonucleoprotein complex binding | 3.71e-15 | 5.22e-13 | 2.43 |
| turquoise | GO:0016569 | covalent chromatin modification | 6.19e-15 | 6.840000000000001e-13 | 1.9 |
| turquoise | GO:0023051 | regulation of signaling | 6.640000000000001e-15 | 7.290000000000001e-13 | 1.26 |
| turquoise | GO:0031982 | vesicle | 2.24e-14 | 7.32e-13 | 1.35 |
| turquoise | GO:0034622 | cellular protein-containing complex assembly | 9.6e-15 | 1.05e-12 | 1.6 |
| turquoise | GO:0010646 | regulation of cell communication | 1.15e-14 | 1.24e-12 | 1.26 |
| turquoise | GO:0032268 | regulation of cellular protein metabolic process | 1.19e-14 | 1.28e-12 | 1.3 |
| turquoise | GO:0050773 | regulation of dendrite development | 1.21e-14 | 1.29e-12 | 2.22 |
| turquoise | GO:0030532 | small nuclear ribonucleoprotein complex | 4e-14 | 1.29e-12 | 3.15 |
| turquoise | GO:0032879 | regulation of localization | 1.31e-14 | 1.38e-12 | 1.28 |
| turquoise | GO:0044248 | cellular catabolic process | 1.38e-14 | 1.45e-12 | 1.41 |
| turquoise | GO:0051254 | positive regulation of RNA metabolic process | 1.75e-14 | 1.83e-12 | 1.37 |
| turquoise | GO:1903827 | regulation of cellular protein localization | 1.94e-14 | 2.02e-12 | 1.71 |
| turquoise | GO:0043254 | regulation of protein complex assembly | 1.97e-14 | 2.0299999999999996e-12 | 1.76 |
| turquoise | GO:0010557 | positive regulation of macromolecule biosynthetic process | 2.17e-14 | 2.2200000000000002e-12 | 1.35 |
| turquoise | GO:0043025 | neuronal cell body | 7.24e-14 | 2.29e-12 | 1.62 |
| turquoise | GO:0005783 | endoplasmic reticulum | 7.54e-14 | 2.3500000000000003e-12 | 1.37 |
| turquoise | GO:0051960 | regulation of nervous system development | 2.54e-14 | 2.58e-12 | 1.48 |
| turquoise | GO:0032990 | cell part morphogenesis | 2.56e-14 | 2.59e-12 | 1.94 |
| turquoise | GO:0034708 | methyltransferase complex | 8.81e-14 | 2.7000000000000002e-12 | 2.48 |
| turquoise | GO:0044309 | neuron spine | 1.35e-13 | 4.07e-12 | 2.17 |
| turquoise | GO:0044297 | cell body | 1.6100000000000002e-13 | 4.769999999999999e-12 | 1.56 |
| turquoise | GO:0010976 | positive regulation of neuron projection development | 6.16e-14 | 6.179999999999999e-12 | 1.83 |
| turquoise | GO:0051640 | organelle localization | 6.42e-14 | 6.4e-12 | 1.77 |
| turquoise | GO:0061136 | regulation of proteasomal protein catabolic process | 6.48e-14 | 6.4099999999999995e-12 | 2.23 |
| turquoise | GO:0097525 | spliceosomal snRNP complex | 2.35e-13 | 6.8700000000000004e-12 | 3.18 |
| turquoise | GO:0043169 | cation binding | 6.08e-14 | 8.28e-12 | 1.23 |
| turquoise | GO:0031647 | regulation of protein stability | 8.98e-14 | 8.83e-12 | 1.98 |
| turquoise | GO:0051716 | cellular response to stimulus | 9.15e-14 | 8.949999999999999e-12 | 1.28 |
| turquoise | GO:0000209 | protein polyubiquitination | 9.56e-14 | 9.28e-12 | 2.21 |
| turquoise | GO:1901575 | organic substance catabolic process | 1.15e-13 | 1.1099999999999999e-11 | 1.4 |
| turquoise | GO:0098798 | mitochondrial protein complex | 3.9299999999999994e-13 | 1.13e-11 | 1.96 |
| turquoise | GO:0010628 | positive regulation of gene expression | 1.3e-13 | 1.25e-11 | 1.32 |
| turquoise | GO:1903050 | regulation of proteolysis involved in cellular protein catabolic process | 1.55e-13 | 1.48e-11 | 2.11 |
| turquoise | GO:0048858 | cell projection morphogenesis | 1.88e-13 | 1.7800000000000003e-11 | 1.94 |
| turquoise | GO:0097367 | carbohydrate derivative binding | 1.3900000000000001e-13 | 1.84e-11 | 1.31 |
| turquoise | GO:0120114 | Sm-like protein family complex | 6.490000000000001e-13 | 1.84e-11 | 2.86 |
| turquoise | GO:0005684 | U2-type spliceosomal complex | 9.57e-13 | 2.6700000000000002e-11 | 2.95 |
| turquoise | GO:0006357 | regulation of transcription by RNA polymerase II | 3.16e-13 | 2.97e-11 | 1.32 |
| turquoise | GO:0060627 | regulation of vesicle-mediated transport | 3.4399999999999995e-13 | 3.22e-11 | 1.69 |
| turquoise | GO:0009891 | positive regulation of biosynthetic process | 3.9500000000000003e-13 | 3.67e-11 | 1.32 |
| turquoise | GO:0120039 | plasma membrane bounded cell projection morphogenesis | 4.08e-13 | 3.77e-11 | 1.94 |
| turquoise | GO:0031346 | positive regulation of cell projection organization | 4.560000000000001e-13 | 4.19e-11 | 1.7 |
| turquoise | GO:0050684 | regulation of mRNA processing | 4.92e-13 | 4.49e-11 | 2.33 |
| turquoise | GO:0009056 | catabolic process | 5.080000000000001e-13 | 4.5599999999999996e-11 | 1.35 |
| turquoise | GO:0031328 | positive regulation of cellular biosynthetic process | 5.080000000000001e-13 | 4.58e-11 | 1.32 |
| turquoise | GO:0003682 | chromatin binding | 3.570000000000001e-13 | 4.589999999999999e-11 | 1.61 |
| turquoise | GO:1903362 | regulation of cellular protein catabolic process | 5.07e-13 | 4.5999999999999996e-11 | 2.0 |
| turquoise | GO:0042176 | regulation of protein catabolic process | 6.73e-13 | 6e-11 | 1.78 |
| turquoise | GO:0060998 | regulation of dendritic spine development | 7.91e-13 | 7.01e-11 | 2.66 |
| turquoise | GO:0022618 | ribonucleoprotein complex assembly | 8.840000000000001e-13 | 7.790000000000001e-11 | 2.05 |
| turquoise | GO:0043197 | dendritic spine | 3.0800000000000002e-12 | 8.47e-11 | 2.11 |
| turquoise | GO:0005762 | mitochondrial large ribosomal subunit | 3.88e-12 | 1.0399999999999999e-10 | 3.04 |
| turquoise | GO:0000315 | organellar large ribosomal subunit | 3.88e-12 | 1.0500000000000001e-10 | 3.04 |
| turquoise | GO:0010629 | negative regulation of gene expression | 1.23e-12 | 1.08e-10 | 1.35 |
| turquoise | GO:0036503 | ERAD pathway | 1.28e-12 | 1.12e-10 | 2.82 |
| turquoise | GO:0048812 | neuron projection morphogenesis | 1.29e-12 | 1.12e-10 | 1.93 |
| turquoise | GO:0045666 | positive regulation of neuron differentiation | 1.6899999999999999e-12 | 1.46e-10 | 1.68 |

| | | | | | |
|---|---|---|---|---|---|
| turquoise | GO:0031461 | cullin-RING ubiquitin ligase complex | 5.63e-12 | 1.48e-10 | 2.16 |
| turquoise | GO:0043484 | regulation of RNA splicing | 1.76e-12 | 1.51e-10 | 2.31 |
| turquoise | GO:0048167 | regulation of synaptic plasticity | 1.93e-12 | 1.6299999999999998e-10 | 2.0 |
| turquoise | GO:1903311 | regulation of mRNA metabolic process | 1.93e-12 | 1.64e-10 | 1.97 |
| turquoise | GO:0051049 | regulation of transport | 2.1e-12 | 1.77e-10 | 1.31 |
| turquoise | GO:0044089 | positive regulation of cellular component biogenesis | 2.17e-12 | 1.8200000000000002e-10 | 1.64 |
| turquoise | GO:0006412 | translation | 2.29e-12 | 1.9099999999999999e-10 | 1.89 |
| turquoise | GO:0018205 | peptidyl-lysine modification | 2.45e-12 | 2.03e-10 | 1.99 |
| turquoise | GO:0098794 | postsynapse | 7.89e-12 | 2.0500000000000002e-10 | 2.01 |
| turquoise | GO:0009896 | positive regulation of catabolic process | 2.62e-12 | 2.16e-10 | 1.72 |
| turquoise | GO:0071826 | ribonucleoprotein complex subunit organization | 2.65e-12 | 2.17e-10 | 2.0 |
| turquoise | GO:1902680 | positive regulation of RNA biosynthetic process | 2.7300000000000003e-12 | 2.23e-10 | 1.35 |
| turquoise | GO:0045893 | positive regulation of transcription, DNA-templated | 3.0699999999999996e-12 | 2.4899999999999997e-10 | 1.35 |
| turquoise | GO:0033044 | regulation of chromosome organization | 3.11e-12 | 2.51e-10 | 1.78 |
| turquoise | GO:0071011 | precatalytic spliceosome | 1.0800000000000001e-11 | 2.7799999999999997e-10 | 4.47 |
| turquoise | GO:0007005 | mitochondrion organization | 3.69e-12 | 2.96e-10 | 1.89 |
| turquoise | GO:1903508 | positive regulation of nucleic acid-templated transcription | 3.8e-12 | 3.03e-10 | 1.34 |
| turquoise | GO:0044877 | protein-containing complex binding | 2.54e-12 | 3.17e-10 | 1.4 |
| turquoise | GO:0050890 | cognition | 4.14e-12 | 3.29e-10 | 1.81 |
| turquoise | GO:0006793 | phosphorus metabolic process | 4.2699999999999999e-12 | 3.37e-10 | 1.33 |
| turquoise | GO:0031984 | organelle subcompartment | 1.6699999999999998e-11 | 4.22e-10 | 1.88 |
| turquoise | GO:0006413 | translational initiation | 5.43e-12 | 4.27e-10 | 3.06 |
| turquoise | GO:0008144 | drug binding | 3.68e-12 | 4.4699999999999996e-10 | 1.33 |
| turquoise | GO:0043043 | peptide biosynthetic process | 5.9100000000000005e-12 | 4.6199999999999996e-10 | 1.85 |
| turquoise | GO:0006796 | phosphate-containing compound metabolic process | 6.27e-12 | 4.880000000000001e-10 | 1.33 |
| turquoise | GO:0019900 | kinase binding | 4.64e-12 | 5.49e-10 | 1.49 |
| turquoise | GO:0006888 | ER to Golgi vesicle-mediated transport | 8.210000000000002e-12 | 6.35e-10 | 2.38 |
| turquoise | GO:0043604 | amide biosynthetic process | 8.64e-12 | 6.650000000000001e-10 | 1.73 |
| turquoise | GO:0070646 | protein modification by small protein removal | 8.91e-12 | 6.8199999999999999e-10 | 2.34 |
| turquoise | GO:1900006 | positive regulation of dendrite development | 1.1099999999999999e-11 | 8.43e-10 | 2.43 |
| turquoise | GO:0030030 | cell projection organization | 1.16e-11 | 8.77e-10 | 1.46 |
| turquoise | GO:0061695 | transferase complex, transferring phosphorus-containing groups | 3.9500000000000006e-11 | 9.87e-10 | 1.94 |
| turquoise | GO:0010506 | regulation of autophagy | 1.35e-11 | 1.02e-09 | 1.95 |
| turquoise | GO:0061659 | ubiquitin-like protein ligase activity | 9.169999999999999e-12 | 1.06e-09 | 1.96 |
| turquoise | GO:0032880 | regulation of protein localization | 1.48e-11 | 1.11e-09 | 1.43 |
| turquoise | GO:1905369 | endopeptidase complex | 4.8700000000000004e-11 | 1.2e-09 | 2.82 |
| turquoise | GO:1902115 | regulation of organelle assembly | 1.7699999999999998e-11 | 1.32e-09 | 2.05 |
| turquoise | GO:0051129 | negative regulation of cellular component organization | 1.93e-11 | 1.4400000000000002e-09 | 1.5 |
| turquoise | GO:0045934 | negative regulation of nucleobase-containing compound metabolic pro | 2.04e-11 | 1.51e-09 | 1.35 |
| turquoise | GO:0008565 | protein transporter activity | 1.53e-11 | 1.7199999999999999e-09 | 2.78 |
| turquoise | GO:0009895 | negative regulation of catabolic process | 2.47e-11 | 1.82e-09 | 1.84 |
| turquoise | GO:0030234 | enzyme regulator activity | 1.7800000000000003e-11 | 1.95e-09 | 1.44 |
| turquoise | GO:0051098 | regulation of binding | 2.7399999999999997e-11 | 2.0100000000000003e-09 | 1.69 |
| turquoise | GO:0016772 | transferase activity, transferring phosphorus-containing groups | 2.01e-11 | 2.15e-09 | 1.44 |
| turquoise | GO:0048814 | regulation of dendrite morphogenesis | 3.01e-11 | 2.2e-09 | 2.36 |
| turquoise | GO:0044093 | positive regulation of molecular function | 3.45e-11 | 2.5e-09 | 1.34 |
| turquoise | GO:0000139 | Golgi membrane | 1.06e-10 | 2.57e-09 | 2.01 |
| turquoise | GO:0030433 | ubiquitin-dependent ERAD pathway | 3.59e-11 | 2.59e-09 | 2.88 |
| turquoise | GO:1901565 | organonitrogen compound catabolic process | 3.9100000000000006e-11 | 2.8199999999999998e-09 | 1.53 |
| turquoise | GO:0000502 | proteasome complex | 1.3899999999999999e-10 | 3.25e-09 | 2.79 |
| turquoise | GO:0005763 | mitochondrial small ribosomal subunit | 1.38e-10 | 3.27e-09 | 3.71 |
| turquoise | GO:0000314 | organellar small ribosomal subunit | 1.38e-10 | 3.3100000000000004e-09 | 3.71 |
| turquoise | GO:0016197 | endosomal transport | 4.73e-11 | 3.3799999999999995e-09 | 2.07 |
| turquoise | GO:0050769 | positive regulation of neurogenesis | 4.77e-11 | 3.4000000000000003e-09 | 1.56 |
| turquoise | GO:0031400 | negative regulation of protein modification process | 5.1999999999999995e-11 | 3.69e-09 | 1.57 |
| turquoise | GO:0032269 | negative regulation of cellular protein metabolic process | 5.2500000000000006e-11 | 3.71e-09 | 1.42 |
| turquoise | GO:0043543 | protein acylation | 5.55e-11 | 3.9e-09 | 2.1 |
| turquoise | GO:0031331 | positive regulation of cellular catabolic process | 5.83e-11 | 4.08e-09 | 1.74 |
| turquoise | GO:0060589 | nucleoside-triphosphatase regulator activity | 3.96e-11 | 4.15e-09 | 1.8 |
| turquoise | GO:0006457 | protein folding | 7.05e-11 | 4.91e-09 | 2.17 |
| turquoise | GO:0005856 | cytoskeleton | 2.21e-10 | 5.13e-09 | 1.32 |
| turquoise | GO:0061630 | ubiquitin protein ligase activity | 5.31e-11 | 5.420000000000001e-09 | 1.93 |
| turquoise | GO:0031327 | negative regulation of cellular biosynthetic process | 8.07e-11 | 5.589999999999999e-09 | 1.33 |
| turquoise | GO:0009966 | regulation of signal transduction | 9.000000000000001e-11 | 6.21e-09 | 1.23 |
| turquoise | GO:0016241 | regulation of macroautophagy | 9.549999999999999e-11 | 6.559999999999999e-09 | 2.41 |
| turquoise | GO:0032989 | cellular component morphogenesis | 1.0200000000000001e-10 | 6.9799999999999995e-09 | 1.58 |
| turquoise | GO:0051962 | positive regulation of nervous system development | 1.09e-10 | 7.41e-09 | 1.52 |
| turquoise | GO:0005769 | early endosome | 3.59e-10 | 8.209999999999999e-09 | 1.91 |
| turquoise | GO:0004674 | protein serine/threonine kinase activity | 8.26e-11 | 8.26e-09 | 1.63 |
| turquoise | GO:0051248 | negative regulation of protein metabolic process | 1.26e-10 | 8.51e-09 | 1.39 |
| turquoise | GO:0065008 | regulation of biological quality | 1.31e-10 | 8.84e-09 | 1.2 |
| turquoise | GO:0050821 | protein stabilization | 1.5800000000000002e-10 | 1.06e-08 | 2.06 |
| turquoise | GO:0016579 | protein deubiquitination | 1.6099999999999999e-10 | 1.07e-08 | 2.36 |
| turquoise | GO:0031399 | regulation of protein modification process | 1.68e-10 | 1.1200000000000001e-08 | 1.29 |
| turquoise | GO:0031248 | protein acetyltransferase complex | 5.09e-10 | 1.14e-08 | 2.41 |
| turquoise | GO:0008135 | translation factor activity, RNA binding | 1.1599999999999999e-10 | 1.14e-08 | 2.62 |
| turquoise | GO:1902493 | acetyltransferase complex | 5.09e-10 | 1.1499999999999999e-08 | 2.41 |
| turquoise | GO:0098791 | Golgi subcompartment | 5.29e-10 | 1.1699999999999999e-08 | 1.84 |
| turquoise | GO:0035097 | histone methyltransferase complex | 5.39e-10 | 1.18e-08 | 2.39 |
| turquoise | GO:1902532 | negative regulation of intracellular signal transduction | 1.8700000000000003e-10 | 1.24e-08 | 1.57 |
| turquoise | GO:0000381 | regulation of alternative mRNA splicing, via spliceosome | 2.0500000000000002e-10 | 1.35e-08 | 2.8 |
| turquoise | GO:0051253 | negative regulation of RNA metabolic process | 2.18e-10 | 1.4299999999999999e-08 | 1.35 |
| turquoise | GO:0007611 | learning or memory | 2.38e-10 | 1.56e-08 | 1.78 |
| turquoise | GO:0031625 | ubiquitin protein ligase binding | 1.7e-10 | 1.62e-08 | 1.76 |
| turquoise | GO:0044452 | nucleolar part | 8.110000000000001e-10 | 1.75e-08 | 1.96 |
| turquoise | GO:0032386 | regulation of intracellular transport | 2.7000000000000005e-10 | 1.76e-08 | 1.68 |
| turquoise | GO:2000058 | regulation of ubiquitin-dependent protein catabolic process | 2.96e-10 | 1.92e-08 | 2.1 |
| turquoise | GO:0071005 | U2-type precatalytic spliceosome | 9.71e-10 | 2.0699999999999997e-08 | 4.43 |
| turquoise | GO:0010720 | positive regulation of cell development | 3.39e-10 | 2.19e-08 | 1.5 |
| turquoise | GO:0019901 | protein kinase binding | 2.55e-10 | 2.3900000000000002e-08 | 1.47 |
| turquoise | GO:0044389 | ubiquitin-like protein ligase binding | 2.66e-10 | 2.44e-08 | 1.73 |
| turquoise | GO:0009890 | negative regulation of biosynthetic process | 4.17e-10 | 2.66e-08 | 1.32 |
| turquoise | GO:0045936 | negative regulation of phosphate metabolic process | 4.15e-10 | 2.66e-08 | 1.54 |
| turquoise | GO:0010563 | negative regulation of phosphorus metabolic process | 4.15e-10 | 2.67e-08 | 1.54 |
| turquoise | GO:0051726 | regulation of cell cycle | 4.21e-10 | 2.6800000000000002e-08 | 1.4 |
| turquoise | GO:0006518 | peptide metabolic process | 4.5600000000000003e-10 | 2.89e-08 | 1.64 |
| turquoise | GO:0016301 | kinase activity | 3.4299999999999995e-10 | 3.09e-08 | 1.45 |
| turquoise | GO:0031330 | negative regulation of cellular catabolic process | 5e-10 | 3.15e-08 | 1.86 |
| turquoise | GO:2000060 | positive regulation of ubiquitin-dependent protein catabolic process | 5.39e-10 | 3.38e-08 | 2.39 |
| turquoise | GO:0005096 | GTPase activator activity | 4.06e-10 | 3.5099999999999997e-08 | 1.85 |
| turquoise | GO:0030695 | GTPase regulator activity | 3.9899999999999997e-10 | 3.52e-08 | 1.82 |
| turquoise | GO:0048024 | regulation of mRNA splicing, via spliceosome | 5.84e-10 | 3.6499999999999996e-08 | 2.34 |
| turquoise | GO:0032434 | regulation of proteasomal ubiquitin-dependent protein catabolic proces | 5.97e-10 | 3.71e-08 | 2.2 |
| turquoise | GO:2000113 | negative regulation of cellular macromolecule biosynthetic process | 6.08e-10 | 3.77e-08 | 1.33 |

| | | | | | |
|---|---|---|---|---|---|
| turquoise | GO:0007610 | behavior | 6.55e-10 | 4.04e-08 | 1.48 |
| turquoise | GO:0005689 | U12-type spliceosomal complex | 2.09e-09 | 4.4100000000000004e-08 | 3.74 |
| turquoise | GO:0005802 | trans-Golgi network | 2.17e-09 | 4.53e-08 | 1.93 |
| turquoise | GO:0031966 | mitochondrial membrane | 2.36e-09 | 4.8600000000000005e-08 | 1.52 |
| turquoise | GO:1903052 | positive regulation of proteolysis involved in cellular protein catabolic pr | 9.300000000000001e-10 | 5.71e-08 | 2.23 |
| turquoise | GO:0051082 | unfolded protein binding | 7.22e-10 | 6.13e-08 | 2.45 |
| turquoise | GO:0016773 | phosphotransferase activity, alcohol group as acceptor | 7.38e-10 | 6.15e-08 | 1.46 |
| turquoise | GO:0048471 | perinuclear region of cytoplasm | 3.09e-09 | 6.3e-08 | 1.47 |
| turquoise | GO:0032182 | ubiquitin-like protein binding | 7.74e-10 | 6.33e-08 | 2.42 |
| turquoise | GO:0003743 | translation initiation factor activity | 9.6e-10 | 7.58e-08 | 2.98 |
| turquoise | GO:0140098 | catalytic activity, acting on RNA | 9.52e-10 | 7.65e-08 | 1.72 |
| turquoise | GO:0045595 | regulation of cell differentiation | 1.59e-09 | 9.720000000000001e-08 | 1.27 |
| turquoise | GO:0010770 | positive regulation of cell morphogenesis involved in differentiation | 1.7600000000000001e-09 | 1.08e-07 | 1.95 |
| turquoise | GO:0070936 | protein K48-linked ubiquitination | 1.87e-09 | 1.13e-07 | 2.82 |
| turquoise | GO:0034248 | regulation of cellular amide metabolic process | 1.88e-09 | 1.14e-07 | 1.66 |
| turquoise | GO:0044427 | chromosomal part | 5.83e-09 | 1.1800000000000001e-07 | 1.38 |
| turquoise | GO:0060999 | positive regulation of dendritic spine development | 2e-09 | 1.2099999999999998e-07 | 2.68 |
| turquoise | GO:0008047 | enzyme activator activity | 1.6300000000000002e-09 | 1.26e-07 | 1.59 |
| turquoise | GO:0034976 | response to endoplasmic reticulum stress | 2.17e-09 | 1.3e-07 | 1.93 |
| turquoise | GO:0022624 | proteasome accessory complex | 6.620000000000001e-09 | 1.32e-07 | 4.17 |
| turquoise | GO:0010008 | endosome membrane | 6.709999999999999e-09 | 1.33e-07 | 1.95 |
| turquoise | GO:0010558 | negative regulation of macromolecule biosynthetic process | 2.27e-09 | 1.36e-07 | 1.31 |
| turquoise | GO:0034703 | cation channel complex | 7.65e-09 | 1.5e-07 | 1.81 |
| turquoise | GO:2001252 | positive regulation of chromosome organization | 2.6e-09 | 1.55e-07 | 1.94 |
| turquoise | GO:0006913 | nucleocytoplasmic transport | 2.65e-09 | 1.57e-07 | 1.92 |
| turquoise | GO:0004672 | protein kinase activity | 2.27e-09 | 1.73e-07 | 1.48 |
| turquoise | GO:0007034 | vacuolar transport | 2.95e-09 | 1.74e-07 | 2.16 |
| turquoise | GO:0090150 | establishment of protein localization to membrane | 2.97e-09 | 1.75e-07 | 2.05 |
| turquoise | GO:1902275 | regulation of chromatin organization | 3.2e-09 | 1.8800000000000002e-07 | 1.9 |
| turquoise | GO:0031056 | regulation of histone modification | 3.37e-09 | 1.97e-07 | 2.01 |
| turquoise | GO:0023057 | negative regulation of signaling | 3.42e-09 | 1.99e-07 | 1.32 |
| turquoise | GO:0010648 | negative regulation of cell communication | 3.4700000000000002e-09 | 2.01e-07 | 1.32 |
| turquoise | GO:0061650 | ubiquitin-like protein conjugating enzyme activity | 2.72e-09 | 2.0399999999999997e-07 | 3.07 |
| turquoise | GO:0051169 | nuclear transport | 3.53e-09 | 2.0399999999999997e-07 | 1.9 |
| turquoise | GO:0007399 | nervous system development | 3.71e-09 | 2.1300000000000001e-07 | 1.61 |
| turquoise | GO:0050770 | regulation of axonogenesis | 3.7200000000000004e-09 | 2.1300000000000001e-07 | 1.86 |
| turquoise | GO:0099023 | tethering complex | 1.1000000000000001e-08 | 2.1300000000000001e-07 | 2.55 |
| turquoise | GO:0043198 | dendritic shaft | 1.23e-08 | 2.3600000000000002e-07 | 2.5 |
| turquoise | GO:0007612 | learning | 4.56e-09 | 2.6e-07 | 1.95 |
| turquoise | GO:0031072 | heat shock protein binding | 3.55e-09 | 2.62e-07 | 2.09 |
| turquoise | GO:0061631 | ubiquitin conjugating enzyme activity | 3.8e-09 | 2.75e-07 | 3.11 |
| turquoise | GO:0061024 | membrane organization | 5.06e-09 | 2.88e-07 | 1.52 |
| turquoise | GO:0019902 | phosphatase binding | 4.13e-09 | 2.95e-07 | 1.86 |
| turquoise | GO:0032008 | positive regulation of TOR signaling | 5.24e-09 | 2.9699999999999997e-07 | 3.15 |
| turquoise | GO:0080135 | regulation of cellular response to stress | 5.44e-09 | 3.07e-07 | 1.47 |
| turquoise | GO:0050790 | regulation of catalytic activity | 6.06e-09 | 3.41e-07 | 1.26 |
| turquoise | GO:0000785 | chromatin | 1.88e-08 | 3.56e-07 | 1.55 |
| turquoise | GO:0051656 | establishment of organelle localization | 6.85e-09 | 3.84e-07 | 1.71 |
| turquoise | GO:0044403 | symbiont process | 7.13e-09 | 3.97e-07 | 2.18 |
| turquoise | GO:0016032 | viral process | 7.13e-09 | 3.98e-07 | 2.18 |
| turquoise | GO:0005840 | ribosome | 2.13e-08 | 3.9899999999999996e-07 | 1.83 |
| turquoise | GO:1903364 | positive regulation of cellular protein catabolic process | 7.52e-09 | 4.15e-07 | 2.07 |
| turquoise | GO:1901800 | positive regulation of proteasomal protein catabolic process | 7.5e-09 | 4.16e-07 | 2.24 |
| turquoise | GO:0033267 | axon part | 2.2999999999999998e-08 | 4.2799999999999997e-07 | 1.54 |
| turquoise | GO:0050807 | regulation of synapse organization | 8.15e-09 | 4.48e-07 | 1.87 |
| turquoise | GO:0032271 | regulation of protein polymerization | 8.22e-09 | 4.51e-07 | 1.83 |
| turquoise | GO:0000123 | histone acetyltransferase complex | 2.54e-08 | 4.69e-07 | 2.33 |
| turquoise | GO:0006417 | regulation of translation | 8.740000000000001e-09 | 4.78e-07 | 1.68 |
| turquoise | GO:0032535 | regulation of cellular component size | 9.63e-09 | 5.24e-07 | 1.71 |
| turquoise | GO:0051668 | localization within membrane | 9.72e-09 | 5.27e-07 | 2.13 |
| turquoise | GO:0050775 | positive regulation of dendrite morphogenesis | 1e-08 | 5.420000000000001e-07 | 2.72 |
| turquoise | GO:0003729 | mRNA binding | 8.120000000000001e-09 | 5.7e-07 | 1.76 |
| turquoise | GO:0008328 | ionotropic glutamate receptor complex | 3.3e-08 | 6.03e-07 | 2.62 |
| turquoise | GO:0034660 | ncRNA metabolic process | 1.23e-08 | 6.64e-07 | 1.57 |
| turquoise | GO:0044448 | cell cortex part | 4.24e-08 | 7.66e-07 | 1.84 |
| turquoise | GO:0018107 | peptidyl-threonine phosphorylation | 1.46e-08 | 7.83e-07 | 2.43 |
| turquoise | GO:0043022 | ribosome binding | 1.23e-08 | 8.49e-07 | 2.66 |
| turquoise | GO:0005770 | late endosome | 5.0599999999999996e-08 | 9.06e-07 | 1.93 |
| turquoise | GO:0051247 | positive regulation of protein metabolic process | 1.74e-08 | 9.33e-07 | 1.27 |
| turquoise | GO:0007409 | axonogenesis | 1.77e-08 | 9.42e-07 | 2.0 |
| turquoise | GO:0035257 | nuclear hormone receptor binding | 1.3899999999999999e-08 | 9.49e-07 | 1.97 |
| turquoise | GO:0031334 | positive regulation of protein complex assembly | 1.8300000000000002e-08 | 9.73e-07 | 1.78 |
| turquoise | GO:0065002 | intracellular protein transmembrane transport | 2.2200000000000004e-08 | 1.18e-06 | 2.75 |
| turquoise | GO:1903320 | regulation of protein modification by small protein conjugation or remov | 2.2999999999999998e-08 | 1.22e-06 | 1.78 |
| turquoise | GO:0043130 | ubiquitin binding | 2.06e-08 | 1.38e-06 | 2.47 |
| turquoise | GO:0030117 | membrane coat | 7.83e-08 | 1.39e-06 | 2.44 |
| turquoise | GO:0008270 | zinc ion binding | 2.32e-08 | 1.49e-06 | 1.42 |
| turquoise | GO:0003714 | transcription corepressor activity | 2.25e-08 | 1.49e-06 | 1.85 |
| turquoise | GO:0036002 | pre-mRNA binding | 2.3100000000000002e-08 | 1.5e-06 | 3.15 |
| turquoise | GO:0036452 | ESCRT complex | 8.99e-08 | 1.58e-06 | 3.49 |
| turquoise | GO:0061919 | process utilizing autophagic mechanism | 3.26e-08 | 1.71e-06 | 1.85 |
| turquoise | GO:0006914 | autophagy | 3.26e-08 | 1.71e-06 | 1.85 |
| turquoise | GO:0061001 | regulation of dendritic spine morphogenesis | 3.31e-08 | 1.73e-06 | 2.77 |
| turquoise | GO:0098878 | neurotransmitter receptor complex | 9.96e-08 | 1.73e-06 | 2.53 |
| turquoise | GO:0016310 | phosphorylation | 3.35e-08 | 1.74e-06 | 1.35 |
| turquoise | GO:0030424 | axon | 1.04e-07 | 1.79e-06 | 1.52 |
| turquoise | GO:0032270 | positive regulation of cellular protein metabolic process | 3.6e-08 | 1.86e-06 | 1.27 |
| turquoise | GO:0032436 | positive regulation of proteasomal ubiquitin-dependent protein cataboli | 3.95e-08 | 2.04e-06 | 2.3 |
| turquoise | GO:0007006 | mitochondrial membrane organization | 4e-08 | 2.06e-06 | 2.25 |
| turquoise | GO:0071806 | protein transmembrane transport | 4.1399999999999994e-08 | 2.13e-06 | 2.7 |
| turquoise | GO:0034212 | peptide N-acetyltransferase activity | 3.38e-08 | 2.14e-06 | 2.43 |
| turquoise | GO:0019783 | ubiquitin-like protein-specific protease activity | 3.46e-08 | 2.16e-06 | 2.13 |
| turquoise | GO:0030135 | coated vesicle | 1.26e-07 | 2.16e-06 | 1.97 |
| turquoise | GO:0044665 | MLL1/2 complex | 1.3900000000000001e-07 | 2.3e-06 | 3.2 |
| turquoise | GO:0071339 | MLL1 complex | 1.3900000000000001e-07 | 2.32e-06 | 3.2 |
| turquoise | GO:1903293 | phosphatase complex | 1.3800000000000002e-07 | 2.33e-06 | 2.75 |
| turquoise | GO:0030833 | regulation of actin filament polymerization | 4.61e-08 | 2.35e-06 | 1.92 |
| turquoise | GO:0018105 | peptidyl-serine phosphorylation | 4.5999999999999995e-08 | 2.35e-06 | 1.89 |
| turquoise | GO:0008287 | protein serine/threonine phosphatase complex | 1.3800000000000002e-07 | 2.35e-06 | 2.75 |
| turquoise | GO:0140030 | modification-dependent protein binding | 3.8900000000000004e-08 | 2.36e-06 | 1.99 |
| turquoise | GO:0036459 | thiol-dependent ubiquitinyl hydrolase activity | 3.83e-08 | 2.36e-06 | 2.19 |
| turquoise | GO:0004843 | thiol-dependent ubiquitin-specific protease activity | 4e-08 | 2.4e-06 | 2.25 |
| turquoise | GO:0007030 | Golgi organization | 4.75e-08 | 2.41e-06 | 2.1 |
| turquoise | GO:0017038 | protein import | 4.88e-08 | 2.47e-06 | 2.02 |

| | | | | | |
|---|---|---|---|---|---|
| turquoise | GO:0006470 | protein dephosphorylation | 5.0700000000000004e-08 | 2.56e-06 | 1.9 |
| turquoise | GO:0010608 | posttranscriptional regulation of gene expression | 5.24e-08 | 2.63e-06 | 1.56 |
| turquoise | GO:0006605 | protein targeting | 5.59e-08 | 2.8e-06 | 1.9 |
| turquoise | GO:0032006 | regulation of TOR signaling | 5.9299999999999995e-08 | 2.96e-06 | 2.2 |
| turquoise | GO:0016311 | dephosphorylation | 6.63e-08 | 3.3e-06 | 1.78 |
| turquoise | GO:0101005 | ubiquitinyl hydrolase activity | 5.62e-08 | 3.33e-06 | 2.16 |
| turquoise | GO:0031300 | intrinsic component of organelle membrane | 2.1100000000000002e-07 | 3.44e-06 | 1.8 |
| turquoise | GO:0007010 | cytoskeleton organization | 6.96e-08 | 3.46e-06 | 1.37 |
| turquoise | GO:0045732 | positive regulation of protein catabolic process | 7.51e-08 | 3.72e-06 | 1.77 |
| turquoise | GO:0009968 | negative regulation of signal transduction | 8.130000000000001e-08 | 4.01e-06 | 1.3 |
| turquoise | GO:0046755 | viral budding | 8.4e-08 | 4.13e-06 | 4.36 |
| turquoise | GO:0031396 | regulation of protein ubiquitination | 9.13e-08 | 4.48e-06 | 1.81 |
| turquoise | GO:1902911 | protein kinase complex | 2.77e-07 | 4.5e-06 | 2.02 |
| turquoise | GO:0016050 | vesicle organization | 1.06e-07 | 5.19e-06 | 1.78 |
| turquoise | GO:0042391 | regulation of membrane potential | 1.0699999999999999e-07 | 5.19e-06 | 1.51 |
| turquoise | GO:0003713 | transcription coactivator activity | 8.880000000000001e-08 | 5.19e-06 | 1.67 |
| turquoise | GO:0032281 | AMPA glutamate receptor complex | 3.24e-07 | 5.2e-06 | 3.09 |
| turquoise | GO:0008306 | associative learning | 1.1800000000000001e-07 | 5.75e-06 | 2.12 |
| turquoise | GO:0005761 | mitochondrial ribosome | 3.9e-07 | 6.17e-06 | 3.54 |
| turquoise | GO:1903507 | negative regulation of nucleic acid-templated transcription | 1.28e-07 | 6.19e-06 | 1.3 |
| turquoise | GO:0000313 | organellar ribosome | 3.9e-07 | 6.22e-06 | 3.54 |
| turquoise | GO:1902554 | serine/threonine protein kinase complex | 4.0399999999999996e-07 | 6.33e-06 | 2.1 |
| turquoise | GO:0001558 | regulation of cell growth | 1.37e-07 | 6.63e-06 | 1.51 |
| turquoise | GO:0099175 | regulation of postsynapse organization | 1.3900000000000001e-07 | 6.7e-06 | 2.3 |
| turquoise | GO:0044325 | ion channel binding | 1.1599999999999999e-07 | 6.7e-06 | 1.96 |
| turquoise | GO:1902679 | negative regulation of RNA biosynthetic process | 1.4e-07 | 6.72e-06 | 1.3 |
| turquoise | GO:0045892 | negative regulation of transcription, DNA-templated | 1.42e-07 | 6.8e-06 | 1.3 |
| turquoise | GO:0016363 | nuclear matrix | 4.42e-07 | 6.88e-06 | 2.14 |
| turquoise | GO:0042393 | histone binding | 1.2699999999999999e-07 | 7.16e-06 | 1.82 |
| turquoise | GO:0047485 | protein N-terminus binding | 1.26e-07 | 7.19e-06 | 1.97 |
| turquoise | GO:2001257 | regulation of cation channel activity | 1.57e-07 | 7.4999999999999999e-06 | 1.88 |
| turquoise | GO:0008064 | regulation of actin polymerization or depolymerization | 1.58e-07 | 7.53e-06 | 1.83 |
| turquoise | GO:0045862 | positive regulation of proteolysis | 1.73e-07 | 8.18e-06 | 1.59 |
| turquoise | GO:0006473 | protein acetylation | 1.73e-07 | 8.19e-06 | 2.01 |
| turquoise | GO:0007613 | memory | 1.7600000000000001e-07 | 8.27e-06 | 1.92 |
| turquoise | GO:0018393 | internal peptidyl-lysine acetylation | 1.8600000000000002e-07 | 8.74e-06 | 2.13 |
| turquoise | GO:0045944 | positive regulation of transcription by RNA polymerase II | 1.8899999999999999e-07 | 8.83e-06 | 1.29 |
| turquoise | GO:1901799 | negative regulation of proteasomal protein catabolic process | 2.01e-07 | 9.38e-06 | 2.6 |
| turquoise | GO:0030832 | regulation of actin filament length | 2.05e-07 | 9.52e-06 | 1.82 |
| turquoise | GO:0019866 | organelle inner membrane | 6.26e-07 | 9.67e-06 | 1.51 |
| turquoise | GO:0005743 | mitochondrial inner membrane | 6.75e-07 | 1.0300000000000001e-05 | 1.53 |
| turquoise | GO:0042326 | negative regulation of phosphorylation | 2.39e-07 | 1.11e-05 | 1.49 |
| turquoise | GO:0035254 | glutamate receptor binding | 2.0199999999999998e-07 | 1.12e-05 | 2.36 |
| turquoise | GO:0007007 | inner mitochondrial membrane organization | 2.47e-07 | 1.1400000000000001e-05 | 3.35 |
| turquoise | GO:0015931 | nucleobase-containing compound transport | 2.55e-07 | 1.17e-05 | 1.83 |
| turquoise | GO:0030427 | site of polarized growth | 7.78e-07 | 1.18e-05 | 1.74 |
| turquoise | GO:1901214 | regulation of neuron death | 2.62e-07 | 1.21e-05 | 1.55 |
| turquoise | GO:0031058 | positive regulation of histone modification | 2.68e-07 | 1.23e-05 | 2.11 |
| turquoise | GO:0007264 | small GTPase mediated signal transduction | 2.75e-07 | 1.25e-05 | 1.64 |
| turquoise | GO:0008213 | protein alkylation | 2.77e-07 | 1.26e-05 | 2.02 |
| turquoise | GO:0006479 | protein methylation | 2.77e-07 | 1.26e-05 | 2.02 |
| turquoise | GO:0051338 | regulation of transferase activity | 2.8e-07 | 1.27e-05 | 1.35 |
| turquoise | GO:0050657 | nucleic acid transport | 2.8300000000000003e-07 | 1.27e-05 | 1.92 |
| turquoise | GO:0050658 | RNA transport | 2.8300000000000003e-07 | 1.28e-05 | 1.92 |
| turquoise | GO:0008022 | protein C-terminus binding | 2.4e-07 | 1.31e-05 | 1.68 |
| turquoise | GO:0030426 | growth cone | 8.77e-07 | 1.32e-05 | 1.74 |
| turquoise | GO:0051099 | positive regulation of binding | 3.06e-07 | 1.3699999999999998e-05 | 1.74 |
| turquoise | GO:0034702 | ion channel complex | 9.31e-07 | 1.39e-05 | 1.58 |
| turquoise | GO:0018210 | peptidyl-threonine modification | 3.19e-07 | 1.4300000000000002e-05 | 2.21 |
| turquoise | GO:1903312 | negative regulation of mRNA metabolic process | 3.25e-07 | 1.4499999999999998e-05 | 2.26 |
| turquoise | GO:0060076 | excitatory synapse | 1.05e-06 | 1.56e-05 | 2.65 |
| turquoise | GO:0098984 | neuron to neuron synapse | 1.1e-06 | 1.6100000000000002e-05 | 3.37 |
| turquoise | GO:0016407 | acetyltransferase activity | 2.98e-07 | 1.6100000000000002e-05 | 2.03 |
| turquoise | GO:1903322 | positive regulation of protein modification by small protein conjugation | 3.7299999999999997e-07 | 1.66e-05 | 1.94 |
| turquoise | GO:0043175 | RNA polymerase core enzyme binding | 3.37e-07 | 1.8100000000000003e-05 | 2.92 |
| turquoise | GO:0043393 | regulation of protein binding | 4.11e-07 | 1.82e-05 | 1.67 |
| turquoise | GO:0032273 | positive regulation of protein polymerization | 4.1299999999999995e-07 | 1.82e-05 | 2.01 |
| turquoise | GO:0018209 | peptidyl-serine modification | 4.3600000000000004e-07 | 1.92e-05 | 1.77 |
| turquoise | GO:0044430 | cytoskeletal part | 1.35e-06 | 1.9600000000000002e-05 | 1.24 |
| turquoise | GO:0044419 | interspecies interaction between organisms | 4.5899999999999997e-07 | 2.02e-05 | 1.7 |
| turquoise | GO:0031965 | nuclear membrane | 1.41e-06 | 2.0399999999999998e-05 | 1.65 |
| turquoise | GO:0005085 | guanyl-nucleotide exchange factor activity | 3.88e-07 | 2.05e-05 | 1.75 |
| turquoise | GO:0048583 | regulation of response to stimulus | 4.77e-07 | 2.09e-05 | 1.15 |
| turquoise | GO:0050806 | positive regulation of synaptic transmission | 4.91e-07 | 2.14e-05 | 1.75 |
| turquoise | GO:0051236 | establishment of RNA localization | 5e-07 | 2.18e-05 | 1.89 |
| turquoise | GO:0033157 | regulation of intracellular protein transport | 5.19e-07 | 2.25e-05 | 1.7 |
| turquoise | GO:0051427 | hormone receptor binding | 4.32e-07 | 2.26e-05 | 1.78 |
| turquoise | GO:0032412 | regulation of ion transmembrane transporter activity | 5.27e-07 | 2.2800000000000002e-05 | 1.67 |
| turquoise | GO:1900449 | regulation of glutamate receptor signaling pathway | 5.3e-07 | 2.2899999999999998e-05 | 2.56 |
| turquoise | GO:0000124 | SAGA complex | 1.6e-06 | 2.3e-05 | 4.29 |
| turquoise | GO:1990351 | transporter complex | 1.62e-06 | 2.3100000000000002e-05 | 1.54 |
| turquoise | GO:0018394 | peptidyl-lysine acetylation | 5.420000000000001e-07 | 2.33e-05 | 2.07 |
| turquoise | GO:0006475 | internal protein amino acid acetylation | 5.420000000000001e-07 | 2.34e-05 | 2.07 |
| turquoise | GO:1990778 | protein localization to cell periphery | 5.5e-07 | 2.35e-05 | 1.74 |
| turquoise | GO:0031252 | cell leading edge | 1.67e-06 | 2.37e-05 | 2.36 |
| turquoise | GO:0042802 | identical protein binding | 4.7299999999999996e-07 | 2.44e-05 | 1.22 |
| turquoise | GO:0000118 | histone deacetylase complex | 1.74e-06 | 2.45e-05 | 2.29 |
| turquoise | GO:0120111 | neuron projection cytoplasm | 1.86e-06 | 2.59e-05 | 2.59 |
| turquoise | GO:0051493 | regulation of cytoskeleton organization | 6.16e-07 | 2.6300000000000002e-05 | 1.43 |
| turquoise | GO:0008180 | COP9 signalosome | 1.94e-06 | 2.68e-05 | 2.78 |
| turquoise | GO:1905269 | positive regulation of chromatin organization | 6.620000000000001e-07 | 2.8199999999999998e-05 | 2.02 |
| turquoise | GO:1901698 | response to nitrogen compound | 6.65e-07 | 2.8300000000000003e-05 | 1.37 |
| turquoise | GO:0016571 | histone methylation | 6.94e-07 | 2.9299999999999997e-05 | 2.16 |
| turquoise | GO:0010639 | negative regulation of organelle organization | 6.92e-07 | 2.9299999999999997e-05 | 1.51 |
| turquoise | GO:0016482 | cytosolic transport | 7.159999999999999e-07 | 3.01e-05 | 1.96 |
| turquoise | GO:0070063 | RNA polymerase binding | 5.92e-07 | 3.0299999999999998e-05 | 2.5 |
| turquoise | GO:0009314 | response to radiation | 7.3e-07 | 3.07e-05 | 1.5 |
| turquoise | GO:0010821 | regulation of mitochondrion organization | 7.37e-07 | 3.09e-05 | 1.92 |
| turquoise | GO:0032550 | purine ribonucleoside binding | 6.75e-07 | 3.41e-05 | 1.53 |
| turquoise | GO:0001883 | purine nucleoside binding | 6.970000000000001e-07 | 3.48e-05 | 1.52 |
| turquoise | GO:0009628 | response to abiotic stimulus | 8.33e-07 | 3.48e-05 | 1.32 |
| turquoise | GO:0000993 | RNA polymerase II complex binding | 7.070000000000001e-07 | 3.4899999999999995e-05 | 2.99 |
| turquoise | GO:0061733 | peptide-lysine-N-acetyltransferase activity | 7.1799999999999999e-07 | 3.5100000000000006e-05 | 2.36 |
| turquoise | GO:1901407 | regulation of phosphorylation of RNA polymerase II C-terminal domain | 8.5e-07 | 3.54e-05 | 4.72 |

| | | | | | |
|---|---|---|---|---|---|
| turquoise | GO:0002039 | p53 binding | 7.5e-07 | 3.63e-05 | 2.29 |
| turquoise | GO:0050793 | regulation of developmental process | 9.54e-07 | 3.96e-05 | 1.18 |
| turquoise | GO:2000785 | regulation of autophagosome assembly | 9.95e-07 | 4.1200000000000005e-05 | 2.86 |
| turquoise | GO:0007265 | Ras protein signal transduction | 1.02e-06 | 4.210000000000001e-05 | 1.69 |
| turquoise | GO:0071007 | U2-type catalytic step 2 spliceosome | 3.11e-06 | 4.270000000000001e-05 | 2.97 |
| turquoise | GO:0098693 | regulation of synaptic vesicle cycle | 1.05e-06 | 4.3200000000000007e-05 | 2.4 |
| turquoise | GO:0001650 | fibrillar center | 3.28e-06 | 4.48e-05 | 1.85 |
| turquoise | GO:0032947 | protein-containing complex scaffold activity | 9.62e-07 | 4.6e-05 | 2.11 |
| turquoise | GO:0019207 | kinase regulator activity | 9.75e-07 | 4.6200000000000005e-05 | 1.73 |
| turquoise | GO:0017048 | Rho GTPase binding | 9.9e-07 | 4.6399999999999996e-05 | 1.77 |
| turquoise | GO:0005798 | Golgi-associated vesicle | 3.46e-06 | 4.6899999999999995e-05 | 2.11 |
| turquoise | GO:0048168 | regulation of neuronal synaptic plasticity | 1.14e-06 | 4.7e-05 | 2.29 |
| turquoise | GO:0016791 | phosphatase activity | 1.05e-06 | 4.85e-05 | 1.63 |
| turquoise | GO:0043414 | macromolecule methylation | 1.19e-06 | 4.8700000000000005e-05 | 1.7 |
| turquoise | GO:0016592 | mediator complex | 3.62e-06 | 4.8700000000000005e-05 | 2.7 |
| turquoise | GO:0043549 | regulation of kinase activity | 1.22e-06 | 4.99e-05 | 1.35 |
| turquoise | GO:0005686 | U2 snRNP | 3.73e-06 | 4.99e-05 | 3.31 |
| turquoise | GO:0120036 | plasma membrane bounded cell projection organization | 1.22e-06 | 5e-05 | 1.38 |
| turquoise | GO:0004402 | histone acetyltransferase activity | 1.1e-06 | 5.03e-05 | 2.36 |
| turquoise | GO:0022898 | regulation of transmembrane transporter activity | 1.26e-06 | 5.13e-05 | 1.63 |
| turquoise | GO:0048585 | negative regulation of response to stimulus | 1.28e-06 | 5.18e-05 | 1.23 |
| turquoise | GO:0016573 | histone acetylation | 1.3e-06 | 5.2000000000000004e-05 | 2.07 |
| turquoise | GO:0019220 | regulation of phosphate metabolic process | 1.3e-06 | 5.21e-05 | 1.22 |
| turquoise | GO:0044088 | regulation of vacuole organization | 1.3e-06 | 5.2300000000000004e-05 | 2.75 |
| turquoise | GO:0034765 | regulation of ion transmembrane transport | 1.29e-06 | 5.2300000000000004e-05 | 1.45 |
| turquoise | GO:0001882 | nucleoside binding | 1.18e-06 | 5.3500000000000006e-05 | 1.5 |
| turquoise | GO:0006468 | protein phosphorylation | 1.34e-06 | 5.3700000000000004e-05 | 1.35 |
| turquoise | GO:0007041 | lysosomal transport | 1.37e-06 | 5.479999999999999e-05 | 2.09 |
| turquoise | GO:0051174 | regulation of phosphorus metabolic process | 1.39e-06 | 5.5299999999999996e-05 | 1.22 |
| turquoise | GO:0006515 | protein quality control for misfolded or incompletely synthesized protein | 1.43e-06 | 5.6999999999999996e-05 | 3.09 |
| turquoise | GO:0000387 | spliceosomal snRNP assembly | 1.46e-06 | 5.79e-05 | 2.89 |
| turquoise | GO:0031301 | integral component of organelle membrane | 4.53e-06 | 5.9699999999999994e-05 | 1.73 |
| turquoise | GO:0098590 | plasma membrane region | 4.53e-06 | 6.01e-05 | 1.27 |
| turquoise | GO:0042594 | response to starvation | 1.57e-06 | 6.19e-05 | 1.79 |
| turquoise | GO:1903363 | negative regulation of cellular protein catabolic process | 1.59e-06 | 6.25e-05 | 2.15 |
| turquoise | GO:0072655 | establishment of protein localization to mitochondrion | 1.6e-06 | 6.29e-05 | 2.41 |
| turquoise | GO:0032279 | asymmetric synapse | 4.89e-06 | 6.4e-05 | 3.41 |
| turquoise | GO:0032549 | ribonucleoside binding | 1.48e-06 | 6.64e-05 | 1.51 |
| turquoise | GO:0070585 | protein localization to mitochondrion | 1.72e-06 | 6.73e-05 | 2.32 |
| turquoise | GO:0034504 | protein localization to nucleus | 1.72e-06 | 6.74e-05 | 1.87 |
| turquoise | GO:0018024 | histone-lysine N-methyltransferase activity | 1.59e-06 | 7.08e-05 | 2.66 |
| turquoise | GO:0032784 | regulation of DNA-templated transcription, elongation | 1.86e-06 | 7.25e-05 | 2.59 |
| turquoise | GO:0005813 | centrosome | 5.59e-06 | 7.269999999999999e-05 | 1.4 |
| turquoise | GO:0034470 | ncRNA processing | 1.91e-06 | 7.41e-05 | 1.54 |
| turquoise | GO:0035770 | ribonucleoprotein granule | 5.74e-06 | 7.41e-05 | 1.65 |
| turquoise | GO:1903543 | positive regulation of exosomal secretion | 1.91e-06 | 7.42e-05 | 3.78 |
| turquoise | GO:0030119 | AP-type membrane coat adaptor complex | 5.98e-06 | 7.670000000000001e-05 | 2.36 |
| turquoise | GO:1902495 | transmembrane transporter complex | 6.02e-06 | 7.670000000000001e-05 | 1.52 |
| turquoise | GO:1903649 | regulation of cytoplasmic transport | 2.01e-06 | 7.77e-05 | 3.15 |
| turquoise | GO:0005682 | U5 snRNP | 6.15e-06 | 7.780000000000001e-05 | 3.54 |
| turquoise | GO:0070461 | SAGA-type complex | 6.39e-06 | 8.04e-05 | 3.08 |
| turquoise | GO:0032409 | regulation of transporter activity | 2.09e-06 | 8.06e-05 | 1.61 |
| turquoise | GO:0042177 | negative regulation of protein catabolic process | 2.11e-06 | 8.1e-05 | 1.89 |
| turquoise | GO:0030120 | vesicle coat | 6.52e-06 | 8.14e-05 | 2.62 |
| turquoise | GO:0034709 | methylosome | 6.88e-06 | 8.540000000000002e-05 | 4.25 |
| turquoise | GO:0099601 | regulation of neurotransmitter receptor activity | 2.29e-06 | 8.809999999999999e-05 | 2.46 |
| turquoise | GO:0060548 | negative regulation of cell death | 2.36e-06 | 9.04e-05 | 1.28 |
| turquoise | GO:0008080 | N-acetyltransferase activity | 2.06e-06 | 9.1e-05 | 2.08 |
| turquoise | GO:0006890 | retrograde vesicle-mediated transport, Golgi to ER | 2.4e-06 | 9.159999999999999e-05 | 2.68 |
| turquoise | GO:0010941 | regulation of cell death | 2.42e-06 | 9.209999999999999e-05 | 1.22 |
| turquoise | GO:0043085 | positive regulation of catalytic activity | 2.44e-06 | 9.259999999999999e-05 | 1.28 |
| turquoise | GO:0006352 | DNA-templated transcription, initiation | 2.45e-06 | 9.279999999999999e-05 | 2.41 |
| turquoise | GO:0005525 | GTP binding | 2.16e-06 | 9.42e-05 | 1.51 |
| turquoise | GO:1901983 | regulation of protein acetylation | 2.49e-06 | 9.429999999999999e-05 | 2.16 |
| turquoise | GO:0051592 | response to calcium ion | 2.51e-06 | 9.46e-05 | 1.87 |
| turquoise | GO:0030008 | TRAPP complex | 7.74e-06 | 9.48e-05 | 3.94 |
| turquoise | GO:0010508 | positive regulation of autophagy | 2.52e-06 | 9.49e-05 | 1.95 |
| turquoise | GO:0030123 | AP-3 adaptor complex | 7.74e-06 | 9.54e-05 | 3.94 |
| turquoise | GO:0033555 | multicellular organismal response to stress | 2.57e-06 | 9.640000000000001e-05 | 2.19 |
| turquoise | GO:0008542 | visual learning | 2.62e-06 | 9.83e-05 | 2.22 |
| turquoise | GO:0016239 | positive regulation of macroautophagy | 2.66e-06 | 9.91e-05 | 2.28 |
| turquoise | GO:0034968 | histone lysine methylation | 2.66e-06 | 9.940000000000002e-05 | 2.28 |
| turquoise | GO:1901566 | organonitrogen compound biosynthetic process | 2.72e-06 | 0.000101 | 1.32 |
| turquoise | GO:0001933 | negative regulation of protein phosphorylation | 2.78e-06 | 0.000103 | 1.47 |
| turquoise | GO:0006366 | transcription by RNA polymerase II | 2.86e-06 | 0.000105 | 1.77 |
| turquoise | GO:0070972 | protein localization to endoplasmic reticulum | 2.82e-06 | 0.000105 | 2.6 |
| turquoise | GO:0045859 | regulation of protein kinase activity | 2.88e-06 | 0.000106 | 1.36 |
| turquoise | GO:0050686 | negative regulation of mRNA processing | 2.88e-06 | 0.000106 | 2.8 |
| turquoise | GO:0042054 | histone methyltransferase activity | 2.45e-06 | 0.000106 | 2.41 |
| turquoise | GO:0032838 | plasma membrane bounded cell projection cytoplasm | 8.77e-06 | 0.00010700000000000001 | 2.41 |
| turquoise | GO:0017137 | Rab GTPase binding | 2.54e-06 | 0.00010800000000000001 | 1.77 |
| turquoise | GO:0004721 | phosphoprotein phosphatase activity | 2.52e-06 | 0.00010800000000000001 | 1.73 |
| turquoise | GO:0019003 | GDP binding | 2.66e-06 | 0.00011200000000000001 | 2.28 |
| turquoise | GO:0042578 | phosphoric ester hydrolase activity | 2.7000000000000004e-06 | 0.000113 | 1.51 |
| turquoise | GO:0019789 | SUMO transferase activity | 2.77e-06 | 0.000113 | 3.22 |
| turquoise | GO:0031503 | protein-containing complex localization | 3.08e-06 | 0.000113 | 1.88 |
| turquoise | GO:0008092 | cytoskeletal protein binding | 2.75e-06 | 0.00011399999999999999 | 1.29 |
| turquoise | GO:0006376 | mRNA splice site selection | 3.11e-06 | 0.00011399999999999999 | 2.97 |
| turquoise | GO:0008219 | cell death | 3.2e-06 | 0.000117 | 1.34 |
| turquoise | GO:0010243 | response to organonitrogen compound | 3.34e-06 | 0.000122 | 1.36 |
| turquoise | GO:0031175 | neuron projection development | 3.48e-06 | 0.000126 | 1.54 |
| turquoise | GO:0035556 | intracellular signal transduction | 3.73e-06 | 0.000135 | 1.24 |
| turquoise | GO:0061077 | chaperone-mediated protein folding | 3.74e-06 | 0.000135 | 2.41 |
| turquoise | GO:0031060 | regulation of histone methylation | 3.77e-06 | 0.000136 | 2.16 |
| turquoise | GO:1903051 | negative regulation of proteolysis involved in cellular protein catabolic p | 3.9e-06 | 0.00014 | 2.19 |
| turquoise | GO:0001099 | basal RNA polymerase II transcription machinery binding | 3.5000000000000004e-06 | 0.00014 | 2.46 |
| turquoise | GO:0008170 | N-methyltransferase activity | 3.46e-06 | 0.00014 | 2.11 |
| turquoise | GO:0036464 | cytoplasmic ribonucleoprotein granule | 1.1599999999999999e-05 | 0.00014 | 1.64 |
| turquoise | GO:0001098 | basal transcription machinery binding | 3.5000000000000004e-06 | 0.00014099999999999998 | 2.46 |
| turquoise | GO:1901409 | positive regulation of phosphorylation of RNA polymerase II C-terminal | 4.02e-06 | 0.000144 | 4.72 |
| turquoise | GO:1905475 | regulation of protein localization to membrane | 4.08e-06 | 0.000146 | 1.76 |
| turquoise | GO:0005741 | mitochondrial outer membrane | 1.27e-05 | 0.000153 | 1.71 |
| turquoise | GO:0030307 | positive regulation of cell growth | 4.32e-06 | 0.000154 | 1.7 |
| turquoise | GO:0009267 | cellular response to starvation | 4.44e-06 | 0.00015800000000000002 | 1.81 |

| | | | | | |
|---|---|---|---|---|---|
| turquoise | GO:0033673 | negative regulation of kinase activity | 4.46e-06 | 0.00015900000000000002 | 1.64 |
| turquoise | GO:0046540 | U4/U6 x U5 tri-snRNP complex | 1.3699999999999998e-05 | 0.00016299999999999998 | 2.95 |
| turquoise | GO:0097526 | spliceosomal tri-snRNP complex | 1.3699999999999998e-05 | 0.000164 | 2.95 |
| turquoise | GO:0008023 | transcription elongation factor complex | 1.4000000000000001e-05 | 0.000165 | 2.36 |
| turquoise | GO:0019001 | guanyl nucleotide binding | 4.26e-06 | 0.000167 | 1.47 |
| turquoise | GO:0055029 | nuclear DNA-directed RNA polymerase complex | 1.4400000000000001e-05 | 0.000168 | 2.31 |
| turquoise | GO:0032561 | guanyl ribonucleotide binding | 4.26e-06 | 0.000168 | 1.47 |
| turquoise | GO:0051087 | chaperone binding | 4.44e-06 | 0.000172 | 1.97 |
| turquoise | GO:0009416 | response to light stimulus | 4.91e-06 | 0.000174 | 1.55 |
| turquoise | GO:0030838 | positive regulation of actin filament polymerization | 4.95e-06 | 0.000175 | 2.09 |
| turquoise | GO:0007033 | vacuole organization | 5.01e-06 | 0.000177 | 1.93 |
| turquoise | GO:0044092 | negative regulation of molecular function | 5.05e-06 | 0.000178 | 1.28 |
| turquoise | GO:0001222 | transcription corepressor binding | 4.89e-06 | 0.000188 | 3.41 |
| turquoise | GO:0040008 | regulation of growth | 5.34e-06 | 0.000188 | 1.33 |
| turquoise | GO:0008276 | protein methyltransferase activity | 5.22e-06 | 0.000199 | 2.11 |
| turquoise | GO:0035303 | regulation of dephosphorylation | 5.67e-06 | 0.000199 | 1.72 |
| turquoise | GO:0140110 | transcription regulator activity | 5.38e-06 | 0.00020299999999999997 | 1.24 |
| turquoise | GO:0046983 | protein dimerization activity | 5.41e-06 | 0.00020299999999999997 | 1.23 |
| turquoise | GO:0098815 | modulation of excitatory postsynaptic potential | 5.98e-06 | 0.00020800000000000001 | 2.36 |
| turquoise | GO:0000122 | negative regulation of transcription by RNA polymerase II | 5.98e-06 | 0.00020899999999999998 | 1.3 |
| turquoise | GO:0043967 | histone H4 acetylation | 5.98e-06 | 0.00020899999999999998 | 2.36 |
| turquoise | GO:2000756 | regulation of peptidyl-lysine acetylation | 6.07e-06 | 0.00021 | 2.21 |
| turquoise | GO:0022411 | cellular component disassembly | 6.06e-06 | 0.00021099999999999998 | 1.69 |
| turquoise | GO:0034314 | Arp2/3 complex-mediated actin nucleation | 6.15e-06 | 0.000212 | 3.54 |
| turquoise | GO:1903541 | regulation of exosomal secretion | 6.15e-06 | 0.000212 | 3.54 |
| turquoise | GO:1903829 | positive regulation of cellular protein localization | 6.21e-06 | 0.00021400000000000002 | 1.53 |
| turquoise | GO:0019903 | protein phosphatase binding | 5.7999999999999995e-06 | 0.00021600000000000002 | 1.76 |
| turquoise | GO:0031365 | N-terminal protein amino acid modification | 6.39e-06 | 0.00021899999999999998 | 3.08 |
| turquoise | GO:0045292 | mRNA cis splicing, via spliceosome | 6.39e-06 | 0.00022 | 3.08 |
| turquoise | GO:1902903 | regulation of supramolecular fiber organization | 6.49e-06 | 0.00022200000000000003 | 1.48 |
| turquoise | GO:0042254 | ribosome biogenesis | 6.54e-06 | 0.000223 | 2.04 |
| turquoise | GO:0000407 | phagophore assembly site | 1.99e-05 | 0.00023199999999999997 | 3.01 |
| turquoise | GO:0017157 | regulation of exocytosis | 6.86e-06 | 0.000233 | 1.69 |
| turquoise | GO:0043162 | ubiquitin-dependent protein catabolic process via the multivesicular bod | 6.88e-06 | 0.000234 | 4.25 |
| turquoise | GO:0072659 | protein localization to plasma membrane | 7.07e-06 | 0.000239 | 1.71 |
| turquoise | GO:0005516 | calmodulin binding | 6.62e-06 | 0.000244 | 1.68 |
| turquoise | GO:1903358 | regulation of Golgi organization | 7.26e-06 | 0.000245 | 3.71 |
| turquoise | GO:1901836 | regulation of transcription of nucleolar large rRNA by RNA polymerase I | 7.26e-06 | 0.000245 | 3.71 |
| turquoise | GO:0051028 | mRNA transport | 7.38e-06 | 0.000248 | 1.92 |
| turquoise | GO:0000428 | DNA-directed RNA polymerase complex | 2.2100000000000002e-05 | 0.000251 | 2.27 |
| turquoise | GO:0000307 | cyclin-dependent protein kinase holoenzyme complex | 2.2100000000000002e-05 | 0.00025299999999999997 | 2.27 |
| turquoise | GO:0005665 | RNA polymerase II, core complex | 2.2e-05 | 0.00025299999999999997 | 3.46 |
| turquoise | GO:0099055 | integral component of postsynaptic membrane | 2.2e-05 | 0.000254 | 3.46 |
| turquoise | GO:0042800 | histone methyltransferase activity (H3-K4 specific) | 7.26e-06 | 0.000265 | 3.71 |
| turquoise | GO:0002209 | behavioral defense response | 8.2e-06 | 0.000275 | 2.47 |
| turquoise | GO:0072599 | establishment of protein localization to endoplasmic reticulum | 8.19e-06 | 0.000275 | 2.74 |
| turquoise | GO:0007632 | visual behavior | 8.29e-06 | 0.000277 | 2.12 |
| turquoise | GO:1990928 | response to amino acid starvation | 8.77e-06 | 0.000293 | 2.41 |
| turquoise | GO:0098772 | molecular function regulator | 8.34e-06 | 0.000303 | 1.21 |
| turquoise | GO:0035258 | steroid hormone receptor binding | 8.43e-06 | 0.000303 | 2.0 |
| turquoise | GO:0043603 | cellular amide metabolic process | 9.26e-06 | 0.000308 | 1.35 |
| turquoise | GO:0035601 | protein deacylation | 9.4e-06 | 0.00031 | 2.24 |
| turquoise | GO:0030100 | regulation of endocytosis | 9.36e-06 | 0.00031099999999999997 | 1.56 |
| turquoise | GO:0098732 | macromolecule deacylation | 9.4e-06 | 0.00031099999999999997 | 2.24 |
| turquoise | GO:0048813 | dendrite morphogenesis | 9.38e-06 | 0.00031099999999999997 | 2.32 |
| turquoise | GO:0051052 | regulation of DNA metabolic process | 9.46e-06 | 0.000312 | 1.43 |
| turquoise | GO:2001234 | negative regulation of apoptotic signaling pathway | 9.49e-06 | 0.000312 | 1.6 |
| turquoise | GO:0006469 | negative regulation of protein kinase activity | 9.51e-06 | 0.000312 | 1.65 |
| turquoise | GO:0005635 | nuclear envelope | 2.84e-05 | 0.00032 | 1.69 |
| turquoise | GO:0031398 | positive regulation of protein ubiquitination | 9.83e-06 | 0.000322 | 1.9 |
| turquoise | GO:0006458 | 'de novo' protein folding | 9.89e-06 | 0.000323 | 2.64 |
| turquoise | GO:0005885 | Arp2/3 protein complex | 3.07e-05 | 0.000342 | 3.86 |
| turquoise | GO:0008540 | proteasome regulatory particle, base subcomplex | 3.07e-05 | 0.000344 | 3.86 |
| turquoise | GO:0016410 | N-acyltransferase activity | 9.83e-06 | 0.00035099999999999997 | 1.9 |
| turquoise | GO:0006915 | apoptotic process | 1.0800000000000002e-05 | 0.00035299999999999996 | 1.34 |
| turquoise | GO:0098978 | glutamatergic synapse | 3.37e-05 | 0.00037400000000000004 | 2.31 |
| turquoise | GO:0051648 | vesicle localization | 1.15e-05 | 0.000375 | 1.8 |
| turquoise | GO:0031669 | cellular response to nutrient levels | 1.1900000000000001e-05 | 0.000384 | 1.68 |
| turquoise | GO:0012501 | programmed cell death | 1.2e-05 | 0.00039 | 1.33 |
| turquoise | GO:0051650 | establishment of vesicle localization | 1.24e-05 | 0.000401 | 1.84 |
| turquoise | GO:0017069 | snRNA binding | 1.1400000000000001e-05 | 0.00040199999999999996 | 2.55 |
| turquoise | GO:1905897 | regulation of response to endoplasmic reticulum stress | 1.25e-05 | 0.000404 | 2.11 |
| turquoise | GO:0006338 | chromatin remodeling | 1.29e-05 | 0.00041500000000000006 | 1.81 |
| turquoise | GO:0022603 | regulation of anatomical structure morphogenesis | 1.3300000000000001e-05 | 0.00042800000000000005 | 1.26 |
| turquoise | GO:0044798 | nuclear transcription factor complex | 3.97e-05 | 0.000436 | 1.57 |
| turquoise | GO:0099568 | cytoplasmic region | 3.97e-05 | 0.00043799999999999997 | 1.57 |
| turquoise | GO:0019887 | protein kinase regulator activity | 1.27e-05 | 0.00044699999999999997 | 1.71 |
| turquoise | GO:0007015 | actin filament organization | 1.47e-05 | 0.00047000000000000004 | 1.59 |
| turquoise | GO:0006367 | transcription initiation from RNA polymerase II promoter | 1.4999999999999999e-05 | 0.00047999999999999996 | 2.66 |
| turquoise | GO:0005622 | intracellular | 4.41e-05 | 0.000481 | 1.29 |
| turquoise | GO:0051100 | negative regulation of binding | 1.52e-05 | 0.00048499999999999997 | 1.68 |
| turquoise | GO:0055038 | recycling endosome membrane | 5.07e-05 | 0.00055 | 2.36 |
| turquoise | GO:0022406 | membrane docking | 1.7899999999999998e-05 | 0.00057 | 2.08 |
| turquoise | GO:0043087 | regulation of GTPase activity | 1.8e-05 | 0.00057 | 1.5 |
| turquoise | GO:0090575 | RNA polymerase II transcription factor complex | 5.379999999999999e-05 | 0.00058 | 1.63 |
| turquoise | GO:0048524 | positive regulation of viral process | 1.87e-05 | 0.000593 | 1.98 |
| turquoise | GO:0018022 | peptidyl-lysine methylation | 1.89e-05 | 0.000599 | 2.11 |
| turquoise | GO:0021957 | corticospinal tract morphogenesis | 1.9e-05 | 0.0006 | 4.72 |
| turquoise | GO:0031401 | positive regulation of protein modification process | 1.91e-05 | 0.000601 | 1.24 |
| turquoise | GO:0098936 | intrinsic component of postsynaptic membrane | 5.68e-05 | 0.000605 | 3.25 |
| turquoise | GO:0099699 | integral component of synaptic membrane | 5.68e-05 | 0.000609 | 3.25 |
| turquoise | GO:1902531 | regulation of intracellular signal transduction | 1.9699999999999998e-05 | 0.0006190000000000001 | 1.2 |
| turquoise | GO:0045597 | positive regulation of cell differentiation | 2e-05 | 0.000628 | 1.25 |
| turquoise | GO:0001662 | behavioral fear response | 2.0600000000000003e-05 | 0.0006450000000000001 | 2.42 |
| turquoise | GO:0030176 | integral component of endoplasmic reticulum membrane | 6.08e-05 | 0.0006450000000000001 | 1.81 |
| turquoise | GO:0001764 | neuron migration | 2.11e-05 | 0.00066 | 1.74 |
| turquoise | GO:0048278 | vesicle docking | 2.14e-05 | 0.000667 | 2.19 |
| turquoise | GO:1902117 | positive regulation of organelle assembly | 2.1899999999999997e-05 | 0.000682 | 2.02 |
| turquoise | GO:0005793 | endoplasmic reticulum-Golgi intermediate compartment | 6.52e-05 | 0.000688 | 2.08 |
| turquoise | GO:0070603 | SWI/SNF superfamily-type complex | 6.7e-05 | 0.0007019999999999999 | 1.97 |
| turquoise | GO:0030880 | RNA polymerase complex | 7.26e-05 | 0.000753 | 2.14 |
| turquoise | GO:0019867 | outer membrane | 7.36e-05 | 0.000755 | 1.6 |
| turquoise | GO:0032592 | integral component of mitochondrial membrane | 7.26e-05 | 0.0007570000000000001 | 2.14 |
| turquoise | GO:0031968 | organelle outer membrane | 7.36e-05 | 0.000759 | 1.6 |

| | | | | | |
|---|---|---|---|---|---|
| turquoise | GO:0005789 | endoplasmic reticulum membrane | 7.509999999999998e-05 | 0.000766 | 1.45 |
| turquoise | GO:0010977 | negative regulation of neuron projection development | 2.48e-05 | 0.000769 | 1.65 |
| turquoise | GO:0005815 | microtubule organizing center | 7.609999999999999e-05 | 0.000773 | 1.31 |
| turquoise | GO:0032502 | developmental process | 2.51e-05 | 0.000779 | 1.1 |
| turquoise | GO:0042147 | retrograde transport, endosome to Golgi | 2.5300000000000002e-05 | 0.000782 | 2.05 |
| turquoise | GO:0070013 | intracellular organelle lumen | 7.92e-05 | 0.000791 | 1.47 |
| turquoise | GO:0031967 | organelle envelope | 7.890000000000001e-05 | 0.000792 | 1.61 |
| turquoise | GO:0006839 | mitochondrial transport | 2.58e-05 | 0.000796 | 1.9 |
| turquoise | GO:0031975 | envelope | 7.890000000000001e-05 | 0.000797 | 1.61 |
| turquoise | GO:0048731 | system development | 2.6300000000000002e-05 | 0.000811 | 1.3 |
| turquoise | GO:0051084 | 'de novo' posttranslational protein folding | 2.65e-05 | 0.000813 | 2.58 |
| turquoise | GO:0034243 | regulation of transcription elongation from RNA polymerase II promoter | 2.65e-05 | 0.0008150000000000001 | 2.58 |
| turquoise | GO:0017148 | negative regulation of translation | 2.6700000000000002e-05 | 0.0008179999999999999 | 1.79 |
| turquoise | GO:0043248 | proteasome assembly | 2.71e-05 | 0.0008289999999999999 | 3.63 |
| turquoise | GO:2000311 | regulation of AMPA receptor activity | 2.75e-05 | 0.00084 | 2.83 |
| turquoise | GO:0031345 | negative regulation of cell projection organization | 2.78e-05 | 0.0008470000000000001 | 1.59 |
| turquoise | GO:0010507 | negative regulation of autophagy | 2.8600000000000004e-05 | 0.000868 | 2.1 |
| turquoise | GO:0034762 | regulation of transmembrane transport | 2.94e-05 | 0.0008880000000000001 | 1.35 |
| turquoise | GO:0039702 | viral budding via host ESCRT complex | 2.94e-05 | 0.000889 | 4.2 |
| turquoise | GO:0005884 | actin filament | 9.01e-05 | 0.000891 | 1.88 |
| turquoise | GO:0032486 | Rap protein signal transduction | 2.94e-05 | 0.000891 | 4.2 |
| turquoise | GO:0071204 | histone pre-mRNA 3'end processing complex | 8.99e-05 | 0.000893 | 4.72 |
| turquoise | GO:0042734 | presynaptic membrane | 9.09e-05 | 0.0008939999999999999 | 1.94 |
| turquoise | GO:0031668 | cellular response to extracellular stimulus | 2.9999999999999997e-05 | 0.0009019999999999999 | 1.6 |
| turquoise | GO:0010469 | regulation of signaling receptor activity | 2.9999999999999997e-05 | 0.0009029999999999999 | 1.8 |
| turquoise | GO:0034705 | potassium channel complex | 9.53e-05 | 0.000932 | 1.84 |
| turquoise | GO:1905037 | autophagosome organization | 3.15e-05 | 0.000942 | 2.15 |
| turquoise | GO:0051056 | regulation of small GTPase mediated signal transduction | 3.15e-05 | 0.000944 | 1.53 |
| turquoise | GO:0004659 | prenyltransferase activity | 2.71e-05 | 0.0009460000000000001 | 3.63 |
| turquoise | GO:0034045 | phagophore assembly site membrane | 9.95e-05 | 0.000968 | 3.54 |
| turquoise | GO:0051966 | regulation of synaptic transmission, glutamatergic | 3.28e-05 | 0.0009789999999999998 | 2.01 |
| purple | GO:0005764 | lysosome | 2.2899999999999997e-21 | 2.17e-18 | 20.0 |
| purple | GO:0000323 | lytic vacuole | 2.2899999999999997e-21 | 4.34e-18 | 20.0 |
| purple | GO:0005773 | vacuole | 2.99e-20 | 1.8899999999999996e-17 | 17.57 |
| purple | GO:0005615 | extracellular space | 1.39e-18 | 6.59e-16 | 7.82 |
| purple | GO:0044421 | extracellular region part | 1.44e-17 | 5.47e-15 | 6.68 |
| purple | GO:0005576 | extracellular region | 6.57e-13 | 2.0799999999999998e-10 | 5.84 |
| purple | GO:0009986 | cell surface | 2.65e-12 | 7.190000000000001e-10 | 9.84 |
| purple | GO:0006955 | immune response | 2.1099999999999997e-12 | 3.2200000000000004e-08 | 8.96 |
| purple | GO:0002376 | immune system process | 4.589999999999999e-11 | 3.5e-07 | 5.76 |
| purple | GO:0048584 | positive regulation of response to stimulus | 1.8599999999999998e-10 | 9.45e-07 | 4.43 |
| purple | GO:0002474 | antigen processing and presentation of peptide antigen via MHC class | 2.9399999999999997e-10 | 1.12e-06 | 127.66 |
| purple | GO:0006952 | defense response | 7.059999999999999e-10 | 2.15e-06 | 6.75 |
| purple | GO:0050896 | response to stimulus | 1.6899999999999997e-09 | 4.2999999999999995e-06 | 2.75 |
| purple | GO:0002684 | positive regulation of immune system process | 2.0600000000000003e-09 | 4.48e-06 | 6.89 |
| purple | GO:0002682 | regulation of immune system process | 7.5e-09 | 1.27e-05 | 5.28 |
| purple | GO:0048002 | antigen processing and presentation of peptide antigen | 8.63e-09 | 1.32e-05 | 39.62 |
| purple | GO:0045807 | positive regulation of endocytosis | 7.070000000000001e-09 | 1.3500000000000001e-05 | 19.51 |
| purple | GO:0045087 | innate immune response | 3.15e-08 | 4.36e-05 | 10.46 |
| purple | GO:0098552 | side of membrane | 1.87e-07 | 4.429999999999999e-05 | 8.65 |
| purple | GO:0048583 | regulation of response to stimulus | 7.5e-08 | 8.79e-05 | 2.88 |
| purple | GO:0002252 | immune effector process | 8.5e-08 | 9.259999999999999e-05 | 9.41 |
| purple | GO:0005515 | protein binding | 2.06e-08 | 9.279999999999999e-05 | 1.91 |
| purple | GO:0050778 | positive regulation of immune response | 7.480000000000001e-08 | 9.499999999999999e-05 | 8.16 |
| purple | GO:0030984 | kininogen binding | 6.71e-08 | 0.000101 | 287.24 |
| purple | GO:0019882 | antigen processing and presentation | 1.0199999999999999e-07 | 0.000103 | 26.41 |
| purple | GO:0009897 | external side of plasma membrane | 5.38e-07 | 0.00011399999999999999 | 9.09 |
| purple | GO:0044877 | protein-containing complex binding | 1.05e-07 | 0.000118 | 5.13 |
| purple | GO:0004197 | cysteine-type endopeptidase activity | 1.5199999999999998e-07 | 0.000136 | 24.71 |
| purple | GO:0042277 | peptide binding | 6.3e-08 | 0.00014199999999999998 | 11.72 |
| purple | GO:0050776 | regulation of immune response | 2.03e-07 | 0.000193 | 6.52 |
| purple | GO:0030100 | regulation of endocytosis | 2.23e-07 | 0.0002 | 12.51 |
| purple | GO:0033218 | amide binding | 3.34e-07 | 0.00025 | 9.63 |
| purple | GO:0006950 | response to stress | 3.33e-07 | 0.00028199999999999997 | 3.27 |
| purple | GO:0051239 | regulation of multicellular organismal process | 4.3100000000000003e-07 | 0.00032900000000000003 | 2.96 |
| purple | GO:0002478 | antigen processing and presentation of exogenous peptide antigen | 4.2299999999999996e-07 | 0.00033999999999999997 | 63.83 |
| purple | GO:0019885 | antigen processing and presentation of endogenous peptide antigen v | 5.839999999999999e-07 | 0.000424 | 164.13 |
| purple | GO:0042612 | MHC class I protein complex | 2.73e-06 | 0.000519 | 104.45 |
| purple | GO:0006956 | complement activation | 7.92e-07 | 0.000549 | 29.01 |
| purple | GO:0019884 | antigen processing and presentation of exogenous antigen | 1.08e-06 | 0.000715 | 51.06 |
| purple | GO:0010543 | regulation of platelet activation | 1.41e-06 | 0.000768 | 47.87 |
| purple | GO:1903555 | regulation of tumor necrosis factor superfamily cytokine production | 1.5e-06 | 0.000788 | 16.77 |
| purple | GO:0042590 | antigen processing and presentation of exogenous peptide antigen via | 1.4e-06 | 0.000789 | 127.66 |
| purple | GO:0048518 | positive regulation of biological process | 1.39e-06 | 0.000814 | 2.17 |
| purple | GO:0032101 | regulation of response to external stimulus | 1.38e-06 | 0.000842 | 6.09 |
| purple | GO:0032680 | regulation of tumor necrosis factor production | 1.37e-06 | 0.0008730000000000001 | 17.02 |
| purple | GO:0072376 | protein activation cascade | 1.83e-06 | 0.0009289999999999999 | 24.55 |
| pink | GO:0043005 | neuron projection | 2.62e-10 | 4.96e-07 | 3.84 |
| pink | GO:0097458 | neuron part | 8.46e-08 | 8.03e-05 | 2.83 |
| pink | GO:0120025 | plasma membrane bounded cell projection | 1.99e-07 | 0.000126 | 2.78 |
| pink | GO:0042995 | cell projection | 1.29e-06 | 0.000612 | 2.5 |
